# A Chemically Defined Synthetic Cell Capable Of Growth And Replication

**DOI:** 10.64898/2026.07.01.735724

**Authors:** Nathaniel J. Gaut, Christopher Deich, Brock Cash, Tanner Hoog, Aaron E. Engelhart, Katarzyna P. Adamala

## Abstract

Cells are the fundamental unit of life. Yet there is no natural cell for which all its life-essential functions are understood. Here we demonstrate a complete cell cycle for a synthetic cell undergoing selection, with genome replication, growth, resource acquisition via feeding, and genetically encoded division. The cell is encoded via a 90kb genome that includes functions needed for resource uptake, transcription, translation, growth, genome replication, and division. The resulting synthetic cell is sufficiently encouraging to support routinization of synthetic cell engineering workflows, and will ultimately underlie diverse applications across all of biotechnology.

## Introduction

One of the most ambitious and fascinating goals of bioengineering is to build a biochemical system that could cross the threshold from chemistry to life. ^1–3^.

Significant progress has been made towards construction of a minimal living synthetic cell^4,5^, including approaches based on minimizing natural living organism^6^, and efforts starting with non-living components^7^. Yet, no synthetic cell created from chemically-defined non-living components, has been shown to implement most life essential functions, be fully controllable, and described in full operational detail.^8^

Here, we present a **synthetic minimal cell with fully defined 90kbp genome, undergoing multiple generations of cell cycle, selection and reproduction**. The resulting gives insights into the minimum components necessary for life^9^. Such a cell may also ultimately inspire a chassis that can be adapted for a variety of purposes^10^, framing a foundation for fully artificial organisms.^11,12^

Our work addresses a key challenge in constructing minimal cells exhibiting life-like properties: coupling of growth and division to gene expression. The synthetic cell presented here contains a chemically defined cell-free translation system encapsulated in liposomes. We then combine several previously demonstrated technologies, including genome replication, with new demonstrations of genetically encoded growth and division, selection, and competition for resources.

To start, the **central dogma** of biology can be reconstituted using two major methods: TxTl (translation system based on whole cell extracts^13,14^) and PURE (Protein Synthesis using Recombinant Elements^15^). Unlike TxTl, PURE is a defined reaction mixture, with all the components and their concentrations known. Thus, we used PURE, except for early experiments establishing the feeding mechanism (TxTl is used on **Figure 2** and SI figures related to **Figure 2**, PURE is used in the rest of the work).

**Figure 1.**
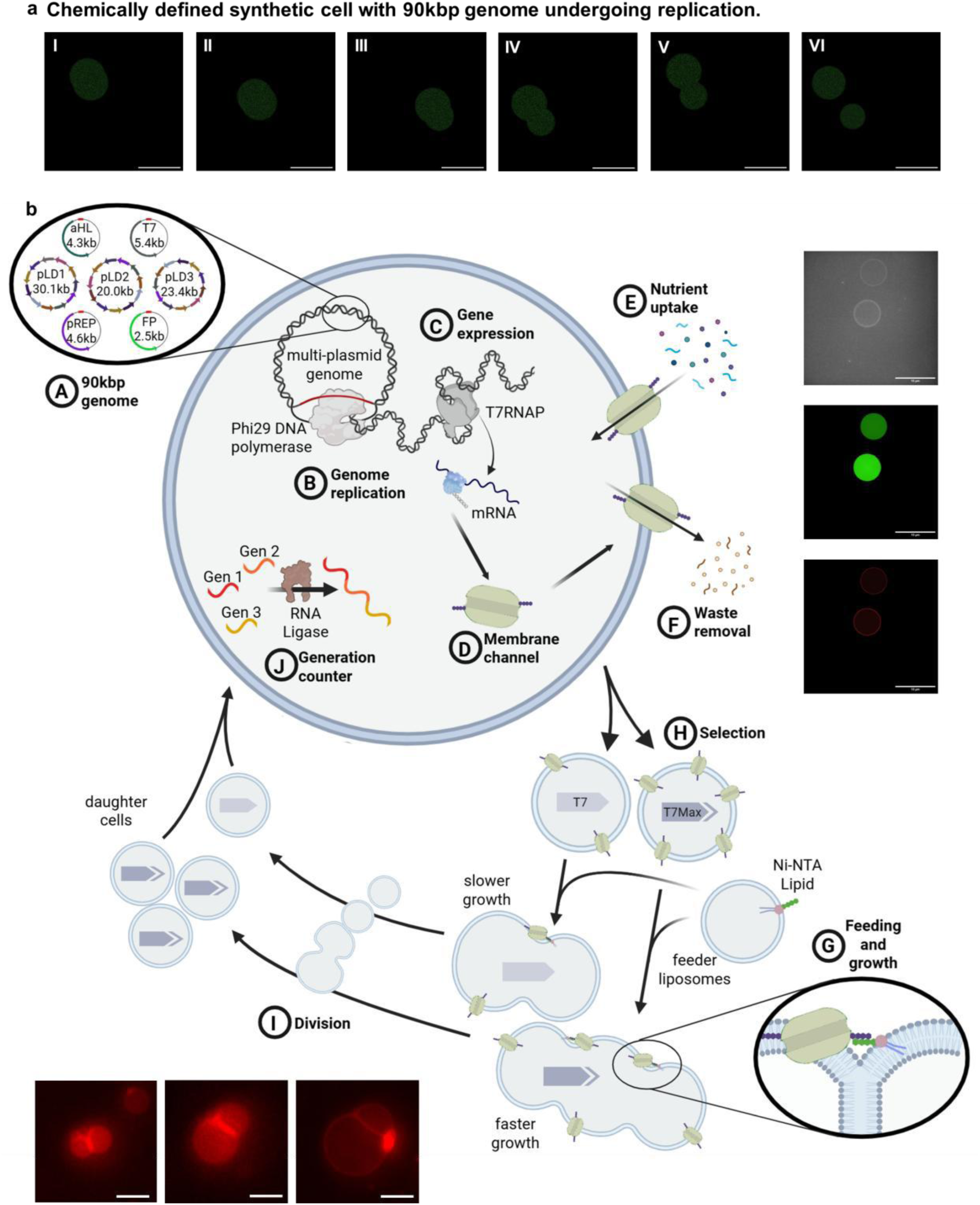
Cell cycle of synthetic cells with 90kbp genome, undergoing selection replication. The cells are assembled from individually purified natural components. **a**: Fluorescent microscopy of synthetic cell undergoing division. The cytoplasm of synthetic cell is expressing GFP. Scale bar is 10µm. Additional imaging is on **Figure 6**. **b**: The synthetic cells contain a multipartite DNA genome (**A**). The cell cycle comprises genome replication (**B**), protein expression (**C**), coupling translation of the membrane protein αHL (**D**) with nutrient uptake (**E**), waste removal (**F**), and genetically encoded feeding and growth (**G**). The feeding causes growth of the cell membrane (utilizing lipids from feeder liposomes) and replenishment of the cellular machinery (from the lumen of feeder liposomes). The feeding is possible through interaction of αHL expressed inside synthetic cells with the tag on the incoming feeder liposome. Thus, we demonstrate coupling of protein expression (αHL expression, which enables feeding) with growth of synthetic cells. The schematic insert on this figure illustrates the beginning of the fusion of the two membranes: synthetic cell membrane with feeder liposome membrane, as a result of Ni-NTA lipid tag interaction with αHL protein. Feeding and growth resulting from expression of protein inside the cell also enables competition between stronger and weaker feeders, and stronger feeders produce more offspring, enabling selection (**H**). The last step of the cell cycle is division of the compartment, either mechanical or genetically encoded division(**I**). We also developed a generation counter, to monitor rounds of feeding and division (**J**). The microscopy images on this figure show synthetic cell after 5 generations of growth and mechanical division using brightfield, GFP to visualize lumen and red dye for membrane (top right panel), and red membrane dye to visualize genetically encoded division (bottom left panel). The scale bar is 5µm.

**Figure 2.**
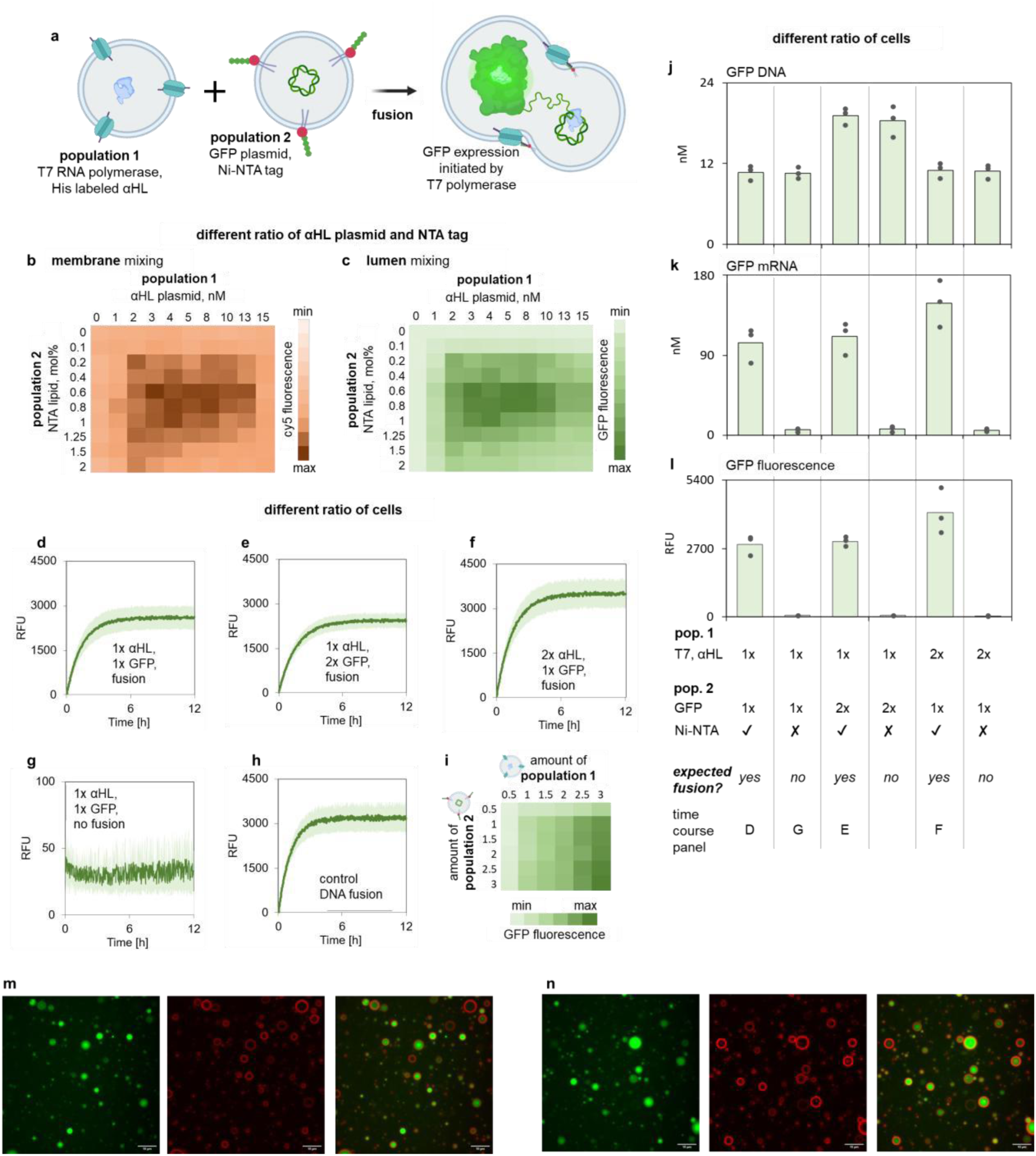
Genetically encoded feeding and growth of synthetic cells. **a:** Two populations of synthetic cells are mixed. Population 1 contains T7 RNA polymerase and TxTl machinery expressing αHL. Population 2 contains the plasmid encoding GFP under the control of a T7 promoter and Ni-NTA lipids in the membrane. The His tag on the αHL (in population 1) binds to the Ni-NTA lipids (in population 2), inducing membrane fusion. After lumens mixed, the T7 RNA polymerase from population 1 enables transcription and expression of GFP from a plasmid in population 2. **Different ratio of αHL plasmid and NTA tag:** **b** and **c**: The fusion efficiency depends on both the amount of the αHL plasmid in population 1, and the concentration of the Ni-NTA lipids in population 2. Fusion efficiency is measured as membrane mixing represented by FRET intensity between Cy5 and Cy7 (**b**, more Cy5 fluorescence correlates with more fusion), and lumen mixing represented by GFP expression (**c**, more GFP fluorescence correlates with more fusion). Individual data points and statistical analysis for GFP fluorescence is on **Figure S6**, for Cy5 FRET fluorescence on **Figure S7**. After fusion, selected samples were purified on a size exclusion column to measure content leakage, data on **Figure S3**. A calibration curve for all FRET membrane mixing measurements is on **Figure S76**. Data for the stability of the liposomes with membranes containing a different percent of Ni-NTA lipids are on **Figure S8** and the data for the stability of liposomes with an increasing amount of αHL are on **Figure S9**. **Different ratio of cells:** **d** - **h**: Example of individual time courses of GFP expression in fusion experiments. **d** - **f**: The amount of either population 1 containing αHL or population 2 containing GFP plasmid was varied. 1x is 1mM total lipid concentration and 2x is 2mM total lipid concentration. **g**: “No fusion” conditions indicate a lack of Ni-NTA lipids on population 2. **h**: The positive control uses DNA fusion tags instead of the αHL and Ni-NTA pair. Shaded area indicates SEM, n=3 independently prepared and processed liposome samples. The Y axis on panels in this section (except negative control **g**) is the same, to allow for quick visual comparison of the data. **i**: Fusion induced GFP expression depends on the ratio of population 1 containing αHL vs. population 2 containing GFP. All values are averaged with n=3 independently prepared and processed liposome samples. Individual GFP fluorescence data points and statistical analysis are on **Figure S12**. **j** and **k**: RT qPCR and qPCR measurements of the GFP gene, **l**: GFP fluorescence from the same fusion samples as **j** and **k** with all measurements taken after a 12-hour incubation. The graphical legend below panels **j**, **k**, and **l** marks the ratio of population 1 (αHL, T7 RNAP) to population 2 (GFP plasmid, Ni-NTA) and the presence or absence of a fusion event (absence of fusion means the absence of Ni-NTA lipids in population 2). Box and whiskers data points on panels **j** and **k** are prepared with n=3 independently prepared and processed liposome samples. Individual data points and statistical analysis are on **Figure S13**. Dots on top of bars on panel **L** indicate all individual data points, the bars are the average value. **m**: Two examples of microscopy images of liposomes after fusion. The green channel shows GFP expression, the red channel shows Rhodamine membrane dye, and the third panel in each grouping is composite of the red and green channel. The liposomes are from experiments under the same fusion conditions as data on panel **d**, with images collected after 12h of incubation. All protein expression data on this figure were collected using whole cell lysate (TxTl).

**DNA replication** in liposomes and extracts has been achieved with the complex bacterial replisome^16,17^, or by a more streamlined method of TTcDR (Transcription-Translation coupled DNA Replication)^18^. The latter is based on the replicase from the *Bacillus subtilis* phage Phi29 ^19^. We used the Phi29 replicase to carry out rolling circle amplification of the plasmids constituting the genome of our synthetic cell^18,20^.

**Feeding**, supply or acquisition of material and energy, is essential for life. Liposomes can be grown by fusion, using peptide^21^ and nucleic acid tags^22–24^, lipid composition ^25^, UV,^26^ or membrane tension^27^. Synthetic cells can also grow via synthesis of lipids inside the liposome^28–31^, coupled with DNA replication^32^, or with lipids from outside the liposome^33,34^. **We implemented feeding by fusion of nanovesicles directed by peptide tags.** The targeting of fusion is directed by a protein encoded and expressed by the synthetic cell; growth is thus regulated from the genome, **coupling metabolic activity to growth and fitness**.

**Selection** is another hallmark of life. Artificial selection has been demonstrated in liposomes ^35^, or in water-oil emulsions ^36^. Here, **we show coupling of the selection trait to synthetic cell growth, enabling selection via differences in the synthetic cell genome**. A mutation that results in production of more protein responsible for growth, enables cells with that mutation to grow bigger. More growth results in more daughter cells, thus spreading the mutation through the population.

Prior analysis has speculated that a minimal **genome** for a living cell could be as small as 113kbp.^37^ The synthetic cells presented here encode a 90kbp genome.

**Self-replication** is another goal of synthetic cell engineering^38,39^. Several mechanisms for achieving growth and division have been proposed^40–42^. Recent progress demonstrated reconstitution of protein cytoskeleton^43–46^, polymer hybrid vesicles^47^, and membrane deformation based on nucleic acids^48,49^, DNA-peptide structures^50^, and other chemistries^51,52^. Here, utilizing mechanisms based on membrane fission induced by surface protein crowding^53,54^, **we have achieved genetically encoded synthetic cell division** in the absence of cytoskeleton.

Taken together, we demonstrate **the first minimal cell with a cell cycle, genetically encoded growth and division, all coupled to selection and competition**. The synthetic cells undergo gene expression via PURE, genome replication via Phi29, and genetically encoded processes of membrane growth, feeding, and division. The introduction of mutation increasing expression of the protein needed for feeding results in faster growth and more daughter cells. This allows for the spread of advantageous mutation through the population, demonstrating selection. (**Figure 1**)

We demonstrated **five generations of the overall cell cycle** including DNA replication, feeding, and growth (**Figure 3**), first tested with mechanical division - to ensure high efficiency of replication. Utilizing this cell cycle, we introduced an advantageous mutation, increasing expression of the protein responsible for growth. **Over several generations, the cells containing the beneficial mutation produced more daughter cells,** demonstrating the spread of the advantageous mutation in the population, and thus **an example of selection** (**Figure 4**). When cells compete for food in a resource-limited environment, faster growing cells produce more offspring than the slower growers as food becomes scarcer – illustrating **competition for resources driving faster selection**. (**Figure 5**) Then we engineered a **genetically encoded division mechanism**, enabling direct link of the genotype of synthetic cell to the number of offspring. (**Figure 6**) Finally, coupling genetically encoded growth and division resulted in **synthetic cells that can feed, grow, and divide.**

**Figure 3.**
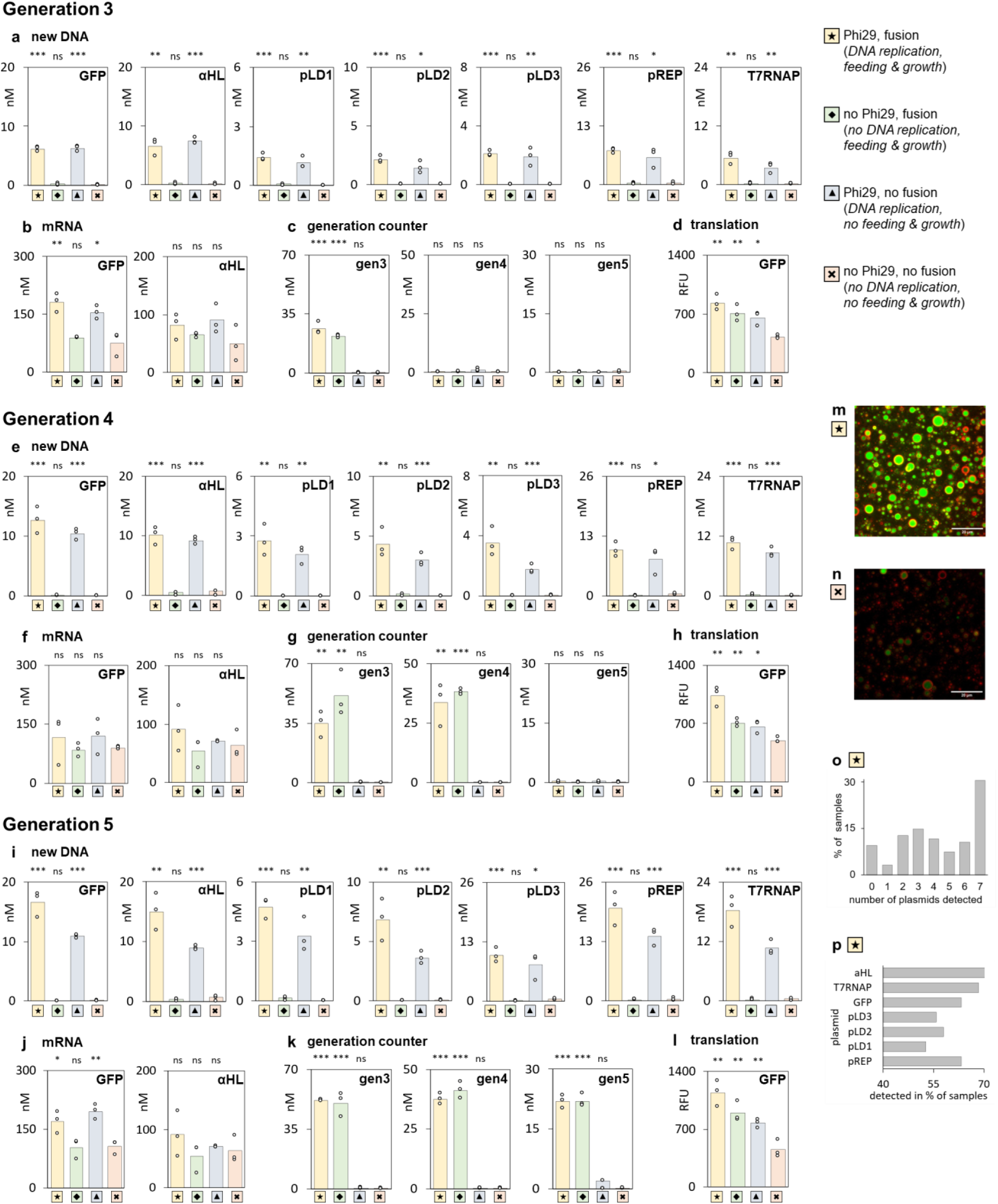
Five generations of cell cycle with 90kbp genome. Synthetic cells were undergoing five generations of cell cycle, and the samples were analyzed after three, four, and five generations. The data shown on this figure represents experiments with mechanical division after each generation. All experiments were performed in four separate variants: the complete cell cycle with Phi29 genome replication and with feeder liposomes (symbol 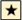); without Phi29 genome replication, but with feeding (symbol 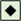; with Phi29 for genome replication, but without feeding (symbol 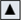); and without either Phi29 genome replication or feeding (symbol 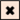). **a**, **e** and **i**: qPCR quantification of new DNA, one gene from each plasmid, indicating Phi29 genome replication. The samples were treated with DpnI, removing the original plasmid DNA, so only newly synthesized DNA was detectable. All individual genes on every plasmid of the genome were similarly quantified after generation 5, data on **Figure S28**. **b**, **f** and **j**: RT qPCR quantification of the mRNA abundance. **d**, **h** and **l**: GFP fluorescence measured for all samples at the end of the cell cycle for the indicated generation. Dots on top of the bars indicate all the individual data points, the bars are the average value. Flow cytometry analysis of GFP samples is on **Figure S30**. **c**, **g** and **k**: quantification of the generation counter, detecting the counter product of the indicated number of generations. The samples were processed with either the primer for generation three (labeled gen3, detecting three, four, or five generations), the primer for generation four (labeled gen4, detecting three or four generations), or the primer for generation five (labeled gen5, detecting only the results of five generations). **m** and **n**: Fluorescence microscopy of synthetic cell samples, imaged after five generations. **m**: complete cell cycle, including both Phi29 genome replication and feeder liposomes, **n**: no genome replication and no feeding. Scale bar is 20µm. The green channel is GFP fluorescence, and the red channel is rhodamine membrane dye. More images are on **Figure S31**. The feeder vesicles are not visible on microscopy images because of lack of membrane or lumen fluorescent label. **o** and **p**: Abundance of each plasmid after 5 generations of cell cycle with growth and replication (symbol 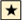) measured in individual cells. Data from single cell amplification and qPCR analysis of abundance of each of the individual plasmids (see **Table S6** for the list of plasmids) in a synthetic cell genome. **o**: Number of cells containing given number of plasmids from the 7-plasmid genome. **p**: Percent of single cell samples containing each plasmid. Individual data points for each plasmid are on **Figure S32**. On all panels, the graphs represent 3 independently prepared and processed liposome samples, individual data points are on **Figure S34**. Statistical significance analysis showing P values for data demonstrated on this figure is in **Table S9**. Cell leakage analysis after 5 generations is on **Figure S35**. The cartoon cells are a visual guide only, not an accurate description of content of each cell (each cell contains multi-plasmid genome, multiple copies of ribosomes, polymerases, and generation counters). All Y axes on qPCR panels are in nM. The panels indicating the same measurement for three different generations (for example new GFP DNA on panels **a**, **e**, and **i**) have the same Y axis scale, to enable direct comparison. All protein expression data on this figure were collected using PURE system.

**Figure 4.**
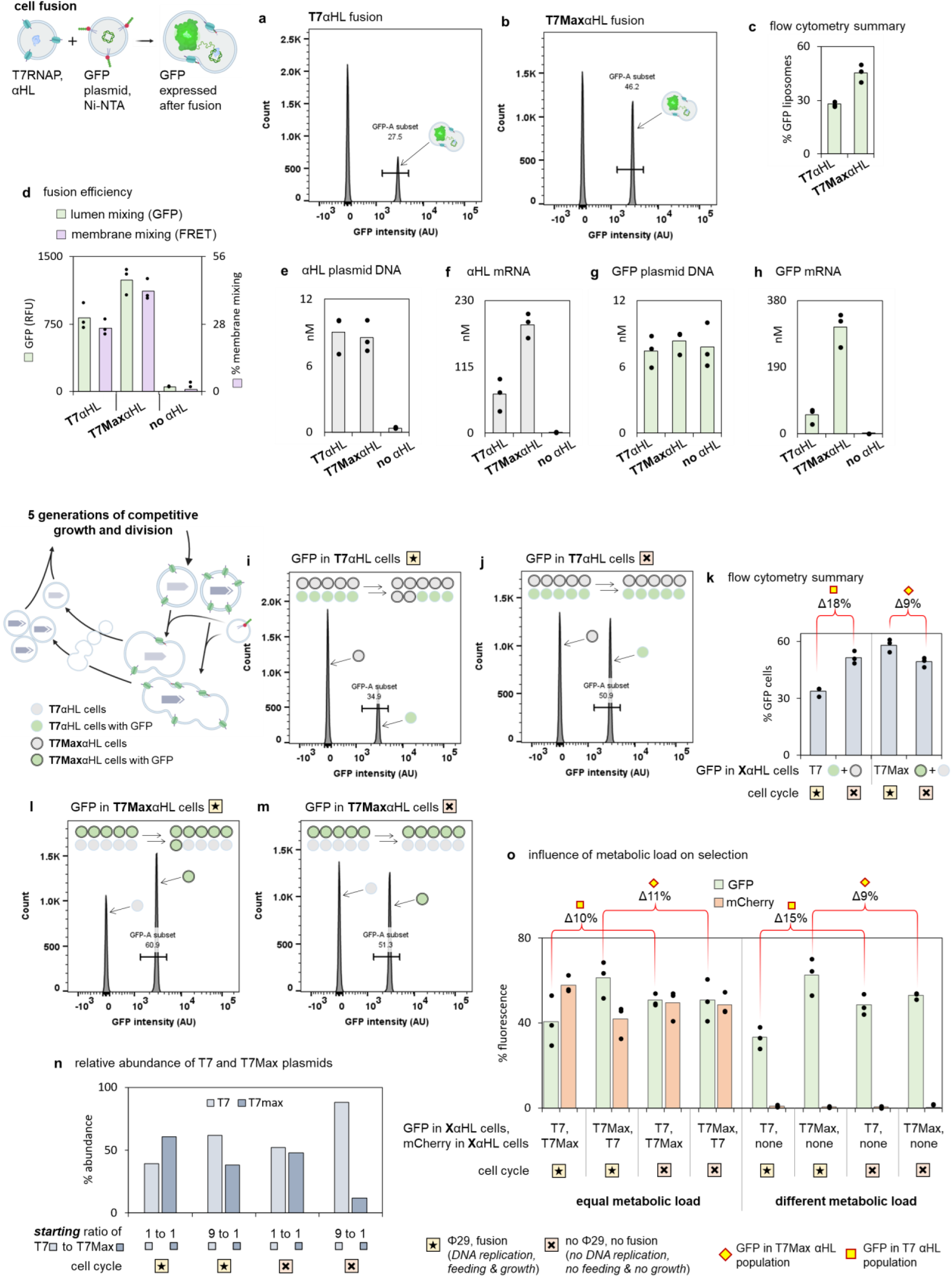
Selection in populations of synthetic cells. **a** - **h**: **Fusion efficiency controlled by strength of αHL expression**. Fusion between two populations of synthetic cells, according to the same setup as shown on **Figure 2**, with lumen fusion indicated by GFP expression and membrane fusion indicated by Cy5 signal. **a** - **c**: Flow cytometry analysis of samples prepared with either T7 αHL (panel **a**) or T7Max αHL (panel **b**), after fusion incubation. Full flow cytometry data are on **Figure S49**. **c**: summary of the flow cytometry results, showing percentage of liposomes expressing GFP in samples with “weaker” T7 αHL and with “stronger” T7Max αHL. All plasmids other than αHL were under the T7 promoter, the T7Max was only used in αHL plasmid. “T7Max cells” indicate cells with αHL under T7Max, with all other plasmids in that cell under regular T7. **d** - **h**: characterization of liposome fusion with T7 αHL, T7Max αHL, and negative control without αHL. **d**: GFP signal (green bars) indicating lumen mixing, and % membrane mixing (gray bars), measured after 12h incubation. **e** and **g**: abundance of plasmid DNA (**E** for αHL plasmid and **G** for GFP plasmid) in the same fusion experiments as shown on panel **d**. **f** and **h**: abundance of mRNA (**F** for αHL gene and **H** for GFP gene) in the same fusion experiments as shown on panel **d**. **i** - **o**: **Selection over 5 generations of cell cycle.** All samples were prepared as described in the cell cycle experiments shown on **Figure 3**. **I** - **m**: Flow cytometry analysis after 5 generations of cell cycle with two competing populations in each sample. Four types of samples were prepared: cells with T7 αHL plasmid (symbol 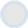), cells with T7 αHL plasmid and GFP plasmid (symbol 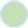), cells with T7Max αHL plasmid (symbol 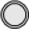), and cells with T7Max αHL plasmid and GFP plasmid (symbol 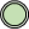). Two types of cells were mixed, at equal ratio, for each experiment. In each case, T7 and T7Max cells were mixed, with one of the populations containing GFP reporter. After either 5 generations of complete cell cycle (symbol 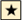), or incubation without DNA replication and without feeding and division (symbol 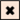) samples were analyzed using flow cytometry. Experiments with GFP reporter in T7 αHL cells: after DNA replication and division on panel **i**, and without DNA replication and division on panel **j**. Experiments with GFP reporter in T7Max αHL cells: after DNA replication and division on panel **l**, and without DNA replication and division on panel **m**. Full flow cytometry data are on **Figure S50**. **k**: summary of the flow cytometry data, showing percentage of liposomes containing GFP in each of the 4 selection samples from panels **i**, **j**, **l** and **m**. **n**: Relative abundance of T7 and T7Max promoter sequences in samples after 5 rounds of complete cell cycle. Two types of synthetic cells were prepared for this experiment: cells with T7 αHL plasmid and cells with T7Max αHL plasmid. For each experiment, those two initial populations were mixed at ratio indicated in panel legend (equal 1 to 1 ratios, or samples where T7 population was 90% of the total abundance with the T7Max being 10%). All resulting samples had identical total lipid concentration, but different ratios of T7 to T7Max populations. After either 5 generations of full cell cycle (symbol 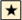), or incubation without DNA replication and without feeding and division (symbol 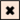) the abundance of T7 and T7Max sequences was quantified via Illumina sequencing (Methods section 16, and **Figure S53** and **Figure S54**). **o**: selection experiments with all cells carrying a fluorescent protein of similar size (normalized metabolic load). Experiments performed similarly to those on panels **i**-**m**, but this time both T7 and T7Max cells carried fluorescent protein (GFP or mCherry, as indicated in the legend under graph on panel **o**). After 5 generations, fluorescence of both GFP and mCherry was measured, and normalized to calibration curves indicating what percent of each population contains this fluorescent protein (see **Figure S55** for raw fluorescence data and **Figure S56** for calibration curves, **Table S10** for statistical analysis). The red bracket on panels **k** and **o** indicates the difference between the percent of population containing T7 and T7Max (all from starting 1 to 1 ratio). Experiments under similar set of conditions are indicated with yellow symbol: 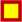 samples with GFP in T7 αHL population, and 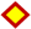 samples with GFP in T7Max αHL population. Data on all panels represent n=3 of independently prepared and processed liposome samples. Dots on top of bars indicate individual data points, the bars are the average value. Statistical analysis of data presented on this figure is in **Table S10.** All protein expression data on this figure were collected using PURE system.

**Figure 5.**
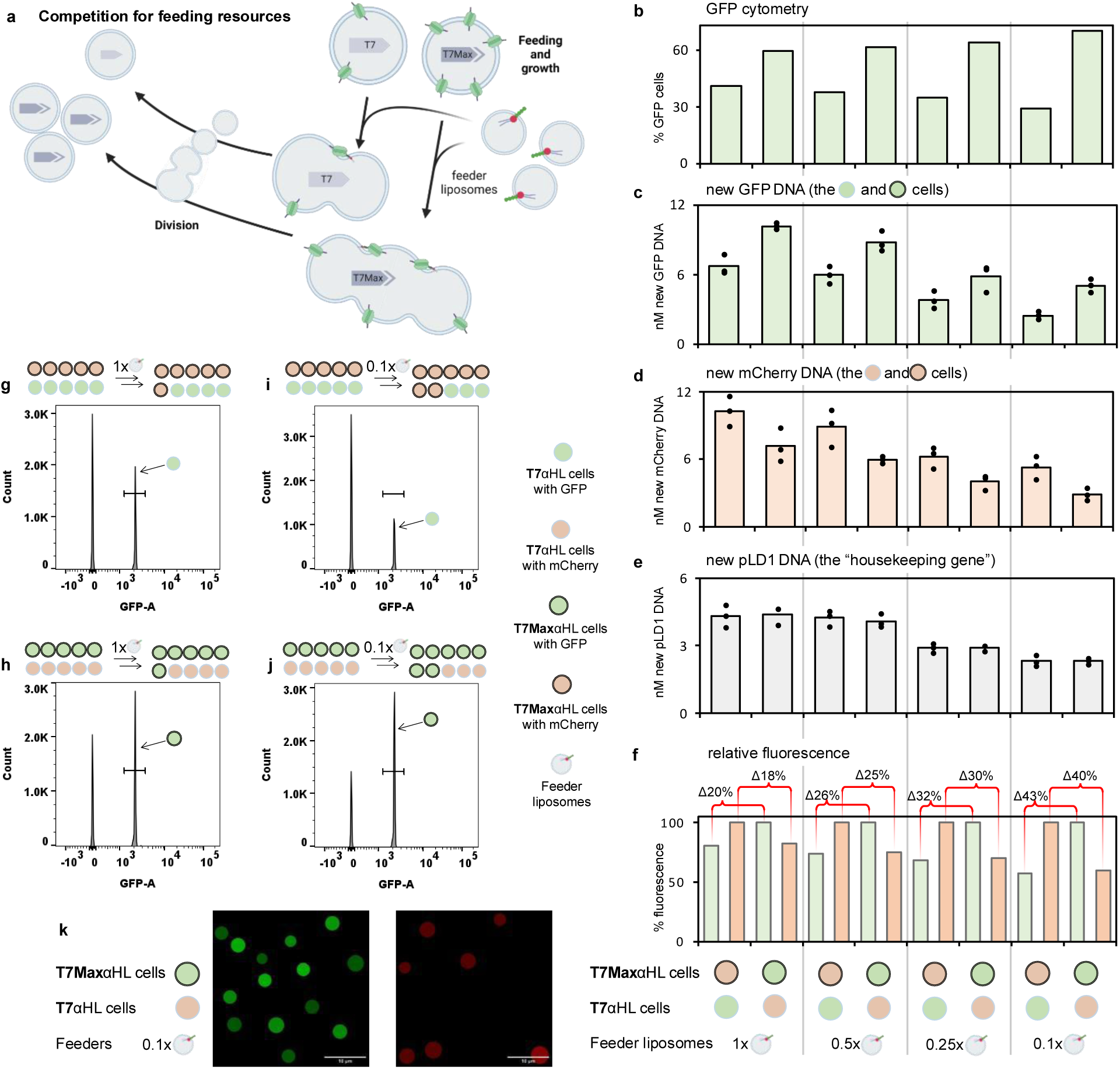
Competition for resources over multiple generations. **a**: Two populations of synthetic cells, with αHL under T7 (weak) or T7Max (strong) promoter, go through 5 cycles of growth and division. The strong growing cells (T7Max) fuse with more feeder liposomes than the weak growers (T7), resulting in more offspring carrying the strong (T7Max) gene after each round of division. If the amount of feeder liposomes is restricted, the cells compete for the limited feeding resource and strong growers win. **b** – **f**: Two populations of synthetic cells were mixed in each competition experiment. One population expressed green marker (GFP) and the other population expressed red marker (mCherry). Each experiment was performed in two variants: either mCherry was in the T7Max αHL population (symbol 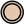) and GFP was in the T7 αHL population (symbol 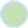), or the reverse with the GFP in the T7Max αHL population (symbol 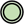) and mCherry in the T7 αHL population (symbol 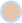). Both variants of each experiment had five generations of growth and feeding, with varying amount of feeder liposomes. The 1x feeder liposomes means the same conditions as cell cycle experiments on **figure 3**. The 0.5x, 0.25x and 0.1x feeder liposomes refer to proportionally less feeder liposomes than in the original 1x conditions. After 5 generations of growth and division, samples were analyzed by flow cytometry and qPCR. Changes in plasmid concentration over each generation were also quantified, **Figure S58**. Summary of changes between feeding regimes is on **Figure S57**. **b**: Flow cytometry analysis, quantifying what percentage of the total population are cells with the GFP marker. Full flow cytometry traces are on **Figure S61**. **c** – **e**: qPCR quantifying amount of newly synthesized DNA. **c**: GFP gene, **d**: mCherry gene, **e**: pLD1 gene (control “housekeeping” plasmid, identical in both T7 and T7Max populations). **f**: Relative fluorescence of mCherry and GFP, after 5 generations of growth and division. Absolute fluorescence values, individual data points and statistical analysis are on **Figure S62**. **g** – **j**: Example flow cytometry traces for individual experiments. **G**: mCherry in the T7Max αHL population, 1x feeder liposomes, **h**: GFP in the T7Max αHL population, 1x feeder liposomes, i: mCherry in the T7Max αHL population, 0.1x feeder liposomes, j: GFP in the T7Max αHL population, 0.1x feeder liposomes. **k**: Microscopy images showing cells under conditions of 1x feeding, with GFP in the T7Max αHL population (symbol 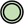) and mCherry in the T7 αHL population (symbol 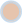). Scale bar is 10µm. The feeder vesicles are not visible on microscopy images because they lack membrane or lumen fluorescent label. Data on all panels represent n=3 of independently prepared and processed liposome samples. Dots on top of bars indicate individual data points, the bars are the average value. Statistical analysis for data on panels **b** – **f** is in **Table S11**. All protein expression data on this figure were collected using PURE system (Methods section 2).

**Figure 6.**
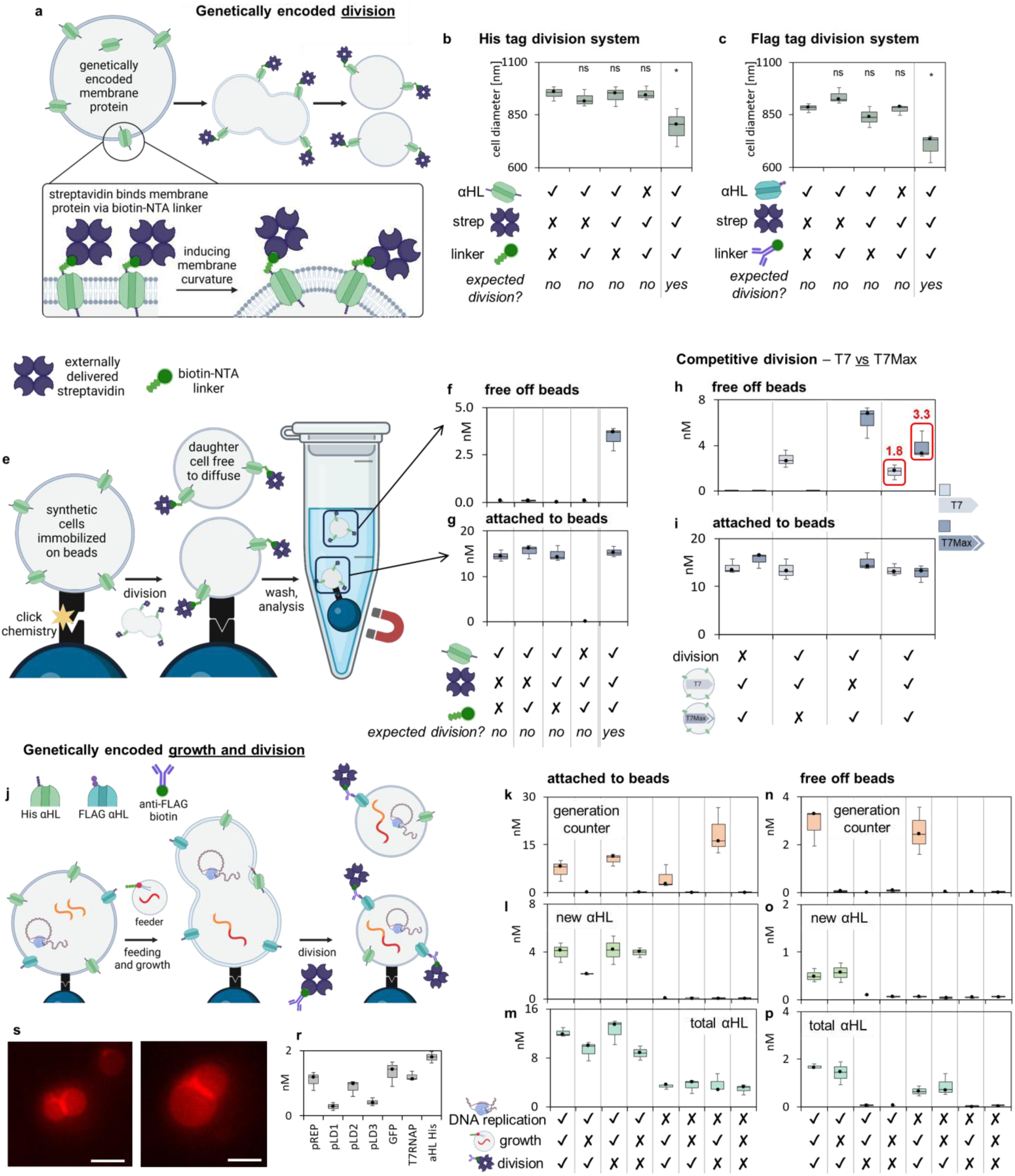
Genetically encoded replication of synthetic cells. **Genetically encoded synthetic cell division.** **a**: synthetic cells express tagged membrane protein αHL; streptavidin and linker are added on the outside of the cells, binding to the αHL tag on the surface of the cells. The streptavidin induces curvature in the membrane, leading to cell division. Two systems were tested: αHL His tagged with Ni-NTA linker to bind streptavidin, and αHL FLAG tagged with biotin conjugated FLAG antibody to bind streptavidin. **b**: Synthetic cell size after division with the His tag system, under conditions excluding one of the essential division components (αHL, streptavidin, or the Ni-NTA linker), and with all division elements present. Individual data points are in **Figure S64**. **c**: Synthetic cell size after division, using FLAG tag system, under conditions excluding one of the essential division components (αHL, streptavidin, or the biotin FLAG antibody linker), and with all division elements present. Individual data points are in **Figure S65**. Flow cytometry data for division experiments are on **Figure S63**. **e-i**: Division of immobilized cells, and competition between two populations of cells. **e**: Synthetic cells are immobilized on magnetic beads. Streptavidin and Ni-NTA linker are added to the outside of the cells, as described on panel **a**. If division occurs, the daughter cells are free to separate from the beads. After the experiment, free synthetic cells are collected separately from the synthetic cells that remained on the beads. **f** and **g**: quantification of αHL DNA in synthetic cells freed from the beads (the “daughter cells”, panel **f**) and cells that remained attached to the beads (the “parent” cells, panel **g**), in experiments excluding one of the essential division components (αHL, streptavidin or the Ni-NTA linker), and with all division elements present. Individual data points are in **Figure S66**. **h** and **i**: Competitive division. Synthetic cells containing αHL under “slow” promoter T7 and αHL under “fast” promoter T7Max are dividing in the same reaction. Samples were analyzed for the presence of unique DNA marker (different for T7 and T7Max cells). Label “division” indicates either presence of streptavidin and Ni-NTA linker (division label “yes”), or absence of both streptavidin and linker (labeled “no” division). In each experiment, the sample contained either a mix of both synthetic cells containing T7 and T7Max αHL in equal amounts, or only one of the populations (only T7 or only T7Max). After incubation and division, unique DNA marker was quantified, identifying T7 and T7Max populations in “daughter” cells free off the beads (panel **h**) and in cells that remained attached to beads (panel **i**). Individual data points and statistical analysis are in **Figure S67**. **Genetically encoded growth and division** **j**: Synthetic cells were immobilized on beads. The cells expressed two types of αHL: His tag labeled, and FLAG tag labeled. The synthetic cells contained entire 90kbp genome (as shown on **Figure 3**), and growth was induced by addition of feeding liposomes with single generation counter marker. Following growth, division was induced by attaching streptavidin to the surface of synthetic cells via biotin conjugated FLAG antibody. After division, the “daughter” cells separated from the beads and the “parent” cells still attached to beads were collected and analyzed separately. Experiments were performed under conditions with or without DNA replication (i.e., +/- Phi29), with or without growth (feeder liposomes +/- Ni-NTA fusion tag), and with or without division (+/- biotin FLAG antibody). **k** and **n**: Generation counter oligo detection, indicating lumen fusion with feeder liposomes, measured in cells that remained attached to beads (panel **k**) and in cells free off the beads (panel **n**). **l** and **o**: quantification of newly synthesized αHL DNA, in cells attached to beads (panel **l**) and in cells free off the beads (panel **o**). **m** and **p**: measurement of total αHL DNA (not just newly synthesized), in cells attached to beads (panel **m**) and in cells free off the beads (panel **p**). **r**: qPCR detection of each plasmid from the synthetic cell genome in the “daughter” cell fraction free off the beads, in samples that did undergo feeding and division and DNA replication via Phi29. Individual data points for panels **k**-**p** are in **Figure S68**, statistical analysis is in **Table S12**, and individual data points for panel **r** on **Figure S69**. **s**: Fluorescent microscopy of synthetic cells after a cycle of genetically encoded growth and division. Scale bar is 2µm. Additional imaging is on **Figure S73**. All protein expression data on this figure were collected using PURE system (Methods section 2).

## Results and Discussion

Synthetic cells in this work are liposomes containing DNA genome encoded across seven plasmids, and *in vitro* protein expression system. Unless otherwise stated, all proteins are expressed inside the synthetic cell.

The synthetic cells presented herein have a defined chemical composition at the time of formation, with known concentration of components making up the cell, starting from the PURE system).

Our synthetic cells divide without cytoskeleton. Division is first achieved by mechanical extrusion through a membrane filter (**Figures 3, 4** and **5)**. Ultimately, we demonstrate genetically encoded division as a result of membrane protein interactions (**Figure 6)**. The summary of major findings is in **Table S1**.

### Genetically encoded liposome fusion leads to growth

Our synthetic cells have a very limited metabolism^55^ and cannot make ribosomes;^56^ complete metabolic independence will require a larger genome ^37^. To engineer a complete cell cycle with a smaller genome, we needed a method for feeding enzymes, small molecules, ribosomes, and lipids, to grow and sustain synthetic cells. We achieved that using small “feeder” liposomes as a source of both lipids for membrane growth, and other molecules. The small molecule nutrients were replenished both by feeder liposomes, and by membrane transport from external medium. We engineered a method of fusion of synthetic cells with feeder liposomes, based on expression of a protein inside the cell, thus making the **growth and feeding genetically encoded**.

To engineer genetically encoded liposome fusion, we first needed proof of principle: evidence that a small peptide tag on the membrane will enable fusion. We used a 6xHis tag embedded in an externally delivered trans-membrane peptide domain (Supplementary text section 1, “His tag mediated liposome fusion leads to growth”).

After establishing that 6xHis tags embedded in the membrane can facilitate fusion, we needed a way for the His tag made inside synthetic cells to be displayed on the surface.

A membrane pore α-hemolysin (αHL) is often used to exchange small molecules between the inside and outside of synthetic cells ^57–61^. The αHL pore expresses robustly in cell-free systems, inserting into the liposome and spanning the whole membrane ^63,64^. This enables presenting internally expressed peptide tags on the surface of synthetic cells.

### Modified αHL expressed inside synthetic cells facilitates fusion

We inserted a 6x-His tag into a trans-membrane loop of αHL, before Asp129 (DNA sequences are in **Table S7**), a location consistent with recently reported functionalization of αHL to display other tags.^62^

To test liposome fusion with His-αHL, we prepared two populations of synthetic cells: population 1 contained T7 RNA polymerase (T7RNAP) and αHL plasmid, and population 2 contained GFP plasmid and Ni-NTA lipid tags. Both populations also contained cell-free translation. αHL expression results in His tag presented on the surface of population 1, and the Ni-NTA lipid tag from population 2 binds to this His tag, initiating fusion. Upon liposome fusion, T7RNAP from population 1 facilitates expression of GFP plasmid from population 2. Lumen mixing can be monitored via GFP expression, and membrane growth can be monitored via membrane FRET dye signal (**Figure 2a**).

After mixing equal amounts of population 1 (T7RNAP, αHL) with population 2 (GFP plasmid, Ni-NTA), we observed both GFP expression (**Figure 2c**) and increased FRET donor signal (**Figure 2b**), indicating membrane growth and lumen mixing. Fusion efficiency was dependent on both the amount of αHL and Ni-NTA. Increasing the concentrations of both resulted in increased fusion efficiency, up to 10nM αHL plasmid, and we observed decreased fusion efficiency above 1 mol% of Ni-NTA lipid (**Figure 2b** and **c**).

The decrease in fusion efficiency at higher Ni-NTA concentrations can be explained by decreased liposome stability (**Figure S1**). We speculate that the decrease in fusion efficiency at higher αHL concentrations was due, in part, to an increase in leakage of the liposome contents through more membrane pores (**Figure S3**).

### Fusion efficiency depends on amount of αHL

We investigated further improvement by mixing population 1 (T7RNAP, αHL) and population 2 (GFP plasmid, Ni-NTA) at different liposome ratios. The GFP expression increased with a higher amount of population 1, but not with higher amount of population 2 (**Figure 2i**), indicating that αHL liposomes carrying T7RNAP were the rate limiting reagent in the GFP production. This phenomenon was advantageous for the intended use of this fusion mechanism: in our synthetic cell cycle, cells make αHL while the Ni-NTA “feeder liposomes” will be used in excess. This might be compared to natural feeding: there is usually more prey (i.e., feeder liposomes) than predators (i.e., synthetic cells).

We compared the time course of GFP expression at different αHL and Ni-NTA liposome ratios (**Figure 2d-f**) to the time course of GFP production in a positive control fusion experiment facilitated by DNA tags (**Figure 2h**). The DNA tag system has been previously used to facilitate synthetic cell fusion^34^. At identical total liposome concentrations, the DNA tag-induced fusion had similar efficiency as the αHL – Ni-NTA fusion. The negative “no fusion” control were samples containing no Ni-NTA tags on population 2, resulting in no measurable GFP expression (**Figure 2g**).

To further characterize the fusion system, we quantified the DNA and mRNA post-fusion. We prepared population 1 containing T7RNAP and αHL at 3nM plasmid, and population 2 containing GFP plasmid and Ni-NTA at 0.6mol%. We performed fusion experiments mixing the two populations, using either 1x (1mM total lipid concentration) or 2x (2mM total lipid concentration) of each population. After fusion and incubation, we measured GFP fluorescence, and quantified amount of GFP mRNA and DNA. As expected, experiments with 2x of population 2 (carrying GFP plasmid) had twice as much GFP DNA as samples with 1x population 2 (**Figure 2j**). Transcription of GFP occurred in all samples that underwent fusion (**Figure 2k**), confirming the fluorescence data showing GFP expression in samples after fusion (**Figure 2l**).

Liposome fusion is possible only if both 1) αHL is expressed in one population, and 2) Ni-NTA lipids are present on the other liposome population (**Figure S5)**. We have visualized the liposomes after fusion, with lumen GFP expression and red membrane dye (**Figure 2m**).

### Summary

Overall, our experiments demonstrate that **His tagged αHL paired with Ni-NTA lipids induce liposome fusion, which leads to mixing of the lumen of synthetic cells**. The fusion efficiency depends on the expression of His tagged αHL, and on the amount of Ni-NTA lipids in the membrane. Because the αHL is expressed inside synthetic cells, this is a fusion system that depends on protein encoded in the genome of the cells.

For the cell cycle experiments described below, we used 3nM αHL plasmid in starting synthetic cell population, and 0.6mol% Ni-NTA in the membrane of feeder liposomes.

### Five generations of synthetic cell cycle

Cell cycle, rounds of feeding and growth, is one of the hallmarks of life. We combined genetically encoded feeding and growth, DNA replication, and mechanical division, into a synthetic cell cycle.

We built a synthetic minimal cell encapsulating a 90kbp genome, divided into eight plasmids (**Table S6**). The genome contained proteins needed for the translation system^65^, Phi29 DNA polymerase for genome replication, His tagged αHL membrane pore, T7RNAP gene for transcription, and GFP reporter gene.

Gene expression was achieved through a PURE *in vitro* translation system modified to accommodate the Phi29 polymerase reaction^20^ (Methods section 2). We optimized the external solution, called the synthetic cell media, to increase yield of translation (Supplementary text section 4, “Synthetic cell culture media”). The presence of an external solution containing all necessary small molecules prevents leakage from αHL pores from diluting the synthetic cell contents.

The feeder liposomes contain all the enzymes and small molecules needed for synthetic cell metabolism (RNA polymerase, Phi29, PURE system) but no DNA. The feeder liposome membranes are labeled with 0.6mol% Ni-NTA lipids, to enable fusion with the His tagged αHL synthetic cells. Feeders provide replenishment for all the components of synthetic cell metabolism, and provide lipids for growth of the cell membrane. Cells were first divided using mechanical extrusion, to provide high division yields (with lower yield genetically encoded division presented later in this paper).

### Cell cycle results in replication of DNA, transcription and translation in daughter cells

Each cell cycle starts with a 12-hour incubation of cells in the presence of feeder liposomes. After 12 hours, synthetic cells are mechanically divided (via extrusion), and mixed with a fresh population of feeders, starting a new generation. Feeder liposomes at each generation carry a generation counter oligo. After three, four, and five generations, we measured: GFP fluorescence, mRNA abundance for GFP and αHL genes, abundance of newly synthesized DNA for each plasmid of the 90kbp genome, and the generation counter analysis (**Figure 3**).

Newly synthesized DNA was detected in cells after each generation. To ensure that we only detect new DNA arising from Phi29 genome replication, samples were first digested with DpnI. We measured the amounts of all individual plasmids in the cell genome: GFP, αHL, pLD1, pLD2, pLD3, pREP and T7RNAP. New DNA was only detected in samples where Phi29 replicase was present (**Figure 3a, e and i**).

In each generation cells also produced mRNA (**Figure 3b, f, j**) and GFP (**Figure 3d, h, l**; Supporting Text section **3**).

### The same “lineage” of synthetic cells feeds in every generation

We developed a generation counter to track feeding events and provide direct evidence that the same “parent” synthetic cell fused with at least one feeder liposome in each generation (Supplementary text section 2, “The generation counter”). A starting population of cells was created with all five-generation counter “top” strands, and each population of feeder liposomes contained one “bottom” strand for the appropriate generation. After each generation, we processed samples with a primer specific to the product of three, four, and five generations of the counter. For example, using the primer for three generations we detected generation counter in cells fed at generation 3, whereas no Gen3 product was detected in samples without feeding (**Figure 3c**). Similarly, analysis with the primer matching the product of four generations showed product in the fed samples after four and five generations, and no product after three generations or without feeding (**Figure 3g**). With the primer for the product of five generations (labeled Gen5 primer) we detected the expected product only in the samples with all five generations of feeding, and no product in earlier generations, or in samples without feeding (**Figure 3k**).

### All plasmids are inherited, and daughter cells are stable

After five generations, we performed single cell quantification of all plasmids from the 90kbp genome. 30% of all analyzed cells contained the complete genome - all seven plasmids (**Figure 3o** and **3p**).

Despite the lack of a cytoskeleton and a DNA segregation mechanism, a significant fraction of daughter cells underwent five cycles of growth and replication inheriting the complete multi-partite genome.

We visualized the liposomes after five generations of growth and division, with samples after complete cell cycle (**Figure 3m)** and samples without genome replication and without feeding (**Figure 3n)**.

As a control, we removed feeder liposomes by dialysis after each generation, to ensure that previous feeders are not available in the next generation. The removal of feeders after each generation did not significantly affect the amount of DNA and mRNA after five generations (**Figure S25**).

To confirm that liposomes remain stable, we measured content leakage after five generations. We detect no leakage of plasmid, mRNA or ribosomal RNA after five generations of growth and division, (**Figure S35**). The major components of the cell that are too big to pass through αHL channels remain inside the liposomes after five generations.

### Summary

The generation counter measurements demonstrate that **there are synthetic cells that fused with feeder liposome during all five rounds of feeding**. The DNA quantification confirmed that **new genetic material is synthesized during the cell cycle**, and **all plasmids of the 90kbp genome are replicated**. A summary of trends in mRNA, DNA, and generation counter abundance is in **Table S4**.

### Selection

Selection and competition are hallmarks of biological life. Darwinian evolution is possible in synthetic cells^66^, however without coupling of genotype to growth and number of offspring, the potential for natural selection would be limited. Having engineered a genetically encoded growth mechanism, we recognized the potential for demonstrating an example of selection process coupling genotype to phenotype through multiple cycles of growth and division.

### More αHL leads to more growth and more daughter cells after division

In our system synthetic cell feeding depends on the amount of αHL in the membrane: more protein results in more fusion with feeder liposomes (**Figure 2**). To engineer selection through feeding and growth, we introduced a mutation increasing expression of αHL. The mutation was in the αHL gene promoter, using a stronger variant of T7 RNA promoter, called T7Max.^67^ This represents a mutation that is potentially beneficial to cells, if more αHL gene product results in more feeding, more growth, and in turn leads to more daughter cells after division.

To explore whether a stronger promoter might cause more efficient fusion, we used a mixed populations of synthetic cells and observed GFP expression (indicating lumen mixing) and FRET (indicating membrane mixing) (**Figure 2**). In each experiment, one population of cells contained αHL plasmid, T7RNAP, and translation system, while the other population was labeled with Ni-NTA lipids and carried GFP plasmid plus translation system, but lacked T7RNAP. After mixing each cell type at equal ratios, we analyzed samples using flow cytometry. In samples with αHL under the control of regular T7 promoter, 28% of liposomes showed GFP fluorescence (**Figure 4a** and **c**), while in samples with αHL under control of stronger T7Max, 45% of the liposomes were fluorescent (**Figure 4b** and **c**). Both GFP (indicating lumen mixing) and Cy5 signals (indicating membrane mixing) increased more in cells with αHL under T7Max promoter, indicating greater fusion efficiency (**Figure 4d**). To confirm the fluorescent data, we quantified the amount of DNA and mRNA for GFP and for αHL genes. The DNA concentration for both αHL and GFP was the same in all samples (**Figure 4e** and **g**). The αHL and GFP mRNA abundance was higher in samples with the stronger promoter (**Figure 4f** and **h**), which aligned with GFP fluorescence data, showing more mRNA in samples with αHL under T7Max and samples with higher fusion efficiency for stronger promoter.

Overall, at the same DNA concentration, **liposomes with αHL under stronger T7Max promoter exhibit higher fusion efficiency**. Encouraged by this, we proceeded with selection experiments.

In all selection experiments, two populations of synthetic cells were prepared separately, one encapsulating αHL under regular T7 promoter, and one with αHL under stronger T7Max promoter. All cells contained the 90kbp genome, PURE translation system and Phi29 genome replication system.

Cells contained GFP plasmid in only one of the populations: either in T7 αHL population, or the T7Max αHL population. In each experiment, two populations were mixed: always one population with GFP was mixed with one population without GFP. After five generations of genome replication, growth and division, samples were analyzed using flow cytometry. In cells with the weaker T7 αHL promoter, the GFP liposomes constituted 34% of the total – down from the initial 50% at the start of the experiment (**Figure 4i** and **k**). In the reverse experiment, where GFP gene was in population with αHL under the stronger T7Max promoter, we found that 58% of liposomes contained GFP after five generations (**Figure 4l** and **k**). After just five generations, cells with stronger T7Max αHL outcompete cells with weaker T7 αHL. **This demonstrates a case of selection in populations of synthetic cells: a beneficial mutation that enables more efficient feeding** (the stronger T7Max promoter) **results in more offspring.**

### Metabolic load affects growth and division

We noted that **the weaker cells lose more than the stronger cells gain**. Since the experiments are analogous, the loss of weaker cells should match the gain of stronger cells. Yet cells with GFP in T7 αHL lost 18% of share in total population, while cells with GFP in the T7Max αHL gained only 9%. This could be explained by **uneven metabolic loads**. Synthetic cell metabolism is not very robust, so the added load of expressing GFP reporter might have burdened the cells enough to create a noticeable difference.

When GFP was in T7 αHL cells (the weaker growers), the effect was cumulative: those cells were both handicapped by the weaker promoter, and burdened by GFP expression. In the reverse experiment, where GFP was in T7Max αHL cells, the stronger promoter still allowed those cells to gain in numbers, but slower than their unburdened counterparts in the previous experiment.

To rule out that the difference in abundance is not caused by one of the plasmids being faster to amplify, we compared Phi29 replication yields for T7 and T7Max plasmids. There was no significant difference in replication yields (**Figure S52**), indicating that Phi29 does not favor one or the other plasmid, and thus does not affect the outcome of the selection experiments.

Next, we directly tested whether the added metabolic load (expressing fluorescent reporter) affects fitness of the cells. We repeated the generation experiments, this time with both populations carrying a fluorescent protein marker. The marker was either GFP or mCherry; both proteins are of similar size. We started with an equal ratio of the GFP and mCherry populations, with either fluorescent protein paired with either T7 or T7Max αHL (see legend under **Figure 4o**). After normalizing the fluorescence data to the calibration curve (Methods section 18), we found that when both populations expressed a fluorescent protein marker, the difference in gain of T7Max (11%) and in loss of T7 (10%) was nearly identical. **The observed coupling of metabolic load to fitness illustrates how relatively complex, population-level regulatory mechanisms can already be observed in synthetic cells**.

We used sequencing to confirm that at the end of the experiment one genome indeed increases in abundance over the other. We prepared samples with equal starting ratio of T7 to T7Max. To better mimic the situation when new mutation slowly emerges and rises in the population, we also prepared samples with 9-to-1 ratio of T7 (the “old genome”) to T7Max (the “new mutation”). After five generations of cell cycle, genomes were sequenced (Methods section 16). We performed SNP/INDEL detection analysis, with T7 as the “original” sequence and the T7Max as the “variant”. (**Figure 4n**) In samples without growth and division, the sequence abundance closely matched the starting ratios (1 to 1 or 9 to 1, respectively). In samples with the five generations of complete cell cycle, the T7Max gene abundance increased. Cells that started at equal ratio, ended up with 61% of T7Max (**Figure 4o**). When T7Max started as only 10% of the original population, after five generations the “new mutation” of T7Max already constituted 38% of the population. **This process is similar to the spread of beneficial mutation through selection in live cells.**

### Competition for feeding resources

Encouraged by the experiments showing that faster growing cells can gain a larger share of the population, we investigated if such advantage persists in a resource limited environment. In all experiments described above, the feeder liposomes were used in excess of cells. To mimic resource limiting conditions, we mixed two cell populations: fast growing cells with T7Max αHL and slower growing cells with T7 αHL, and we proceeded with cell cycle experiments using varying amount of feeder liposomes. (**Figure 5a**) We used 1x (the same amount of feeder liposomes as used in previous experiments, 2mM total lipid concentration), and progressively less feeder liposomes: 0.5x (1mM total lipid in feeder liposomes), 0.25x (0.5mM lipid) or 0.1x (0.2mM lipid). Each feeding conditions were applied to two sets of experiments. In one set, the T7Max αHL cells carried GFP and the T7 αHL cells carried mCherry (the genomes were otherwise identical). In the other set of experiments, the fluorescent marker proteins were reversed: the T7Max αHL cells carried mCherry and the T7 αHL cells carried GFP. The proportion of T7Max cells to T7 cells was 1-to-1 at the start of the experiment.

After five generations, the flow cytometry quantified green cells in each sample. Under the 1x feeding conditions, T7Max αHL cells that carried GFP were 60% of the total population (from a starting point of 50%), while when GFP was in the T7 αHL cells, the GFP cells were 41% of the total population. This result was consistent with the competition results shown earlier (**Figure 4**). In the resource limited feeding experiments, with 0.5x, 0.25x and 0.1x feeder liposomes, the **faster growing T7Max cells gained progressively more advantage over slower growing T7 cells**. In the most resource limited feeding experiment, with 0.1x feeder liposomes, 70% of total population was fast growing GFP cells, compared to only 29% slow growing GFP cells (**Figure 5b, g, h, i, j**). **Under resource limited conditions, the fast-growing cells gain progressively more advantage over the slow growing cells.**

We used qPCR to quantify the amount of new DNA. After DpnI digest (to remove the genome originally inserted into synthetic cells) we quantified new GFP, mCherry and pLD1 DNA. The pLD1 was used as a “housekeeping control,” because this plasmid was present in both fast and slow growing populations. As expected, the results matched the flow cytometry data. Each sample had more of the fluorescent reporter gene present in the T7MAX fast growing cells: when T7Max αHL cells carried GFP, we found more GFP gene (**Figure 5c**), when T7Max αHL cells carried mCherry, we found more mCherry gene (**Figure 5d**). The amount of pLD1 in all samples was similar under each feeding condition (**Figure 5e**).

The total amount of new DNA in cells with less feeder liposomes was lower than in cells with access to more feeders. This indicates that the lower amount of feeder liposomes indeed created resource limited conditions, where no cells were able to grow as much as they would if food wasn’t scarce. This created true competition for resources, and the fast growers were the winners.

Directly comparing the number of GFP cells, and the amount of new DNA, in fast growing T7Max cells vs slow growing T7 cells, shows that under all feeding conditions T7Max cells grow faster, with greater gains under the most resource limited 0.1x feeding conditions (**Figure 5b-f**). In this minimal non-autonomous system, we already see the emergence of the maxim “**the rich get richer during global crisis.**” As feeder liposomes were scarcer, the fast growers managed to gain more advantage over the slow growers.

### Genetically encoded division

Natural cells divide because of forces of the cytoskeleton. In synthetic cells, we do not yet have a functional cytoskeleton.^40,42^ Thus, to achieve division we relied on a physical membrane deformation mechanism. Specifically, protein crowding on the surface of liposomes can induce division of liposomes^68^,69. So inspired e utilized our ability to display peptide tags on the surface of synthetic cells to engineer a simple genetically-encoded division mechanism.

### Modified αHL can induce division

Synthetic cells expressing His tag labeled αHL were incubated with Ni-NTA biotin linker, and streptavidin. The binding of Ni-NTA to His tag, and binding of biotin to streptavidin, enabled attachment of streptavidin to the surface of synthetic cells (**Figure 6a**). Based on protein crowding principles, we expected streptavidin to induce membrane curvature and division, much like was previously demonstrated with GFP ^69^.

Initial experiments confirmed that synthetic cells with streptavidin attached to the surface are, on average, smaller than synthetic cells that did not have either αHL, streptavidin or Ni-NTA linker (**Figure 6b**).

Mindful of potential disadvantages of the size measurement using dynamic light scattering in heterogeneous populations like synthetic cells, we characterized division by more quantitative methods. We immobilized synthetic cells on magnetic beads using click chemistry (Methods section 20). If division occurs, the “parent” cells should remain attached to beads, and “daughter” cells can separate from the beads. (**Figure 6e**) After incubation with streptavidin and Ni-NTA linker, we collected the cells still attached to the beads (the “parent” fraction), and the cells separated from the beads (the “daughter” fraction).

As expected, in all samples containing cells with αHL, we detected αHL DNA in the “parent” cells still attached to the beads (**Figure 6g**). In the “daughter” cell fraction, we only detected αHL DNA in the sample with all division elements present: αHL, streptavidin and Ni-NTA linker (**Figure 6f**). Only if the cells divide was a detectable amount of daughter cell genome found free off the beads, confirming division, **demonstrating synthetic cell division induced by expression of a protein inside the cell.**

### Stronger expression of αHL leads to more daughter cells

Next, we took advantage of the slow (T7) and fast (T7Max) promoter system described earlier, to investigate competitive division. Two populations of synthetic cells were mixed and immobilized on beads: one contained αHL under the slow T7 promoter, the other population contained αHL under the fast T7Max promoter. Since T7 and T7Max are difficult to differentiate directly by qPCR, we used a unique DNA marker to label the T7 and T7Max cells (Methods section 21). After division, all “parent” fractions, cells still attached to the beads, as expected contained either T7, T7Max, or both markers (**Figure 6i**). Off bead daughter containing markers for T7 and T7Max were observed only if division was enabled by the presence of streptavidin and linker (**Figure 6h**). In samples where both T7 and T7Max cells were present, the amount of T7 marker in daughter cells was lower than the amount of T7Max marker (1.8nM vs 3.3nM), indicating **competitive division driven by genetic differences among synthetic cells --** the faster T7Max cells produced more daughter cells than the slower T7 cells.

### Genetically encoded growth and division can be combined

To **combine growth and division**, we prepared synthetic cells encapsulating the entire 90kbp genome, and two αHL plasmids: the His tagged and the FLAG tagged αHL. Those cells were immobilized on magnetic beads and fed with Ni-NTA labeled feeder liposomes. After feeding, biotin FLAG antibody linker and streptavidin were added to induce division (**Figure 6j**). We tested all possible combinations of DNA replication (in the presence or absence of Phi29), growth (feeder liposomes with or without Ni-NTA lipids), and division (with or without biotin FLAG antibody). Both “daughter” free off beads and “parent” attached to beads cells were analyzed. In all “parent” samples we detected, as expected, a generation counter oligo in samples that were fed (**Figure 6k**). αHL DNA was detected in all “parent” samples (**Figure 6m**), while newly synthesized αHL DNA was detected only in samples with Phi29 (**Figure 6l**). In daughter cells, the generation counter oligo and DNA were detected only in samples that did undergo division – confirming that division is necessary to produce daughter cells. Generation counter was detected in daughter cells that had feeder liposomes and divided (**Figure 6n**), αHL DNA was found in all daughter cells that divided (**Figure 6p**), and newly synthesized αHL DNA was found, as expected, only in daughter cells with Phi29 and division (**Figure 6o**). The αHL quantification here does not distinguish between FLAG and His tagged variants – the data shows total αHL gene.

We used qPCR to confirm that daughter cells inherited the multi-partite genome. Each of the plasmids comprising the 90kbp genome was detected in the daughter cell fraction that went through DNA replication, feeding and division (**Figure 6r)**.

This experiment demonstrated that **synthetic minimal cells with 90kbp genome can undergo both genetically encoded growth and division**.

### Complete synthetic cell cycle with selection

The synthetic cell cycle shown here represents key steps towards the development of artificial life. We demonstrated that **synthetic cells can undergo cycles of genetically encoded feeding**, **coupled with replication of genetic material**, **and followed by division into new daughter cells**. We engineered a genetically encoded mechanism for the growth and division of synthetic cells, and we demonstrated that this genetically encoded growth mechanism can be used to spread a beneficial mutation through the population, resulting in **selection**. We demonstrated how in a resource limited environment the advantages of the faster growing population increase, as food availability decreases, illustrating **competition for resources**. We demonstrated a genetically encoded mechanism for the division of synthetic cells that does not require cytoskeleton. We have combined the feeding and division, demonstrating that **synthetic cells can grow and divide via genetically encoded mechanisms.**

All living cells require external intervention to support life. For example, if a cell culture plate does not get passaged into fresh media, the cells die. Similarly for synthetic cells, intervention might entail feeding (via feeder liposomes) and providing streptavidin necessary for division. This work presents the first example of a **cell cycle, selection and competition, and reproduction, in minimal synthetic cells** created from non-living building blocks. The identity and concentration of each building block is known, and can be controlled. This brings us closer to **defining the possible minimal chemical composition of life.** A minimal cell displaying key hallmarks of life, like cell cycle and reproduction, also enables building an accurate computational model of life, improving on already advanced minimal non-living cell models.^70^ Our cell is not as capable or complex as living minimal cells, while the function of all its essential genes are fully defined.

The selection demonstrated here is directly coupled with feeding and growth, like in the case of Darwinian evolution in nature. Cells that carry the mutation enabling more efficient growth (stronger promoter for αHL) produce more daughter cells. Over several generations of cell cycle, the population distribution changes, with increased amount of the cells carrying the advantageous mutation. This contrasts with artificial selection methods demonstrated *in vitro* before, where external selection procedure (like pull down column) was needed. Synthetic cells presented here undergo selection: **the beneficial mutation results in more offspring**. Since the beneficial mutation did not arise spontaneously in the population, but was instead introduced artificially, this is different from the natural Darwinian evolution. Next step towards more robust synthetic life will be to turn the selection we just demonstrated into a true Darwinian evolution, by enabling spontaneous rise of mutations.

Mechanism presented in this work demonstrates genetically encoded division of synthetic cells, enabling future work in coupling growth, division and evolution. Further work is needed to develop more robust, higher yield and controllable division mechanisms^71,72^. This will likely eventually require the development of a synthetic cytoskeleton. However, our work demonstrates that simple genetically encoded replication is possible in the absence of cytoskeleton. Supplementary text section 9 provides more in-depth discussion on future directions for engineering replication and robust metabolism in synthetic cells.

Overall, **we have demonstrated key milestones towards construction of synthetic life: a complete cell cycle, including growth and division, and selection, in minimal cells with known identity of all components**. This can serve as a chassis for further optimization of synthetic cells undergoing Darwinian evolution, advancing the field towards robust artificial life.

## Acknowledgments

We thank Hannes Mutschler and Kai Libicher for advice on troubleshooting the pREP reactions and for the gift of the pREP plasmid. We thank Vincent Noireaux for advice on T7 RNAP expression and for the generous gift of αHL and T7 RNA polymerase plasmids. We thank Anthony Forster for the gift of the pLD plasmids. We thank Drew Endy for helpful comments and feedback on framing of this work.

Figures were created with elements from BioRender.com. This work was supported by John Templeton Foundation award 61184, the generous gifts from Jeremy Wertheimer, Alfred P. Sloan Foundation awards G-2022-19420 and G-2024-22710, and NSF Awards 2338121 and 2419641.

## Materials and methods

Sequences of primers used in this work are listed in **Table S2**. Sequences of synthetic cell genome plasmids are in **Table S7**.

Wavelengths used in fluorescent measurements in this work are listed in **Table S5**.

For quick reference and finding the methods corresponding to each part of the project, below is a list of methods sections relevant to each main figure of this paper:

Figure 2: methods sections 1, 3, 4, 5, 7, 8, 9, 10, 14, 15, 25, 26, 27

Figure 3: methods sections 1, 3, 4, 5, 6, 11, 12, 13, 14, 15, 16, 23, 24, 26, 27

Figure 4: methods sections 1, 2, 3, 4, 5, 6, 11, 12, 14, 15, 17, 18, 26, 27

Figure 5: methods sections 1, 2, 3, 4, 5, 6, 11, 12, 14, 15, 17, 18, 26, 27

Figure 6: methods sections 1, 2, 3, 4, 6, 11, 12, 14, 15, 19, 20, 21, 22, 26, 27

### Translation systems

Most of this work was done in PURE, with some preliminary experiments in whole cell extract TxTl.

The only part of this work done with whole cell lysate TxTl was data on figure 2, described in the main paper section “Genetically encoded liposome fusion leads to growth”, accompanied by the experiments described in SI text sections 1 and 3 and accompanying SI figures.

PURE translation system was used for all other results, shown on main figures 3, 4, 5 and 6 and all corresponding SI figures and SI text sections. All results described in main paper sections “Five generations of synthetic cell cycle”, “Selection”, “Competition for feeding resources”, and “Genetically encoded division” are obtained with PURE.

### Methods table of content

1. General conditions and replicates
2. pREP Reaction
3. Plasmids and cloning
4. Synthetic cell liposome preparation
5. Fluorescent labeling of liposomes
6. Feeder liposomes
7. Fusion experiments
8. VAMP2 peptide
9. FRET
10. Whole-cell extract TxTl
11. Cell cycle with feeder liposome growth and mechanical division
12. Generation counter
13. Synthetic cell media optimization
14. qPCR
15. RT-qPCR
16. Next Generation Sequencing
17. Flow cytometry
18. Selection and competition
19. Dynamic Light Scattering
20. Immobilization of synthetic cell liposomes on beads
21. Genetically encoded division of synthetic cells
22. Genetically encoded growth and division
23. Single cell plasmid abundance
24. Dialysis of feeder liposomes
25. Western blot
26. Summary of statistical analysis
27. Size exclusion purification

## 1. General conditions and replicates

All experiments shown in this paper were done in triplicate. For liposome experiments, each replicate was prepared from a separate thin film of lipids, through separate liposome formation and separate incubation. All replicates for fluorescence, qPCR, RT-qPCR, sequencing, cytometry, DLS and gel analysis were prepared from independently performed experiments, and in case of a liposome sample undergoing this analysis the replicate was independently prepared starting from separate thin film. Unless otherwise indicated, synthetic cell reactions were incubated at 30°C.

Unless otherwise stated, liposome experiments were performed in PCR tubes with gentle shaking. All experiments were performed under RNAse free regime, with RNAse free equipment.

## 2. pREP Reaction

Most of the protein expression experiments in this project were done with pREP transcription-translation reaction: all experiments with Phi29 genome replication, cell cycle, feeding and growth, were done with pREP.^20^

All plasmids used for pREP reactions were obtained via mini-prep (Epoch 2160250) followed by a second clean up via an PCR clean up kit (Epoch 2360250), then concentrated via lyophilization (see methods section 3 for plasmid preparation information).

This protocol was adapted from a previous publication^20^. The 10x energy mix is stored at −80°C and composed of the following: 700mM potassium Glutamate, 79mM Magnesium Glutamate, 1M HEPES pH 8.0, 250mM Creatine Phosphate (Sigma-Aldrich 27920-5G), 3.75mM Spermidine (Sigma-Aldrich S2526-5G), 5.18g/L *E. coli* tRNAs (Sigma-Aldrich 10109550001), 60mM DTT (Goldbio DTT50). Additionally, it contained 3.6mM of each L-amino acid: Alanine, Arginine, Asparagine, Aspartic acid, Cysteine, Glutamic acid, Glutamine, Glycine, Histidine, Isoleucine, Leucine, Lysine, Methionine, Phenylalanine, Proline, Serine, Threonine, Tryptophan, Tyrosine and Valine.

The 50x rNTP mix is premixed, stored at −80°C and composed of the following: 18.75mM ATP (Larova ATP_1000G), 12.5mM GTP (Larova GTP_1000G), 6.25mM CTP (Larova CTP_1000G), 6.25mM UTP (Larova UTP_1000G).

The following reagents were combined at the following final concentrations: 1x Energy Mix from above, 1x rNTP mix from above, 0.1x PURExpress Solution A (NEB E6800L), 60% of total volume PURExpress Solution B (NEB E6800L), 0.6mM dATP, 0.6mM dGTP, 0.6mM dCTP, 0.6mM dTTP (Denville Scientific CB4420-2), 4mM DTT (Goldbio DTT50), 0.4U/μl NxGen Phi29 (Lucigen 30221-2), and plasmids.

For most experiments, unless otherwise indicated, reactions were incubated at 30°C for 8 hours.

## 3. Plasmids and cloning

Sources of plasmids not cloned in our lab are indicated in the text. We thank Dr. Anthony Forster for a generous gift of pLD1, pLD2 and pLD3 plasmids.^65^ We thank Dr. Vincent Noireaux for generous gift of αHL and T7 RNA polymerase plasmids.

All plasmids under T7Max promoter were cloned into a vector we have previously prepared^67^. This vector contains UTR1 and T500 terminator sequences optimized for bacterial in vitro translation.^73^

All cloning was done using either Gibson assembly (using NEB enzyme kit E2611), or site directed mutagenesis (using NEB Q5 polymerase M0493 and NEB KLD enzyme mix M0554)

In a typical translation experiment, each DNA plasmid was at 10nM final concentration. Unless otherwise stated, when more then one plasmid was used, each was introduced at the same concentration, except in experiments with the whole 90kb genome, where each plasmid was at 2.5nM concentration.

All plasmids were amplified in DH5alpha *E. coli* cells. Plasmids were purified using GenCatch™ Plasmid DNA Mini-Prep Kit (Epoch, 2160250), followed by GenCatch™ PCR Cleanup Kit (Epoch, 2360250) for further purification. This was done based on oral reports from several colleagues who observed higher yields from in vitro translation reactions when using plasmids that were double-purified.

The pREP reactions do not have large amount of “water” volume left after addition of all reagents. To address that problem, if more than 3 plasmids were used, the plasmid stock solutions were mixed, lyophilized, and resuspended in small volume of water (usually 3μl for a 30μl final volume reaction).

## 4. Synthetic cell liposome preparation

Unless otherwise specified, each synthetic cell liposome sample used in the experiments contained 1mM total lipids (this is called “1x” liposomes in all mixing experiments). Liposomes were prepared with DOPE (1,2-dioleoyl-sn-glycero-3-phosphoethanolamine, Avanti 850725P), DOPC (1,2-dioleoyl-sn-glycero-3-phosphocholine, Avanti 850375P) and cholesterol (Avanti, 700100P) at a 1:3:1 molar ratio. Cholesterol was not included in the “total lipid” calculation, so each 1mM liposome sample contained 0.2 mM DOPE, 0.75mM DOPC and 0.25mM cholesterol.

Liposomes were prepared according to the oil extrusion protocol described earlier^35,74^. Stock solutions of all lipids in chloroform were prepared at 1mg/ml of each lipid or cholesterol. Those stock solutions were mixed in amber glass vials to create thin film aliquots for each liposome sample. In a typical experiment, to create 30μl liposomes at 1mM lipid, a thin film aliquot was prepared by mixing 5.58μl DOPE stock, 17.7μl DOPC stock and 2.9μl cholesterol stock. All chloroform stocks were transferred using glass syringes. Chloroform was removed from each vial by drying under gentle flow of nitrogen gas for 6h. Each vial was capped and stored in −20°C until use.

To prepare liposomes, the thin film was suspended in 200μl mineral oil and incubated at 60°C with vigorous shaking for 3 hours, followed by 10 minutes of sonication in a bath sonicator and overnight incubation with vigorous shaking (on a vortexer with continuous shaking attachment) at room temperature. Then, the mineral oil samples were cooled to 4°C while shaking (on a vortexer in a 4°C cold room).

For each sample, 30μL of internal liposome solution was added to the mineral oil lipid suspension and vortexed for 5 minutes at 4°C. That solution was next carefully layered on top of 2 0μL of centrifuge buffer (100mM HEPES, 200mM glucose, pH 8). Samples were carefully transferred to a chilled centrifuge and centrifuged at 18,000rcf at 4°C for 15 minutes. Next, the oil layer was very carefully removed using a gel loading pipette tip (a regular pipette tip would work too, but we preferred gel loading capillary tips for better precision).

Next, with a fresh pipette tip, the liposomes were carefully removed from the bottom of the centrifuge tube and transferred to 250μL of pre-chilled wash buffer (100mM HEPES, 250mM glucose, pH 8).

Samples were centrifuged at 12,000rcf at 4°C for 5 minutes. After that centrifugation step, any residual mineral oil visible on top of the wash buffer was removed, the liposomes were transferred to a new tube with a fresh pipette tip, and resuspended in the final reaction external media buffer. All the liposome preparations steps were performed at 4°C (in the cold room).

### Liposome sizes

For all liposomes in this work, the starting size was guided by the extrusion during initial preparation. For synthetic cell cycle experiments, liposomes were extruded through 2μm membrane (methods section 11). Feeder liposomes were extruded through membrane with 0.4μm pores (methods section 6). For DLS liposomes were extruded through 0.8μm membrane (methods section 19).

## 5. Fluorescent labeling of liposomes

If liposomes were labeled with a fluorescent membrane dye, the dye stock solution in chloroform was added to the thin lipid film chloroform mix. Unless otherwise stated, fluorescent dyes were used at 0.1mol% of total lipid (excluding chloroform), so a sample of 1mM liposomes contained 1μM dye. If two dyes were used in the same liposome population, each was at 0.1mol% (so total 0.2mol% dye).

All dyes were used from stock chloroform solution of 100μg/ml, adding appropriate amount of stock to the chloroform mixture of lipids in the process of preparation of thin film (see methods section 4 above).

The dyes used to label liposomes in this project, and the stock amounts used to prepare sample of 30μl of 1mM liposomes, were:

NBD (NBD-PE, (N-(7-Nitrobenz-2-Oxa-1,3-Diazol-4-yl)-1,2-Dihexadecanoyl-sn-Glycero-3-Phosphoethanolamine, Thermo Fisher N360), 2.86μl stock for each thin film;

Rhodamine (Lissamine™ Rhodamine B 1,2-Dihexadecanoyl-sn-Glycero-3-Phosphoethanolamine, Thermo Fisher L1392), 4.0μl stock for each thin film;

Cy5 (Cy5-PE, 1,2-dioleoyl-sn-glycero-3-phosphoethanolamine-N-(Cyanine 5), Avanti 810335C), 3.79μl stock for each thin film;

Cy7 (Cy7-PE, 1,2-dioleoyl-sn-glycero-3-phosphoethanolamine-N-(Cyanine 7), Avanti 810337C, 3.98μl stock for each thin film.

## 6. Feeder liposomes

The feeder liposomes for the generations of growth and division experiments were prepared according to the liposome preparation protocol described above (Methods section 4). The feeder liposomes were made from the same lipids (DOPE, DOPC, cholesterol at 1:3:1 molar ratio) as all other liposomes in this work, with the addition of Ni-NTA lipids (1,2-dioleoyl-sn-glycero-3-[(N-(5-amino-1-carboxypentyl)iminodiacetic acid)succinyl] nickel salt, Avanti 790404P) at the molar percentage described in each experiment.

The molar percent was calculated relative to total lipid (not including cholesterol). For example, in a 2mM liposome sample with 0.6mol% Ni-NTA lipid contained 12μM Ni-NTA lipid. The Ni-NTA lipids were added to the initial thin film chloroform mixture, using 0.1mg/ml chloroform stock. For example, for 30μl of 2mM liposome sample with 0.6mol% Ni-NTA lipid, 3.8μl of Ni-NTA chloroform stock was added to the thin film mixture in chloroform.

Feeder liposomes, after liposome formation, were extruded through a membrane with 0.4μm pores, using Avanti mini-extruder (Avanti, 610023) and PC Membranes 0.4μm (Avanti, 610007), using 9 passages in each extrusion. After extrusion, the liposomes were equilibrated for 10 minutes with gentle shaking on an orbital shaker at 4°C (in the cold room), before use in fusion experiments.

For cell cycle experiments, each feeder liposome population contained 0.6mol% Ni-NTA lipid, 4nM of one of the “bottom” oligos for the generation counter (see methods section 12), and a complete 1x reaction mixture for pREP reaction (methods section 2). Unless otherwise stated, 0μl of 2mM of feeder liposomes were used for each generation.

## 7. Fusion experiments

For all the fusion experiments, liposomes were prepared as described above (methods sections 4 and 6). All fusion samples were prepared at 1mM total lipid concentration, unless otherwise stated in the description of the experiment.

For samples with externally added fusion tags, the liposomes were decorated with fusion tags according to the previously described protocols^75^. See section below for Vamp-2 protein details. The samples were incubated with fusion tags for 30 minutes at room temperature, tumbling gently on an orbital shaker.

For samples with αHL as the fusion tag, the synthetic cells were prepared as described above and incubated for three hours at 30°C with gentle shaking on an orbital shaker inside of an incubator.

After the initial incubation with fusion tags, or incubation to express αHL, the liposomes were mixed at the indicated ratios. In most experiments, equal volumes of 1mM total lipid samples were mixed, unless indicated otherwise in the experimental description. The fusion samples were incubated for 12 hours at 30°C with gentle shaking. After the incubation, samples were transferred to a 96-well plate and fluorescence was recorded. If RT PCR, qPCR, or Western Blot analysis was needed, the liposomes were lysed with 0.1% Triton-X 100 (Acros Organics, AC215682500).

In a typical fusion experiment, the liposome samples were mixed with gentle pipetting and incubated for 6h (unless otherwise indicated) at 30°C with gentle shaking (on a shaker inside incubator). The mixing experiments were done in PCR tubes.

Unless otherwise stated, each liposome sample used in fusion experiment was at 1mM total lipid (or 1mM + Ni-NTA and/or dye, if used).

## 8. VAMP2 peptide

The His tag labeled 6xHis-VAMP2 peptide was purchased through a custom synthesis order from GeneScript, prepared via solid phase peptide synthesis. The peptide sequence was the VAMP (synaptobrevin) trans-membrane domain of the E3-VAMP2 peptide used in liposome embedding experiments before, HHHHHHGRKYWWKNLKMMIILVICAIILIIIIVYFST.^75^

The peptide was delivered as a lyophilized powder, which was resuspended in 50mM HEPES pH 8.0 to create a 0μM stock solution. The solution of peptide was aliquoted in μL portions, to avoid repeated freeze-thaw cycles.

Unless otherwise indicated, the VAMP peptide to lipid molar ratio was 1:500. For example, samples of 1mM lipid contained 2μM VAMP peptide.

To label liposomes, we used a previously described procedure ^75^: after preparation of liposomes, typically a 0μL samples of 1mM total lipid was mixed with 1.2μL of peptide stock. The samples were mixed by vigorous pipetting for a minimum of 1 minute. The VAMP labeling was done at 4°C (in the cold room), to avoid initiating translation reactions inside the liposomes.

## 9. FRET

Membrane growth was measured using FRET experiments. Two different FRET pairs were used: either NBD-Rhodamine, or Cy5-Cy7. Generally, NBD-rhodamine was used when there was no GFP signal measured in the same samples, and Cy5-Cy7 was used when GFP (spectrally overlapping with NBD) was produced in the samples.

The labeling of liposomes with fluorescent dye is described in methods section 5.

Membrane growth experiments were monitored by tracking wavelength of the donor dye in each FRET pair, which was NBD and Cy5 for the pairs used in this project. See **Figure S14** for the general scheme of FRET membrane growth experiment. One population of liposomes was labeled with both FRET dyes, the other population did not carry membrane dyes. Upon fusion and membrane growth, FRET dyes were diluted in the membrane, decreasing FRET effect and increasing measurable donor fluorescence.

FRET signal was calibrated to % lipid mixing by preparing control samples of liposomes with decreasing amounts of FRET dyes corresponding to dilution upon membrane mixing. Sample representing 0% membrane mixing was 1mM liposomes with 0.1mol% of each of the FRET dyes (so total 0.2mol% FRET dyes). Sample representing 50% lipid mixing was 1mM liposomes with 0.05mol% each dye, etc.

All control liposomes were prepared from thin lipid films prepared as described above (see sections 4 and 5), then samples were extruded through 0.4μm pores, using Avanti mini-extruder (Avanti, 610023) and PC Membranes 0.4μm (Avanti, 610007). Fluorescence was measured using wavelengths listed in Table S5. The FRET calibration is on **Figure S76**.

## 10. Whole-cell extract TxTl

Most of the protein expression experiments in this project were done with pREP transcription-translation reaction (see methods section 2). Due to high cost and lower protein yields of pREP compared to the whole cell extract TxTl, we used TxTl for experiments where pREP was not required: establishing and characterizing the genetically encoded synthetic cell growth system. <u>TxTl was only used</u> <u>to collec data demonstrated on</u> Figure 2 <u>and its’ corresponding SI figures</u>.

Protein expression experiments with TxTl were performed using whole cell *E. coli* extract according to the protocol described before^76^. To grow cells for the extract, 2xYT+P was prepared by mixing 124 g 2xYT powder, 160ml potassium phosphate dibasic 1M solution, 88ml potassium phosphate monobasic 1M solution and 3.752L water (to total 4L), sterilized in the autoclave on liquid cycle and stored at 4°C until use. Rosetta 2(DE3) cells were grown in 2xYT+P media until an OD of 1.5 to 2 was reached. The culture was then chilled and washed twice with cold S30A buffer (10.88g Mg-glutamate powder, 24.39g K-glutamate powder, 50ml of 2M Tris base, and pH adjusted to 7.4 with glacial acetic acid). The cells were lysed using sonication, with energy settings according to the previously described protocol ^77^.

The lysate was centrifuged at 18,000rcf at 4°C for 30 minutes and the supernatant was shaken at 2 0rpm at 7°C for one hour for the “runoff reaction.” The lysate was then centrifuged again at 18,000rcf at 4°C for 30 minutes and the supernatant, the final cell extract, was aliquoted and stored at - 80°C.

Due to the batch-to-batch variability between different preparations of the cell extract, all data reported on the same figure in this paper were prepared using the same batch of cell extract.

The final TxTl reaction was prepared by mixing cell extract, energy mix, amino acid mix, and salt mix to the following final concentrations: 130mM potassium glutamate, 10mM ammonium acetate, 10mM magnesium glutamate, 1.5mM ATP and GTP, 0.9mM CTP and UTP, 0.068mM folinic acid, 200ug/mL of *E. coli* tRNA mixture, 0.33mM nicotinamide adenine dinucleotide (NAD), 0.26mM coenzyme-A (CoA), 1.5mM spermidine, 4mM sodium oxalate, 0.75mM cAMP, 30mM 3-PGA, 50mM HEPES pH 8, 2mM of each of the 20 amino acids in a KOH amino acid stock.

Unless otherwise stated in the experimental description, the complete TxTl reaction was incubated for 12 hours at 30°C.

## 11. Cell cycle with feeder liposome growth and mechanical division

For each growth and division experiment in the 5-generation cell cycle, liposomes were prepared as described above (methods section 4). At the beginning of the experiment, each sample of liposomes (1mM total lipid concentration, 30uL) was incubated for 12 hours at 30°C. This was considered generation 0. The sample was then mixed with 30μL of 2mM feeder liposomes (see methods section 6 for preparation), starting generation 1. The mixture of the starting population and the feeder liposomes of generation 1 was cultured, with gentle tumbling, at 30°C for 12 hours.

For each generation, feeder liposomes contained 4nM of one of the “bottom” oligos for the generation counter (see methods section 12), and a complete 1x reaction mixture for pREP reaction (methods section 2), without any DNA template. Feeder liposomes were labeled with Ni-NTA lipids in the membrane (methods section 6).

At the start of cell cycle experiment, the “generation 0” synthetic cells contained 1nM of each of the “top” strands for the generation counter (methods section 12), a complete pREP reaction (methods section 2), and DNA templates (typically 10nM each, see methods section 3).

Typical 0μl lumen of “generation 0” synthetic cells for a cell cycle experiment contained: complete 1x pREP reaction (methods section 2), 4μl of plasmid mix (methods section), 1μl of a mix of top strands for generation counter and 1μl T4 DNA Ligase (methods section 12).

An orbital shaker placed inside an incubator was used for the incubation steps after addition of feeder liposomes. After 12 hours at 30°C, the samples were extruded using a mini-extruder (Avanti, 610023) with a 2μm membrane (Whatman, 76457-184). This concluded generation 1.

The extruded sample was then mixed with another 30μl of 2mM total lipid concentration of feeder liposomes, and this started generation 2. This, and each following generation, was similarly cultured for 12 hours at 30°C with gentle shaking.

Starting after generation 3, after each 12-hour incubation followed by extrusion, samples were analyzed for fluorescence, abundance of DNA, mRNA and generation counter. First, fluorescence of GFP was measured using a plate reader (Molecular Devices SpectraMax). Then, liposomes were lysed with 0.1% Triton-X 100 (Acros Organics, AC215682500), 0.5M stock EDTA was added to final concentration of 10mM, and sample was incubated at 65°C for 15 minutes to heat inactivate the synthetic cell enzymes.

Samples for analysis of newly synthesized DNA abundance were digested with DpnI (NEB R0176) using 5 units of enzyme per reaction in 1x rCutSmart buffer (NEB B6004), incubated 30min at 37°C, followed by heat inactivation for 15min at 80°C. Then all samples were concentrated by lyophilization, and resuspended in 1 μl of water. Those samples were used for subsequent analysis of abundance of DNA, mRNA and generation counter RNA (methods section 15).

The cell cycle experiments shown in this work included fluorescent proteins expressed by the synthetic cells (GFP in cell cycle experiments, GFP and mCherry in cell cycle competition experiments). To confirm that the presence of fluorescent proteins doesn’t change the membrane dynamics and cell cycle results, we repeated 5 generations of cell cycle experiments with synthetic cells without GFP. We analyzed new DNA and mRNA abundance, comparing it to the results from a similar experiment in presence of GFP, results are shown on **Figure S40**.

We also tested membrane stability to small molecule leakage in presence of fluorescent proteins. Synthetic cells were created with either GFP or mCherry plasmid. The GFP cells contained 1mM of water soluble Cy5.5 (Cyanine5.5 carboxylic acid, Luminprobe 17090), and the mCherry cells contained 1mM calcein. After 12h incubation at 30°C, cells were purified (methods section 27) to measure leakage of the small molecule dye. No leakage was observed with either of the fluorescent proteins, data on **Figure S41** for cells with GFP protein and Cy5.5 small molecule dye, and **Figure S42** for cells with mCherry protein and calcein small molecule dye.

To investigate the mRNA stability, DNA stability and fluorescent protein accumulation over the course of many hours of synthetic cell “life cycle”, we performed control experiments with cells containing GFP plasmid and αHL plasmid. Synthetic cells were prepared with 5nM GFP plasmid and 5nM αHL plasmid.

Cells were incubated in the synthetic cell culture media (see Supplementary text section 4) for 30 hours. The GFP mRNA, DNA and GFP fluorescence were measured at indicted time points, see **Figure S43**.

### The rationale for mechanical division in cell cycle experiments

In this work, we used more reliable mechanical division to decouple the variables: we wanted to optimize conditions for the cell cycle of growth and DNA replication, and to investigate the competition and selection processes enabled by the genetically encoded feeding mechanism. The mechanical division, being an older and widely used technique for dividing liposomes, ensures complete division and uniform size of daughter cells.

Mechanical division was used in experiments shown on Figures 3, 4 and 5.

For figure 6, the genetically encoded division was used instead of mechanical division, resulting in daughter cells that are less uniform in size (as seen in microscopy images on Figure 6 and **Figure S73**).

## 12. Generation counter

The sequences of all generation counter oligos are in **Table S3**. All generation counter oligos were purchased from IDT DNA as custom synthesized RNA oligos and used without further purification.

The generation counter reactions require T4 DNA ligase to ligate the “sticky ends” of the generation counter oligos (see **Figure S45** for the generation counter reaction scheme). The T4 DNA ligase typically is used in buffer containing: 50mM Tris-HCl pH 7.5, 10mM MgCl2, 1mM ATP, 10mM DTT (per NEB information for product B0202S). However, since this reaction will be required to work in pREP reaction conditions in cell cycle experiments, we have tested and optimized the reaction using pREP buffer conditions.

In each generation counter validation experiment, 1nM of each generation counter oligo was mixed. Each 20μl reaction thus contained 0.02pmoles of each generation counter oligo (unless otherwise stated, in negative control experiments where some oligos were omitted). All generation counter oligos were stored as separate 20nM stocks in water. For each experiment, 1μl of each stock was mixed, and the resulting mixture of generation counter oligos was lyophilized and resuspended in 2μl water. This mixture was used in the generation counter reaction.

Each 20μl reaction contained 1μl T4 DNA Ligase (NEB, M0202), 1nM of each generation counter oligo (from the mixture described above), 0.6mM dATP, 0.6mM dGTP, 0.6mM dCTP, 0.6mM dTTP (Denville Scientific CB4420-2), 1x pREP energy mix (see methods section 2) containing 70mM potassium glutamate, 7.9mM magnesium glutamate, 0.1M HEPES pH 8.0, 25mM creatine phosphate (Sigma-Aldrich 27920-5G), 0.375mM spermidine (Sigma-Aldrich S2626-5G), 0.5g/L *E. coli* tRNAs (Sigma-Aldrich 10109550001), 10mM DTT (Goldbio DTT50), 0.3775mM ATP (Larova ATP_1000G), 0.25mM GTP (Larova GTP_1000G), 0.125mM CTP (Larova CTP_1000G) and 0.125mM UTP (Larova UTP_1000G).

Most of those components were not essential for the generation counter ligation. Those components were included so that the reaction conditions match the final application conditions in cell cycle experiments.

The reactions were incubated at 30°C for 12h. 30°C is not optimal temperature for those ligation reactions, but since this was the temperature required in the cell cycle experiments with pREP, this was the temperature we used to optimize the reaction.

After completion of the reaction, the generation counter products were analyzed using RT-qPCR (see methods section 15). For each rt-qPCR reaction, 4. μl of the 20μl generation counter ligation reaction was used as a template.

In cell cycle liposome experiments (methods section 11), each starting “parent” synthetic cell population contained 1nM of each of the “top” strands for counter for each generation. Those strands were mixed from the 20nM stock, lyophilized and resuspended in 1μl, before adding to the liposome lumen solution (see methods section 4 for liposome preparation). The feeder liposomes (methods section 6) contained 4nM of one of the “bottom” generation counter oligos, one for each generation those feeder liposomes were used in. The concentration of “bottom” oligo in feeder liposomes was higher than earlier used 1nM to account for volume dilution when smaller feeder liposome fuses with larger synthetic cell.

We quantified the abundance of full length generation counter product in single cell experiments after 3, 4 and 5 generation (**Figure S37**).

## 13. Synthetic cell media optimization

Each sample in the media optimization experiments was prepared with 1mM of liposomes (methods section 4), 10nM of αHL and GFP plasmids, and the internal pREP mixture as described above (methods section 2).

The external media mixture was prepared as indicated in the results **Figure S15**, where 1x indicated the energy mix, amino acid mix, and salt mix of the concentrations and compositions described below, and the rage of 0.5x and 1.5x are the 1x scaled appropriately.

The 1x energy mix contained: 1.5mM ATP and GTP, 0.9mM CTP and UTP, 0.068mM folinic acid, 200 ug/mL of *E. coli* tRNA mixture, 0.33mM nicotinamide adenine dinucleotide (NAD), 0.26mM coenzyme-A (CoA), 1.5mM spermidine, 4mM sodium oxalate, 0.75mM cAMP, 30mM 3-PGA, 50mM HEPES pH 8.

The 1x amino acid mix contained: 2mM Alanine, 2mM Arginine, 2mM Asparagine, 2mM Aspartic acid, 2mM Cysteine, 2mM Glutamic acid, 2mM Glutamine, 2mM Glycine, 2mM Histidine, 2mM Isoleucine, 2mM Leucine, 2mM Lysine, 2mM Methionine, 2mM Phenylalanine, 2mM Proline, 2mM Serine, 2mM Threonine, 2mM Tryptophan, 2mM Tyrosine, 2mM Valine.

The 1x salt mix contained: 130mM potassium glutamate, 10mM ammonium acetate, and 10mM magnesium glutamate.

For each media optimization experiment, 0μl of 1mM liposomes were diluted in 100μl “media” mixture. Samples were incubated at 30°C for 12 hours. After incubation, GFP fluorescence was recorded using plate reader (Molecular Devices SpectraMax).

## 14. qPCR

Liposome samples for analysis were diluted (μl of sample into 12. μl) with the inclusion of 5ug of RNase A (AG Scientific R-2000-100MG) and incubated at 37°C for 1 hour.

A 20μl OneTaq PCR was set up with NEB OneTaq 2X Master Mix with Standard Buffer (NEB M0482L) with 2μl of the sample as template and following the NEB recommended protocol with the inclusion of 1x of Chai Green Dye (Chai R01200S). qPCR was read and analyzed on either a Chai Open qPCR machine (single or dual channel) or CFX Opus 384 System.

Calibration curves for qPCR quantification are on **Figure S75**.

## 15. RT-qPCR

Liposome samples for analysis were diluted (μl of sample into 12. μl) with the inclusion of 1x Turbo DNase Buffer, and 1U of Turbo DNase (Invitrogen AM2238), and incubated at 37°C for 1 hour.

A 10μl reverse transcription reaction was primed via mixing of 4. μl sample, 0. μl of 2μM primer (the corresponding forward or reverse primer), 0. μl of 10mM dNTPs (Denville Scientific CB4420-2), and 1μl water. This was incubated at 6 °C for 1 minute, followed by at least minutes on ice. Then 2μl of x Superscript IV RT buffer, 0. μl of 100mM DTT, 0. μl of 200U/μl of Superscript IV RT (Invitrogen 18090010), and 0. μl 40U/μl RNase Inhibitor Murine (NEB M0 14S) were added.

Samples were incubated at 2°C for 10 minutes, followed by 80°C for 10 minutes. A 20μl OneTaq PCR was set up with NEB OneTaq 2X Master Mix with Standard Buffer (NEB M0482L) with 2μl of the sample as template and following the NEB recommended protocol with the inclusion of 1x of Chai Green Dye (Chai R01200S). qPCR was read and analyzed on either a Chai Open qPCR machine (single or dual channel) or CFX Opus 384 System.

## 16. Next Generation Sequencing

After 5 generations of growth and replication, samples were lysed, digested with DpnI (to remove all original DNA) and amplified via PCR, which also installed partial Illumina library adapters.

Samples were sequenced using Azenta Amplicon-EZ sequencing protocol using Illumina NGS sequencing. Each cell cycle sample DNA was amplified with primers flanking the promoter region on the αHL plasmid (See **Table S2** for sequences).

This amplification installed partial Illumina adapters, for the forward sequencing read: ’-ACACTCTTTCCCTACACGACGCTCTTCCGATCT- ’ and for the reverse sequencing read: ’-GACTGGAGTTCAGACGTGTGCTCTTCCGATCT- ’.

Amplified DNA was purified on agarose gel and the concentration of the amplicon was measured using PicoGreen dsDNA assay kit (Thermo P7589). Samples were analyzed using SNO/INDEL detection algorithm, using sequence of T7Max promoter as reference sequence.

All samples provided over 75000 usable reads.

## 17. Flow cytometry

Each sample for flow cytometry was prepared by first dialysis (see Methods section 24) to remove feeder vesicles, then diluting liposomes to 0.1mM total lipid concentration in PBS. This was done so all samples contain a similar number of particles (liposomes) per unit of volume. Liposomes in PBS were assessed for GFP fluorescence (488nm excitation, 525nm-550nm bandpass emission filter) at the University of Minnesota Flow Cytometry Resource with a BD LSR II Flow cytometer (Figure 4) or BD FACSymphony™ A3 Cell Analyzer (figure 5). Liposomes were measured at a rate of 150-250 events per second until 10,000 events were recorded in the designated “liposome” gate.

Flow cytometry data was analyzed using FlowJo version 10.8.1 software. Samples were analyzed in a 5mL Polystyrene Round-Bottom Tube (Falcon, REF 352054).

The phosphate buffered saline (PBS) used here was pH 7.4, containing 137mM sodium chloride, 2.7mM potassium chloride, 10mM sodium phosphate dibasic, 1.8mM potassium phosphate monobasic.

Voltages for each measurement: FSC (586V), SSC (255V), and GFP (220V).

Each dot on the plot is defined as an “event”. It’s termed this way because not every event is going to be a sample of interest. An event can be any kind of contamination-dust, single bacteria cell, a clump of proteins, etc. Therefore, a negative and positive control are necessary for flow, because of the need to trial and error finding actual data points. In the liposome measurements shown in this paper, we look at size (FSC), complexity (SSC), and fluorescence (GFP-A).

FSC-forward scatter-size. Sensor is directly across from the laser origin. Measures absence of light as the event passes through the laser. Larger events will block the light from hitting the sensor for an extended period of time.

SSC-side scatter-complexity. Sensor is perpendicular to the laser path. Measures light that was diffracted from its path when the event passed through the laser. More light will hit this sensor if the event is more “complex” aka has components that scatter the light.

FSC and SSC have three components to their measurements:

W-width. time it takes for the liposome to pass through the laser H-height. Maximum intensity of the duration of the width.

A-area. Width times the height of the event

## 18. Selection and competition

The selection and competition experiments using 5 generations of cell cycle were performed according to the general cell cycle protocol (see methods section 11). The starting “1x” population of synthetic cells was 30μL of 1mM liposomes, as described for cell cycle protocol. If two different populations of synthetic cells were mixed for a competition experiment, we used 1 μL of each of the populations.

The “1x” feeder liposomes were 0μL of 2mM feeder liposomes (see methods section 6 for preparation protocol). In competition for resources experiments, where less feeder liposomes were available, we kept the total volume of the sample constant, while varying the concentration of feeder liposomes.

Therefore, “0. x” feeder liposome conditions mean addition of 0μL of 1mM feeder liposomes, “0.2 x” is 0μL of 0. mM feeder liposomes, and “0.1x” is 0μL of 0.2mM feeder liposomes.

All competition and selection experiments were incubated through the cell cycle protocol as described above (methods section 11).

### Normalizing GFP and mCherry fluorescence in selection metabolic load experiments

To estimate what percent of population carries each of the fluorescent proteins, we performed calibration experiments: 5 generations of cell cycle, or no cell cycle, with varying ratio of cells with mCherry and with GFP, with the same αHL in both populations. This resulted in calibration curves translating measured green (GFP) and red (mCherry) fluorescence to relative percent of the synthetic cell population carrying that fluorescent protein gene (see **Figure S56**).

## 19. Dynamic Light Scattering

The DLS experiments were performed using Microtrac NanoFlex DLS Particle Analyzer. The data was analyzed using Microtrac Flex software, assuming the liposomes were spherical. Each sample was prepared by diluting liposomes to approximately 0.1mM total lipid concentration in PBS. This was done so all samples contain a similar number of particles (liposomes) per unit of volume.

All synthetic cell liposomes for DLS experiments were prepared as described in Methods section 4, and then before the start of the experiment the liposomes were extruded through 0.8μm extruder membrane. This was to ensure uniformed size distribution at the beginning of the experiment, mindful of the limitations of DLS technique in analyzing heterogenous samples.

Prior to each sample analysis, a baseline of zero was set using a blank solution (PBS). The average values were obtained based on six individual runs of 120s each.

We recognize the limitations of using DLS: synthetic cell liposomes, particularly ones that undergo spontaneous curvature changes and division, will likely be a heterogenous population, while DLS measurements are the most reliable homogenous samples. Thus, the DLS measurements were used as an initial test for synthetic cell division, to be confirmed with more quantitative measurements in bead experiments.

The only experiments in this paper that used DLS were the division tests.

## 20. Immobilization of synthetic cell liposomes on beads

To immobilize synthetic cell liposomes, the liposomes were prepared with 1mol% of 16:0 azidocaproyl PE (1,2-dipalmitoyl-sn-glycero-3-phosphoethanolamine-N-(6-azidohexanoyl), Avanti 870126P). The liposomes were prepared using the same method used for all other experiments in this project (methods section 4). Each 0μl sample of 1mM liposomes labeled with 1mol% of the azido lipid contains 10μM azido lipid. The azido lipid was added during the preparation of thin film, mixing 2. 4μl of the 0.1mg/ml stock with the rest of the lipids in chloroform.

DBCO tagged magnetic beads (Kerafast FCC433) were washed three times with 50mM HEPES pH 8. The manufacturer’s reported bead loading is 30-50nmol DBCO groups per mg. In each experiment, 5mg beads (approximately 250nmol DBCO) were mixed with 30μl liposomes (0.3nmol azide, although since both inner and outer leaflet are labeled, only half of the total azide groups is available for the conjugation). Azide labeled liposomes were mixed with the DBCO beads and incubated at 30°C for 8 hours. The samples were gently mixed by shaking. During the incubation, the translation reaction inside synthetic cells was also started.

To confirm immobilization rates, beads of three control samples were pulled down (using magnetic PCR tube rack), and the solution from above the beads was collected. After collecting the “supernatant” above the beads, 20μl of 50mM HEPES pH 8 with 0.1% Triton-X 100 (Acros Organics, AC215682500) was added to lyse the liposomes. The lysed liposome solution was also collected. Both the solution from above liposome-beads and the lysed liposome solutions were lyophilized dry, and re-suspended in 2μl 50mM HEPES pH 8. Both solutions were used as template in qPCR reaction with primers for αHL plasmid (present in the liposomes). In all three control samples, αHL plasmid was detected in the samples from lysed liposomes at quantities corresponding to the initial concentration of αHL plasmid (10nM plasmid) – indicating αHL carrying liposomes were immobilized on beads, and no detectable αHL was found in any of the samples from solution collected from above the beads with liposomes (indicating that no detectable free liposomes were left in the solution). This confirmed that the immobilization protocol with large excess of beads to liposome azide tag results in immobilization of all available liposomes on the beads.

Since large excess DBCO beads was used, there are unreacted DBCO groups that could bind to azide on newly divided cells, trapping new daughter cells and defeating the purpose of the experimental design (which assumes daughter cells will be free and separated from beads). To avoid this, we have blocked the unreacted DBCO groups before the division experiments. After initial 8h incubation to couple

synthetic cells to the beads, 7μl of a 50mM stock solution of 3-Azido-L-alanine (Jena, CLK-AA003) in 50mM HEPES pH 8 was added (350nmol azide), and incubated for 4h. After the blocking reaction, samples were ready for division experiments.

## 21. Genetically encoded division of synthetic cells

Purified streptavidin (BioLegend, 280302) manufacturer stock of 1.0 mg/ml (19μM) was diluted in 50mM HEPES pH 8 to a working stock concentration of 5μM and stored at 4°C. Biotin-NTA linker (Biotin nitrilotriacetic acid, VWR 90074) was resuspended in 100mM HEPES pH 8 to a stock concentration of 20μM and stored at 4°C.

1ml of 1nM liposomes of 1 micron diameter is approximately 6.71E+10 liposomes^58^, which is approximately 0.003 picomoles of liposome particles in 30μl sample. To allow an approximate excess of 1000 streptavidin particles per liposome, each 30μl sample needs to contain 3 picomoles of streptavidin, giving 0.1μM concentration of streptavidin. The biotin-NTA linker was used in 4x excess (to account for most of the biotin linker being bound to streptavidin, which has 4 binding sites for biotin), resulting in 0.4μM of biotin linker in each sample. For each 30μl synthetic cell liposome sample, 0.6μl of 5μM streptavidin stock and 0.6μl of 20μM biotin linker stock were used. The biotin linker was pre-incubated with streptavidin at 30°C for 10 minutes.

For all division experiments, synthetic cells were prepared as for the cell cycle experiments (methods section 11), but without the generation counter oligos.

For each division experiment with synthetic cells free in solution (not immobilized), 0μl synthetic cells were mixed with 1.2μl of the pre-incubated streptavidin – biotin linker solution. Samples were incubated for 12h at 30°C with gentle shaking (in PCR tube on a shaker inside incubator). After incubation, the cells were diluted to 0.1mM total lipid concentration in PBS and analyzed using dynamic light scattering measurements (methods section 19).

For each division experiment with immobilized synthetic cells, 5mg beads with immobilized 0μl synthetic cells (after azide blocking) were mixed with 1.2μl of the pre-incubated streptavidin – biotin linker solution. Samples were incubated for 12h at 30°C with gentle shaking (in PCR tube on a shaker inside incubator).

After incubation, magnetic beads were pulled down (using magnetic PCR plate), and the solution from above beads was gently removed. The beads were washed with 50μL of 100mM HEPES pH 8, and the wash solution was combined with the initially removed solution, creating the “daughter cells” fraction. The liposomes in this combined “daughter cells” fraction were lysed with 0.1% Triton-X 100 (Acros Organics, AC215682500) for 10 minutes, lyophilized (to concentrate), and resuspended in μl water.

The magnetic beads with remaining cells still attached were resuspended in 100μL of 100mM HEPES pH 8 and liposomes were lysed with 0.1% Triton-X 100 for 10 minutes. The lysed liposome solution was removed from the beads, lyophilized (to concentrate), and resuspended in μl water. The “free off beads daughter fraction” and “attached to beads fraction” solutions were analyzed using qPCR (methods section 14).

In competitive growth experiments, two populations of synthetic cells were mixed: each population on 5mg beads with immobilized 0μl synthetic cells, the cells were identical except one containing T7Max αHL gene and the other T7 αHL gene. The competitive division samples were incubated and processed as described above, except all wash and resuspension volumes were doubled.

Two plasmids were used as unique DNA markers to track the T7 and T7Max populations in samples where those two types of cells were mixed for competitive division. To avoid adding extra metabolic burden, we chose two plasmids that did not express in bacterial system: a GFP plasmid and mCherry plasmid, both under control of a mammalian CMV promoter. Those plasmids were used here only to mark otherwise identical synthetic cells, since T7 and T7Max promoters are similar enough that it’s difficult to reliably differentiate them with qPCR primers. Both marker plasmids were encapsulated in synthetic cells at 10nM, the CMV-GFP plasmid in T7 cells and the CMV-mCherry plasmid in T7Max cells.

After competitive growth experiments, “daughter fraction” was removed and samples were processed as described above.

## 22. Genetically encoded growth and division

Synthetic cells were prepared with as described earlier (methods section 4) with complete genome, plus 10nM FLAG-tagged αHL, and with 1mol% azido lipid for immobilization (methods section 20). The feeder liposomes were prepared as described earlier, with Ni-NTA and pREP reaction (methods section 6).

Biotin conjugated FLAG tag antibody (Abcam, ab173832) manufacturer stock of 1mg/mL (28μM, assuming 35kDa molecular weight) was diluted to working stock of 20μM and stored at 4°C.

Similar to His-tag based division experiments (methods section 21), for the FLAG tag based division we used 0.1μM final concentration of streptavidin and 0.4μM of biotin FLAG antibody linker in each sample. For each 0μl synthetic cell liposome sample, 0.6μl of μM streptavidin stock and 0.6μl of 20μM biotin FLAG antibody linker stock were used. The antibody linker was pre-incubated with streptavidin at 30°C for 10 minutes.

Feeder liposomes were prepared as previously described (methods section 6). 5mg beads with immobilized 0μl synthetic cells (after azide blocking) were mixed with 1.2μl of the pre-incubated streptavidin – biotin linker solution and 0μl feeder liposomes. In those experiments, the immobilized “parent” synthetic cells carried only generation counter “top” oligos gen1, gen2 and gen, and feeder liposomes carried generation counter “bottom” oligos gen1, gen2 and gen. The presence of full-length generation counter oligo was detected using primers for gen3 (see methods section 12). Despite this being single generation experiment, 3 generation counter oligos were used, to allow formation of a product long enough for unambiguous detection.

Samples were incubated for 12h at 30°C with gentle shaking (in PCR tube on a shaker inside incubator). After incubation, magnetic beads were pulled down (using magnetic PCR plate), and the solution from above beads was gently removed. The beads were washed with 30μL of 100mM HEPES pH 8, and the wash solution was combined with the initially removed solution. The liposomes in this solution were lysed with 0.1% Triton-X 100 (Acros Organics, AC215682500) for 10 minutes, lyophilized (to concentrate), and resuspended in μl water. The magnetic beads with remaining cells still attached were resuspended in 50μL of 100mM HEPES pH 8 and liposomes were lysed with 0.1% Triton-X 100 for 10 minutes. The lysed liposome solution was removed from the beads, lyophilized (to concentrate), and resuspended in μl water. The “free off beads” and “attached to beads” solutions were analyzed using qPCR (methods section 14).

## 23. Single cell plasmid and generation counter product abundance

To estimate distribution of plasmids in daughter cells after growth and division, we first dialyzed the cells, to remove feeder liposomes (see Methods section 24). Cells were then diluted to OD 0.2 normalize cell concentration. Then we proceed to dilute the samples to obtain single cell per experiment. This was done at 4°C (in the cold room), working very fast through the sequential dilution protocol, to avoid leakage from synthetic cells as the dilution approaches critical aggregation concentration (CAC).

To achieve highest possible likelihood of single synthetic cell per reaction, we diluted the samples so that theoretical concentration is approximately 1 synthetic cell per 1μl, and we collected 0.8μl of each aliquot for analysis. This means that it is possible many of the aliquots will be empty (containing no cells). This would skew the plasmid abundance analysis. To ensure that only samples with a cell present will be analyzed, we analyzed the samples for presence of 16S RNA, with the assumption that any sample that has ribosomes has a synthetic cell. Since abundance of ribosomes in each cell was significantly higher than abundance of plasmid DNA copies, it is likely that a daughter synthetic cell might not have any copies of the genome, but as long as it has a cytoplasm, it will have ribosomes.

Each putative single cell aliquot sample (384 in total) were used as a template in reverse transcription, followed by qPCR amplification reaction with *E. coli* 16S RNA primers.^78^

Each initial 16S test 10μl reverse transcription reaction was prepared mixing of 0.8μl of the putative single cell sample, 0. μl of 2μM primer (forward 16S primer, see **Table S2** for all the primer sequences), 0. μl of 10mM dNTPs (Denville Scientific CB4420-2), and 4.7μl water. This was incubated at 6 °C for 1 minute, followed by at least minutes on ice. Then 2μl of x Superscript IV RT buffer, 0. μl of 100mM DTT, 0. μl of 200U/μl of Superscript IV RT (Invitrogen 18090010), and 0. μl 40U/μl RNase Inhibitor Murine (NEB M0314S) were added. Samples were incubated at 52°C for 10 minutes, followed by 80°C for 10 minutes. A 20μl OneTaq PCR was set up with NEB OneTaq 2X Master Mix with Standard Buffer (NEB M0482) with 2μl of the RT sample as template (saving the remaining 8μl in −80°C) and following the NEB recommended protocol with the inclusion of 1x of Chai Green Dye (Chai R01200S). qPCR was read and analyzed on either a Chai Open qPCR machine (single or dual channel) or CFX Opus 384 System.

Of the 384 aliquots initially analyzed, 16S RNA was detected in 204 samples. We have selected 95 samples for further processing, selecting the samples with lowest abundance of 16S RNA (further maximizing chances that those samples contained only one and not more cells).

This is the only experiment in this project where some samples were removed from further processing and analysis based on initial screening.

The 95 samples selected for further processing were used as templates in a multiplex PCR reaction, to amplify the DNA of all 7 plasmids. Following primers were used for detection of polycistronic plasmids: for pLD1 we used primers for HisRS, for pLD2 we used primers for AlaRS, and for pLD3 we used primers for ProRS. A master mix was prepared with all forward and reverse primers, with concentration of 1μM each primer. Each multiplex PCR reaction was prepared using 3.75µl primer stock, 5µl Multiplex PCR 5X Master Mix (NEB, M0284), 8µl of the template (the remaining volume from the above described 16S test reactions) and 8.25µl water. The 40 cycles PCR reaction was performed according to manufacturer’s protocol. After this amplification, the PCR product was used as a template in 7 individual qPCR reactions, one for each plasmid DNA, performed according to the above-described general qPCR protocol (see methods section 14). The results are on **Figure S32** and **Figure S33** and **Figure S37**.

## 24. Dialysis of feeder liposomes

Synthetic cells and feeder liposomes were prepared, and cell cycle experiments were performed, as described earlier (methods section 11). After each 12h generation incubation, samples were dialyzed for 4h against 1μm filter to remove free feeder liposomes.

A liposome dialyzer was prepared by modifying 0.5ml Thermo Scientific Slide-A-Lyzer MINI dialysis devices (similarly to the procedure described earlier^79^ for an older style Slide-A-Lyzer cassettes). The dialysis membrane in each device was replaced with a 1μm Polycarbonate (PCTE) Membrane Filters (Sterlitech, PCT1029320).

Sample of synthetic cells with feeder vesicles after 12h generation incubation was placed in the modified dialysis device, and the bottom tube was filled with 100mM HEPES pH 8. Samples were dialyzed for 4h, then removed from the dialyzer device, extruded (as described in methods section 11) and next generation was started by addition of next population of feeder liposomes.

The 4h dialysis time was chosen based on a calibration experiment, where large 2-3μm liposomes were mixed with small 0.4μm liposomes (large modeling synthetic cells and small liposomes modeling feeder liposomes).

The small “feeder” 0.4μm liposomes were prepared with 10nM GFP plasmid (no transcription and translation system, just plasmid for quantitative detection). The large liposomes were prepared to model “parent” synthetic cells in the growth and feeding experiments, with average diameter of 2-3μm, those liposomes were filled with 10nM αHL plasmid (no transcription and translation system, just plasmid for quantitative detection). Each sample was prepared by mixing 30μl of 1mM lipid liposomes of the small and large type each, gently mixing by flipping the tube, and placing the solution in a dialyzer.

The samples were dialyzed for up to 6 hours. After each hour, a sample was removed from the dialyzer device, lysed with 0.1% Triton-X 100 (Acros Organics, AC215682500), lyophilized (to concentrate) and resuspended in 10μl water. Those samples were analyzed using qPCR (see methods section 14) to quantify amount of αHL plasmid (proxy for concentration of the large synthetic cell liposomes) and amount of GFP plasmid (proxy for concentration of the small model feeder liposomes). Results of this calibration are on **Figure S26**.

## 25. Western blot

Samples for Western blot analysis of protein expression in solution (without vesicles) were mixed directly with loading buffer, liposome samples were first lysed with 0.1% Triton X-100. Samples were mixed 1:1 with 2X SDS loading buffer (100mM Tris HCl, 2.5% SDS, 20% Glycerol, 4% Beta - mercaptoethanol, 0.1% bromophenol blue). Samples in loading buffer were incubated at 95°C for 5 minutes (in a thermocycler), and then loaded on a 37.5:1 Acrylamide:Bis-Acrylamide SDS-Page gel in a Mini-PROTEAN tank (Bio-Rad) according to the manufacturer’s protocol, with Bio-Rad Power Pac 3000 power supply. Gels were run for 60 minutes at 100V in 800 mL of 1X SDS running buffer (25mM Tris, 192mM Glycine, 3.5mM SDS). Gels were then transferred to a 0.2μm nitrocellulose membrane. Transfer was done for 60 minutes at 100V in 1L of 1X transfer buffer (25mM Tris, 192mM Glycine).

After transfer, membrane was incubated with 5% nonfat milk in TBST (20mM Tris, pH 7.4, 150mM NaCl, 0.05% tween) for 60 minutes on a horizontal rocker, then mouse IgG1 anti-his primary antibodies (BioLegend, 652505) were added at 1:5000 dilution. The 5% nonfat milk TBST and mouse IgG1 mixture incubated with the membrane for 60 minutes on a horizontal rocker. After incubation with primary antibodies, the membrane was rinsed three times with TBST followed by three 10 min washes in TBST. The membrane was next added to 5% nonfat milk in TBST containing horseradish peroxidase-conjugate goat anti-mouse IgG1 secondary antibodies (Biolegend 405306) diluted at 1:5000 and incubated on a horizontal rocker for 60 minutes. After incubation with secondary antibodies, the membrane was rinsed three times with TBST followed by three 10 minute washes in TBST.

Blots were developed with SuperSignal (Thermo Scientific, 34577) immunoblotting detection system according to manufacturer’s protocols. Blots were imaged using the ChemiDoc MP Imaging System (Bio-Rad) running Image Lab version 5.2.1.

## 26. Summary of statistical analysis

The main figures present data as heat maps or densely packed plots, leaving limited opportunity to illustrate statistical significance calculations. P values were calculated using 2-tailed equal variance T-test with graphical representation of P values following this convention: ns = P > 0.0; * = P ≤ 0.0, ** = P ≤ 0.01, *** = P ≤ 0.001; n/a when results are undetectable and P value was not calculated. Statistical analysis for data presented on Figure 2 is on **Figure S6** and **Figure S7**, and **Figure S13**.

Statistical analysis for data presented on Figure 3 is in **Table S9**.

Statistical analysis for data presented on Figure 4 is in **Table S10**.

Statistical analysis for data presented on Figure 5 is on **Figure S62** and in **Table S11**.

Statistical analysis for data presented on Figure 6 is on **Figure S67** and **Table S12**.

## 27. Size exclusion purification

Size exclusion purification experiments were used to separate the liposomes from unencapsulated solute fractions.

For each purification experiment, a 10ml plastic column (Poly-Prep Chromatography Column, Biorad 7311550) was filled with Sepharose 4B 45-16 μm bead diameter ethanol slurry (Sigma Aldrich 4B200-1L). The ethanol slurry was allowed to settle by gravitational draining of the solvent, and the column was washed with minimum 4 column volumes of 50mM HEPES pH 8.0. Great care as taken to avoid disturbing the beads on top of the column. After washing, the buffer was allowed to drain until the buffer level nearly touched the top of beads, and liposome sample was loaded on top of the column. During loading, the sample was applied in small drops over the entire surface of the top of the column.

The typical loading volumes were from 50 to 100µl, with the minimal loading volume that in our hands produced good purification results being 20µl.

After loading the sample, 50mM HEPES pH 8.0buffer was immediately applied to cover the top of the column, avoiding drying the sample. The column was eluted under gravity, without speeding up the flow of the buffer. Samples were collected into 96-well plate using fraction collector (Gilson FC203B).

Minimum 48 wells were filled during each purification, enough to ensure all liposomes and free solute fraction were thoroughly eluted.

After purification, samples were analyzed using plate reader, and when needed, samples were recovered from the wells, concentrated by lyophilization, and analyzed further.

## Supplementary Information

### Supplementary text

**Table of content of Supplementary text section**

1. His tag mediated liposome fusion leads to growth
2. The Generation Counter
3. mRNA and protein abundance during cell cycle
4. Synthetic cell culture media
5. Genetically encoded division using FLAG system
6. Fusion system efficiency across multiple generations and between cells
7. The multi-partite genome
8. The translation system cytoplasm of synthetic cells
9. The limitations and ways forward

### Note on translation systems used

The only part of the main figure results done with TxTl was data on figure 2, the experiments described in SI text sections 1 and 3 and accompanying SI figures. All other results, shown on main figures 3, 4, 5 and 6 and corresponding SI figures, were done using PURE translation system.

## 1. His tag mediated liposome fusion leads to growth

*This supplementary text section has information related to section “Genetically encoded liposome fusion leads to growth” of the main paper results, and data on* Figure 2.

The major compositional difference between the peptide and nucleic acid fusion systems implies that any mechanism that can increase binding affinity of the liposome leaflets will induce membrane fusion. We decided to use another well characterized molecular interaction system to bring two liposomes together, the Ni-NTA interaction with a His tag. We hypothesized that if one population of synthetic cell liposomes contains Ni-NTA lipids, and the other population of liposomes is decorated with histidine tags, the Ni-NTA/His tag interaction will bring the outer leaflets of liposomes together, initiating fusion.

To verify that theory, we prepared a VAMP2-His fusion peptide based on the previously used VAMP2 SNARE mimic fusion system^75^. The VAMP2 transmembrane region was synthesized with a 6xHis tag on the N terminus (Methods section 8), and a population of liposomes was decorated with this VAMP2-His protein.

The initial ratio of protein to lipid was 1 to 500 with 1mM liposomes carrying 2µM VAMP2-His. Another population of liposomes was decorated with 18:1 DGS-Ni-NTA lipid at an initial concentration of 0.2 mol%. The VAMP liposomes contain T7 RNA polymerase and a complete TxTl reaction mix, while the Ni-NTA liposomes contain a plasmid encoding GFP under the control of a T7 promoter and a TxTl reaction mix without T7 RNA polymerase.

This design is such that GFP is expressed only if the lumens of those two liposome populations combine (so the GFP plasmid from one liposome population can be transcribed by T7 RNAP from the other population). The membrane of the VAMP liposomes is labelled with a pair of FRET dyes (Cy5 and Cy7) to follow membrane growth. This red-shifted FRET pair (instead of NBD – rhodamine commonly used in liposome growth experiments) was chosen to allow for the monitoring of membrane growth via FRET without spectral interference from the GFP expression used to track lumen mixing. The schematic of that experiment is on **Figure S17**.

In all experiments, we mixed an equal amount of both liposome populations, incubated for 12 hours at 30°C, and then recorded the fluorescence of GFP and Cy5. Higher GFP fluorescence indicates more lumen mixing, while higher Cy5 fluorescence indicates more membrane mixing (since Cy5 is the FRET donor, and diluting the FRET lipids in the newly mixed membrane increases donor fluorescence, while decreasing the acceptor fluorescence, see **Figure S14**).

Upon mixing the liposomes at initial concentrations of 0.2mol% of peptide to lipid and 0.2mol% of Ni-NTA to lipid, we observed GFP production and FRET signal indicating membrane mixing. Encouraged by those results, we investigated both VAMP and Ni-NTA concentration in their respective membranes.

Those experiments revealed that increasing peptide concentration results in more liposome fusion, while increasing Ni-NTA concentration improves fusion only up to about 1mol% of Ni-NTA (higher than 1mol% decreases the fusion efficiency). (**Figure S17**).

We investigated the stability of liposomes with an increased amount of Ni-NTA by testing the retention of calcein, a small fluorescent dye, in liposomes prepared with varying amounts of Ni-NTA. The samples with a higher concentration of Ni-NTA showed an increase in leakage of calcein, independent of the concentration of the VAMP protein in the membrane (**Figure S1**). This indicates that at higher concentrations, the Ni-NTA causes a decrease in the stability of liposomes, despite improved fusion performance. Overall, the VAMP-His results indicate that the His tag and Ni-NTA lipid pair can be used to induce effective fusion of synthetic cell liposomes.

Encouraged by those results, we proceeded to engineering a fusion system that relied on a protein expressed inside synthetic cells, instead of a protein added from the outside after liposome formation. While the VAMP-His fusion could be synthesized inside synthetic cell liposome, that protein doesn’t translocate across the membrane. Therefore, it would be difficult to design a VAMP based mechanism that would couple protein expression inside the synthetic cell with the His tag displayed on the outer membrane. To achieve the coupling of gene expression with membrane tag display, we used modification of αHL membrane protein.

After characterizing the αHL based fusion system (see main results section “Genetically encoded liposome fusion leads to growth”), we confirmed that the αHL can also be synthesized outside of synthetic cells. The αHL protein has two His tags, C-terminal tag and a tag in the trans-membrane loop, so there should be a His tag presented on either side of the membrane.

We set up fusion experiments with one population of synthetic cells carrying T7RNAP, and the other population carrying GFP plasmid and Ni-NTA tags, but this time the external solution also contained complete TxTl reaction and αHL plasmid. The fusion experiments were performed, testing different amount of αHL plasmid and different amount of Ni-NTA lipid tags (**Figure S20**). Both GFP monitored lumen fusion and Cy5 FRET monitored membrane mixing results show that fusion is dependent on both αHL concentration and the concentration of Ni-NTA lipid tag. The results show similar trends to the fusion results with αHL expressed inside the synthetic cell (Figure 2b and **2c**).

## 2. The Generation Counter

*This supplementary text section has information related to section “Five generations of synthetic cell cycle” of the main paper results, and data on* Figure 3.

Demonstrating synthetic cells successfully undergo all five cycles of feeding, growth, and division, is critical to demonstrating a full cell cycle. In previous work on multiple rounds of synthetic cell fusion, we demonstrated that particular cell participated in all fusion events using a complex, multicomponent genetic circuit. That circuit only produced a protein readout if all 10 genes from 9 fusion events were present^34^. Those previous results made us optimistic about the possibility of achieving fusion levels adequate to create a continuous lineage. However, the use of a multi-component reporter pathway was not practical in the cell cycle system. We needed a simpler generation counter. We also rejected any DNA-based designs, to avoid interference with the Phi29 gene replication, discussed in detail later.

We chose an RNA design based on short double stranded RNA oligos with overlapping “sticky ends” (**Figure S45**). The oligos are ligated by a T4 RNA ligase, demonstrated previously to be capable of ligating the “sticky ends” of RNA^80^. The design of overlapping overhangs ensures that the oligos can only ligate in one order, producing a specific ligation product only if all generation counter oligo pairs are present (**Table S3**). The ligation products are detected in an RT qPCR reaction, with the RT primers designed to bind at the end of the third, fourth, or fifth generation oligo product. Generation one and two primers were not used because the product would be too short for the primers to bind without overlap, and too small for efficient qPCR detection. Based on which primers are used (Gen3, 4, or 5), the RT qPCR analysis detects if the generation counter RNA underwent at least that number of ligation events, and thus can determine if the product is the result of at least that many generations. For example, if the generation four (Gen4) primer is used, a product will be detected if four or five fragments were ligated (the product of five ligations also contains the product of four ligations, and so the Gen4 primer will also bind the generation five product) (**Figure S45**).

We first tested the proof of principle of the generation counter, demonstrating that the full-length ligation product can be detected only if all 10 fragments are present. The “top” and “bottom” single stranded oligonucleotides are used to make each double stranded oligos for five generations, for a total of 10 RNA oligonucleotides (**Figure S46**). RT qPCR was performed with Gen5 primers to detect the full-length product. We present the data as fluorescence from the qPCR reaction at round 20 instead of typical Cq values, because samples missing one oligonucleotide lack Cq callouts. We find that showing raw fluorescence is more informative than showing results of 0 for most samples. Only samples where all oligos were present produced measurable qPCR signal, indicating that no product is made unless all 10 oligos (counters for all 5 generations) are present. (**Figure S46**)

Next, we tested the generation counter in liposome fusion experiments. We used two populations of liposomes, with one fusion event instead of five, to first test the generation counter ligation independently from the efficiency of multiple fusion events. One population carried the T4 RNA ligase and the bottom oligos for all five generation counter pairs, while the other population carried the top strand oligos for one, two, three, four, or five generations. Negative controls included a population with no top strands and one with all top strands but no liposome fusion. All samples were also labeled with an NBD – Rhodamine FRET pair, as an independent means to verify membrane fusion. NBD is the donor in the NBD – Rhodamine FRET pair, so successful membrane mixing is indicated by an increase in NBD fluorescence, and a decrease in FRET (**Figure S14**). In all samples where liposome fusion occurred, NBD fluorescence was higher, indicating membrane mixing, while NBD fluorescence remained low in the no fusion control sample (**Figure S47**). After NBD fluorescence was measured, samples were processed for RT qPCR generation counter analysis. Each sample was used for three RT reactions, with primers for generation three, generation four, or generation five (Gen3, Gen4 and Gen5). The qPCR analysis showed no detectable generation counter fusion product in the samples with: no top oligos, no fusion, and in samples with top oligos for only one or two generations. The generation three primer detected ligation product in samples with top oligos for gen 1, 2 and 3 (as well as samples with the oligos for gen 4 and 5). Similarly, the generation four primer detected product in samples with top oligos for generations one through four, and in samples with the top oligos for all five generations. The generation five primer only detected ligation product in samples with all five generation top oligos (**Figure S47**). Those results are consistent with the design of the generation counter, and demonstrate that our generation counter can be used to detect up to five liposome fusion events (a.k.a. generations). **Thus, we developed a generation counter, to confirm the presence of daughter cells that underwent all cycles of growth and division (as opposed to cells that skipped one or more cycles of feeding).**

Herein, we will use the term “generation” to refer to a single cycle of cell growth, followed by incubation and division.

## 3. mRNA and protein abundance during cell cycle

The abundance of mRNA did not significantly correlate with the presence or absence of Phi29 or fusion. Both GFP and αHL mRNA were detected in all samples (Figure 3b, f, j). We speculate that the mRNA abundance was influenced by a few factors that might have opposing effects. Feeding is expected to increase mRNA abundance, by providing more substrate for transcription; however, the Phi29 might have a negative effect on mRNA abundance, by decreasing RNA polymerase access to the DNA template (since T7RNAP and Phi29 compete for DNA occupancy). At the same time, Phi29 replication might also have a positive effect on mRNA abundance, by increasing the total DNA available for transcription.

Future work on characterizing this cell cycle might help elucidate those emergent regulatory mechanisms.

GFP fluorescence presented clearer trends: in samples with DNA replication and feeding, GFP expression increases throughout subsequent generations. In samples with feeding and growth but no genome replication, and in samples without feeding but with genome replication, the GFP expression levels remain similar throughout the experiment (Figure 3d, h, l). This might indicate that our speculative interpretation of mRNA abundance data was correct; feeding and Phi29 replication have both positive and negative effects on mRNA abundance and corresponding protein expression. The GFP expression in samples without Phi29 and without feeding (i.e., no growth and no genome replication) was significantly lower than in all other cases. While this mechanism will require further detailed studies, it might provide an unexpected self-regulating benefit for engineering of a synthetic cell cycle with larger and more complex genomes.

## 4. Synthetic cell culture media

The presence of αHL, while necessary for liposome growth in our system, also causes leakage of synthetic cell components. We tested varying compositions of the external buffer, optimizing the concentrations of amino acids, salts, and energy molecules for gene expression in synthetic cells with αHL channels (**Figure S15**).

In all experiments, 1mM of total lipids synthetic cell liposomes were prepared with 3 nM of both GFP and αHL plasmids. The internal solution was the standard 1x TxTl mix, as described in Methods section 10, and the external solution was prepared as indicated on the figure, with varying amounts of salt, amino acid mix, and energy mix.

All GFP expression values were compared to a negative control experiment with an mCherry plasmid in place of the αHL plasmid, to preserve the multi-plasmid metabolic load on the system. The highest single increase in GFP expression was from an increased energy mix concentration; however, increasing any component of the external mixture resulted in increased GFP expression. Samples with a concentration of any one or more of the tested components below the 1x value for standard TxTl, resulted in less GFP expression than a control sample kept at 1x TxTl via the absence of αHL channels (**Figure S15**).

Overall, keeping the external solution of synthetic cell culture medium, at small molecule concentrations equal to the standard concentration for the protein expression reaction (1x), results in the best performance of the synthetic cell translation system in the presence of αHL membrane channels.

### Nutrient uptake and waste removal

The αHL is a non-specific membrane pore, allowing passage of all molecules under its molecular weight size cut-off. The equilibration through the channel is concentration gradient driven, it’s not active or selective transport. For that reason, the channel serves both for nutrient uptake and for removal of waste products, equilibrating the small molecules from the inside of the synthetic cell with the outside buffer. The volume of the outside buffer is significantly larger than the volume of the lumen of the cells.

## 5. Genetically encoded division using FLAG system

*This supplementary text section has information related to section “Genetically encoded division” of the main paper results, and data on* Figure 6.

The genetically encoded growth requires small fusogenic tag that can be embedded in the feeder liposomes, to facilitate their fusion with the synthetic cell. His tag with its small molecule binding partner of Ni-NTA is the only system we know that fits all required parameters: a small, not terminal peptide tag (so it can be incorporated in trans-membrane loop of αHL and presented on the surface of synthetic cells), and a small binding partner (nickel NTA complex) that can be embedded in the membrane of feeder liposomes by lipid conjugation. All other peptide-based interactions require at least one of the binding partners to be significantly larger. Based on our experience with different fusogenic peptide systems, large binding partner would inhibit fusion of lipids. Therefore, we concluded that His tag system is the only currently available system for genetically encoded uptake of feeder liposomes in our system.

To combine genetically encoded growth with genetically encoded division, we needed to engineer another system for immobilizing the division inducing large protein on the surface of synthetic cells (to keep the His tag free to facilitate fusion with feeder liposomes). We chose a FLAG tag, with its binding partner of FLAG antibody. FLAG tag antibody would likely be too big to induce membrane fusion, but since for division the goal is to attach a large protein to the surface of the liposomes anyway, we speculated that FLAG antibody mediated streptavidin attachment will be effective in inducing division. The results are similar to the His system, as demonstrated by DLS measurements (Figure 6c). We also confirmed division in the bead immobilization division experiments, similar to the His system tests shown on Figure 6f and **6g**. Synthetic cells expressing 10nM αHL FLAG plasmid were prepared in the same way as for the His αHL division experiments, immobilizing the cells on magnetic beads with click chemistry. Streptavidin and biotin conjugated anti-FLAG antibody were added to the cells to induce division (see methods section 22). After 12h incubation, cells still attached to beads were separated from “daughter” cells free off the beads, and both fractions were analyzed for the presence of αHL DNA.

The αHL DNA was detected, as expected, in all samples of cells still attached to beads that had αHL to begin with (**Figure S70** panel **c**). In the free off the beads “daughter cell” fractions, αHL DNA was detected only in the sample with all elements necessary for division (αHL, streptavidin and biotin conjugated FLAG antibody linker). This confirmed that only if the cells divide, the daughter cells can detach from the beads. The amount of αHL DNA detected in daughter cells free off the beads is similar in the His division scheme (Figure 6f) and in the FLAG tag division scheme (**Figure S70** panel **b**). This confirmed that FLAG tag division scheme works with similar efficiency as the His tag division scheme.

Next, we tested the FLAG based division system in a competitive division scenario. Similarly to experiments with the His tag based division: two populations of synthetic cells were mixed. Synthetic cells containing FLAG tagged αHL under “slow” promoter T7 and FLAG tagged αHL under “fast” promoter T7Max were dividing in the same reaction. After division, we collected the “daughter” cells free off beads and the “parent” cells remaining attached to beads. Both fractions were analyzed for the presence of a unique DNA marker (see methods section 21) indicating presence of either 7T population or the T7Max population. The αHL DNA was detected, as expected, in all “parent” samples still attached to the beads that initially contained either T7 or T7Max cells, or both (**Figure S70** panel **E**).

In the samples free off the beads, the “daughter” fractions, αHL DNA was only detected if the division occurred (**Figure S70** panel **d**). Similarly to the results of His tag-based division competition (Figure 6g and **6h**), in the FLAG tag-based division experiments the cells with T7Max αHL produced more detectable “daughter cells” – as expected, the faster T7Max population divided more.

Those tests confirmed that FLAG tag based genetically encoded division system is as viable as the His tagged based system. Having two different tags, His and FLAG, enabled us engineering of synthetic cells capable of genetically encoded feeding, growth and division. The feeding and growth system used the His tag with Ni-NTA linker, and the division system used the FLAG tag with biotin conjugated FLAG antibody linker.

### The His and the FLAG division systems do not interfere with each other

The His division tag doesn’t interfere with FLAG based division, and the FLAG division tag doesn’t interfere with the His based division. We tested two things: cross-reactivity of the tags, and interference of the opposite system tag with division. All results are on **Figure S74**.

To test cross-reactivity, we tested whether cells with αHL-FLAG can undergo division in presence of Ni-NTA linker (no division was observed), and whether cells with αHL-His can undergo division in presence of biotin FLAG antibody linker (also no division was observed).

Then, we tested whether adding the tag of the opposite division system to a matching pair (so adding biotin FLAG antibody linker to system containing αHL-His and Ni-NTA linker, and adding Ni-NTA linker to a system containing αHL-FLAG and biotin FLAG antibody linker) would inhibit division. No interference was noted.

## 6. Fusion system efficiency across multiple generations and between cells

*This supplementary text section has information related to Ni-NTA His tag fusion system used through this paper*.

### Fusion across multiple generations

In each experiment of the multi-generational cell cycle, the αHL concentration is renewed through protein expression, while the Ni-NTA linker concentration is renewed by tags coming in with each portion of feeder liposomes. Therefore, we speculated the efficiency of the fusion after each generation will only be affected by the concentration of αHL plasmid (how many copies of αHL make it into each daughter cell generation). To test this, we set up a fusion efficiency testing experiment similar to the schematic described on Figure 2a, but this time the GFP plasmid was brought in only in feeder liposomes during one generation. The starting synthetic cells contained T7 RNA polymerase and TxTl machinery expressing αHL. Generation of growth and division experiments were performed, as described on Figure 3 and in Methods section 11. The “feeder vesicles” were being added, containing all metabolites as described in Methods section 11, except no generation counter oligos were used here. Instead of generation counter oligo system, we used GFP reconstitution: at one of the generations, the feeder vesicles bring in GFP plasmid under the control of a T7 promoter. After fusion, the T7 RNA polymerase from the synthetic cells enables transcription and expression of GFP from a plasmid brought in by feeder liposome.

As we demonstrated during optimization of the fusion system (data presented on Figure 2 and accompanying supporting materials), the fusion efficiency correlates with the GFP expression yield. Since in the experiments described here the GFP plasmid is brought in by feeder liposomes, quantifying GFP expression after 12h incubation following the addition of feeder liposomes allows us to estimate fusion efficiency. However, comparing GFP expression directly between generations was not informative, because after each generation of growth and division the total amount of cells in the sample increases, so more total protein can be made.

To ensure that GFP expression values are correlated with efficiency of fusion, we needed to normalize cell density in the samples before fusion. To do that, we used a membrane dye. All samples (starting synthetic cells, and all feeder liposomes in generations leading to the experiment) contained 0.01mol% Rhodamine dye. First, samples were dialyzed to remove previous generations feeder liposomes (see Methods section 24). Then, samples were normalized to equal Rhodamine fluorescent value per volume (10uL sample at 5000 RFU). Then, feeder vesicles with GFP plasmid were added, samples were incubated for 12h and GFP fluorescence was measured. Since starting concentrations of synthetic cells was the same regardless of which generation the experiment was performed at, the resulting GFP signal from the sample can be directly compared, to indicate fusion efficiency.

Results are presented on **Figure S36**. The data indicates that, as expected based on previous αHL titrations, fusion efficiency somewhat decreases with each generation. This is most likely because it correlates with αHL plasmid abundance. As indicated on Figure 3p, αHL plasmid is in 71% cells analyzed after 5 generations, making it the most abundant plasmid. Both the GFP fusion efficiency data described here, and plasmid abundance data presented earlier, can be explained by this correlation of αHL abundance with fusion efficiency: the plasmid that is responsible for fusion efficiency will have highest likelihood of persisting through the generations, because the daughter cells which lost αHL will have lower chance of growing in the next round (some αHL protein will persist in the membrane, so fusion is not expected to completely stop, and some αHL mRNA is likely to persist even if the daughter cell did not receive the αHL plasmid).

### Fusion between different cells

A separate issue potentially affecting fusion efficiency is the possibility of fusion between two different synthetic cells – as Ni-NTA tags accumulate during the feeding events. This is distinct from the desirable fusion between a synthetic cell and feeding vesicles.

To test that, we prepared two populations of synthetic cells, and carried out 5 generations of growth and division. The cells contained a genome comprised of T7RNAP, αHL. One population of cells carried GFP plasmid under control of T3 RNA polymerase promoter, while the other population contained T3 RNA polymerase gene under control of T7 promoter. The cells containing T3 plasmid were expressing T3 RNAP during each incubation step (because they contained T7 RNAP needed to express the T3 RNAP), while the cells that contained GFP plasmid under T3 promoter were unable to express the GFP. Expression of GFP was only possible if the cells containing T3 RNAP fused with cells containing the GFP plasmid. This is analogous to previously described T7 / GFP plasmid fusion experiments. The T3 promoter / polymerase pair was used here because the T7RNAP was already used for αHL expression (so it would not have been omitted from one population of cells).

The two populations of synthetic cells (one carrying T3RNAP and one carrying GFP plasmid under T3 promoter) were mixed at the start of the experiment, and generations of growth and division experiments were performed as described earlier. After each generation, GFP fluorescence was measured, to monitor possible fusion between synthetic cells (not fusion between synthetic cell and feeder liposome).

The results are shown on **Figure S38**, along with positive control of GFP fluorescence from GFP under T7 RNAP in synthetic cells that did undergo all 5 cycles of growth and division.

## 7. The multi-part genome

*This supplementary text section discusses the design of the multi-plasmid genome*.

In this work, we chose to encode the entire genome of the synthetic cell on several plasmids, rather than constructing one megaplasmid containing all genes. At first, this decision was driven by practical consideration: designing the cell cycle, we did not know which genes will result in the best performance. Optimizing each element separately was easier when key genes were encoded on smaller plasmids.

After completing the cell cycle experiments, we considered the merits of encoding the entire genome on a single plasmid, but we decided to keep it split between multiple smaller plasmids for both practical and biological considerations. For future engineering of this system, it will be practically easier to modify specific genes, to add or remove components, when the genome remains split. The biological reason for keeping it on independent plasmids is the consideration for partition into daughter cells during division. The fact that genes are divided into multiple, physically separate plasmids, limits the possible influence of stochastic inequalities in plasmid partitioning among daughter cells. Since the gene is encoded on multiple plasmids, and each gene is present at relatively low copy number in each cell (see **Table S8**), during division it is more likely that at least some daughter cells receive at least one copy of each plasmid. If the entire genome was on single plasmid, any accidental fluctuation in genome concentration within the grown cell would potentially have more serious effect on the copy number of all genes in each daughter cell. We speculate that as synthetic cell systems mature and develop further, including development of cytoskeleton (see discussion in section below, Limitations and way forward), the genome will eventually be placed on single large plasmid (aka chromosome). For now, as this is the first demonstration of a complete cell cycle and genetically encoded division, we think more optimization and engineering will be needed to build up robustness of this system. Such engineering will be easier if the genome remains split between multiple plasmids.

### The timing of genome replication and division

In the absence of complex cell cycle control checkpoints, the Phi29 polymerase is active all the time. In the cell cycle with mechanical division, the DNA replication occurs during the incubation time before each division step. In the cells with genetically encoded division, once the streptavidin and division tag (the biotin FLAG antibody linker or the Ni-NTA linker) are added, the division process can begin, and it continues unless tags are removed. In that system the DNA replication occurs concurrently with cell division. In both cases, the daughter cells have been shown to contain newly synthesized DNA (separated from the original DNA used to make first generation of synthetic cells by DpnI digest, see methods section 11).

The quantification of newly synthesized DNA in daughter cells after cell cycle with mechanical division is shown on Figure 3, 5, **S25**, **S28**, and **S32** and **S33**. The quantification of newly synthesized DNA in daughter cells after genetically encoded division is shown on Figure 6, **S67**, **S69**, **S71**, and **S72**.

### The limitations of genome replication

This work uses Phi29 DNA polymerase for replicating of the synthetic cell genome. The advantages of that enzyme are isothermal reaction conditions, activity at the temperature compatible with protein expression, and the fact that active Phi29 enzyme can be expressed *in vitro* (there are no modifications or folding chaperones necessary for activity). However, there are several limitations of this system, including the strand displacement activity, and limited ability to replicate large haploid genomes. In this work, Phi29 was good enough to replicate the relatively small (by the standards of living organisms) multi-partite synthetic genome, but more work is needed to improve synthetic genome replication. To achieve greater robustness of a fully autonomous synthetic cell, a different, better genome replication system will need to be engineered.

### All plasmids are transcribed

The qPCR analysis indicates presence of mRNA from all genes on each plasmid of the multi-partite genome, indicating that all genes (each having its own promoter) are transcribed after 5 generations of cell cycle with growth and division. Results for mRNA abundance for some genes are on Figure 3, and qualitative measurement of presence of all other genes is on **Figure S44**.

### All genes are replicated

We quantified all individual genes from the multi-partite genome after 5 generations of growth and division. Data on Figure 3 and **Figure S28** shows that under conditions enabling genome replication (with Phi29 present), newly synthesized DNA for all genes is detected. Individual data points for that figure are on **Figure S29**.

### Each plasmid is partitioned into daughter cells

Single cell analysis measured abundance of each plasmid after 5 generations of growth and division. **Figure S32** shows abundance of each plasmid in each of the 95 single cell experiments, demonstrating that each plasmid is present in significant number of daughter cells after 5 generations. Summary of that data, showing count by plasmid, is on **Figure S33**.

### The limitations of incomplete partitioning into daughter cells

As shown in data on figures discussed in the paragraph above, not all daughter cells receive full complement of plasmids. This is the main limitation of multi-part genome, one that will hopefully be solved after the genome is optimized and placed on single chromosome. We notice an interesting, although not entirely unexpected pattern: pLD1, pLD2, pLD3 and GFP plasmids are lost at highest rates, while αHL, T7RNAP and pREP are more retained. This could be explained by the relative necessity of those plasmids. Since feeder liposomes bring fresh ribosomes, and GFP has no role in feeding process, losing those plasmids would not affect feeding and mechanical division. Losing membrane channel, RNA polymerase, or DNA polymerase, would probably affect the cell’s ability to grow and express proteins more significantly. However, because PURE system of our synthetic cells has no protein degradation machinery, it is possible the loss of those plasmids in later generations, when the proteins already had a chance to accumulate, might have less detrimental effect.

## 8. The translation system cytoplasm of synthetic cells

*This supplementary text section discusses cell-free translation systems used in this work*.

In engineering liposome encapsulated synthetic cells, the most commonly used liposome lumen solution, a.k.a. the synthetic cell cytoplasm, is a cell-free translation system. Two of such systems are used in this work: the whole-cell lysate TxTl, and the PURE system.

Both TxTl and PURE contain natural ribosomes purified from *E. coli*, together with translation system components like acyltransferases, EF-Tu, and tRNAs. All those components are isolated from *E. coli*.

There are three major differences between TxTl and PURE that were relevant to work presented here: the efficiency, compatibility with genome replication, and the trackability of composition. The TxTl has significantly higher yield than PURE, providing more protein in shorter amount of time, and it is more economical to use. The PURE system, despite lower yields, was used in most experiments presented in this paper, because it is the only translation system currently compatible with genome replication via the Phi29 polymerase system.^18,20^ The PURE system is also the only translation system with a chemically defined composition - all elements of PURE, including proteins, RNA, and small molecules, are separately purified and combined for the reaction.

In a typical protein expression experiment inside synthetic cells used in this work, the expression starts with DNA template transcription via T7 RNA polymerase. The mRNA created in this transcription is immediately used in translation, since our system does not include mRNA splicing and maturation.

Whole-cell lysate TxTl is used on Figure 2 for testing and optimization of the genetically encoded fusion system. We used TxTl to originally optimize the feeding system, since the yields of TxTl reactions are higher and costs of those reactions are lower.

The modified PURE system is used in cell cycle, competition, selection and growth experiments, including Figure 3, 4, **5** and **6**. The PURE system used in this was included some buffer and solutions modifications, described in Methods section 2, pREP reaction. Those modifications, first described in the work using Phi29 to replicate genome in PURE^20^, were necessary to couple transcription, translation and Phi29 mediated genome replication. We are grateful to prof. Hannes Mutschler and dr. Kai Libicher for their generous advice on following their modified PURE protocol necessary for Phi29 reactions.

## 9. The limitations and ways forward

*This supplementary text section expands on the conclusions of the main paper*.

In this work, we present a chemical system made from non-living components, capable of feeding, growth, selection and division. While recognizing this is a significant step forward in the quest to generate living systems from non-living components, we also recognize that significant work remains before synthetic cells are capable of robust independent life and evolution. Below we list the major constraints of the system presented in this paper, and we suggest possible next steps to engineer around or away from those constraints.

### Cellular organization

All natural life has some means of segregating the content of cytoplasm, and most also has means to impart structure into the membrane. This is usually done by cytoskeleton and organelles (including membraneless organelles and cytoplasmic domains). Synthetic cells presented here have no such spatial organization aids. Despite that, we have confirmed that after 5 generations of cell cycle, a significant portion of synthetic cells still contain all 7 plasmids from the 90kbp genome (Figure 3o and Figure 3p).

Future work on this system is needed to investigate genome distribution after the genetically encoded division. We confirmed that all plasmids make it to the daughter cell fraction (Figure 6R), unfortunately relatively low yields of this division mechanism make it difficult to investigate with more granularity.

The abundance of each individual gene in the sample after 5 generations is on **Figure S28**, abundance of each plasmid in single cells is on **Figure S32**, count of cells expressing GFP after each cycle is on **Figure S30**, and abundance of full length generation counter oligo in single cells after each generation is on **Figure S37**.

To achieve more robust, continuous multi-generational replication, the cell cycle and division mechanisms will need to be expanded, to include segregation of cytoplasmic components and the genome. Synthetic cell engineering already provides many technologies that can be utilized for this purpose. There has been a lot of progress in engineering synthetic cell cytoskeleton^40,42^, and in building intracellular structures like membraneless organelles^81–83^. Those tools can be used to segregate genomes into daughter cells, and to ensure proportional distribution of ribosomes and other cytoplasmic components.

Interestingly, while cytoskeleton has been considered essential to replication, partitioning of cellular components, and cell motility, a recent report demonstrates that synthetic cells can achieve chemotaxis, a simple example of motility, without functioning cytoskeleton. ^84^ While this report is not directly related to the work presented in this paper, it provides interesting evidence that a key cellular function can be realized by means other than mimicking a natural system.

### Regeneration of metabolism

It has recently been demonstrated that a PURE system is capable of reconstituting a functional ribosome^56^, and regeneration of metabolic components^55^. Synthetic cells presented in this work have genome containing the proteins necessary to build a translation system (the pLD1, pLD2 and pLD3 plasmids), but we still deliver ribosomes and other elements of the pREP reaction in the feeder liposomes. Those cells are also perfect heterotrophs, with all small molecule substrates being delivered from outside media and via feeding. More work is required to engineer a robust synthetic cell metabolism, capable of self-regeneration of all key enzymes.

### Mutagenesis

Phi29 has reported mutation rates on the order of 10^6^.^85^ In the previous mutation studies, the enzyme is named ‘phi 29 DNA polymerase’, while in this work we use the name Phi29. To the best of our knowledge, it is the same enzyme.

It has been demonstrated that synthetic cells can undergo Darwinian evolution using Phi29 ^66^. In a genome of the size comparable to the one used in our work, for simplicity assuming 100kbp, Phi29 would statistically make one error for every 10 complete replications of the whole genome. While this high-fidelity replication exceeds the fidelity of that found in natural cells by 1-2 orders of magnitude^86^, over enough replication cycles and generations, beneficial mutations could arise and affect the phenotype.

In this work, we introduced mutations artificially, preparing synthetic cells with slow T7 or fast T7Max promoter. While this enabled selection, but not Darwinian evolution – the latter would require that the mutations arise within the cells. More work is needed to engineer this system with built-in evolvability, to enable nature to take over from our curated pool of mutations. There are no mutation repair mechanisms, so most of the spontaneous mutations in promoters and protein coding sequences will likely be lethal to the synthetic cell. However, this might actually be beneficial in the early stages of exploring evolvability of synthetic life. Additionally, synthetic cells contain multiple copies of each gene – they are, using biological term, polyploids.

### Control mechanisms

Gene expression in synthetic cells is not controlled by any sophisticated mechanism. Like in most *in vitro* translation systems, gene expression level depends on the strength of the promoter and the copy number of each gene. For more complex behaviors and robust metabolic resource management, gene expression control mechanisms need to be implemented. Regulation of copy number of genes after replication will also be needed for synthetic cells dividing continuously over unlimited number of generations.

### Biosafety and Biosecurity

Engineering living synthetic cells provides many opportunities to challenge the existing biosafety frameworks, and potentially also creates novel biosecurity hazards. While engineering any pathogenic biological system is, in theory, prohibited under the Biological Weapons Convention, international law remains poorly positioned to deal with regulating emerging fields like ours. The Cartagena Protocols, supplement to the Convention on Biological Diversity, would be better suited to cover this research. Unfortunately, two countries did not ratify the protocols (the United States and the Holy See), and one of those countries is a major center of synthetic cell engineering efforts. Recent conversations about one particular type of synthetic cells, the so-called mirror life of a synthetic cell with opposite stereochemistry^87^, highlights the needs of developing better regulatory framework for governing all synthetic cell technologies.

The synthetic cell system presented in this paper is not robust enough to survive outside of perfect lab conditions. For one, the necessity of adding a large bulky protein and a specialized linker (in this work, streptavidin and either FLAG antibody or Ni-NTA biotin linker) is a nearly ideal auxotrophy: it is difficult to imagine those compounds being available in the environment. Therefore, even if synthetic cells replicating according to the mechanism presented here were to be accidentally or intentionally released, there is no plausible pathway for them to replicate outside of the experimental conditions.

However, this project offers a significant milestone towards evolvability of synthetic cells, making it more likely that more robust, autonomous systems will be available soon. This highlights the urgent need to develop a safety and security framework for future synthetic cell engineering.

### The definition of life

Life is notoriously difficult to define exhaustively and unambiguously. A common question raised in the field of engineering live from non-living components is “How do we know we are done?” The trait most commonly listed as a key property of a living cell is self-replication. However, defining “self” in self-replication remains elusive. One might point out that there is no life that truly is independent of environmental interventions. Synthetic cells engineered in this work need external supply of streptavidin for replication. Most living organisms need external intervention to produce offspring.

Sometimes it’s chemical (as simple as abundance of food), astronomical (mating at specific time of the year), or a mechanical force (certain mycobacteria will divide if poked with a needle ^88^). One might also argue that if self-replication is a trait necessary for life, a human is no alive – none of us can *self*-replicate. This highlights the inadequacy of all existing definitions.

The authors of this paper are partial to a definition quoting Justice Potter Stewart, “I will know it when I see it”.

### Supplementary figures

Each SI figure is labeled, in the upper right corner, with information about the section this data is related to.

For quick reference and finding supplementary figures corresponding to each part of the project, below is a list of SI figures relevant to each main figure of this paper.

Figure 2: SI figures 1-24

Figure 3: SI figures 25-48,

Figure 4: SI figures 49-56

Figure 5: SI figures 57-62

Figure 6: SI figures 63-74

Calibration data for all parts of the project: SI figures 75 and 76

### Table of content of supplementary figures

| SI figure | Title | Related to main figure |
| --- | --- | --- |
| **Figure S1** | Small molecule leakage from liposomes with different NTA and VAMP2 His peptide tag. | Fig. 2 |
| **Figure S2** | Leakage after fusion of NTA lipids with VAMP2 His. | Fig. S1 |
| **Figure S3** | Leakage test for double His labeled αHL fusion experiments. | Fig. 2 |
| **Figure S4** | Leakage test for double His labeled αHL fusion. | Fig. S3 |
| **Figure S5** | FRET data for liposome fusion samples corresponding to the Western Blot analysis of αHL expression | Fig. 2 |
| **Figure S6** | His labeled αHL fusion, GFP data. | Fig. 2 |
| **Figure S7** | His labeled αHL fusion, Cy5 FRET data. | Fig. 2 |
| **Figure S8** | Leakage test for NTA lipids. | Fig. 2 |
| **Figure S9** | Test leakage for αHL expression. | Fig. 2 |
| **Figure S10** | Leakage test for NTA lipids. | Fig. S8 |
| **Figure S11** | Test leakage for αHL expression. | Fig. S9 |
| **Figure S12** | Liposome fusion testing at different liposome ratios. | Fig. 2 |
| **Figure S13** | GFP DNA and mRNA after genetically encoded lumen mixing of synthetic cells | Fig. 2 |
| **Figure S14** | General scheme for liposome FRET size measurement. | Fig. 2 |
| **Figure S15** | Optimization of synthetic cell media | Fig. 2 |
| **Figure S16** | Optimization of synthetic cell media. | Fig. S15 |
| **Figure S17** | VAMP2-His fusion testing. | Fig. 2 |
| **Figure S18** | VAMP2-His fusion testing, GFP. | Fig. S17 |
| **Figure S19** | VAMP2-His fusion testing, Cy5. | Fig. S17 |
| **Figure S20** | Fusion with αHL introduced externally. | Fig. 2 |
| **Figure S21** | Fusion testing with external αHL, GFP signal. | Fig. S20 |
| **Figure S22** | Fusion testing with external αHL, Cy5 signal. | Fig. S20 |
| **Figure S23** | Leakage test for His labeled αHL fusion experiments. | Fig. S20 |
| **Figure S24** | Leakage test for αHL fusion experiments. | Fig. S23 |
| **Figure S25** | Synthetic cells after 5 generations of growth and division with dialysis. | Fig. 3 |
| **Figure S26** | Dialysis time course of removing “feeder” liposomes | Fig. S25 |
| **Figure S27** | Generations of growth and division with dialysis. | Fig. S25 |
| **Figure S28** | All genes in 90kB synthetic cell genome after 5 generations of cell cycle. | Fig. 3 |
| **Figure S29** | Quantification of individual genes in the synthetic cell genome after 5 generations of cell cycle. | Fig. S28 |
| **Figure S30** | GFP fluorescence in synthetic cells | Fig. 3 |
| **Figure S31** | Fluorescence microscopy of synthetic cell samples, imaged after five generations | Fig. 3 |
| **Figure S32** | Abundance of each plasmid after 5 generations of cell cycle. | Fig. 3 |
| **Figure S33** | Summary of single cell analysis of abundance of each plasmid. | Fig. S32 |
| **Figure S34** | Individual data points for 5 generations of growth and division. | Fig. 3 |
| **Figure S35** | Detection of component leakage from cells after 5 generations of growth and division | Fig. 3 |
| **Figure S36** | Fusion efficiency across multiple generations | Fig. 3 |
| **Figure S37** | Full length generation counter oligo after each generation | Fig. 3 |
| **Figure S38** | Investigating fusion between synthetic cells | Fig. 3 |
| **Figure S39** | Cell number changes during cell cycle and replication | Fig. 3 |
| **Figure S40** | Cell cycle without fluorescent proteins | Fig. 3 |
| **Figure S41** | Membrane stability in presence of fluorescent protein GFP | Fig. 3 |
| **Figure S42** | Membrane stability in presence of fluorescent protein mCherry | Fig. 3 |
| **Figure S43** | mRNA and protein time course | Fig. 3 |
| **Figure S44** | Detection of mRNA for each gene after 5 generations of cell cycle | Fig. 3 |
| **Figure S45** | The general scheme of the generation counter. | Fig. 3 |
| **Figure S46** | Detection of the generation counter full-length product. | Fig. S45 |
| **Figure S47** | Generation counter tests in fusion experiments. | Fig. S45 |
| **Figure S48** | Optimizing the ratio of liposome populations and ligase enzyme. | Fig. S47 |
| **Figure S49** | Flow cytometry analysis of fusion reactions, with either T7 αHL or T7Max αHL plasmids. | Fig. 4 |
| **Figure S50** | Flow cytometry analysis of synthetic cells after 5 generations of growth and division. | Fig. 4 |
| **Figure S51** | Flow cytometry analysis of positive and negative control samples after 5 generations of growth and division. | Fig. 4 |
| **Figure S52** | Phi29 replication for T7 αHL and T7Max αHL plasmids. | Fig. 4 |
| **Figure S53** | Amplicon DNA isolated from samples of competing cells after 5 generations. | Fig. 4 |
| **Figure S54** | Example of analysis of NGS data. | Fig. 4 |
| **Figure S55** | Selection experiments with normalized metabolic load. | Fig. 4 |
| **Figure S56** | Calibration curves for GFP and mCherry production after 5 generations of synthetic cell cycle. | Fig. 4 |
| **Figure S57** | Summary of the results of competitive growth | Fig. 5 |
| **Figure S58** | Number of cells containing GFP or mCherry plasmid during the subsequent generations of competition experiments | Fig. 5 |
| **Figure S59** | Abundance of GFP and mCherry plasmid in each individual cell after competitive growth experiments, with 1x feeder liposomes | Fig. 5 |
| **Figure S60** | Abundance of GFP and mCherry plasmid in each individual cell after competitive growth experiments, with 0.1x feeder liposomes | Fig. 5 |
| **Figure S61** | Flow cytometry data from feeding competition experiments | Fig. 5 |
| **Figure S62** | Fluorescence values for mCherry and GFP competitive feeding. | Fig. 5 |
| **Figure S63** | Flow cytometry of synthetic cells before and after division | Fig. 6 |
| **Figure S64** | Size of synthetic cells after genetically encoded division using His tag system. | Fig. 6 |
| **Figure S65** | Size of synthetic cells after genetically encoded division using FLAG tag system | Fig. 6 |
| **Figure S66** | DNA abundance in synthetic cells after genetically encoded division on beads. | Fig. 6 |
| **Figure S67** | DNA abundance in synthetic cells after competitive genetically encoded division on beads. | Fig. 6 |
| **Figure S68** | Individual data points for genetically encoded growth and division. | Fig. 6 |
| **Figure S69** | Detection of each plasmid in the synthetic cell genome after growth and division | Fig. 6 |
| **Figure S70** | Genetically encoded division using FLAG system | Fig. 6 |
| **Figure S71** | DNA abundance in synthetic cells after genetically encoded division on beads with the FLAG tagged αHL | Fig. S70 |
| **Figure S72** | DNA abundance in synthetic cells after competitive genetically encoded division on beads with the FLAG tagged αHL | Fig. S70 |
| **Figure S73** | Fluorescent microscopy of synthetic cell undergoing a cycle of growth and division | Fig. 6 |
| **Figure S74** | Measuring interference and cross-reactivity of His and Flag based division systems | Fig. 6 |
| **Figure S75** | Calibration curves for qPCR experiments. |  |
| **Figure S76** | FRET calibration curves. |  |

---

**Figure S1.**
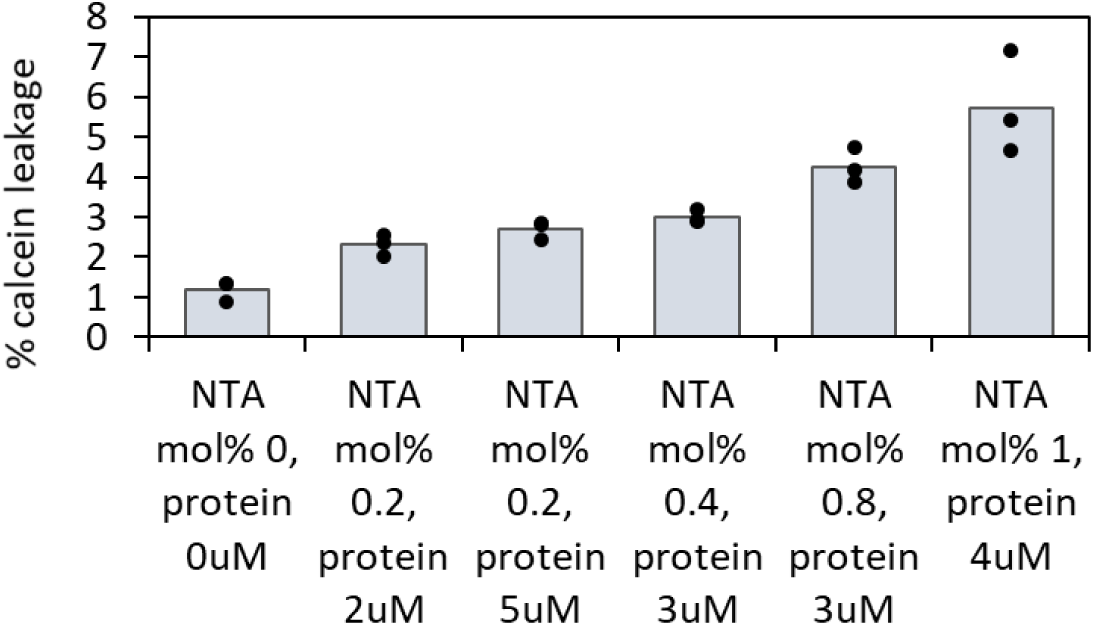
Small molecule leakage from liposomes with different NTA and VAMP2 His peptide tag. Leakage of small molecule calcein was tested in liposomes with NTA lipids and liposomes with VAMP2 His peptide tag (labeled “protein”), after 12h incubation at 0°C. Liposomes contained 1mM calcein at the beginning of the experiment. The dots on top of the bar graphs indicate individual values of three replicates. Examples of individual purification plots for samples summarized on this figure are on **Figure S2**.

**Figure S2.**
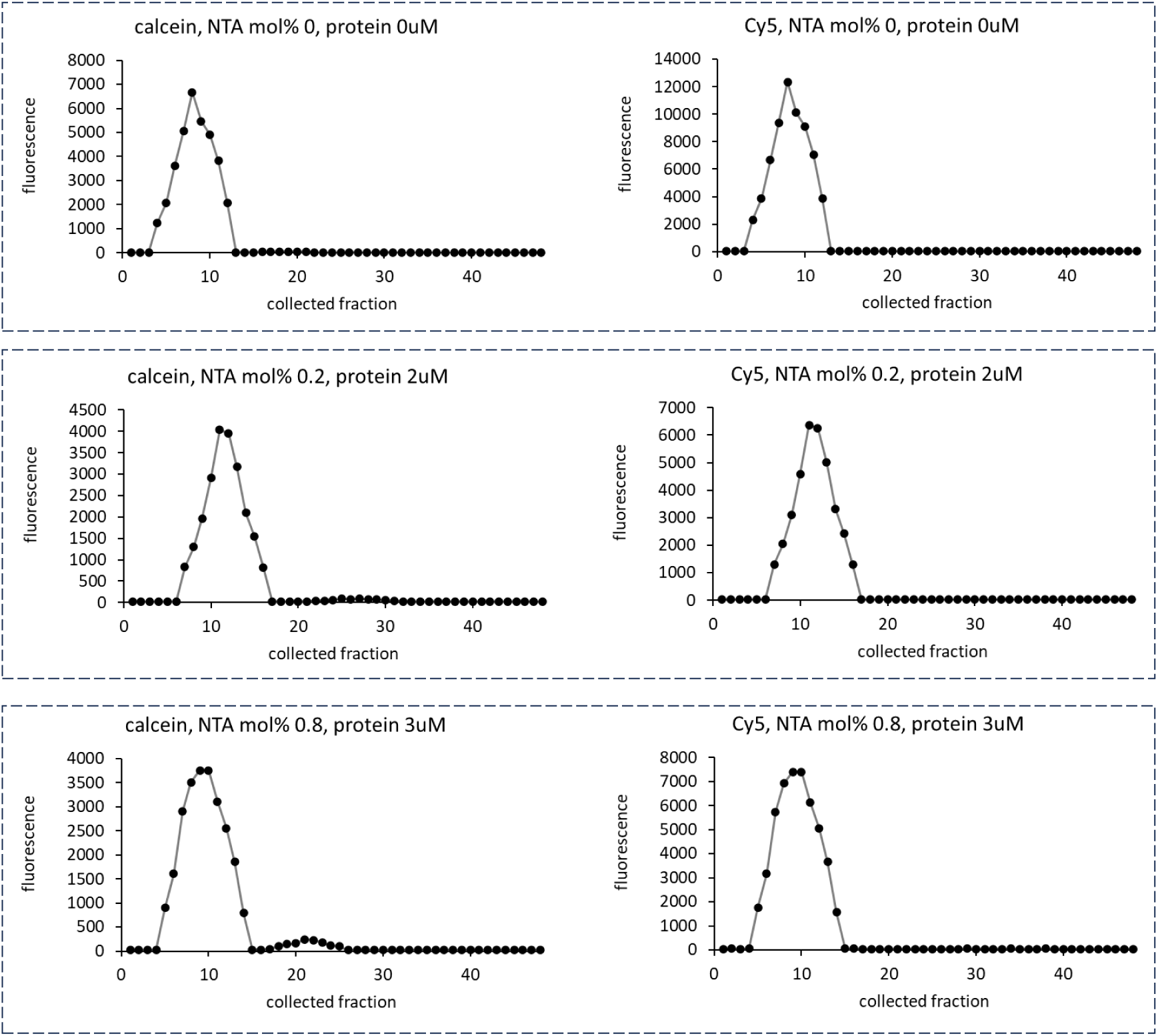
Leakage after fusion of NTA lipids with VAMP2 His. Example plots from size exclusion chromatography purification after fusion of liposomes with NTA lipids and liposomes with VAMP2 His peptide tag (labeled “protein”), after 12h incubation at 0°C. Liposomes contained 1mM calcein at the beginning of the experiment. Each sample was analyzed using two fluorescent measurements: calcein channel and Cy5 channel (see **Table S5** for wavelengths). The calcein channel traces indicate leakage of small molecule and the Cy5 channel (Cy5 lipid dye 1,2-dioleoyl-sn-glycero-3-phosphoethanolamine-N-(Cyanine 5)) indicates which fractions contain liposomes.

**Figure S3.**
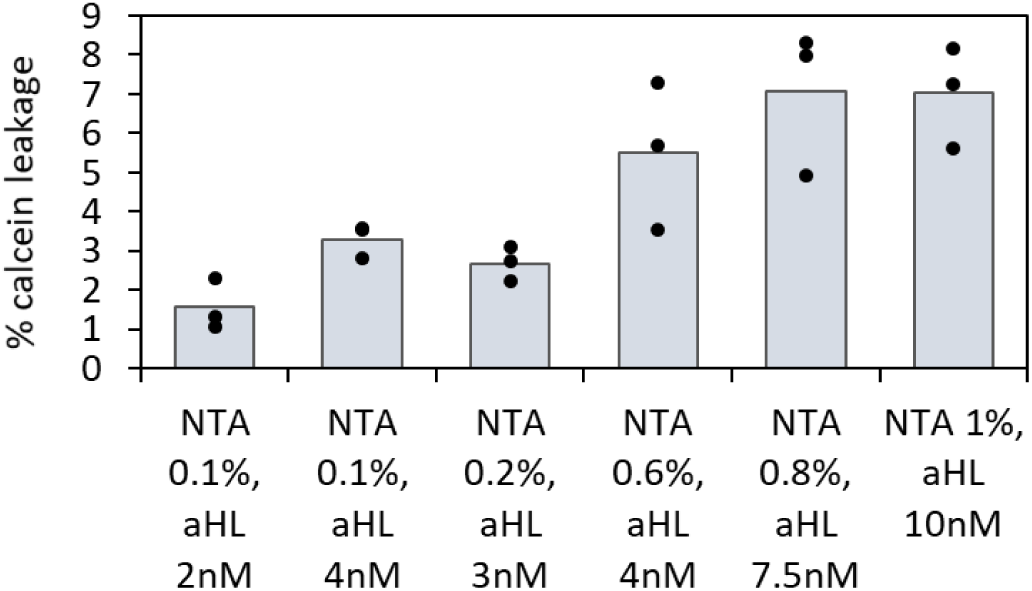
Leakage test for double His labeled αHL fusion experiments. Leakage of small molecule calcein was tested in liposomes with NTA lipids and liposomes with αHL, after 12h incubation at 30°C. Liposomes contained 1mM calcein, and the complete translation mix, αHL and GFP plasmids, at the beginning of the experiment. Examples of individual purification plots for samples summarized on this figure are on **Figure S4**. Dots on top of the bar graphs indicate individual values of three replicates.

**Figure S4.**
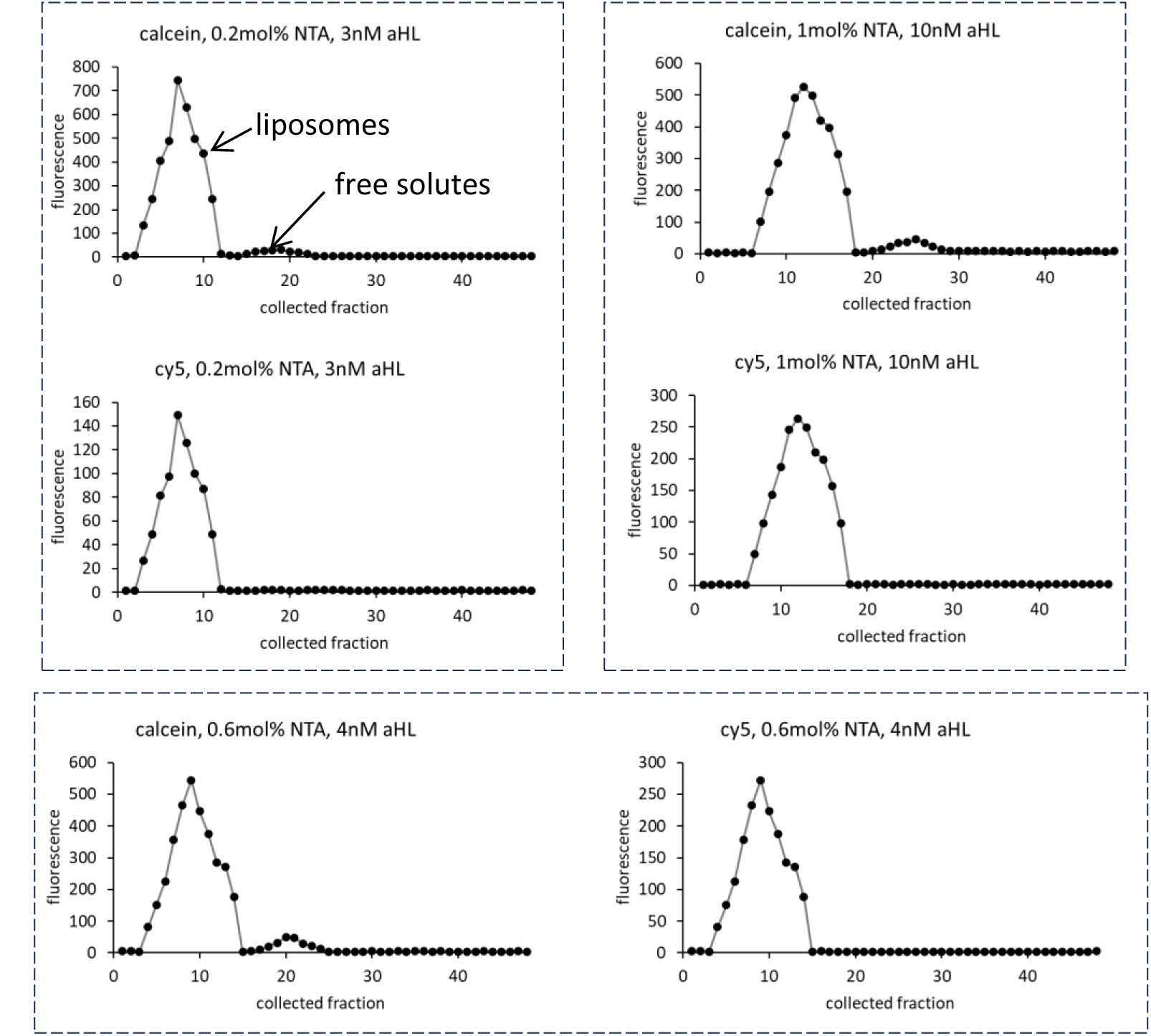
Leakage test for double His labeled αHL fusion. Example plots from size exclusion chromatography purification of synthetic cell liposomes. Leakage of small molecule calcein was tested in liposomes with NTA lipids and liposomes with αHL, after 12h incubation at 30°C. Liposomes contained 1mM calcein at the beginning of the experiment. Summarized data for all samples are on **Figure S3**. Each sample was analyzed using two fluorescent measurements: calcein channel and Cy5 channel (see **Table S5** for wavelengths). The calcein channel traces indicate leakage of small molecule and the Cy5 channel (Cy5 lipid dye 1,2-dioleoyl-sn-glycero-3-phosphoethanolamine-N-(Cyanine 5)) indicates which fractions contain liposomes. This data was collected using TxTl.

**Figure S5.**
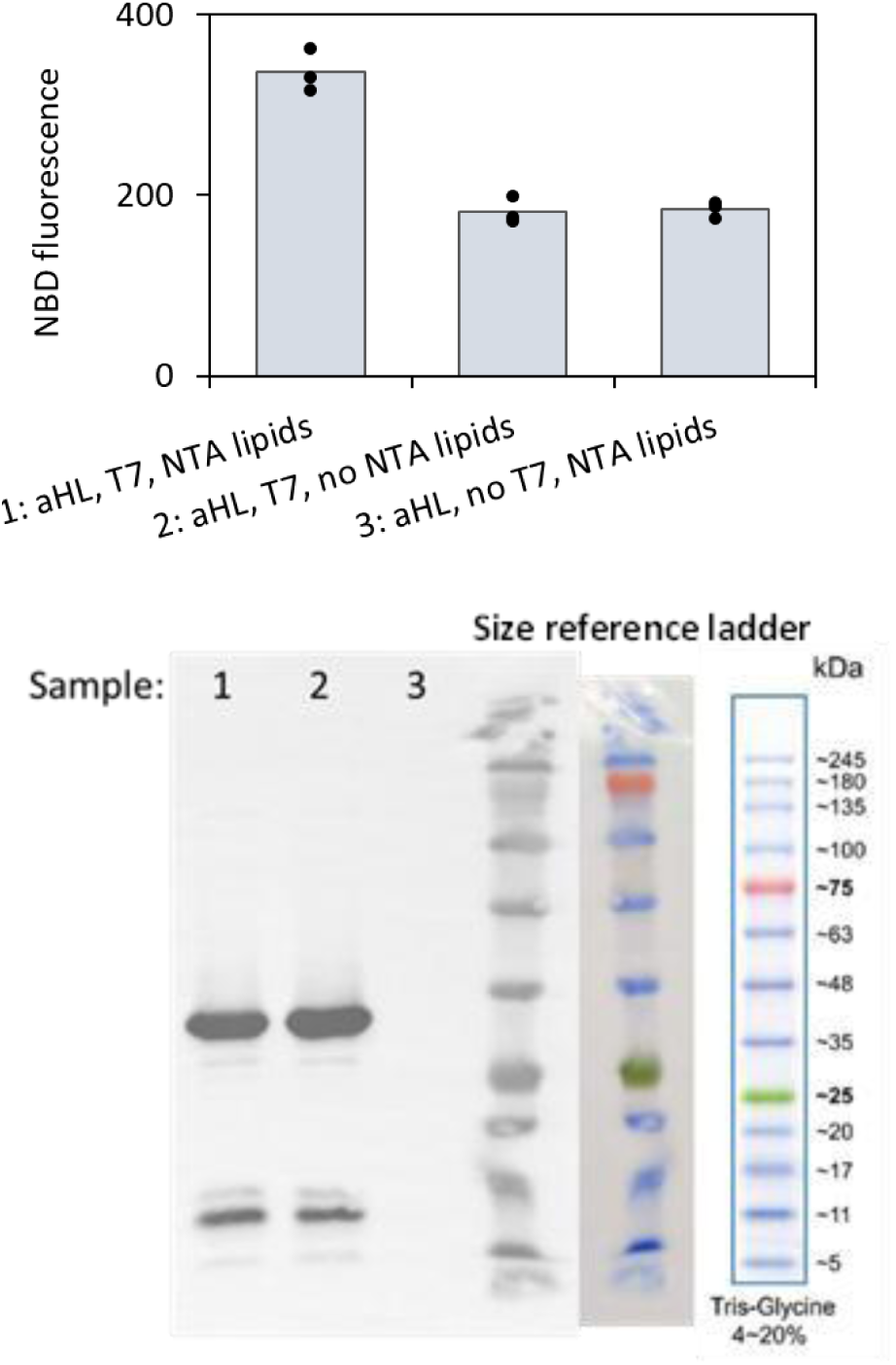
FRET data for liposome fusion samples corresponding to the Western Blot analysis of αHL expression in synthetic cell liposomes. Population 1 of liposomes were prepared as described in Methods section 4, encapsulating *E. coli* TxTl, αHL plasmid and T7 RNA polymerase plasmid. Population 2 was prepared with FRET dye pair (NBD and rhodamine) to trace fusion, and population 2 contained Ni-NTA lipids (or not, as indicated below). The population 1 was incubated at 30°C for 12h to induce αHL expression, followed by mixing with population 2. The FRET signal was analyzed, followed by Western Blot analysis of αHL expression. sample 1: αHL plasmid, T7 plasmid, NTA lipids, expected: fusion and αHL expression sample 2: αHL plasmid, T7 plasmid, no NTA lipids, expected: αHL expression, no fusion sample: αHL plasmid, no T7 plasmid, NTA lipids, expected: no fusion and no αHL expression Dots on top of the bars indicate all the individual data points, the bars are the average value, n=3 independently prepared and processed liposome samples. This data was collected using TxTl.

**Figure S6.**
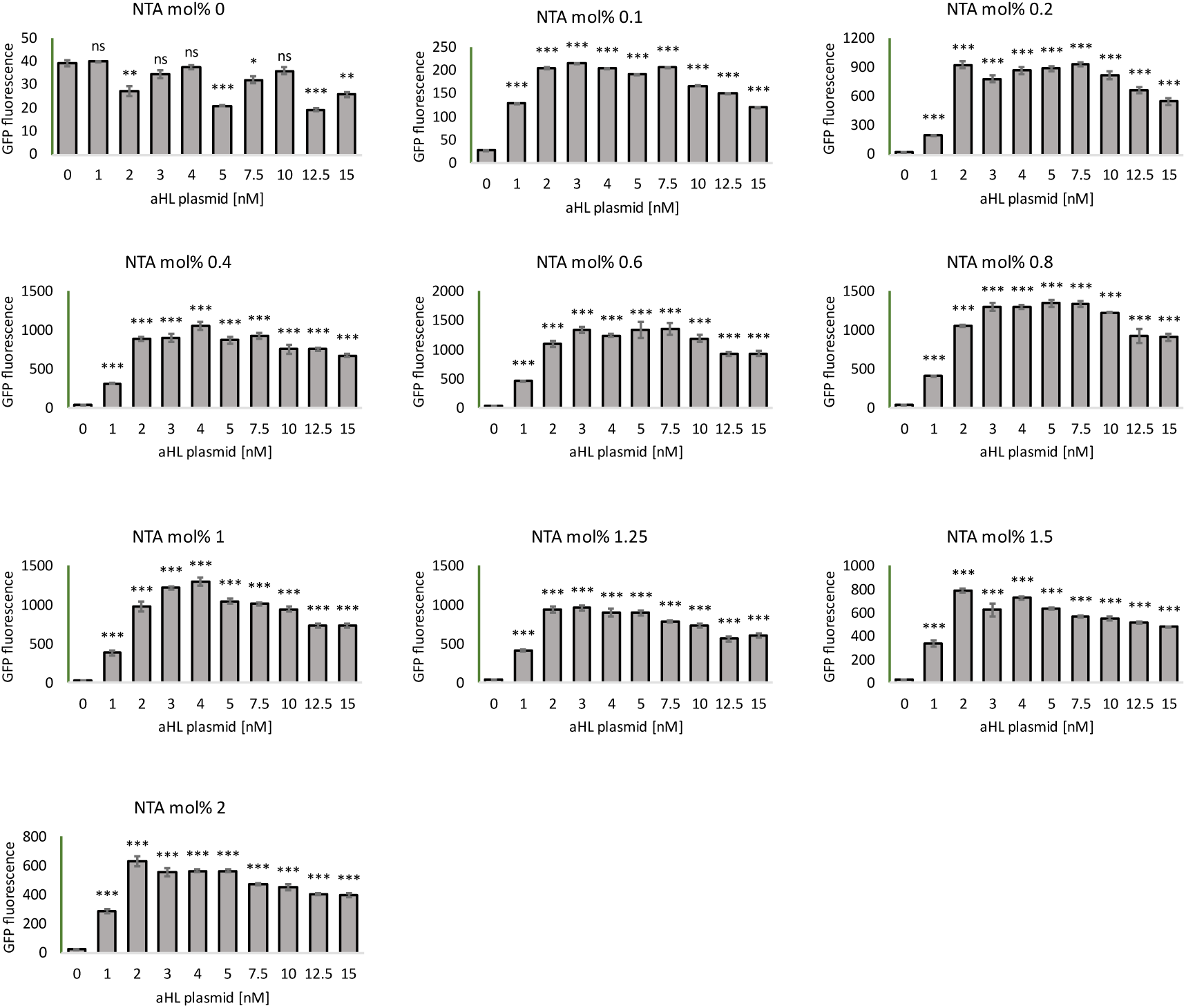
His labeled αHL fusion, GFP data. Individual GFP fluorescence data points for the heatmap on Figure 2. Error bars indicate SEM, n=3 independently prepared and processed liposome samples. Statistical analysis shows P values for each series of NTA concentrations compared to 0mM αHL plasmid. ns = P > 0.0; * = P ≤ 0.0, ** = P ≤ 0.01, *** = P ≤ 0.001. This data was collected using TxTl.

**Figure S7.**
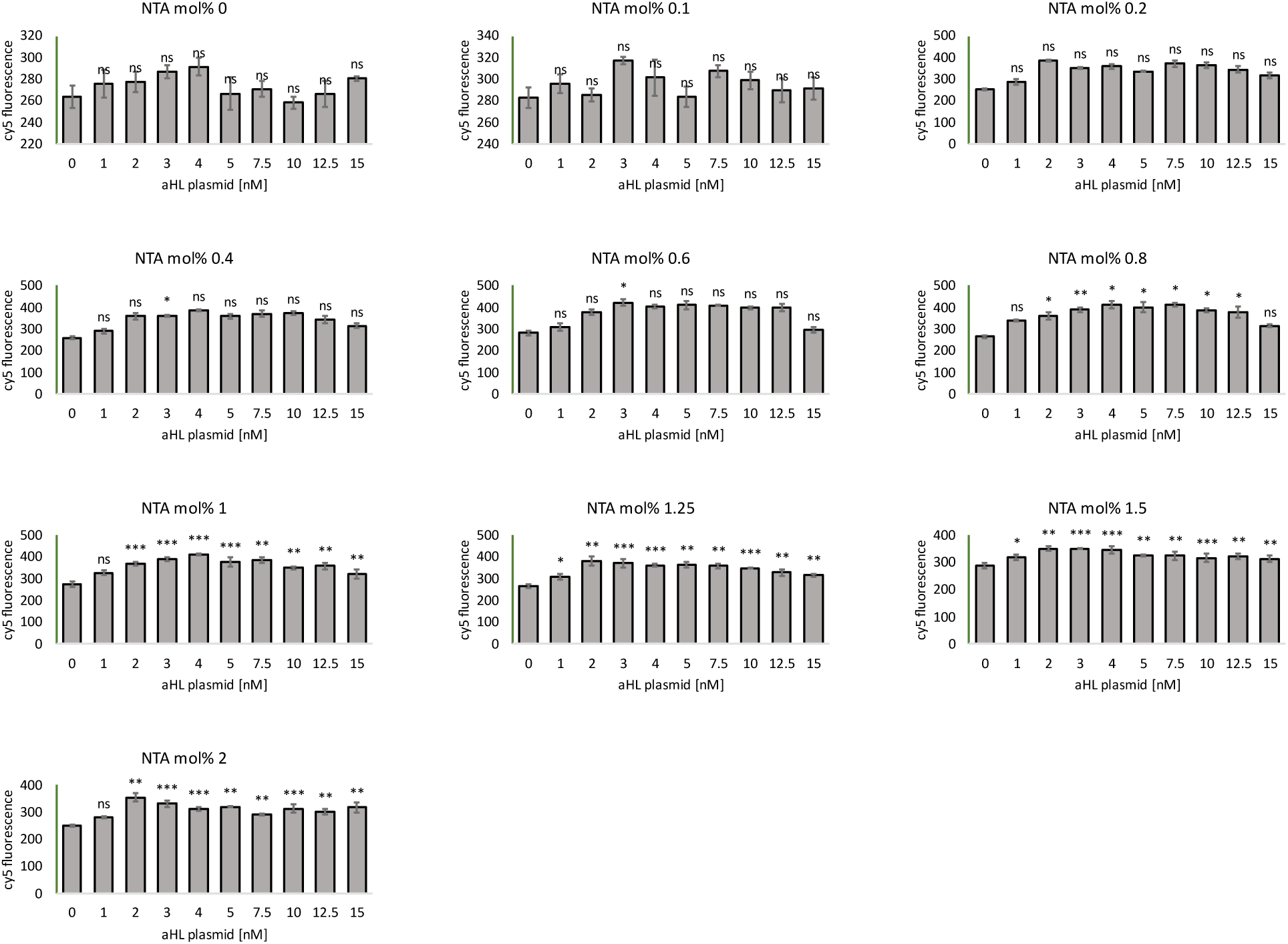
His labeled αHL fusion, Cy5 FRET data. Individual Cy5 FRET fluorescence data points for the heatmap on Figure 2. Error bars indicate SEM, n=3 independently prepared and processed liposome samples. Statistical analysis shows P values for each series of NTA concentrations compared to 0mM αHL plasmid. ns = P > 0.0; * = P ≤ 0.0, ** = P ≤ 0.01, *** = P ≤ 0.001. This data was collected using TxTl.

**Figure S8.**
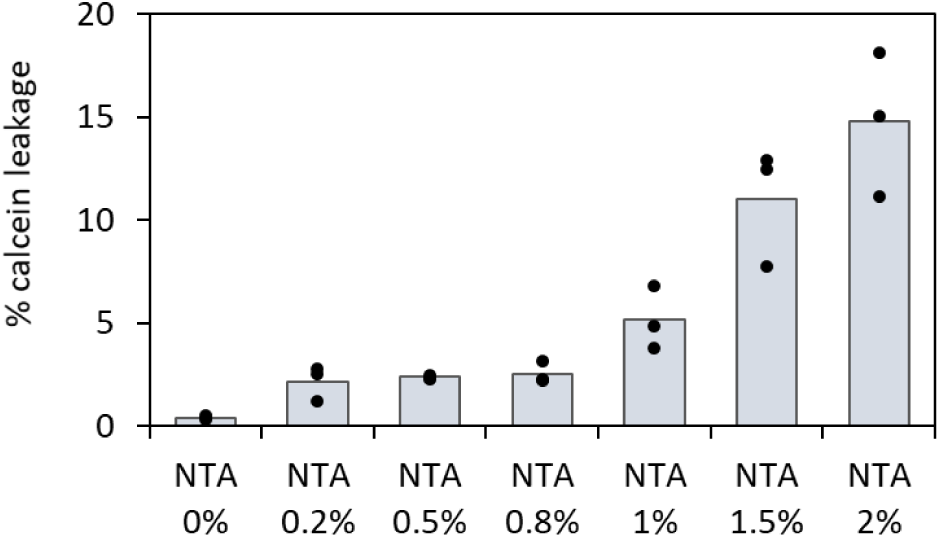
Leakage test for NTA lipids. Testing leakage of calcein from liposomes with varying amount of NTA lipid in the membrane, expressed as mol% of total lipids. Leakage was calculated as % of calcein detected outside of the liposomes, measured via size exclusion chromatography purification after 12h incubation at 30°C. Examples of individual purification plots for samples summarized on this figure are on **Figure S10**. Dots on top of the bar graphs indicate individual values of three replicates.

**Figure S9.**
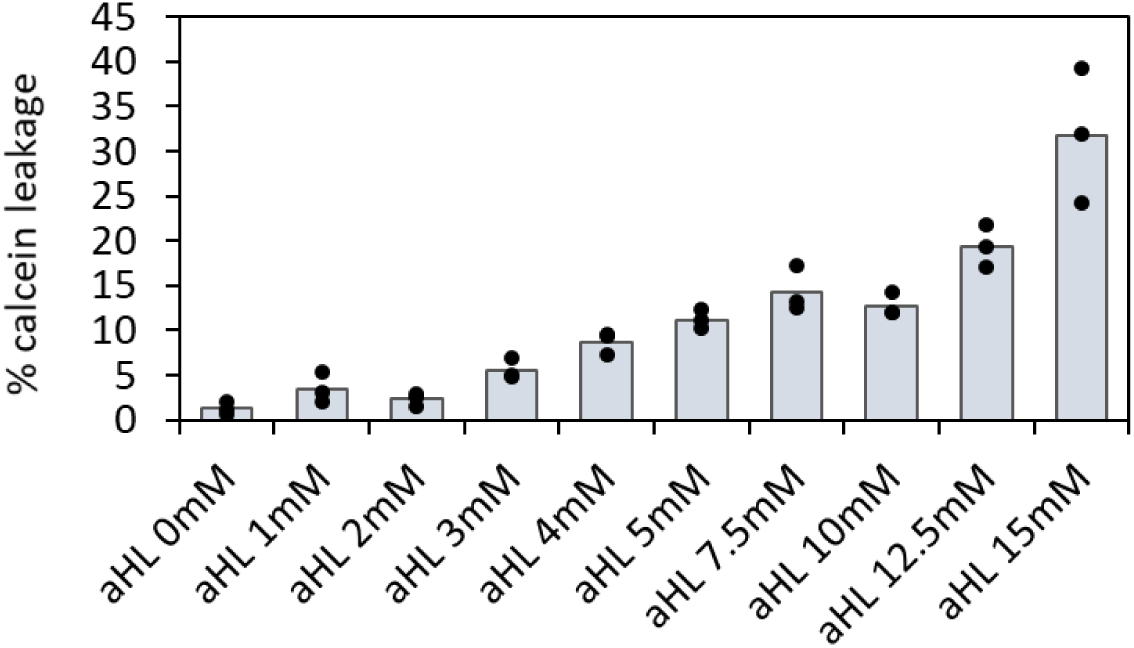
Test leakage for αHL expression. Testing leakage of calcein from liposomes with varying amount of αHL plasmid. Leakage was calculated as % of calcein detected outside of the liposomes, measured via size exclusion chromatography purification after 12h expression incubation at 30°C. Examples of individual purification plots for samples summarized on this figure are on **Figure S11**. Dots on top of the bar graphs indicate individual values of three replicates. This data was collected using TxTl.

**Figure S10.**
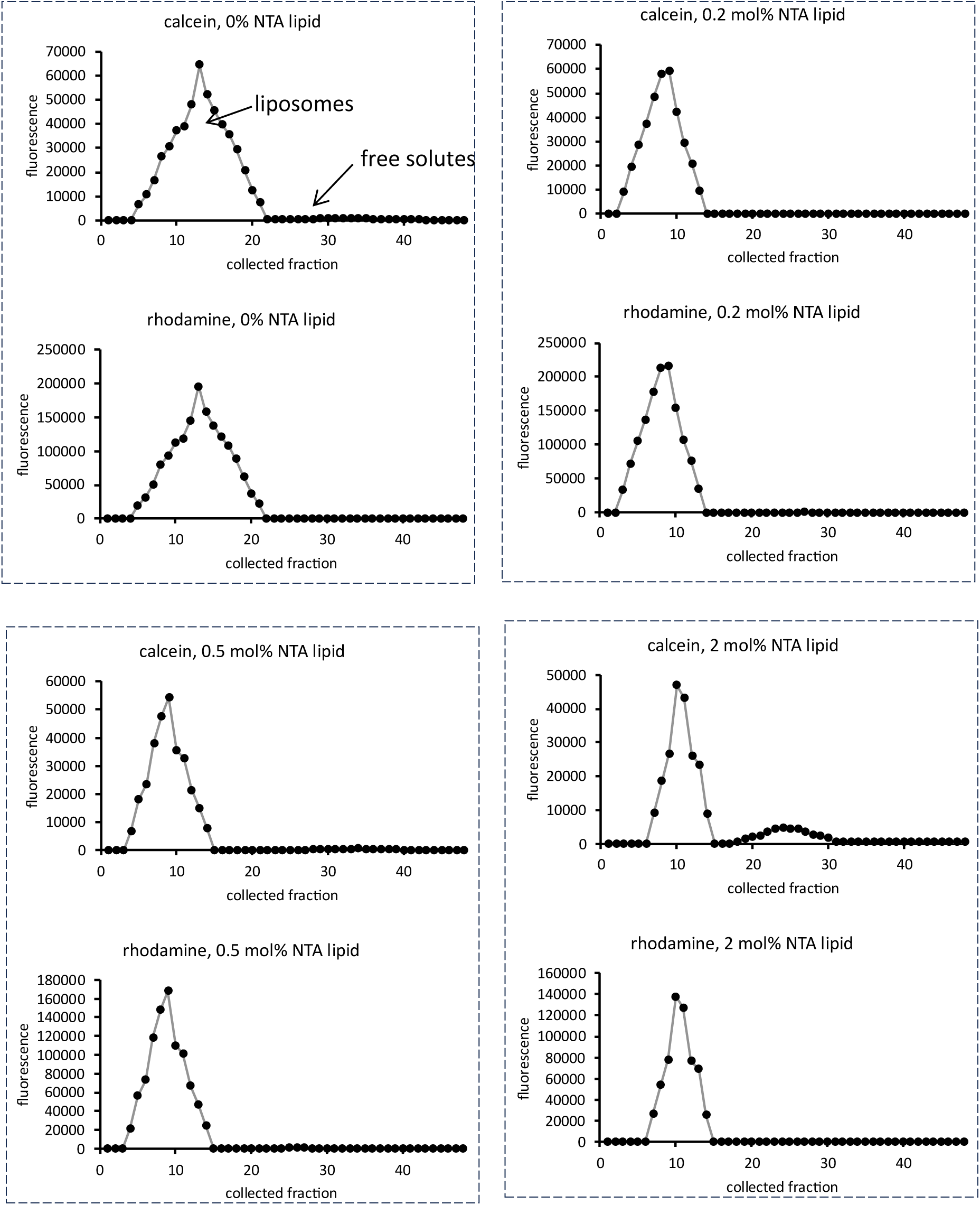
Leakage test for NTA lipids. Example plots from size exclusion chromatography purification of liposomes with varying amount of NTA lipid as mol% of total lipids. Leakage of small molecule calcein was tested in liposomes with NTA lipids, after 12h incubation at 30°C. Liposomes contained 1mM calcein at the beginning of the experiment. Each sample was analyzed using two fluorescent measurements: calcein channel and rhodamine channel (see **Table S5** for wavelengths). The calcein channel traces indicate leakage of small molecules and the rhodamine (red membrane dye, Lissamine Rhodamine B 1,2-Dihexadecanoyl-sn-Glycero-3-Phosphoethanolamine) channel indicates which fractions contain liposomes.

**Figure S11.**
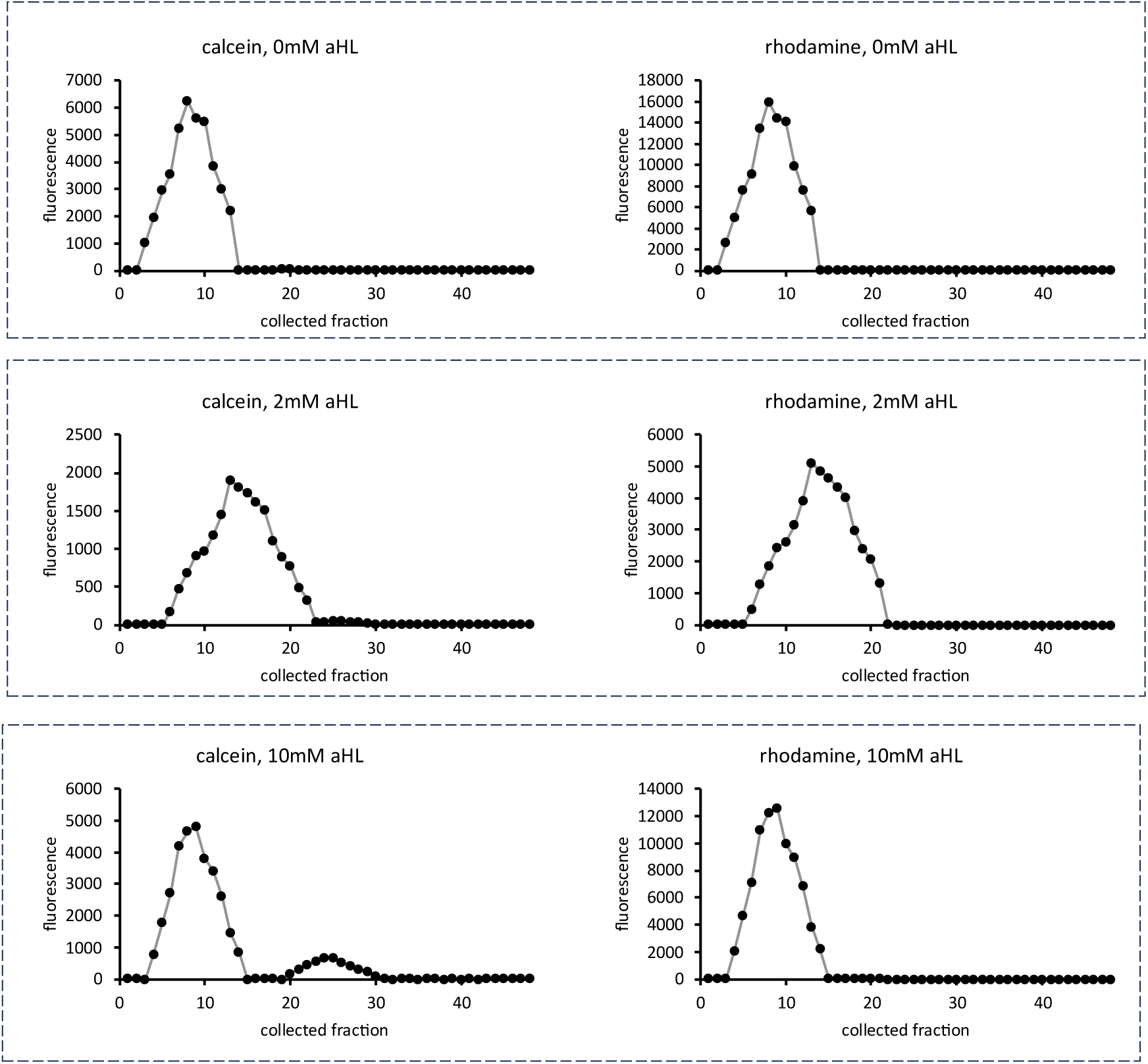
Test leakage for αHL expression. Example plots from size exclusion chromatography purification of liposomes with varying amount of αHL plasmid. Leakage of small molecule calcein was tested after 12h incubation at 30°C. Liposomes contained 1mM calcein at the beginning of the experiment. Each sample was analyzed using two fluorescent measurements: calcein channel and rhodamine channel (see **Table S5** for wavelengths). The calcein channel traces indicate leakage of small molecules and the rhodamine (red membrane dye, Lissamine Rhodamine B 1,2-Dihexadecanoyl-sn-Glycero-3-Phosphoethanolamine) channel indicates which fractions contain liposomes. This data was collected using TxTl.

**Figure S12.**
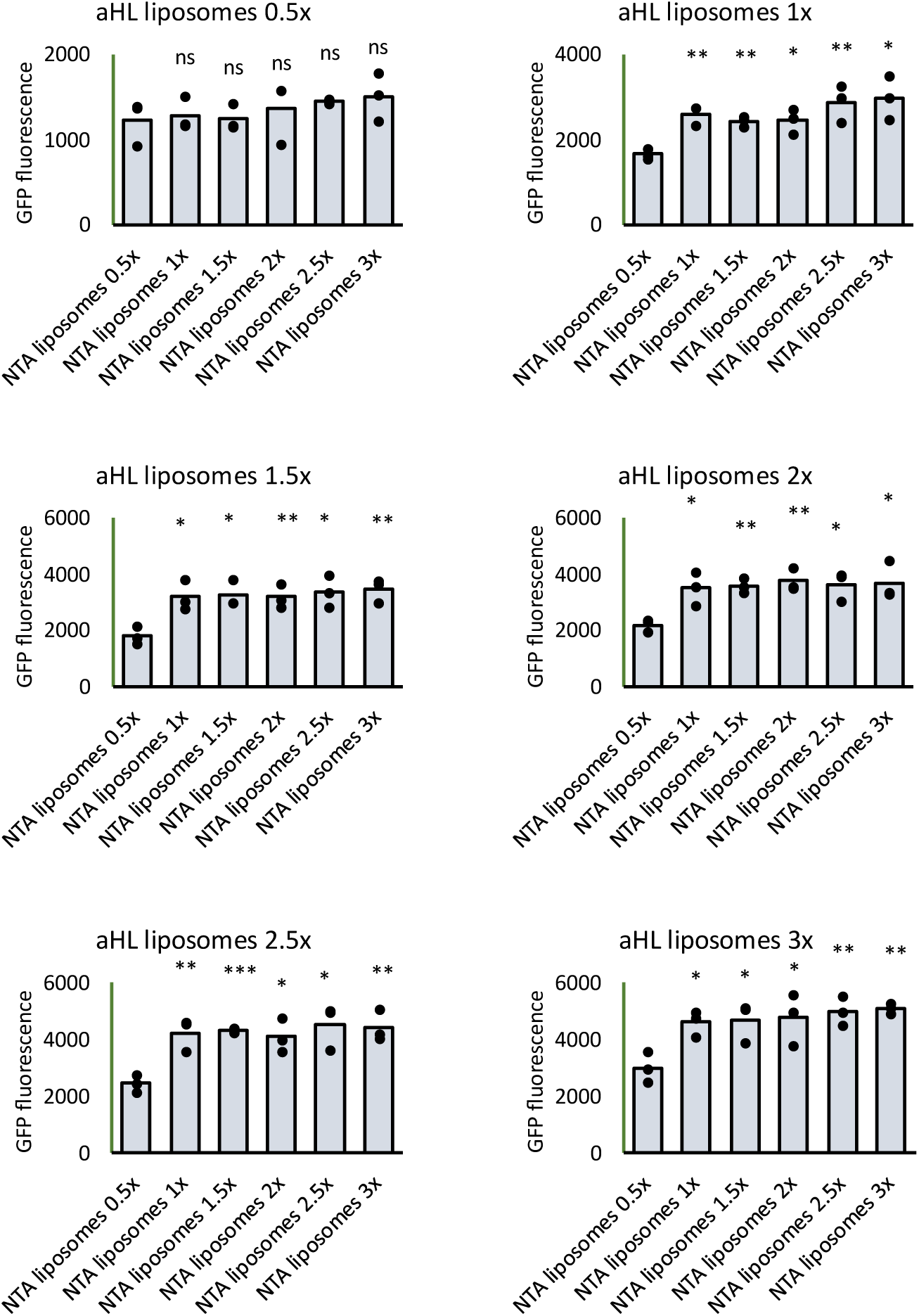
Liposome fusion testing at different liposome ratios. Individual data points for the heat map on Figure 2. Dots on top of the bar graphs indicate individual values of three replicates. Statistical analysis shows P values for each series of liposome concentrations compared to NTA liposomes 0. x. ns = P > 0.0; * = P ≤ 0.0, ** = P ≤ 0.01, *** = P ≤ 0.001. This data was collected using TxTl.

**Figure S13.**
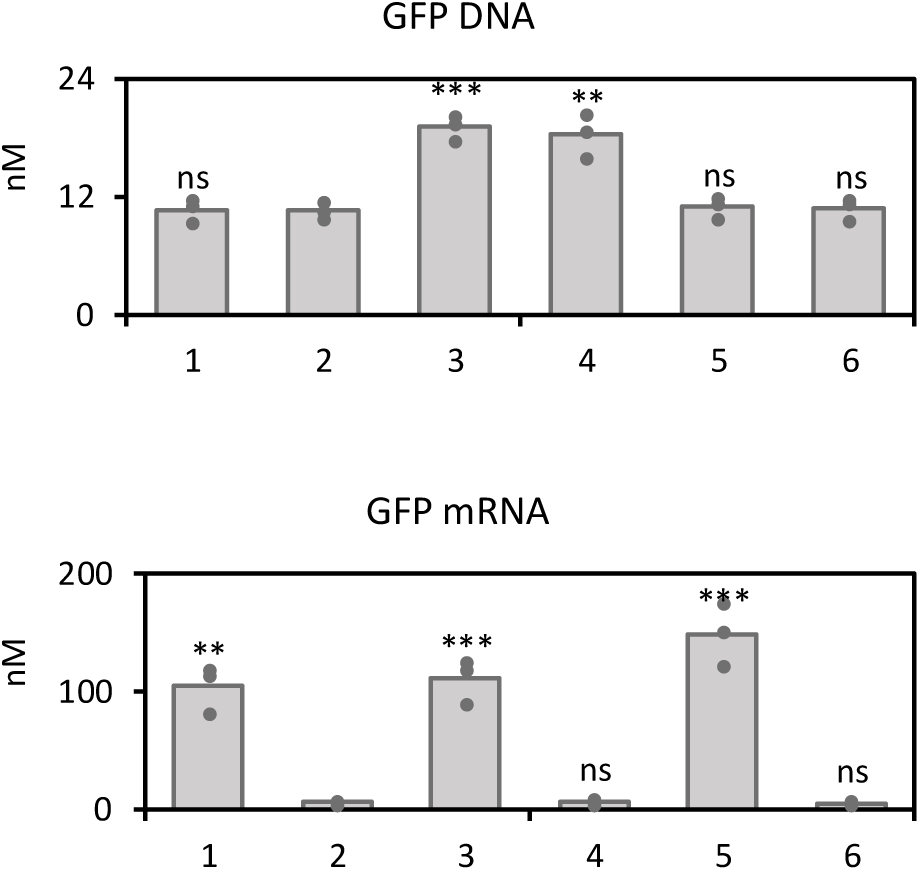
GFP DNA and GFP mRNA after genetically encoded lumen mixing of synthetic cells. Individual data points for quantification of GFP plasmid and GFP mRNA in synthetic cells after fusion event data shown on Figure 2. Samples are in the same order as shown on panels J and K of Figure 2. <u>Sample 1</u>: 1x cells of population 1 (with T7RNAP and αHL), 1x population 2 (with GFP plasmid) with Ni-NTA, expected fusion <u>Sample 2</u>: 1x cells of population 1 (with T7RNAP and αHL), 1x population 2 (with GFP plasmid) without Ni-NTA, no fusion is expected <u>Sample 3</u>: 1x cells of population 1 (with T7RNAP and αHL), 2x population 2 (with GFP plasmid) with Ni-NTA, expected fusion <u>Sample 4</u>: 1x cells of population 1 (with T7RNAP and αHL), 2x population 2 (with GFP plasmid) without Ni-NTA, no fusion is expected <u>Sample 5</u>: 2x cells of population 1 (with T7RNAP and αHL), 1x population 2 (with GFP plasmid) with Ni-NTA, expected fusion <u>Sample 6</u>: 2x cells of population 1 (with T7RNAP and αHL), 1x population 2 (with GFP plasmid) without Ni-NTA, no fusion is expected Bar graphs represent average value, dots are individual data points for each replicate. Statistical analysis shows P values for each series of NTA concentrations compared to negative control experiment of no Ni-NTA in Sample 2. ns = P > 0.0; * = P ≤ 0.0, ** = P ≤ 0.01, *** = P ≤ 0.001. This data was collected using TxTl.

**Figure S14.**
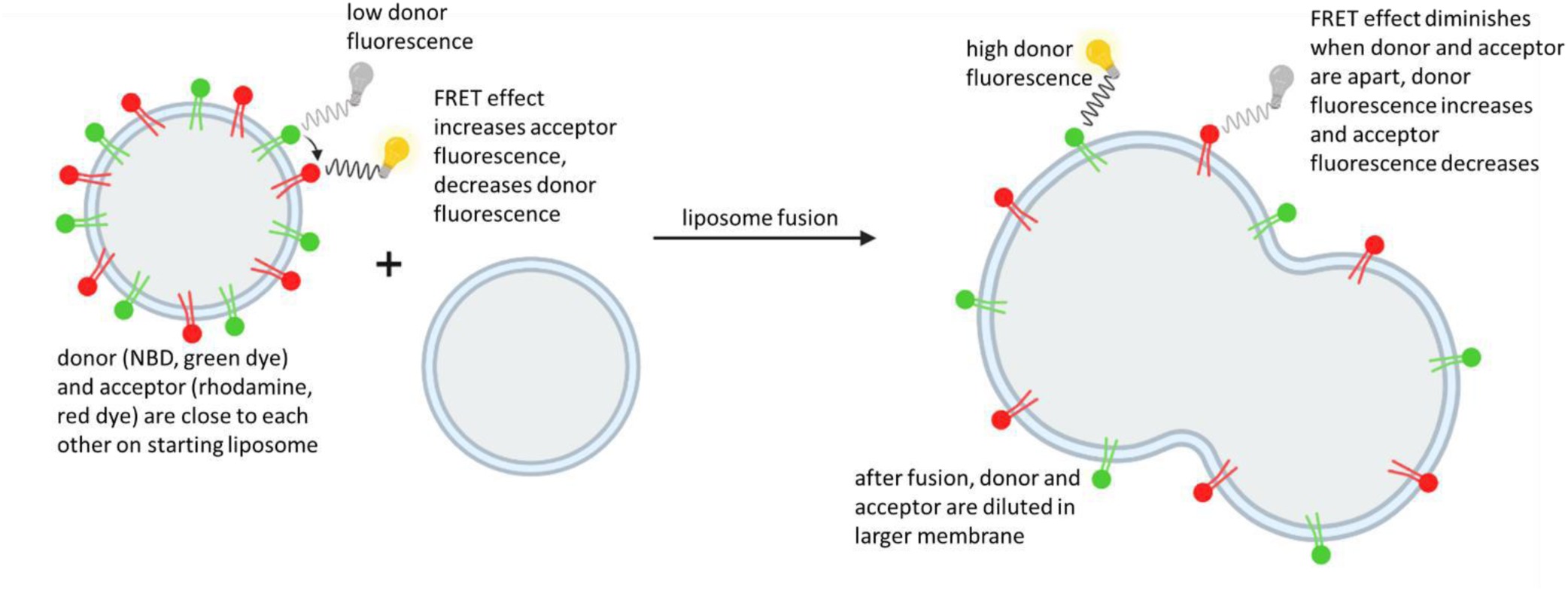
General scheme for liposome FRET size measurement. Liposome fusion experiment monitored by FRET effect, measuring fluorescence of donor dye from the FRET pair. In synthetic cell fusion experiments, one population of cells contains both FRET donor and FRET acceptor, while the other population does not contain any FRET dyes. Before fusion, the FRET dyes are in relatively close proximity, resulting in high FRET effect, which means low donor fluorescence signa and higher acceptor fluorescence signal. After fusion and membrane mixing, the FRET dyes are “diluted” in the now larger membrane, putting the FRET dyes further apart from each other on the larger membrane surface. The efficiency of the FRET effect decreases, which results in increase of donor fluorescence and decrease of acceptor fluorescence. In the fusion experiments in this work, two different pairs of FRET dyes were used. Donor NBD - acceptor Rhodamine, this dye pair was used when no concurrent GFP monitoring was needed (since NBD fluorescence spectrally overlaps with GFP) Donor Cy5 – acceptor Cy7, this dye pair was used to allow for monitoring of GFP fluorescence at the same time. In all FRET experiments, we monitored the fluorescence of the donor dye. Increased donor fluorescence indicated increased membrane mixing.

**Figure S15.**
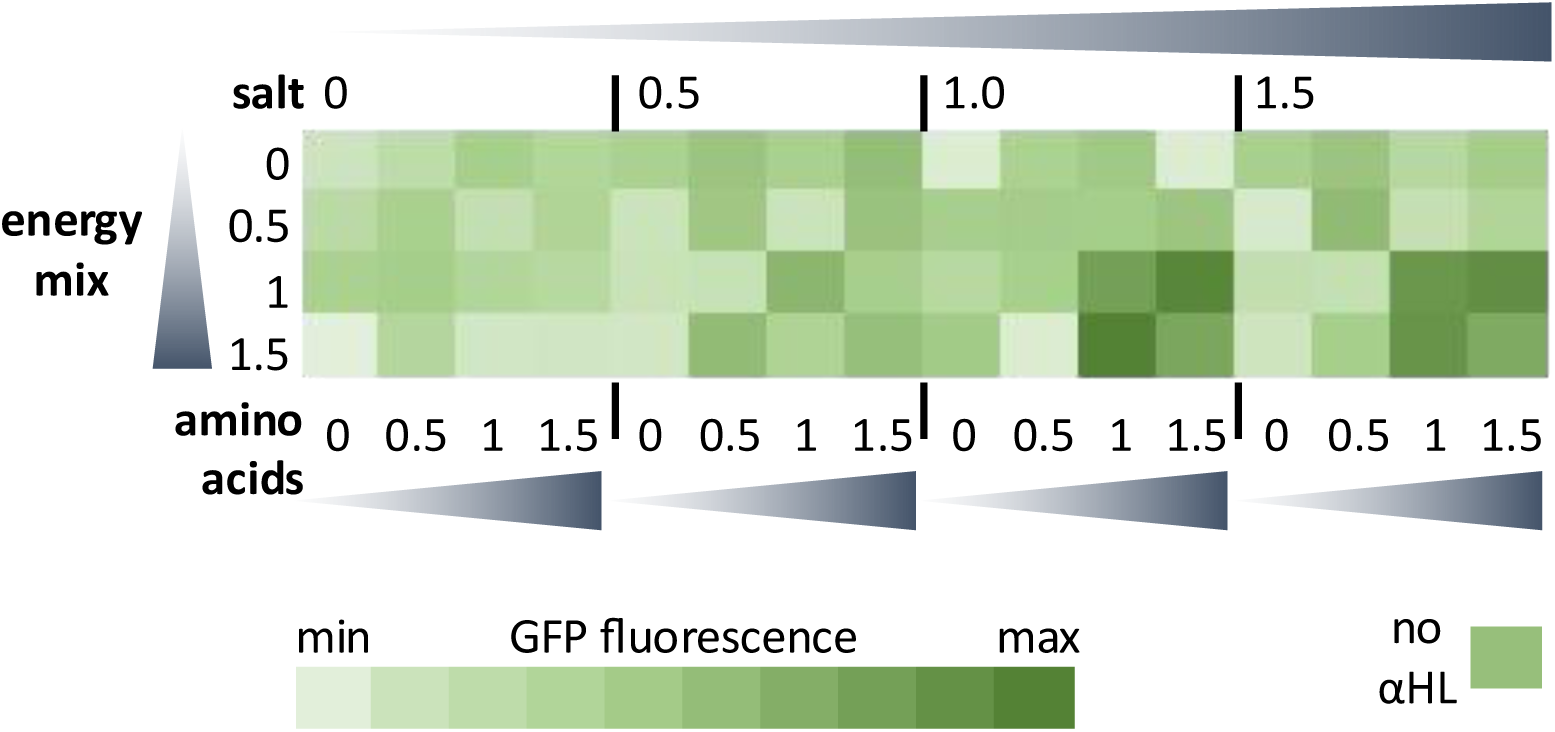
Optimization of synthetic cell media. GFP expression in synthetic cells with 10nM αHL plasmid (and control without αHL), with the external buffer (“media”) contained varying amounts of one of the three cell-free translation system components: “salt”, “amino acids” and “energy mix”. Values on the plot correspond to the standard reaction final concentration of 1x, so 0.5 is 50% final concentration compared to standard reaction conditions, and 1.5 is 150% of the standard reaction conditions concentration (see methods section 13). Samples were incubated at 30°C for 24h. End point GFP fluorescence was recorded. The 1x, or 100%, concentrations used through this project were: “<u>salt</u>” mixture: 1 0mM potassium glutamate, 10mM ammonium acetate, 10mM magnesium glutamate “<u>amino acid</u>” mixture: 2mM of each of the 20 amino acids from a KOH amino acid stock “<u>energy mix</u>” mixture: 1. mM ATP and GTP, 0.9mM CTP and UTP, 0.068mM folinic acid, 200ug/mL of *E. coli* tRNA mixture, 0.33mM nicotinamide adenine dinucleotide (NAD), 0.26mM coenzyme-A (CoA), 1.5mM spermidine, 4mM sodium oxalate, 0.75mM cAMP, 30mM 3-PGA, 50mM HEPES pH 8. The “no αHL” control contained GFP plasmid but no αHL plasmid, and the external media was at highest concentration of all three components (1.5x salt, amino acid and energy mixes). Individual data points for all heatmap points are in **Figure S16**. This data was collected using TxTl.

**Figure S16.**
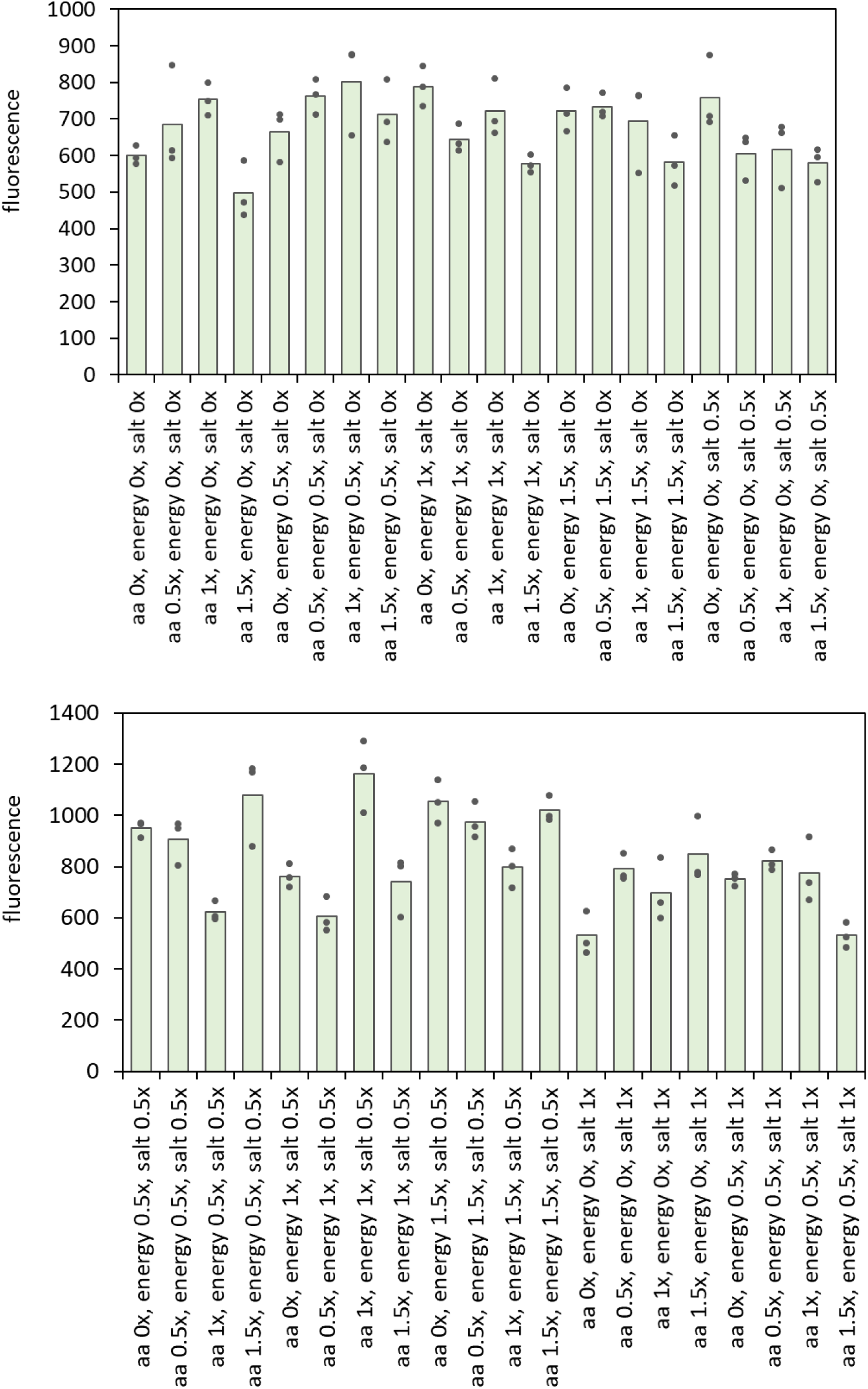
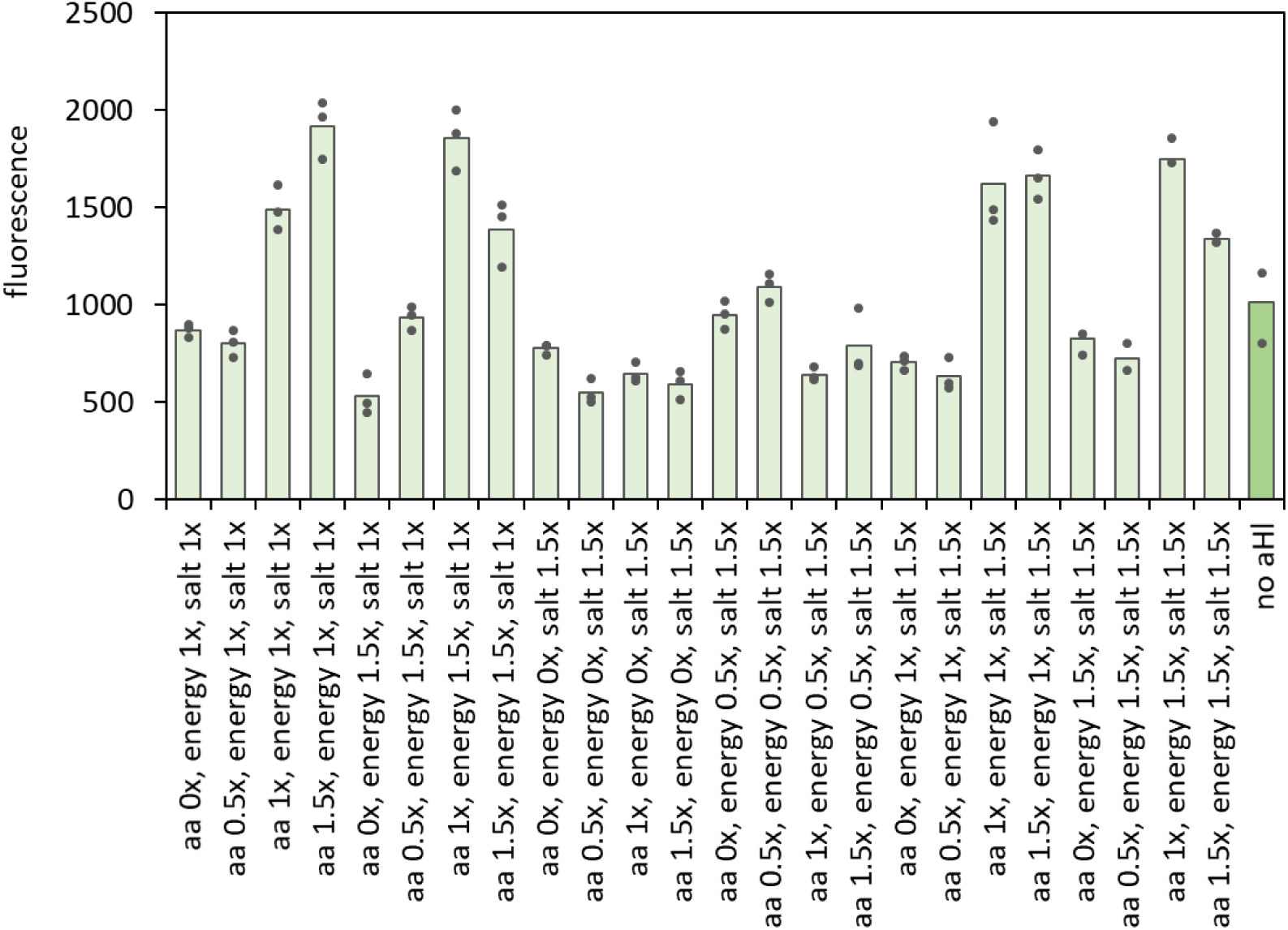
Optimization of synthetic cell media. Individual data points for the heat map on **Figure S15**. aa: amino acids. The “no αHL” control contained GFP plasmid but no αHL plasmid, and the external media was at highest concentration of all three components (1.5x salt, amino acid and energy mixes). Dots on top of the bar graphs indicate individual values of three replicates. This data was collected using TxTl.

**Figure S17.**
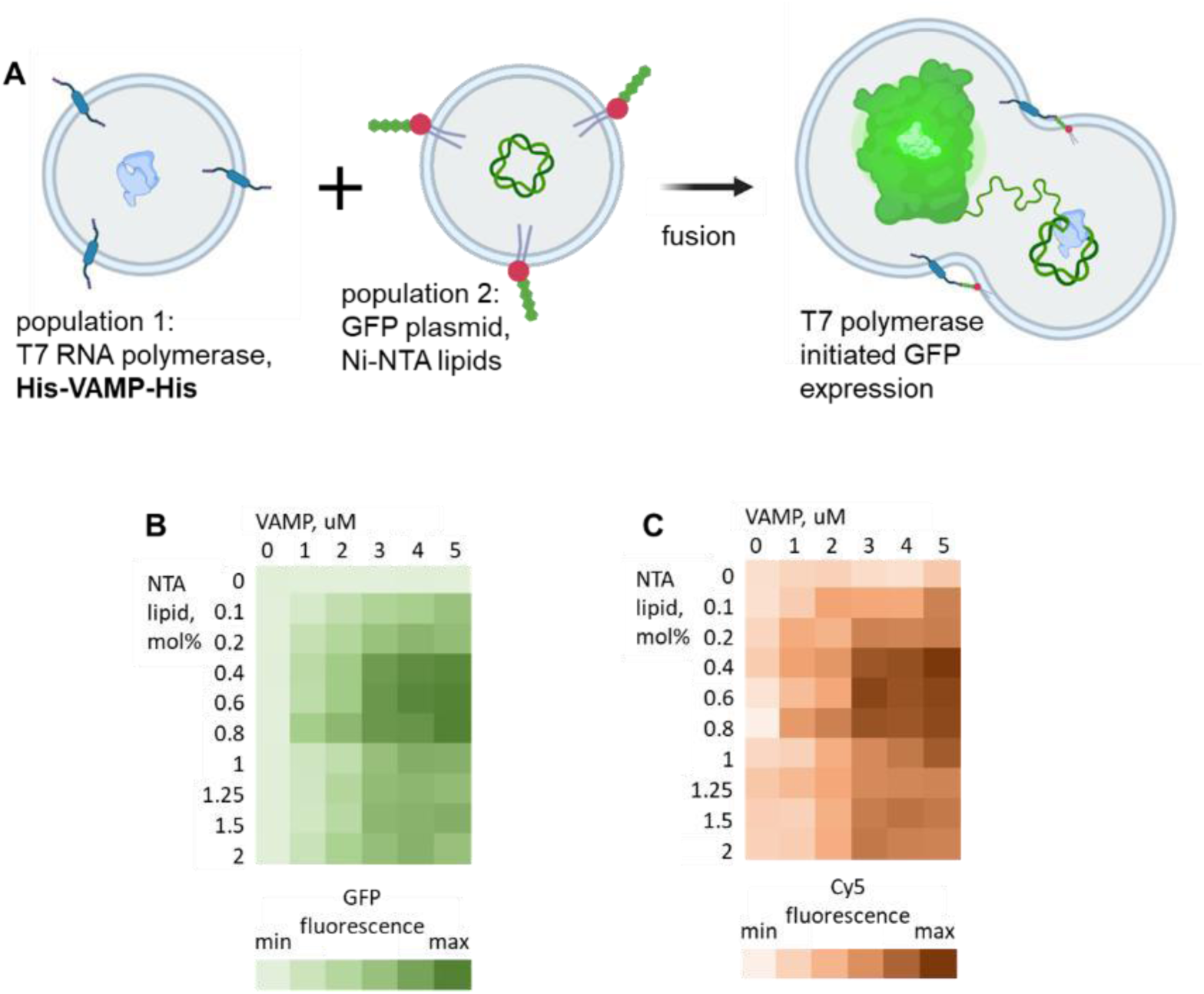
VAMP2-His fusion testing. **A**: Two populations of synthetic cells were mixed, population 1 containing VAMP transmembrane domain labeled with His tag and encapsulating T7 RNA polymerase, and population 2 containing Ni-NTA lipid tags and encapsulating plasmid with GFP under T7 promoter. Only if the lumen of the membranes fuses, GFP could be expressed (T7 RNA polymerase from population 1 is needed to express GFP plasmid in population 2). **B**: Lumen fusion, measured as expression of GFP, for experiments with varying amount of VAMP protein and varying amount of NTA lipid (described as mol% of total lipid). Higher GFP fluorescence means more lumen mixing. **C**: Membrane mixing, measured as FRET interaction between Cy5 and Cy7 FRET pair (see **Figure S14** for general scheme of FRET experiment). Higher Cy5 fluorescence means more membrane mixing. Population 2 was labeled with 0.1mol% of each Cy5 and Cy7 lipid anchored dyes. Individual heat map data points for GFP are on **Figure S18**, and individual heat map data points for Cy5 are on **Figure S19**. This data was collected using TxTl.

**Figure S18.**
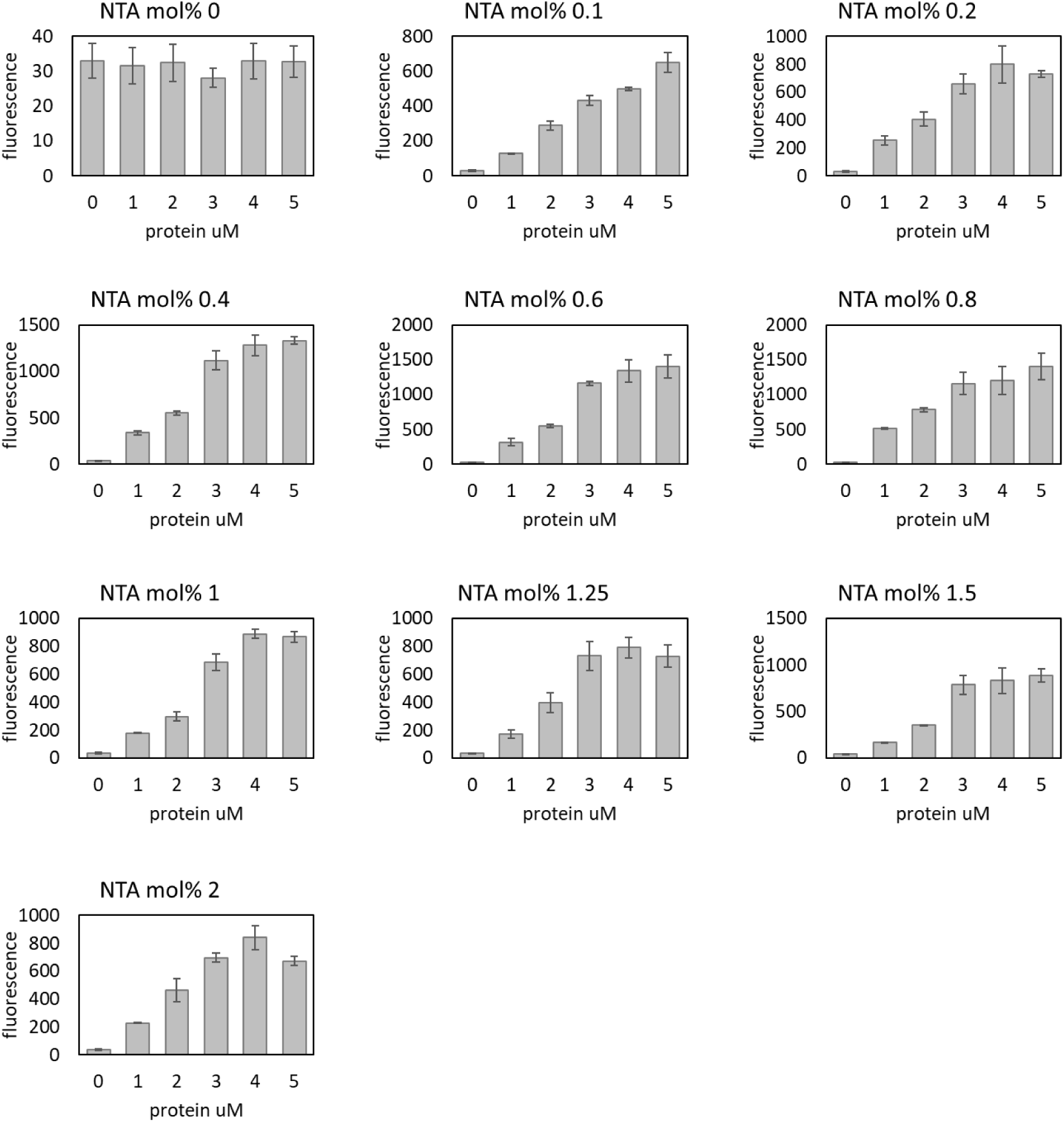
VAMP2-His fusion testing, GFP. Data points for heat map on **Figure S17**, GFP fluorescence (see **Table S5** for wavelengths). Error bars indicate SEM, n=3 independently prepared and processed liposome samples. This data was collected using TxTl.

**Figure S19.**
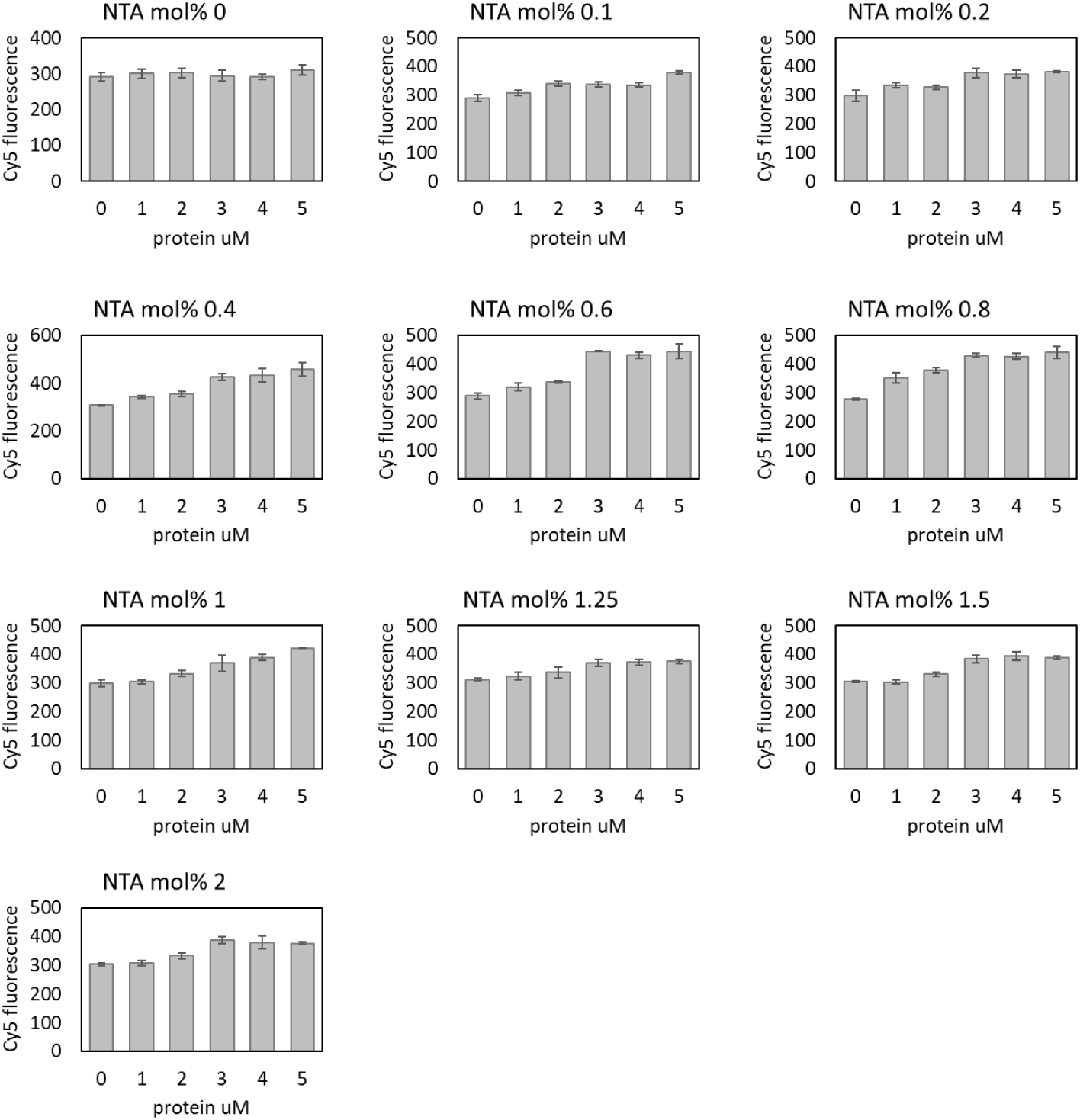
VAMP2-His fusion testing, Cy5. Data points for heat map on **Figure S17**, Cy5 fluorescence (see **Table S5** for wavelengths). Higher Cy5 fluorescence means more membrane mixing. Population 2 was labeled with 0.1mol% of each Cy5 and Cy7 lipid anchored dyes. Error bars indicate SEM, n=3 independently prepared and processed liposome samples. This data was collected using TxTl.

**Figure S20.**
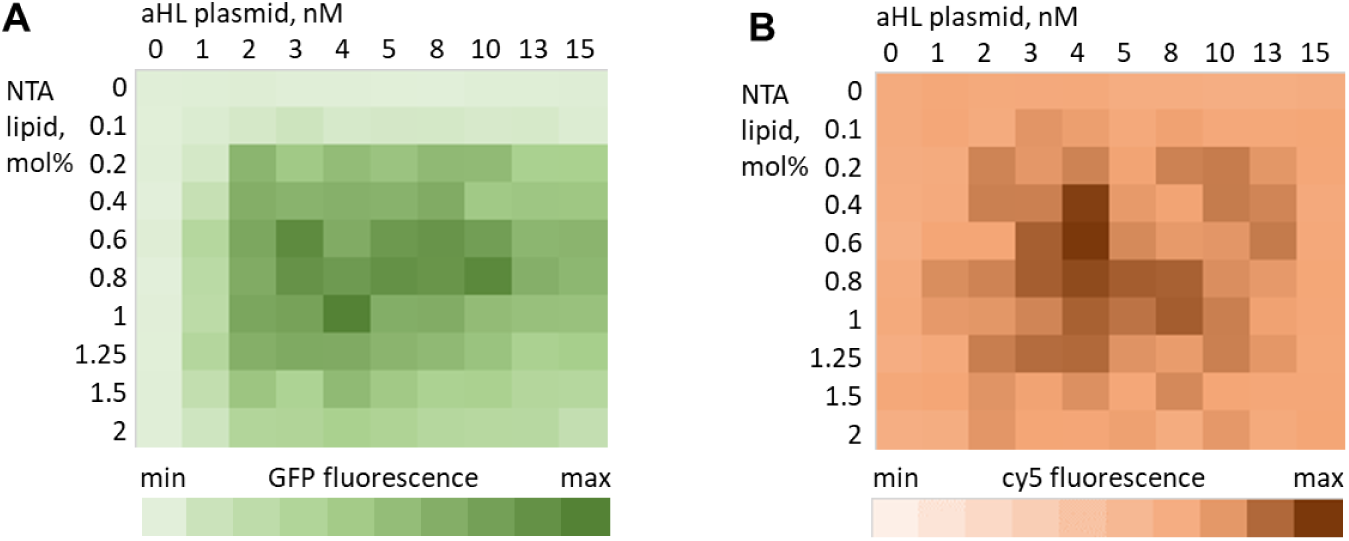
Fusion with αHL introduced externally. Fusion between two populations of synthetic cells, one carrying T7RNAP and the other population carrying GFP DNA and Ni-NTA lipid tag. Lumen fusion was monitored by GFP expression (panel A) and membrane mixing was monitored by Cy5 FRET signal (panel B). Individual heat map data points for GFP are on **Figure S21** and individual heat map data points for Cy5 are on **Figure S22**. After fusion, selected samples were purified on size exclusion column to measure content leakage, data on **Figure S23**. This data was collected using TxTl.

**Figure S21.**
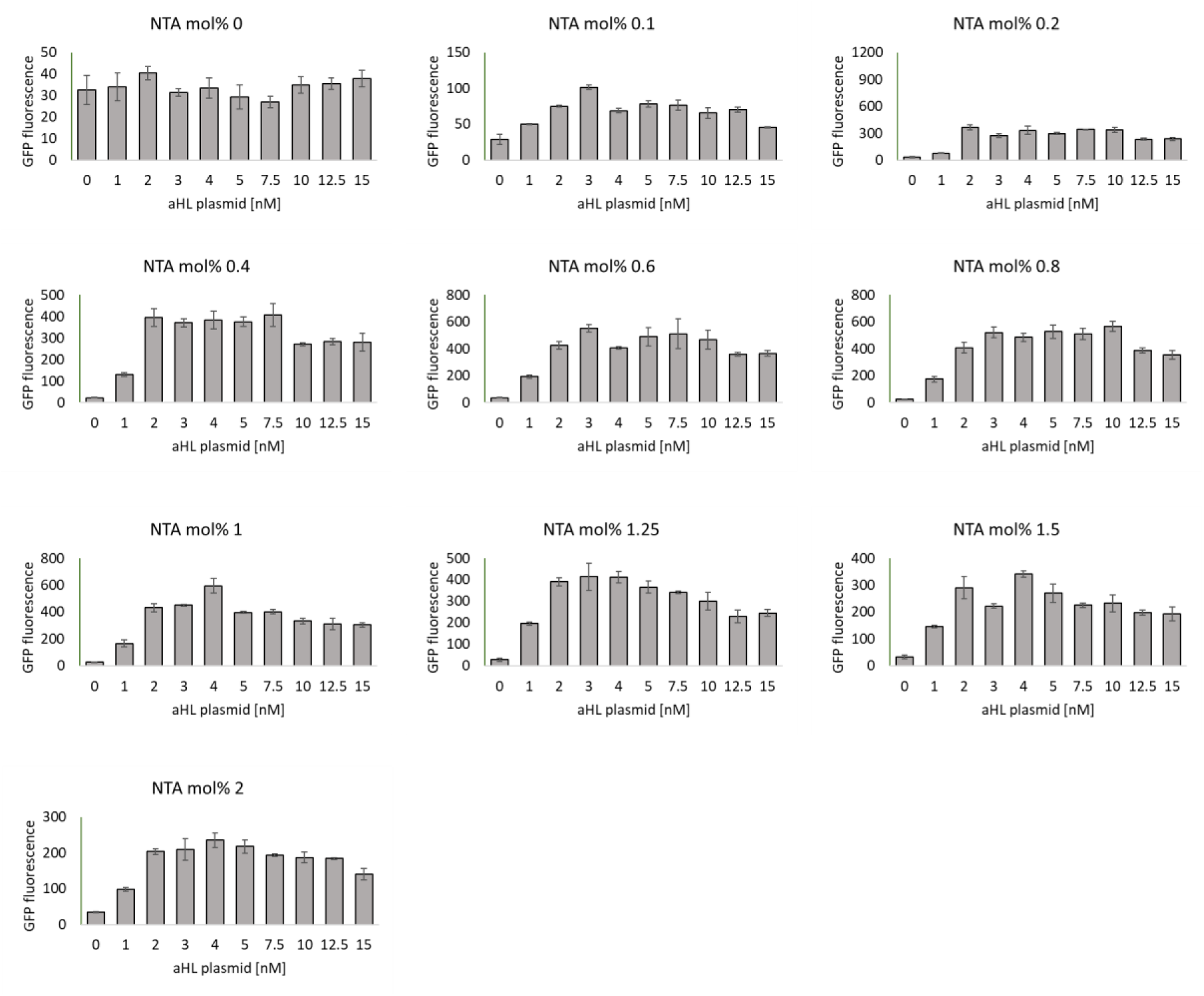
Fusion testing with external αHL, GFP signal. Individual GFP fluorescence data points for the heatmap on **Figure S20**. Error bars indicate SEM, n=3 independently prepared and processed liposome samples. This data was collected using TxTl.

**Figure S22.**
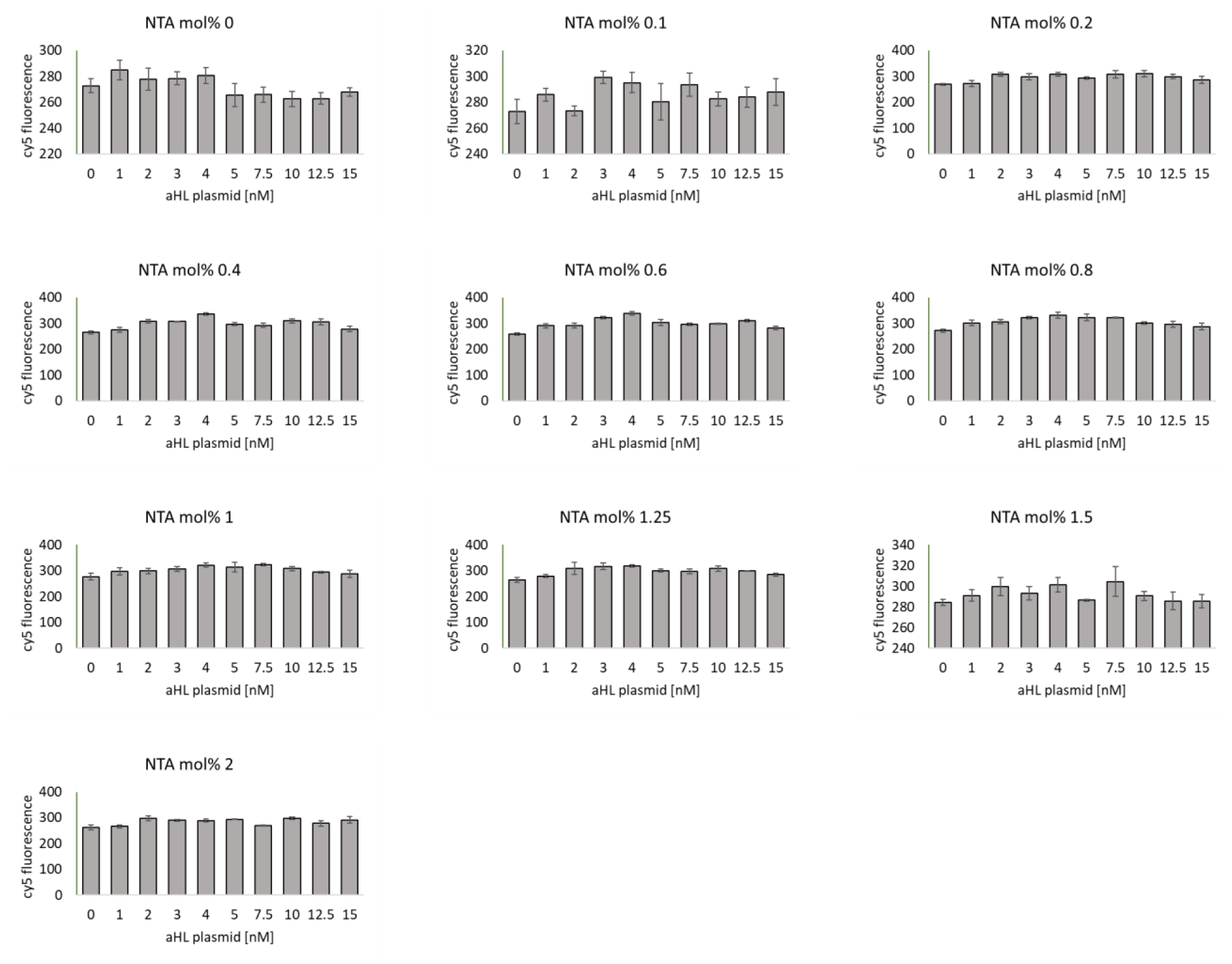
Fusion testing with external αHL, Cy5 signal. Individual Cy5 FRET fluorescence data points for the heatmap on **Figure S20**. Error bars indicate SEM, n=3 independently prepared and processed liposome samples. This data was collected using TxTl.

**Figure S23.**
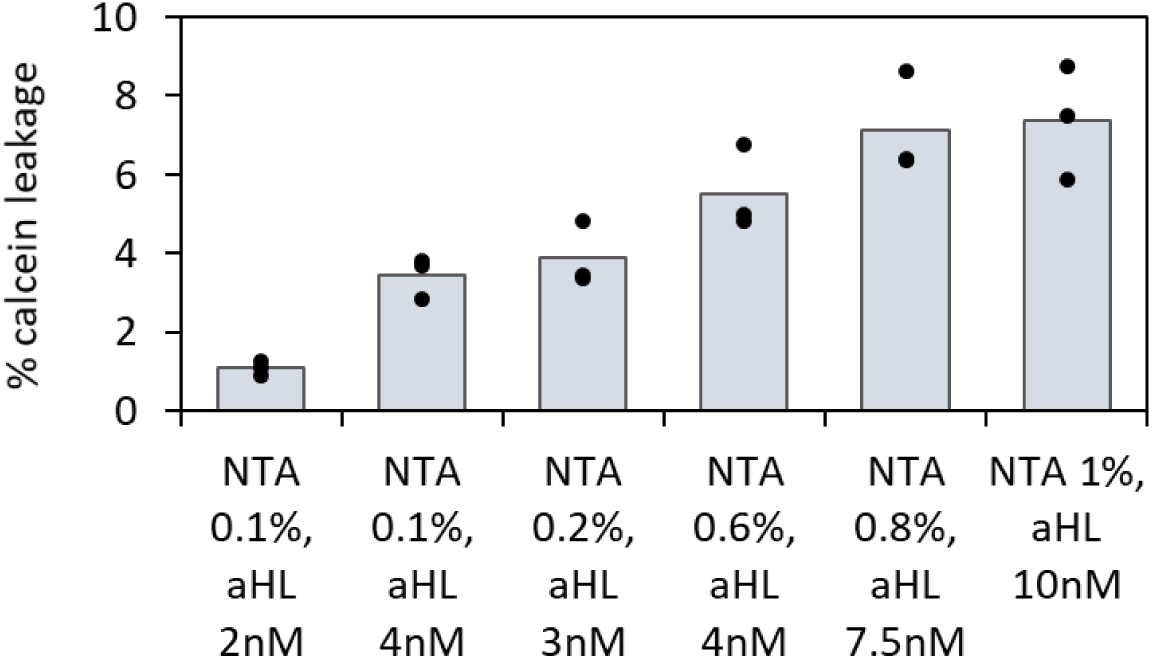
Leakage test for His labeled αHL fusion experiments. Dots on top of the bar graphs indicate individual values of three replicates. Samples shown on this figure are selected from the fusion experiment presented as heatmap data on **Figure S20.** Examples of individual purification plots for samples summarized on this figure are on **Figure S24**. This data was collected using TxTl.

**Figure S24.**
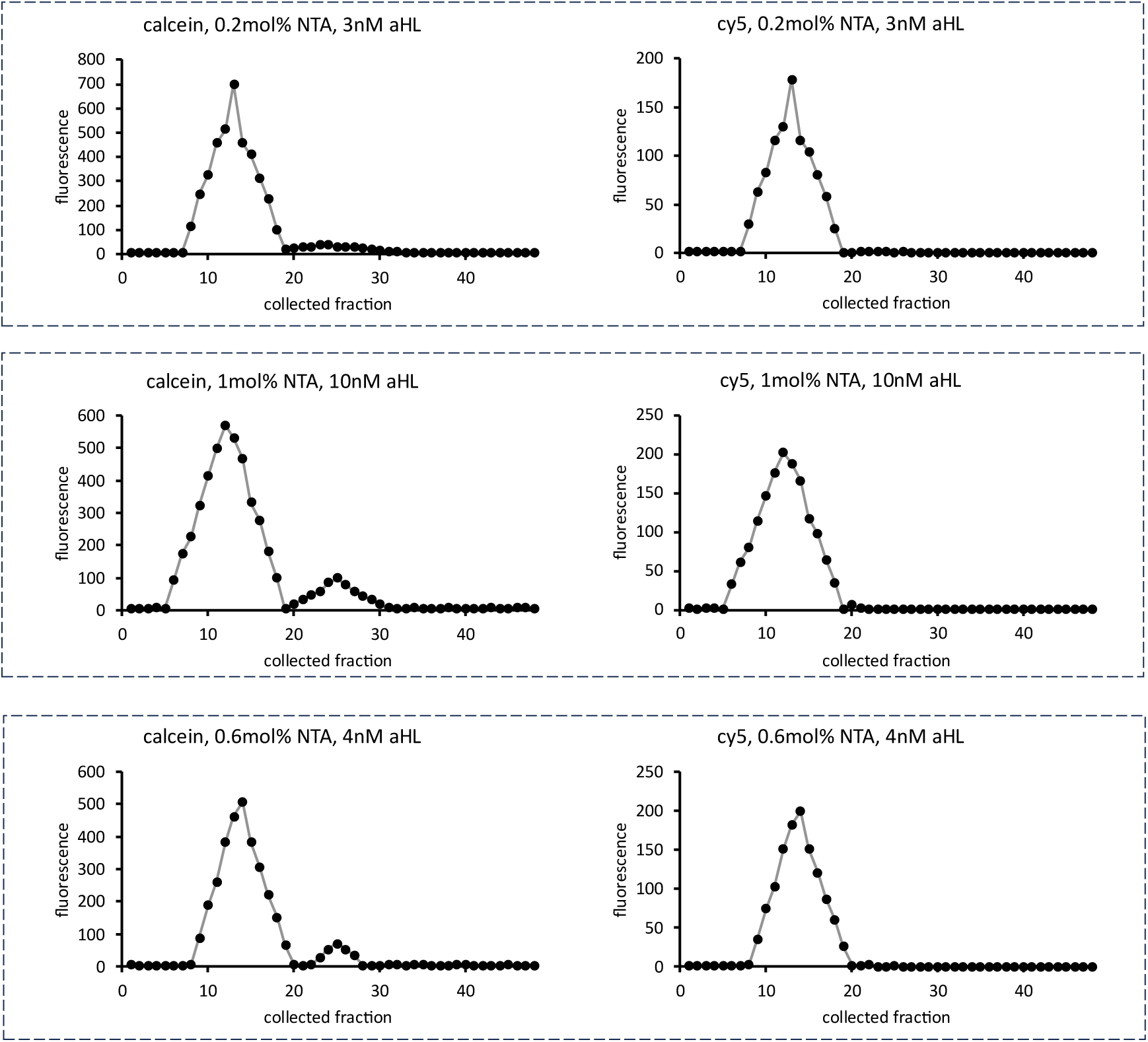
Leakage test for αHL fusion experiments. Example plots from size exclusion chromatography purification of liposomes with varying amount of αHL plasmid, and varying amount of NTA lipid as mol% of total lipids. Leakage of small molecule calcein was tested after 12h incubation at 30°C. Liposomes contained 1mM calcein at the beginning of the experiment. Each sample was analyzed using two fluorescent measurements: calcein channel and Cy5 channel (see **Table S5** for wavelengths). The calcein channel traces indicate leakage of small molecule and the Cy5 channel (Cy5 lipid dye 1,2-dioleoyl-sn-glycero-3-phosphoethanolamine-N-(Cyanine 5)) indicates which fractions contain liposomes. This data was collected using TxTl.

**Figure S25.**
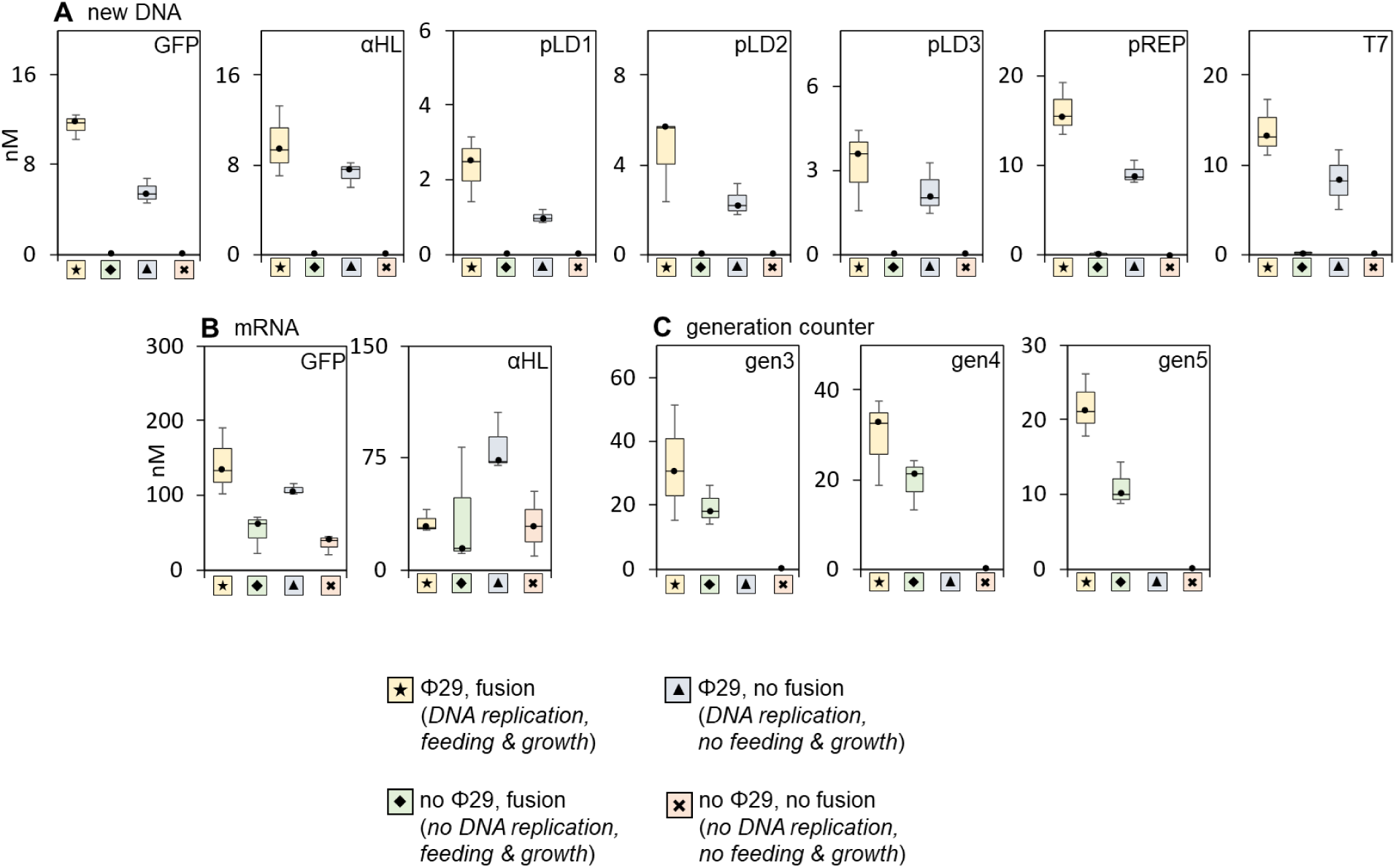
Synthetic cells after 5 generations of growth and division with dialysis removing feeder liposomes. The experiments were performed according to the same protocol as experiments on Figure 3, but after each addition of feeder liposomes and 12h incubation at 30°C, samples were dialyzed for 4h against 1μm filter to remove free feeder liposomes. See Methods section 24 for details. Dialysis time course is on **Figure S26**. **A**: qPCR quantification of new DNA, indicating Phi29 genome replication. The samples were treated with DpnI, removing the original plasmid DNA, so only newly synthesized DNA was detectable. **B**: RT qPCR quantification of the mRNA abundance. **C**: quantification of the generation counter, detecting the counter product of the indicated number of generations. Samples were processed with either primer for generation 3 (labeled gen3, detecting three, four, or five generations), primer for generation 4 (labeled gen4, detecting three or four generations), or the primer for generation 5 (labeled gen5, detecting only the results of five generations). All experiments were performed in four separate variants: the complete cell cycle with Phi29 genome replication and with feeder liposome fusion (symbol 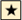); without Phi29 genome replication, but with feeder liposome fusion (symbol 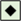); with Phi29 for genome replication, but without feeder liposome fusion (symbol 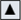); and without either Phi29 genome replication or feeding (symbol 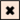). Individual data points for the box and whiskers plots are on **Figure S27**. This data was collected using PURE pREP.

**Figure S26.**
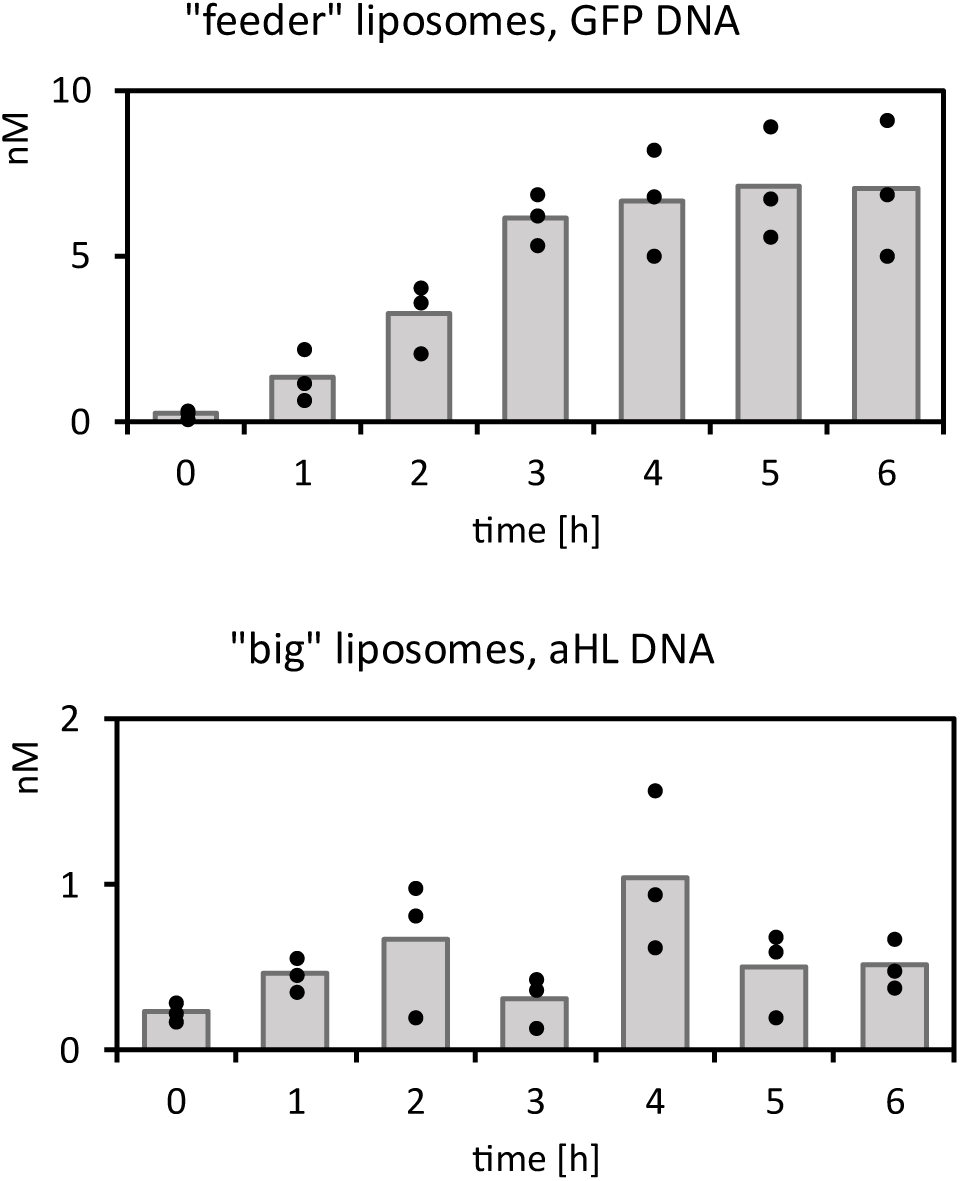
Dialysis time course of removing “feeder” liposomes. The model “feeder” 0.4μm liposomes were prepared with 10nM GFP plasmid (no transcription and translation system, just plasmid for quantitative detection). The “big” liposomes were prepared to model “parent” synthetic cells in the growth and feeding experiments, with average diameter of 2-3uM, those liposomes were filled with 10nM αHL plasmid (no transcription and translation system, just plasmid for quantitative detection). Samples were dialyzed for amount of time indicated on X axis, with time 0 being a sample mixed in dialyzer and immediately collected for analysis. Data shows qPCR abundance of GFP and αHL plasmids in the external dialysis buffer (showing material removed from the original sample). Bar graphs represent average value, dots are individual data points for each replicate. This data was collected using PURE pREP.

**Figure S27.**
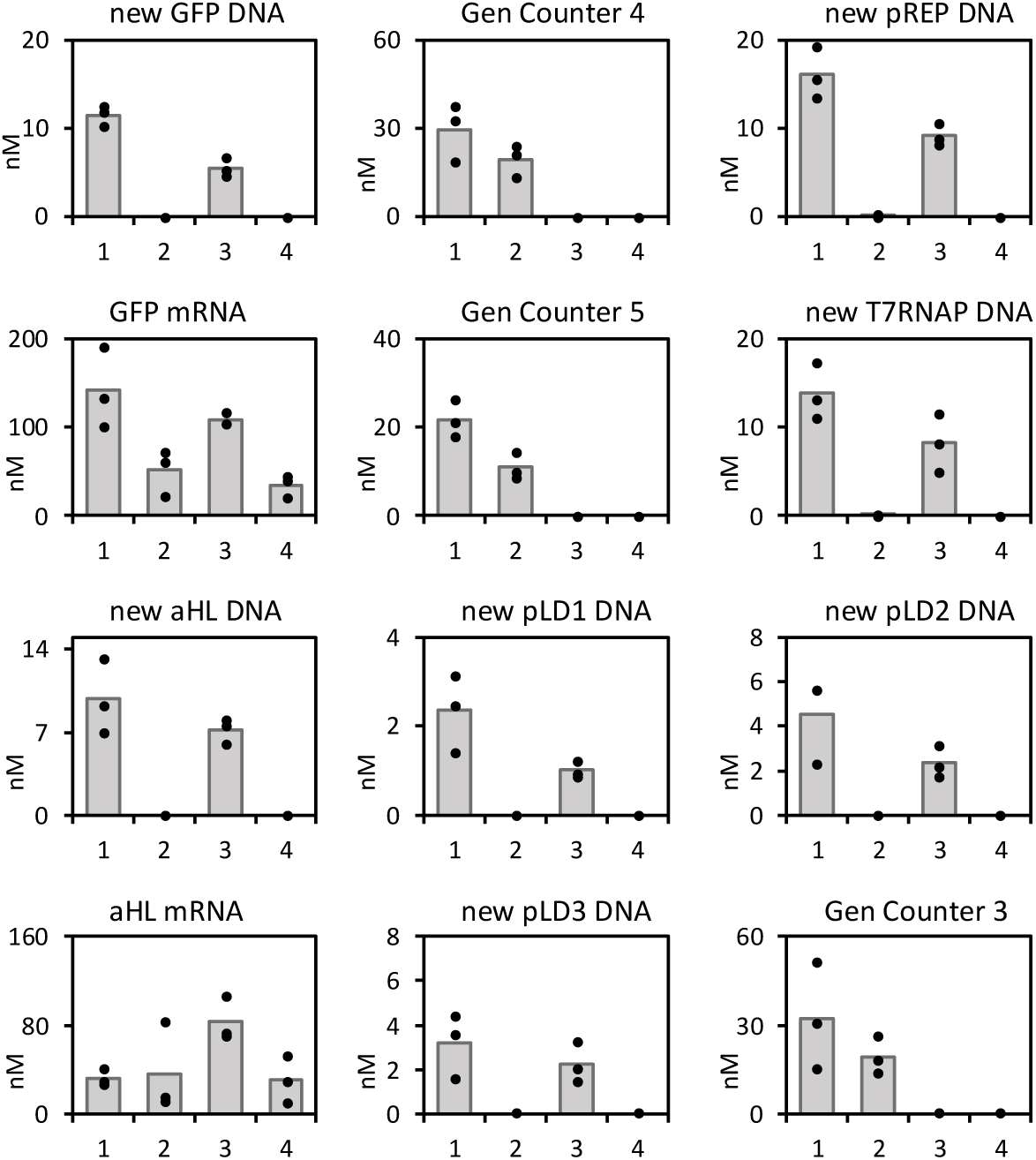
5 Generations of growth and division with dialysis. Individual data points for box and whiskers plots on **Figure S25**, analysis of plasmids, mRNA and generation counter oligo after 5 generations of growth and division with dialysis after each generation. All experiments were performed in four separate variants: the complete cell cycle with Phi29 genome replication and with feeder liposome fusion (symbol 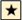, labeled sample 1 on this figure); without Phi29 genome replication, but with feeder liposome fusion (symbol 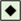, labeled sample 2 on this figure); with Phi29 for genome replication, but without feeder liposome fusion (symbol 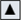, labeled sample 3 on this figure); and without either Phi29 genome replication or feeding (symbol 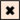, labeled sample 4 on this figure). <u>Sample 1</u>: Phi29, fusion (symbol 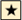 on Figure 3), Generation 5 <u>Sample 2</u>: No Phi29, fusion (symbol 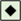 on Figure 3), Generation 5 <u>Sample 3</u>: Phi29, No Fusion (symbol 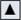 on Figure 3), Generation 5 <u>Sample 4</u>: No Phi29, No Fusion (symbol 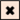 on Figure 3), Generation 5 Bar graphs represent average value, dots are individual data points for each replicate. This data was collected using PURE pREP.

**Figure S28.**
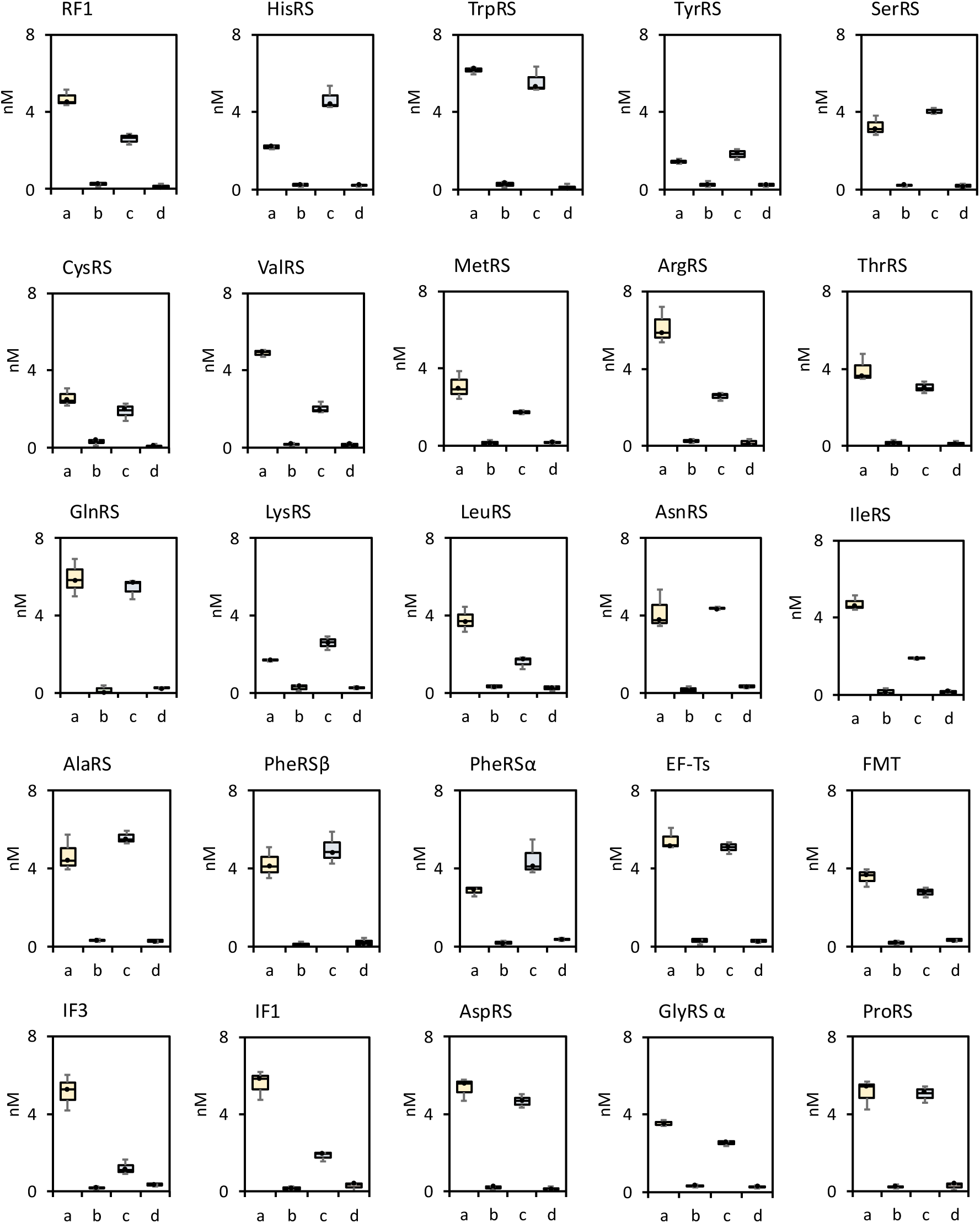
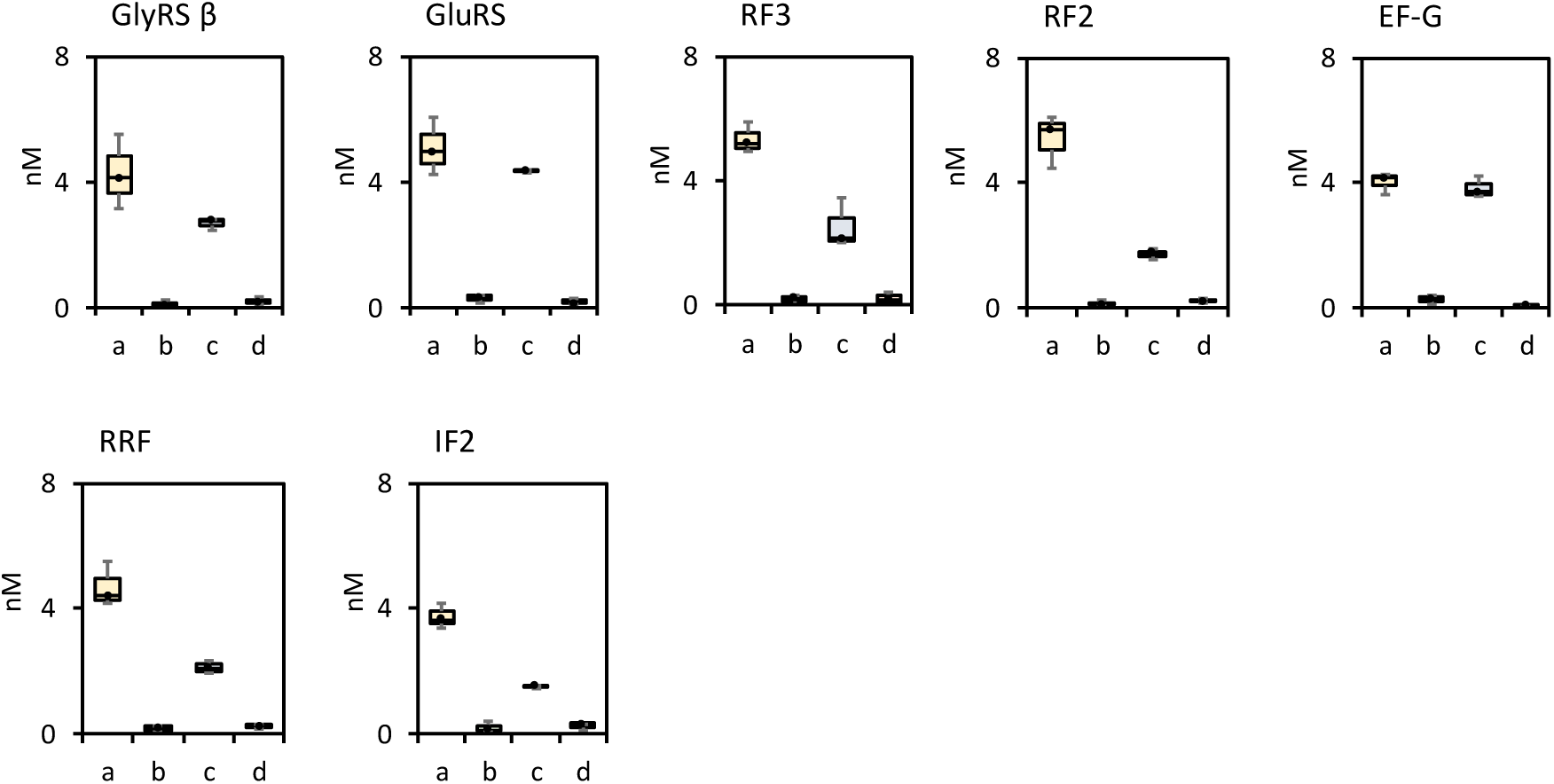
All genes in 90kB synthetic cell genome after 5 generations of cell cycle. Quantification of all genes in the synthetic cell genome undergoing 5 generations of cell cycle (data for genes not included on the Figure 3). For each gene, 4 individual experimental conditions were tested, labeled a, b, c and d on the graphs. <u>sample a</u>: the complete cell cycle with Phi29 genome replication and with feeder liposome fusion (labeled with symbol 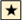 on the main figures); <u>sample b</u>: without Phi29 genome replication, but with feeder liposome fusion (labeled with symbol 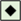 on the main figures); <u>sample c</u>: with Phi29 for genome replication, but without feeder liposome fusion (labeled with symbol 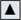 on the main figures); <u>sample d</u>: and without either Phi29 genome replication or feeding (labeled with symbol 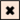 on the main figures). Individual data points for all data on this figure are on **Figure S29**. This data was collected using PURE pREP. ***Related data***: abundance of each individual gene in the sample after 5 generations is on **Figure S28** (with individual data points on **Figure S29**), abundance of each plasmid in single cells is on **Figure S32**, count of cells expressing GFP after each cycle is on **Figure S30**, and abundance of full length generation counter oligo in single cells after each generation is on **Figure S37**.

**Figure S29.**
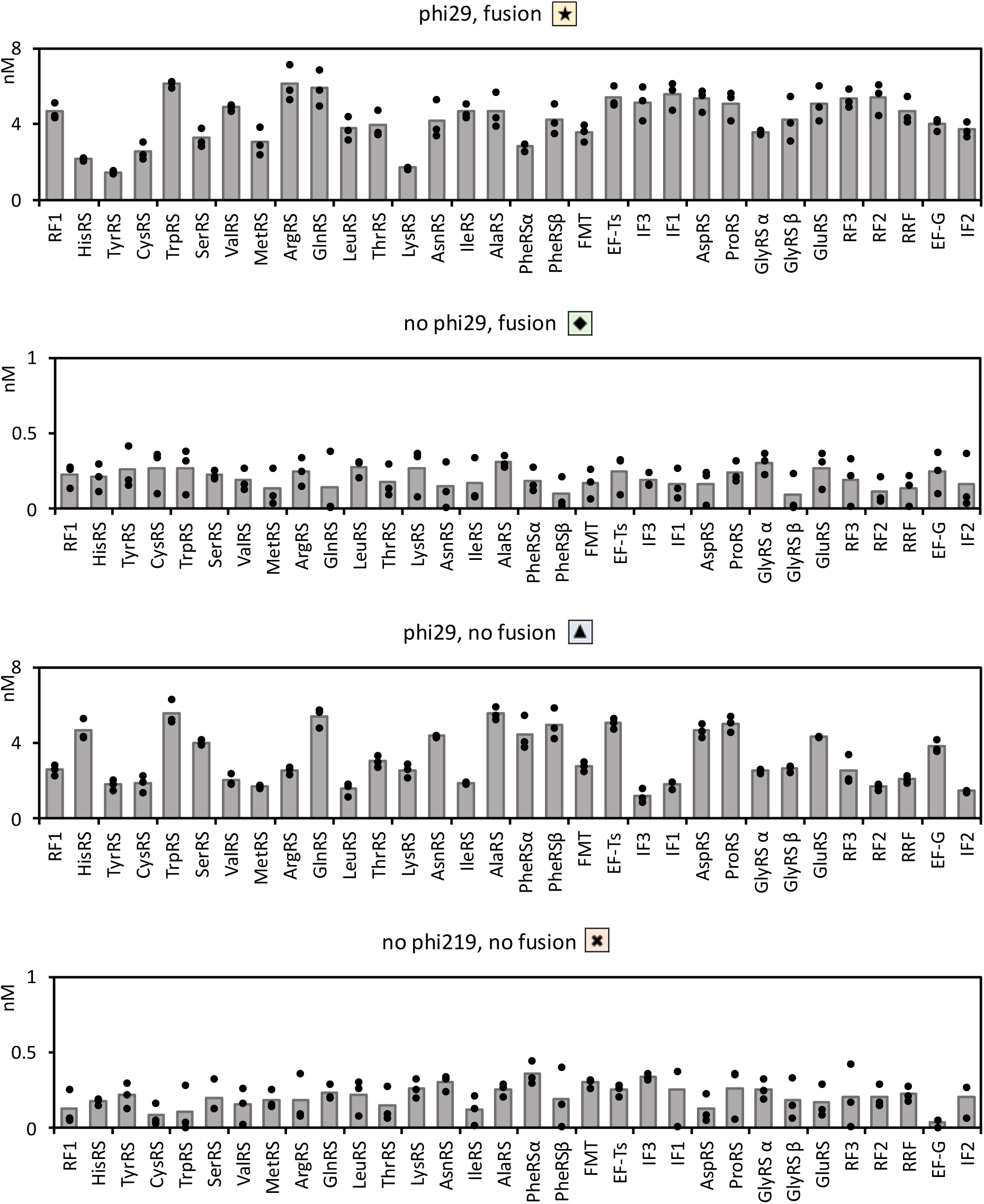
Quantification of individual genes in the synthetic cell genome after 5 generations of cell cycle. Individual data points for qPCR data of quantification of all genes in the synthetic cell genome undergoing 5 generations of cell cycle from **Figure S28**. For each gene, 4 individual experimental conditions were tested: <u>Phi 29, fusion</u>: the complete cell cycle with Phi29 genome replication and with feeder liposome fusion, labeled with symbol 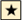 on the main figures, and labeled “a” in box and whiskers plots on **Figure S28**. <u>no Phi29, fusion</u>: without Phi29 genome replication, but with feeder liposome fusion, labeled with symbol 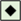 on the main figures, and labeled “b” in box and whiskers plots on **Figure S28**. <u>Phi 29, no fusion</u>: with Phi29 for genome replication, but without feeder liposome fusion (labeled with symbol 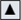 on the main figures, and labeled “c” in box and whiskers plots on **Figure S28**. <u>no Phi 29, no fusion</u>: and without either Phi29 genome replication or feeding (labeled with symbol 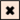 on the main figures, and labeled “d” in box and whiskers plots on **Figure S28**. Dots on top of the bars indicate all the individual data points, the bars are the average value. This data was collected using PURE pREP. ***Related data***: abundance of each individual gene in the sample after 5 generations is on **Figure S28** (with individual data points on **Figure S29**), abundance of each plasmid in single cells is on **Figure S32**, count of cells expressing GFP after each cycle is on **Figure S30**, and abundance of full length generation counter oligo in single cells after each generation is on **Figure S37**.

**Figure S30.**
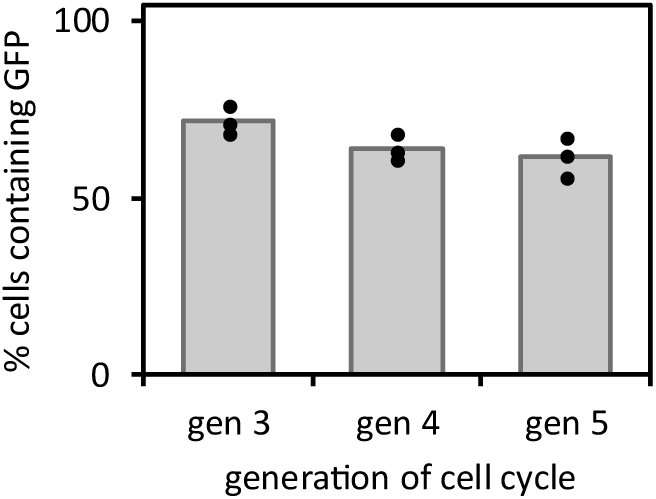
GFP fluorescence in synthetic cells. Flow cytometry measurements indicating what percent of total synthetic cell population contains detectable amount of GFP, measured after 3, 4 and 5 cycles of Phi29 genome replication and with feeder liposome fusion (labeled with symbol 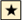 on the main figures). See Methods section 17 “Flow cytometry” for measurement protocol. For each measurement, experiments were performed after the extrusion step, and samples were dialyzed to remove feeder vesicles. Dots on top of the bars indicate all the individual data points, the bars are the average value. This data was collected using PURE pREP. ***Related data***: abundance of each individual gene in the sample after 5 generations is on **Figure S28** (with individual data points on **Figure S29**), abundance of each plasmid in single cells is on **Figure S32**, count of cells expressing GFP after each cycle is on **Figure S30**, and abundance of full length generation counter oligo in single cells after each generation is on **Figure S37**.

**Figure S31.**
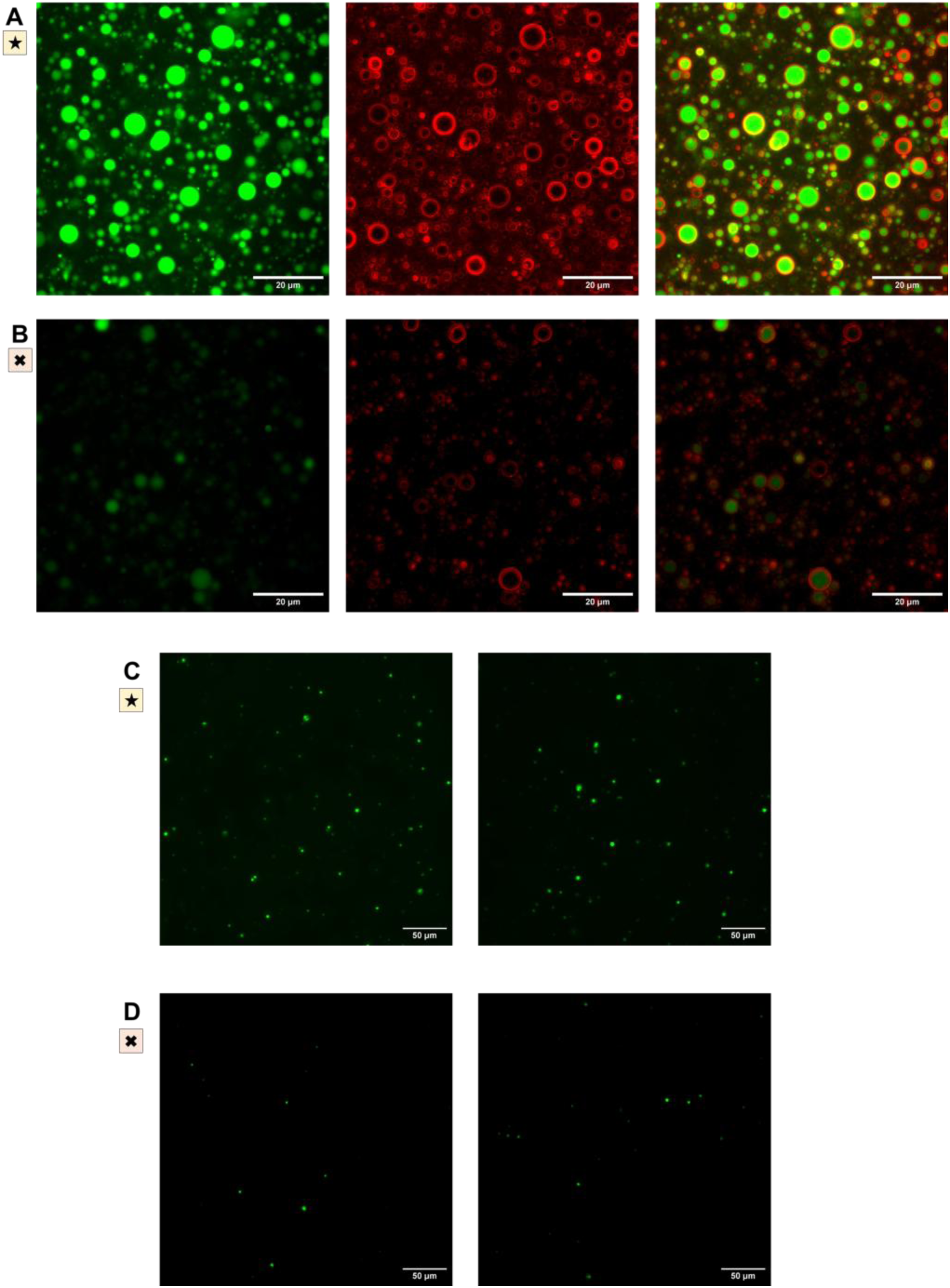
Fluorescence microscopy of synthetic cell samples, imaged after five generations. The green channel is GFP fluorescence, and the red channel is rhodamine membrane dye. Samples are labeled according to the conditions of the cell cycle experiment: the complete cell cycle with Phi29 genome replication and with feeder liposomes (symbol 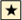); and without either Phi29 genome replication or feeding (symbol 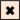). **A**: The complete cell cycle with Phi29 genome replication and with feeder liposomes. The red and green channels and the composite for the image shown on Figure 3M. Scale bar is 20µm. **B**: Samples without either Phi29 genome replication or feeding. The red and green channels and the composite for the image shown on Figure 3N. Scale bar is 20µm. **C**: Two fields of view from lower concentration imaging, samples with the complete cell cycle with Phi29 genome replication and with feeder liposomes. Scale bar is 50µm. **D**: Two fields of view from lower concentration imaging, samples without either Phi29 genome replication or feeding. Scale bar is 50µm. Panels A and B: The samples were imaged at the highest concentration possible, without diluting from the cell cycle experiments (applied in the middle of a cover slip for immediate imaging). Scale bar is 20µm. Panels **C** and **D**: The samples were diluted 100x for imaging. Scale bar is 50µm. This data was collected using PURE pREP.

**Figure S32.**
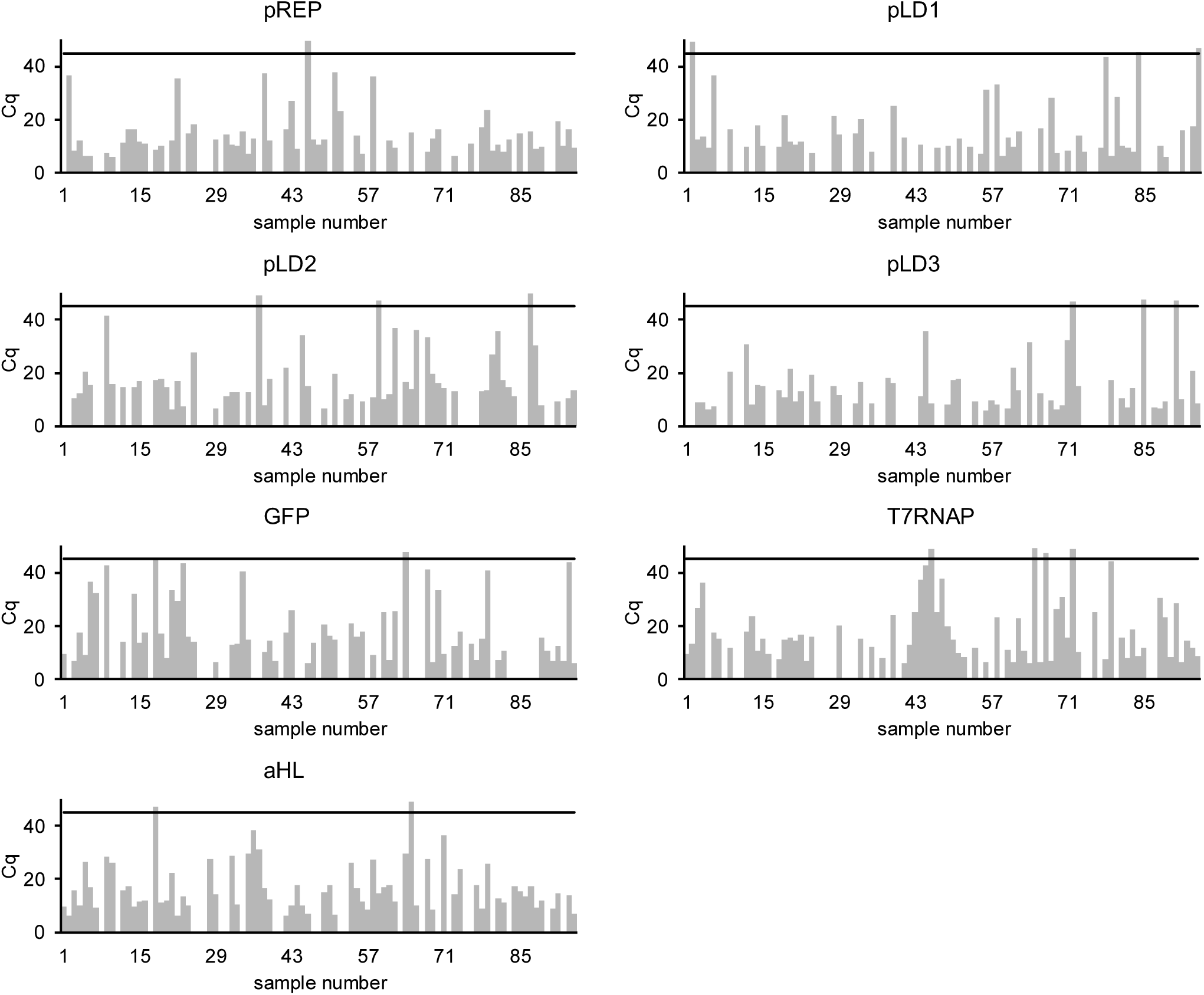
Abundance of each plasmid in single cell experiments after 5 generations of cell cycle. Single cell amplification and qPCR analysis of abundance of each of the individual plasmids (see **Table S6** for the list of plasmids) in a synthetic cell genome, after 5 cycles of growth and replication (condition labeled with symbol 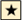 through Figure 3). Black line shows threshold of detection (Cq below 45), plasmid was not considered present if Cq value was above that number. This data was collected using PURE pREP. ***Related data***: abundance of each individual gene in the sample after 5 generations is on **Figure S28** (with individual data points on **Figure S29**), abundance of each plasmid in single cells is on **Figure S32**, count of cells expressing GFP after each cycle is on **Figure S30**, and abundance of full length generation counter oligo in single cells after each generation is on **Figure S37**.

**Figure S33.**
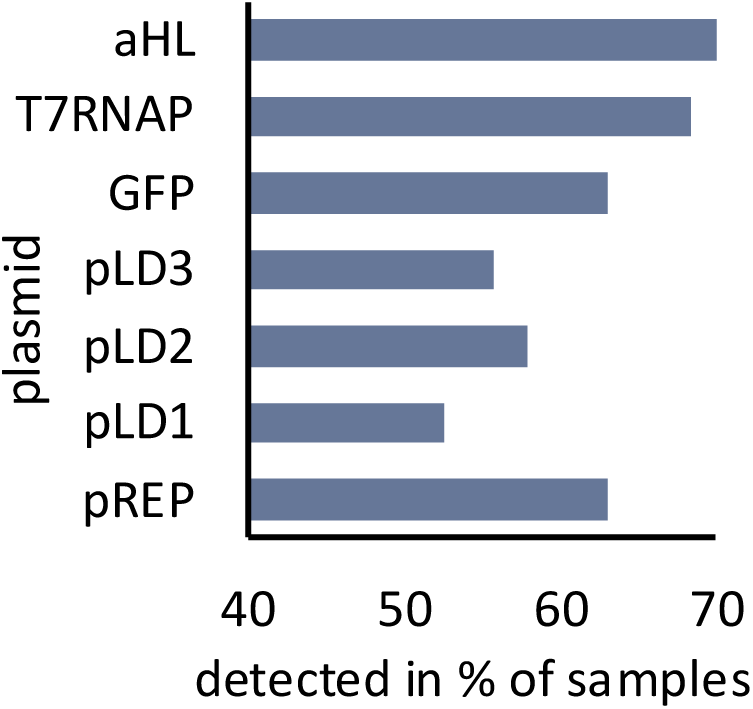
Summary of single cell analysis of abundance of each plasmid. Analysis was done on samples after 5 cycles of growth and division. Total of 95 samples were analyzed. This is summary of individual cell data shown on **Figure S32**. This data was collected using PURE pREP.

**Figure S34.**
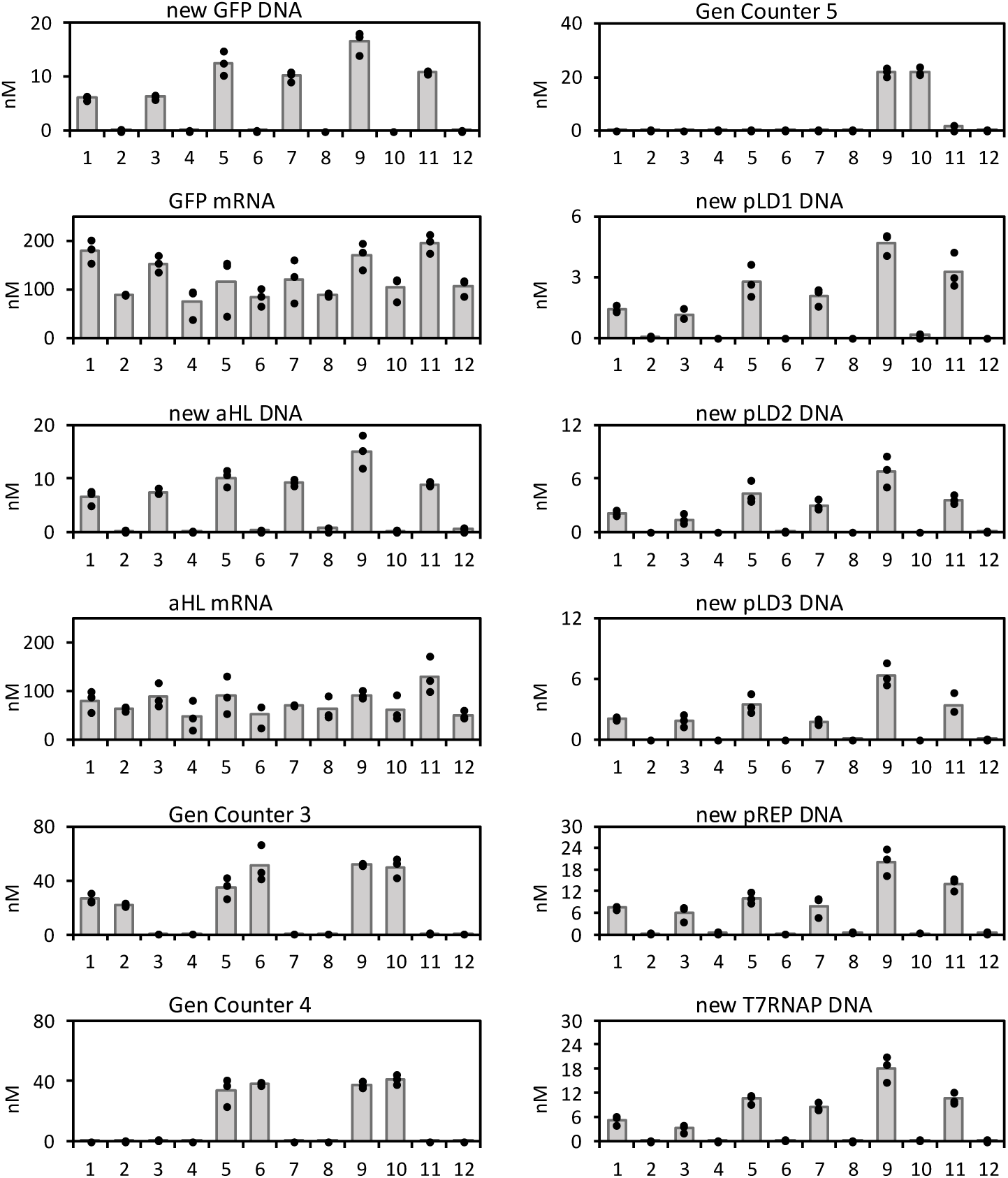
Individual data points for 5 generations of growth and division. Individual data points for data shown on box and whiskers plots on Figure 3. All experiments were performed in four separate variants: the complete cell cycle with Phi29 genome replication and with feeder liposome fusion (symbol 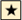, labeled samples 1, 5 and 9 on this figure); without Phi29 genome replication, but with feeder liposome fusion (symbol 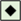, labeled samples 2, 6 and 10 on this figure); with Phi29 for genome replication, but without feeder liposome fusion (symbol 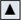, labeled samples 3, 7 and 11 on this figure); and without either Phi29 genome replication or feeding (symbol 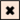, labeled samples 4, 8 and 12 on this figure). **Statistical significance analysis** showing P values is in **Table S9**. <u>Sample 1</u>: Phi29, fusion (symbol 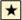 on Figure 3), Generation 3 <u>Sample 2</u>: No Phi29, fusion (symbol 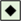 on Figure 3), Generation 3 <u>Sample 3</u>: Phi29, No Fusion (symbol 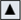 on Figure 3), Generation 3 <u>Sample 4</u>: No Phi29, No Fusion (symbol 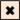 on Figure 3), Generation 3 <u>Sample 5</u>: Phi29, fusion (symbol 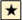 on Figure 3), Generation 4 <u>Sample 6</u>: No Phi29, fusion (symbol 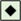 on Figure 3), Generation 4 <u>Sample 7</u>: Phi29, No Fusion (symbol 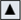 on Figure 3), Generation 4 <u>Sample 8</u>: No Phi29, No Fusion (symbol 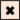 on Figure 3), Generation 4 <u>Sample 9</u>: Phi29, fusion (symbol 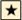 on Figure 3), Generation 5 <u>Sample 10</u>: No Phi29, fusion (symbol 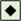 on Figure 3), Generation 5 <u>Sample 11</u>: Phi29, No Fusion (symbol 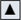 on Figure 3), Generation 5 <u>Sample 12</u>: No Phi29, No Fusion (symbol 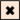 on Figure 3), Generation 5 Bar graphs represent average value, dots are individual data points for each replicate. This data was collected using PURE pREP.

**Figure S35.**
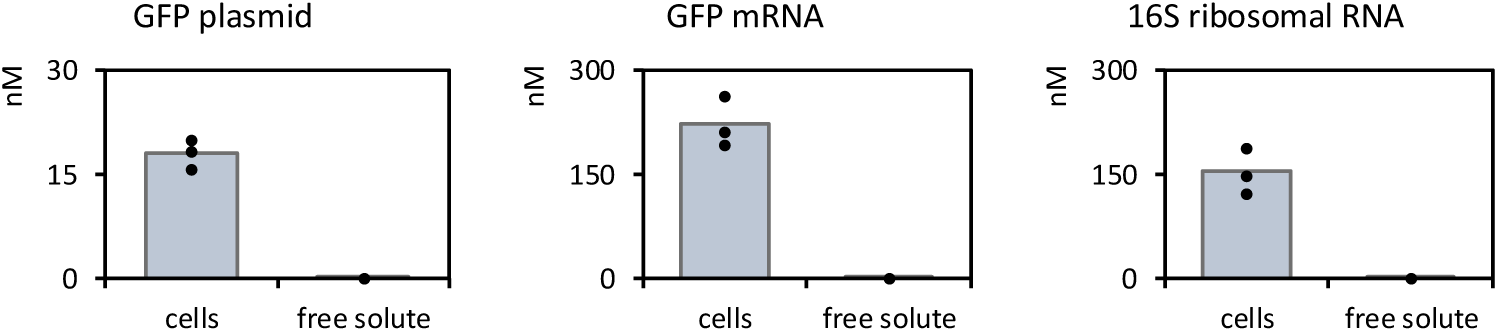
Detection of component leakage from cells after 5 generations of growth and division. Cells undergo 5 generations of complete growth and replication, the complete cell cycle with Phi29 genome replication and with feeder liposomes (symbol 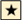). Cells were prepared as described before (Methods section 11). After 5 cycles of growth and replication, and final extrusion, cells were purified on size exclusion chromatography (Methods section 27), then both free solute fraction and the synthetic cell fractions were collected. The fractions were concentrated by lyophilization and resuspended to total volume of 10uL, and processed through a qPCR analysis (Methods section 14) to quantify amount of the GFP plasmid, the GFP mRNA, and the 16S ribosomal RNA. It is important to note that the concentrations reported here do not represent actual concentrations in the original cell sample. The cells were diluted during the column purification and re-concentrated after collecting fractions. This is a qualitative detection test, to check the integrity of the cells (leakage of nucleic acids and ribosomes) after cycles of growth and replication. Due to the low total concentration of GFP in the collected cell fractions and no detectable GFP signal in free solute fraction, fractions containing cells were collected mostly based on the characteristic meniscus appearance of the wells containing cells, and free solute fraction was collected to very generously encompass all fraction from the end of the cell fraction till the end of the run. Results shown on this figure show plasmid abundance analysis for plasmid, mRNA and ribosomal RNA in the synthetic cell fraction and in the free solute fraction. This data was collected using PURE pREP.

**Figure S36.**
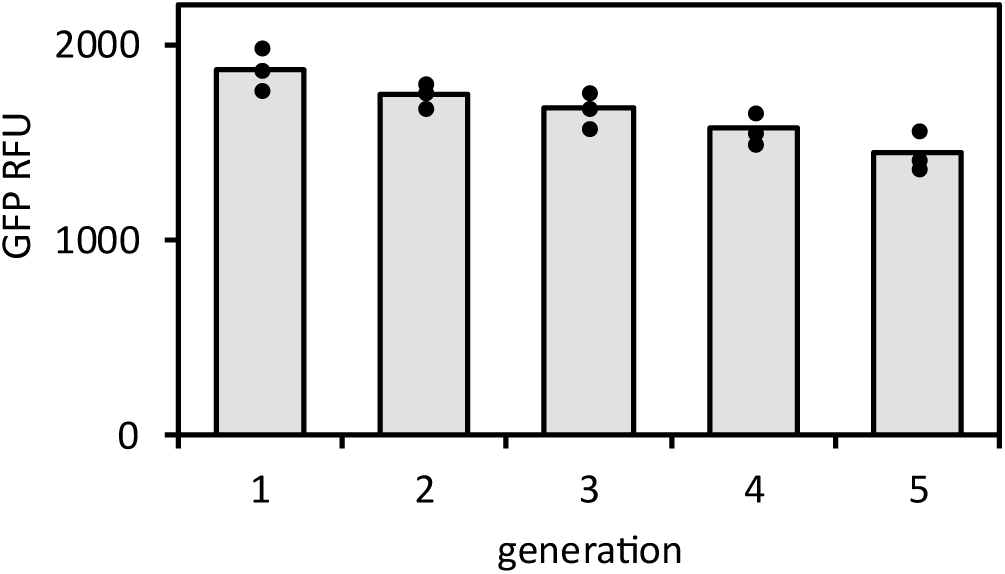
Fusion efficiency across multiple generations. The starting synthetic cells contained T7 RNA polymerase and TxTl machinery expressing αHL, but no GFP plasmid. Generation of growth and division experiments were performed, as described on Figure 3 and in Methods section 11. The “feeder vesicles” were being added, containing all metabolites as described in Methods section 11, and at a generation indicated for testing also contained a GFP plasmid. Cells were dialyzed, normalized to the same cell density (see Supplementary text section 5), and GFP fluorescence was measured to indicate fusion efficiency. Dots on top of the bars indicate all the individual data points, the bars are the average value. This data was collected using PURE pREP.

**Figure S37.**
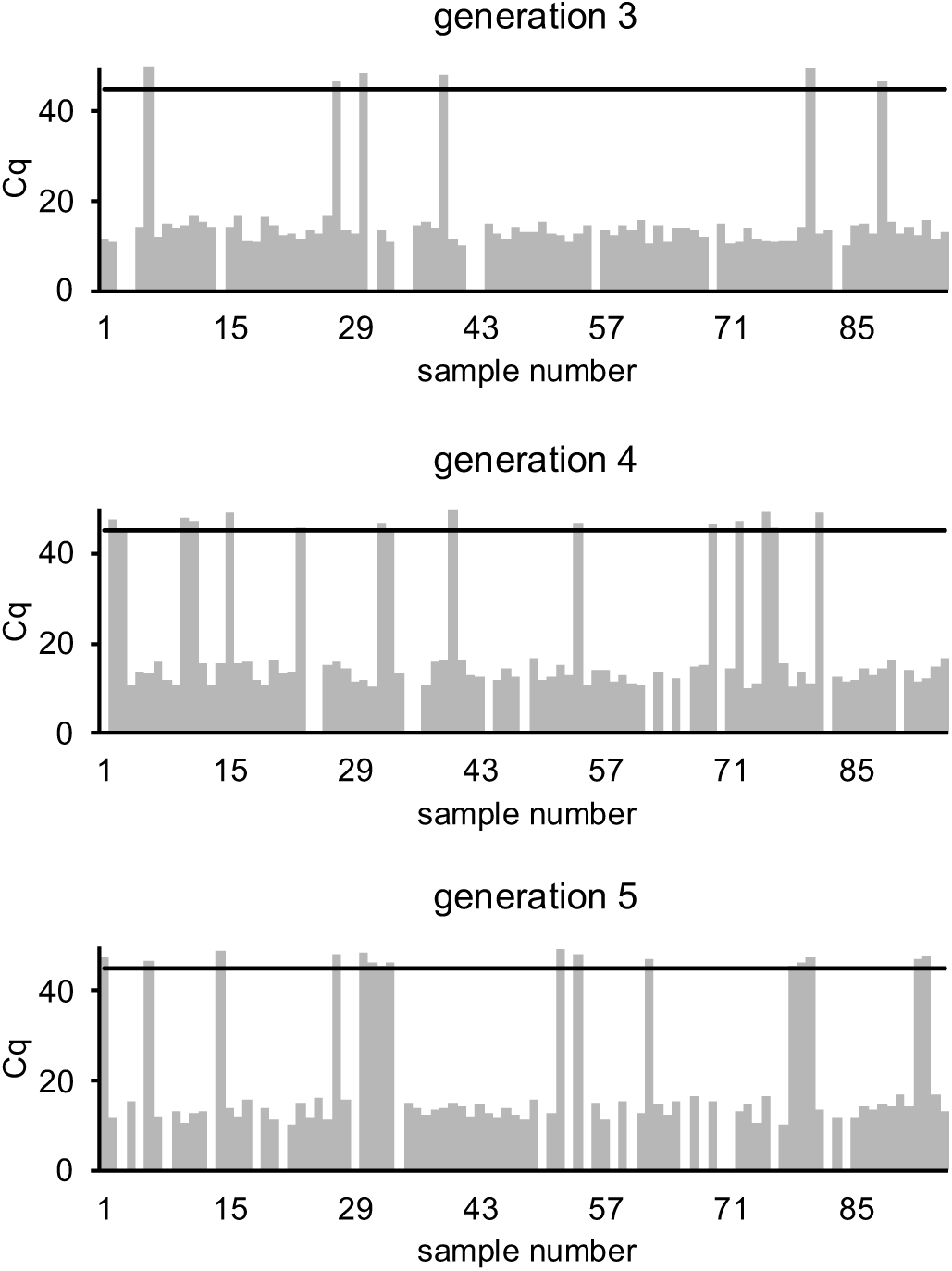
Full length generation counter oligo after each generation. Single cell amplification and qPCR analysis of abundance of full length generation counter oligo, after 3, 4 and 5 cycles of growth and replication (condition labeled with symbol 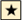 through Figure 3). Black line shows threshold of detection (Cq below 45), oligo was not considered present if Cq value was above that number. In summary, 78 cells collected after generation 3 contained full length generation 3 product, 67 cells collected after generation 4 contained full length generation 4 product, 59 cells collected after generation 5 contained full length generation 5 product. This data was collected using PURE pREP. ***Related data***: abundance of each individual gene in the sample after 5 generations is on **Figure S28** (with individual data points on **Figure S29**), abundance of each plasmid in single cells is on **Figure S32**, count of cells expressing GFP after each cycle is on **Figure S30**, and abundance of full-length generation counter oligo in single cells after each generation is on **Figure S37**.

**Figure S38.**
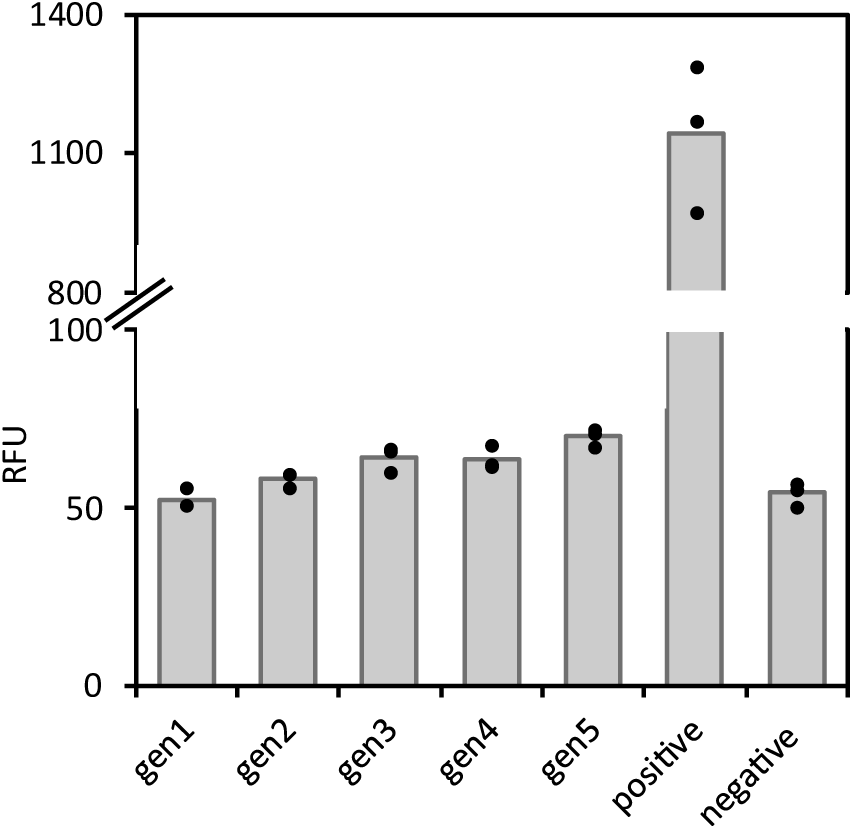
Investigating fusion between synthetic cells. This experiment investigates the possibility of fusion between two synthetic cells (not between synthetic cell and feeder vesicles). Two populations of synthetic cells were prepared. The cells contained a genome comprised of T7RNAP, αHL. One population of cells carried GFP plasmid under control of T RNA polymerase promoter, while the other population contained T3 RNA polymerase gene under control of T7 promoter. Expression of GFP was only possible if the cells containing T3 RNAP fused with cells containing the GFP plasmid. The two populations of synthetic cells (one carrying T3RNAP and one carrying GFP plasmid under T3 promoter) were mixed at the start of the experiment, and generations of growth and division experiments were performed as described earlier. After each generation, GFP fluorescence was measured, to monitor possible fusion between synthetic cells (not fusion between synthetic cell and feeder liposome). Sample labels: gen1, gen2, gen3, gen4, gen5 - samples analyzed after indicated generation; positive - a positive control of GFP fluorescence from GFP under T7 RNAP in synthetic cells that did undergo all 5 cycles of growth and division; negative – synthetic cells after 5 cycles of growth and division without any GFP plasmid. Dots on top of the bars indicate all the individual data points, the bars are the average value. This data was collected using PURE pREP.

**Figure S39.**
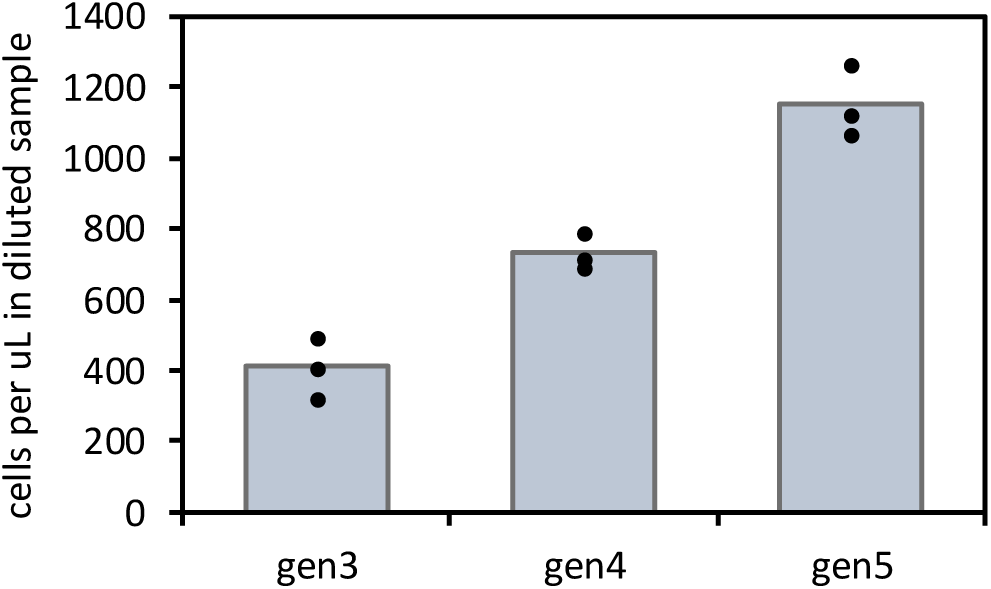
Cell number changes during cell cycle and replication. After each experiment of complete cell cycle (feeding, DNA replication, division, condition labeled with symbol 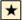 on Figure 3), at the end of the generation (after extrusion, before next feeding), cells were diluted 100,000x (5 serial dilutions) and loaded into a hemocytometer. In each count, cells in three of the 9 large hemocytometer squares were counted. For each experiment, aliquot was prepared from three replicates. Values reported in this figure show average number of cells per 1ul in the diluted sample. This data was collected using PURE pREP.

**Figure S40.**
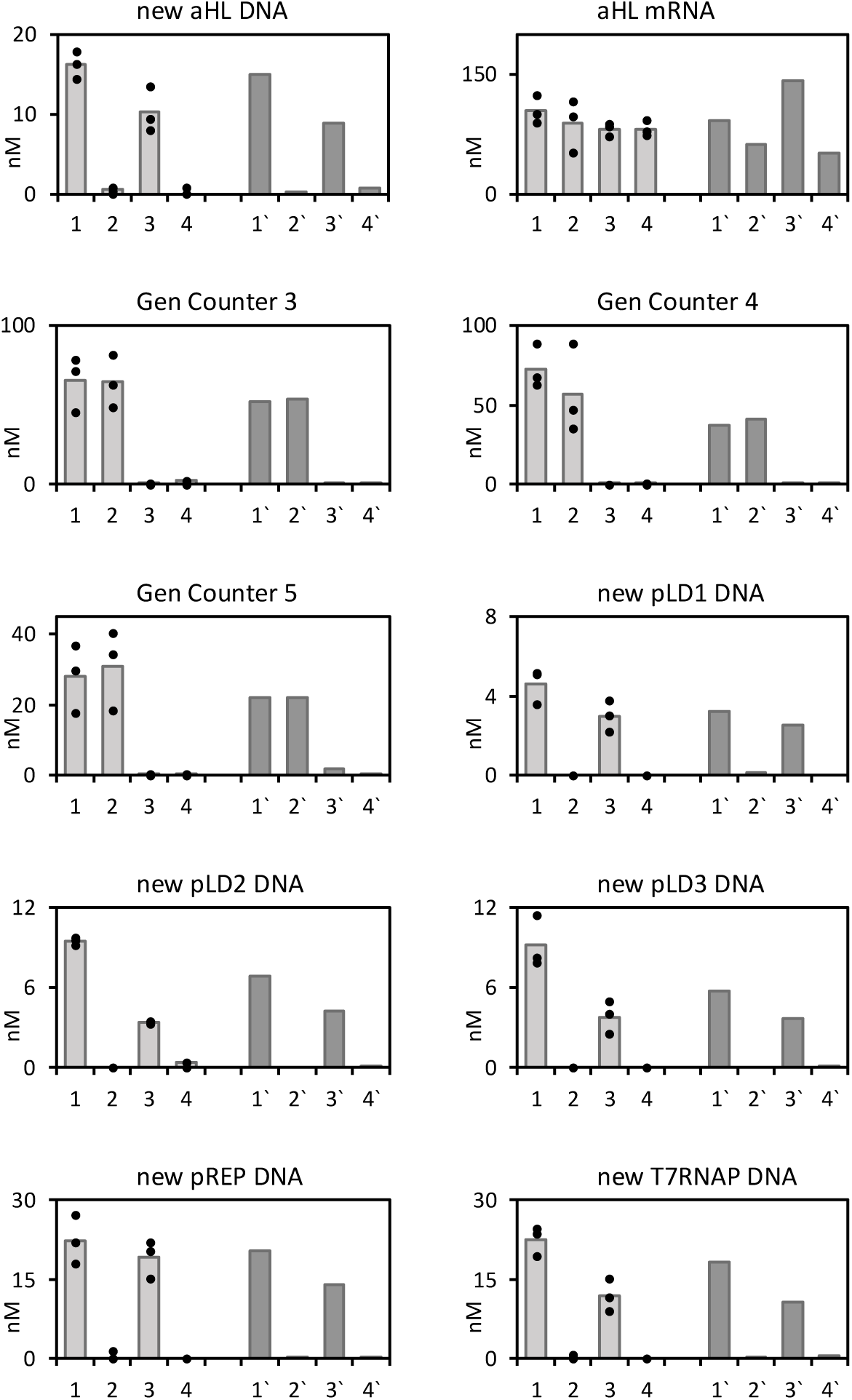
Cell cycle without fluorescent proteins. Results of 5 generations of growth and division experiments, without fluorescent proteins. This experiment demonstrates that the metabolic load from expressing the reporter fluorescent proteins do not significantly affect genome replication through the cell cycle. For easy comparison, the new results (lighter gray) are compared on each graph with results of previous cell cycle experiments where samples contained GFP. Sample labeling: <u>Sample 1</u>: Phi29, fusion (symbol 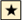on Figure 3), Generation 5 <u>Sample 2</u>: No Phi29, fusion (symbol 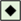 on Figure 3), Generation 5 <u>Sample 3</u>: Phi29, No Fusion (symbol 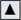 on Figure 3), Generation 5 <u>Sample 4</u>: No Phi29, No Fusion (symbol 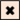 on Figure 3), Generation 5 Darker color, comparison with previous results: <u>Sample 1’</u>: Phi29, fusion (symbol 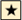 on Figure 3), Generation 5 (results shown on Figure 3) <u>Sample 2’</u>: No Phi29, fusion (symbol 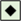 on Figure 3), Generation 5 (results shown on Figure 3) <u>Sample 3’</u>: Phi29, No Fusion (symbol 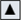 on Figure 3), Generation 5 (results shown on Figure 3) <u>Sample 4’</u>: No Phi29, No Fusion (symbol 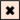 on Figure 3), Generation 5 (results shown on Figure 3) This data was collected using PURE pREP.

**Figure S41.**
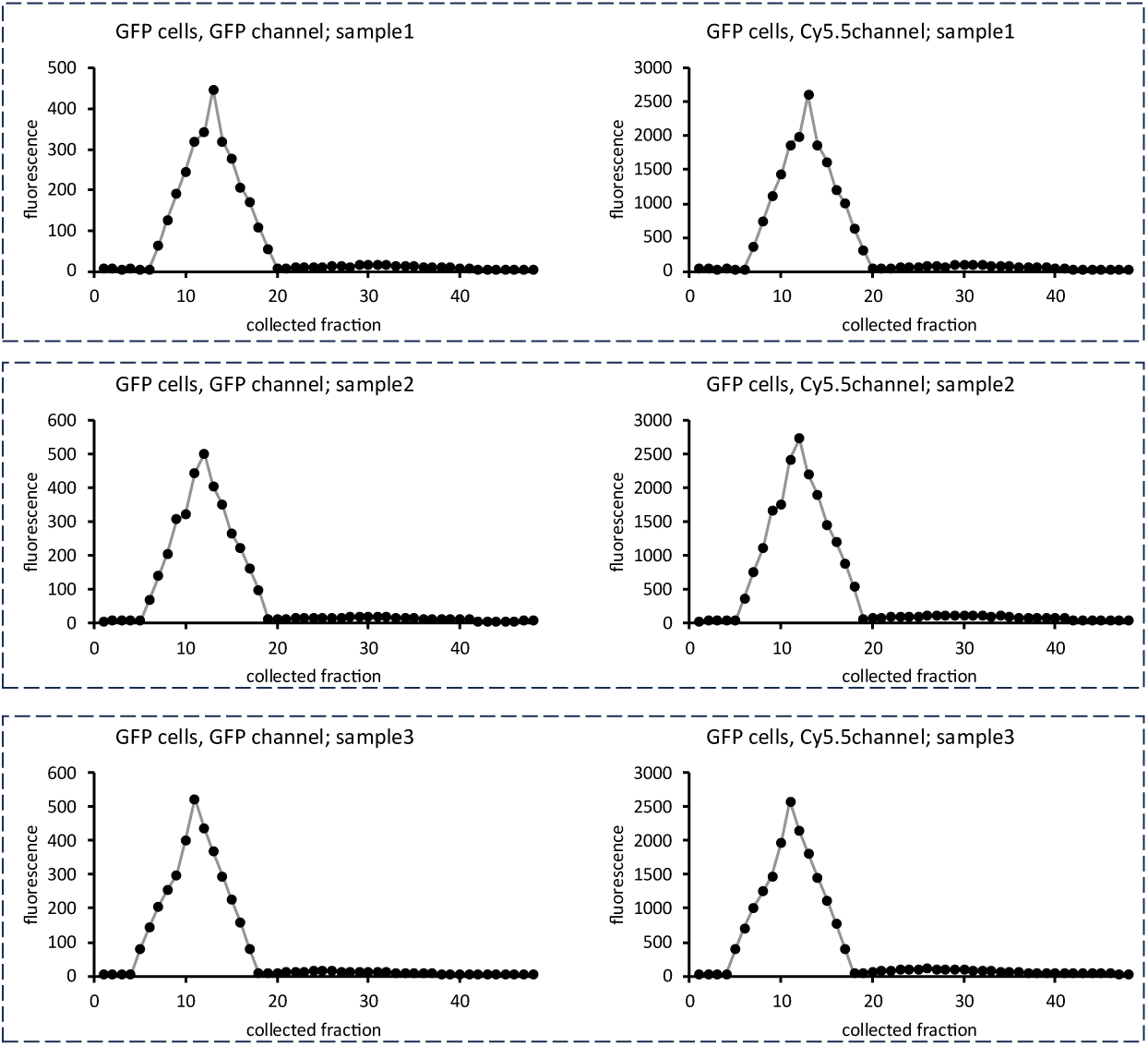
Membrane stability in presence of fluorescent protein GFP. Synthetic cells were created with GFP plasmid. The cells also contained 1mM of Cy5.5. After 12h incubation at 30°C, cells were purified (methods section 27) to measure leakage of the small molecule dye. Each panel shows one purification trace, label indicated type of cells (GFP containing cells on this figure), type of readout of fluorescence (GFP channel or Cy5.5 channel, see **Table S5** for list of wavelengths used), and replicate sample number (3 independent samples). This data was collected using PURE pREP.

**Figure S42.**
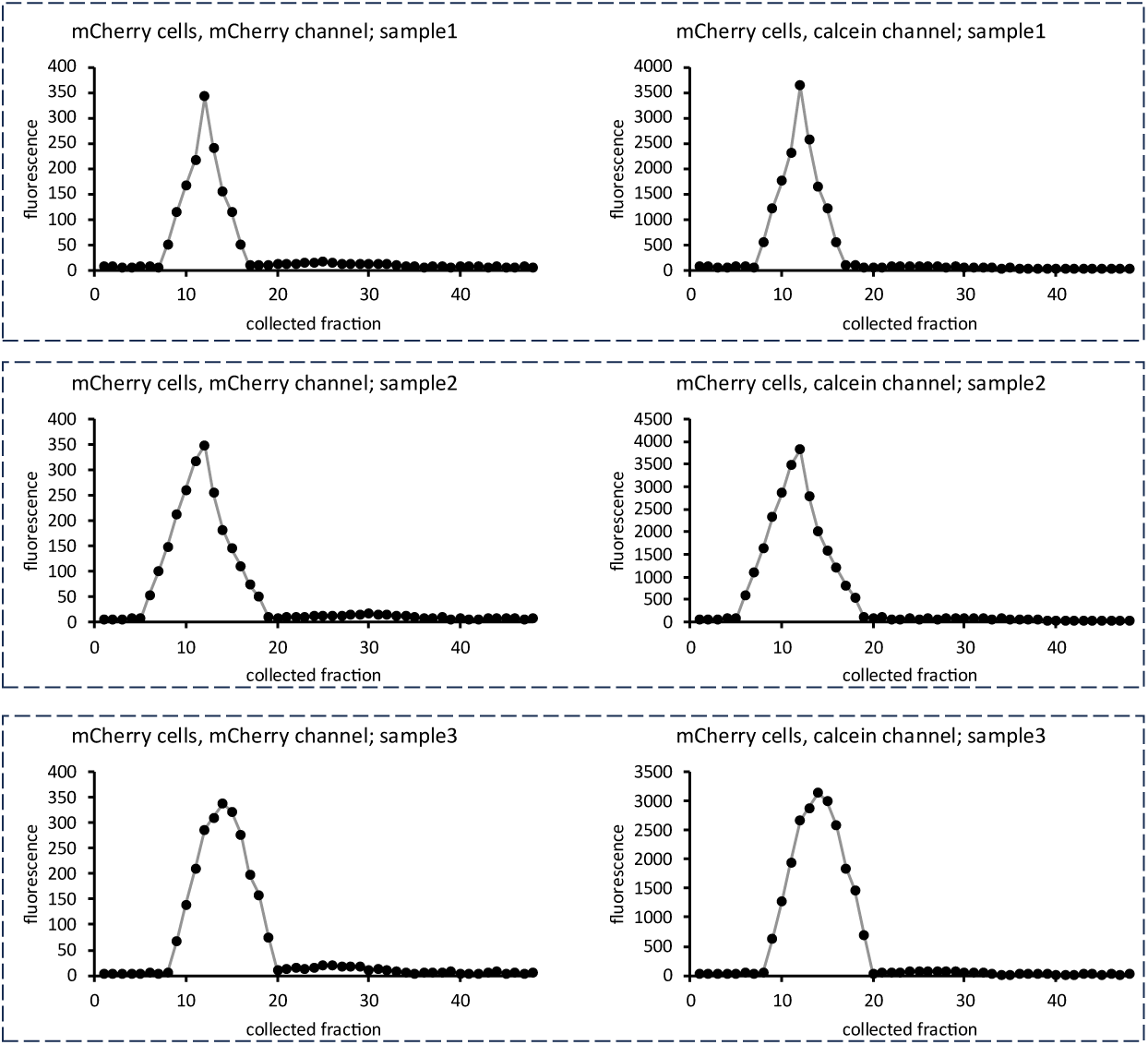
Membrane stability in presence of fluorescent protein mCherry. Synthetic cells were created with mCherry plasmid. The cells also contained 1mM calcein. After 12h incubation at 30°C, cells were purified (methods section 27) to measure leakage of the small molecule dye. Each panel shows one purification trace, label indicated type of cells (mCherry containing cells on this figure), type of readout of fluorescence (mCherry channel or calcein channel, see **Table S5** for list of wavelengths used), and replicate sample number (3 independent samples). This data was collected using PURE pREP.

**Figure S43.**
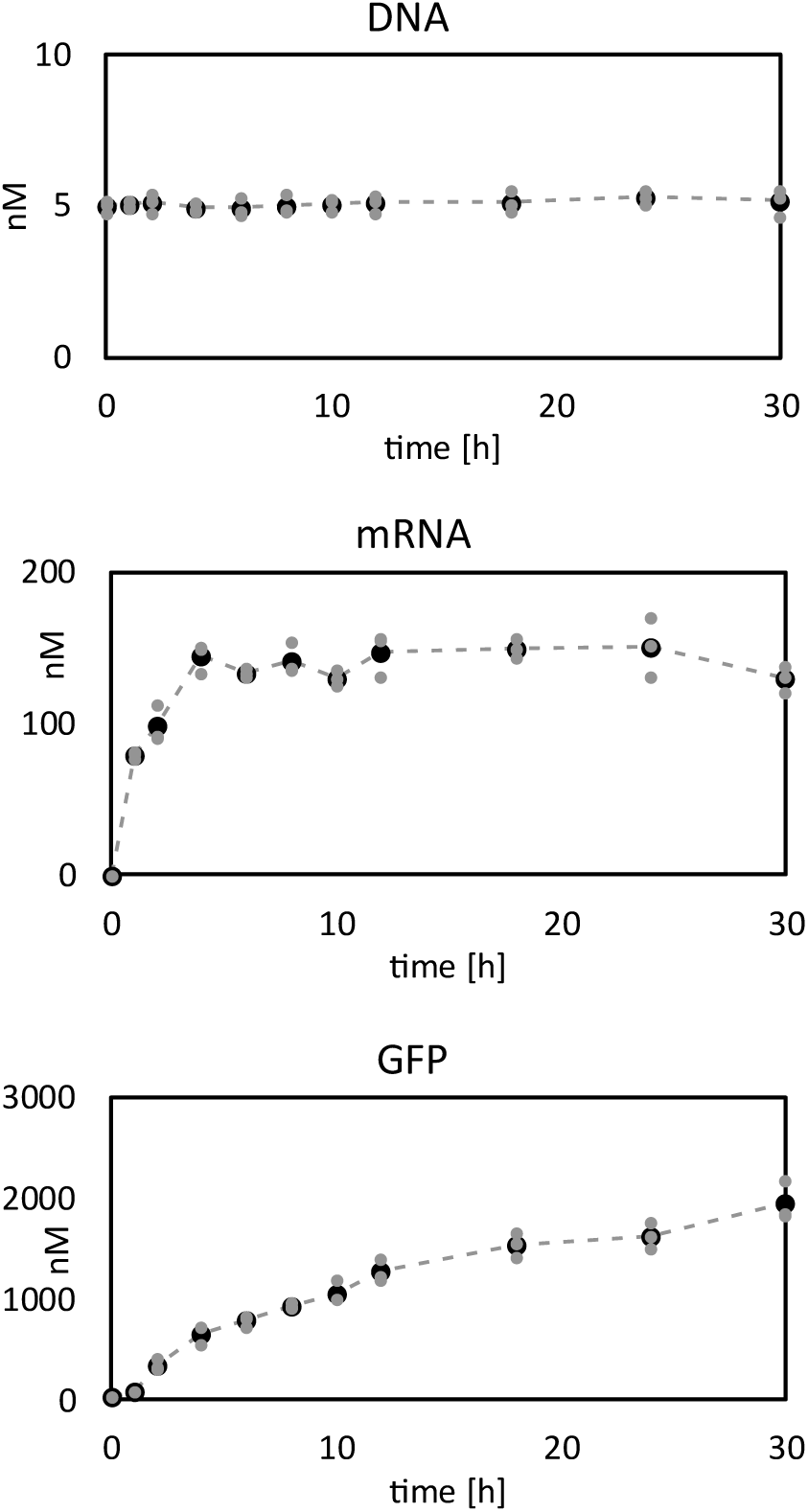
mRNA and protein time course. Synthetic cells were prepared with nM GFP plasmid and nM αHL plasmid. Cells were incubated in the synthetic cell culture media (see Supplementary text section 4) for 30 hours. The GFP mRNA, DNA and GFP fluorescence were measured at indicted time points. The lines connecting the dots are visual guides, not fits. Gray dots show individual data points, black dots are average of three replicates. This data was collected using PURE pREP.

**Figure S44.**
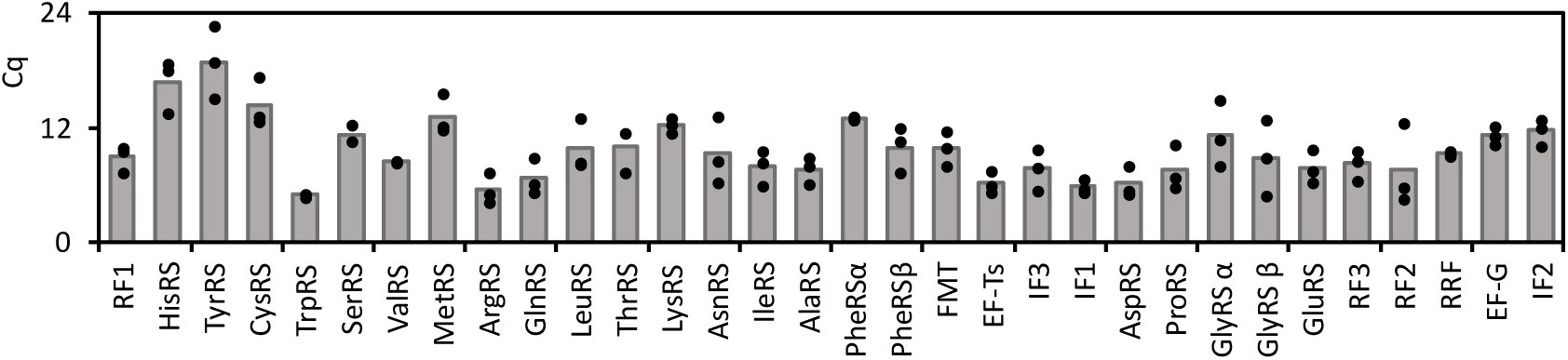
Detection of mRNA for each gene after 5 generations of cell cycle. Quantification of mRNA for all genes in the synthetic cell genome undergoing 5 generations of cell cycle (data for genes not included on the Figure 3). See methods section 15 for Rt-qPCR procedure. This is a qualitative experiment, indicating presence of detectable amount of mRNA for each gene, not the absolute amount of mRNA. Bar graphs indicate average value; dots indicate individual data points from 3 replicates. Values reported on this figure are Cq values from qPCR experiment – **the higher the value, the lower is the abundance of the analyte**. This data was collected using PURE pREP.

**Figure S45.**
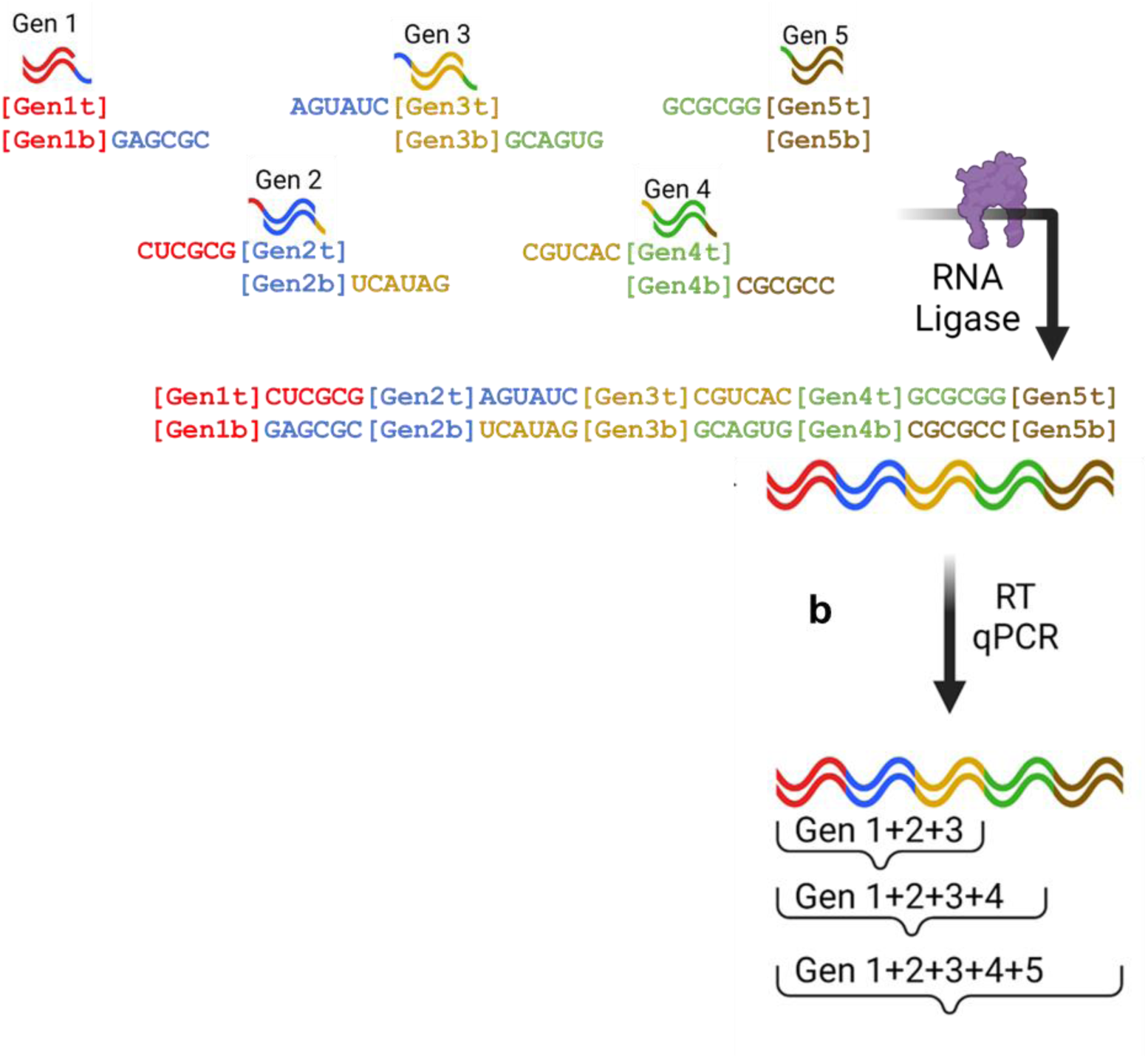
The general scheme of the generation counter. The generation counter consists of five pairs of double stranded RNA with short “sticky” single stranded overhangs. The oligos are labeled: Gen1t for generation 1, top oligo, and Gen1b for generation 1, bottom oligo, etc. Full sequences of RNA oligos used here are in **Table S3**. T4 RNA ligase joins the RNA into one dsRNA product, in the order directed by the complementary overhangs between the generation oligos. The ligation product can then be detected by RT qPCR (see methods section 12). The RT qPCR analysis starts with a pair of primers: the forward primer binding the Gen1 oligo and the reverse primer binding either the Gen3, Gen4, or Gen5 oligo.

**Figure S46.**
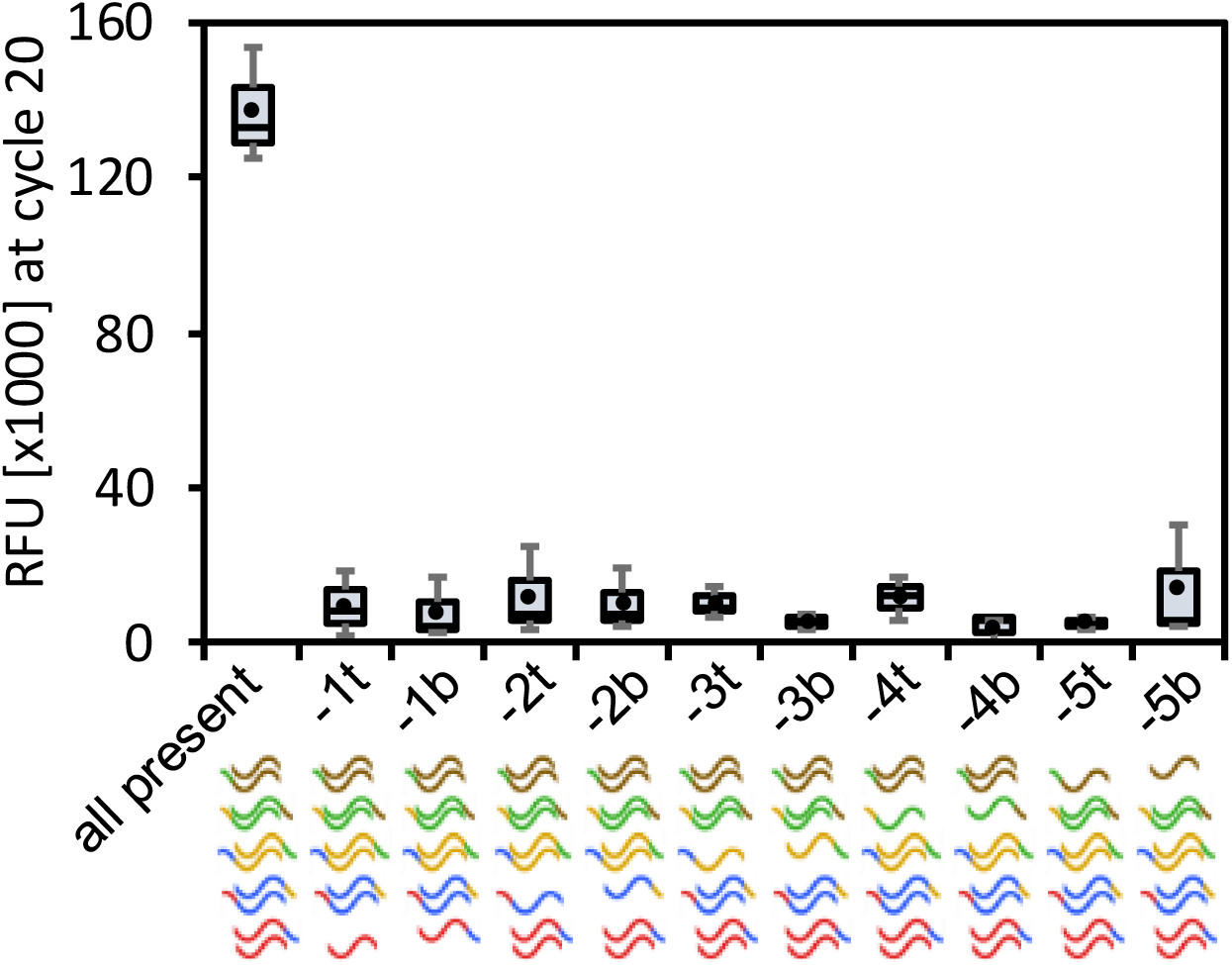
Detection of the generation counter full-length product. The full-length generation counter ligation product can only be produced if all the top and bottom strands of the gen counter oligos are present in the reaction. The graph shows the RT qPCR analysis of the generation counter reaction with all the top and bottom strands present in addition to the reactions missing a specific single strand: top (t) or bottom (b) strand in each generation oligo. The results are shown as fluorescence of the qPCR reaction at cycle 20, because in all the cases of a reaction missing one oligo, there was no Cq call to be made. The graphical legend under the graphs illustrates which generation counter oligo is missing from each experiment, using the color scheme introduced on the generation counter general scheme on **Figure S45**.

**Figure S47.**
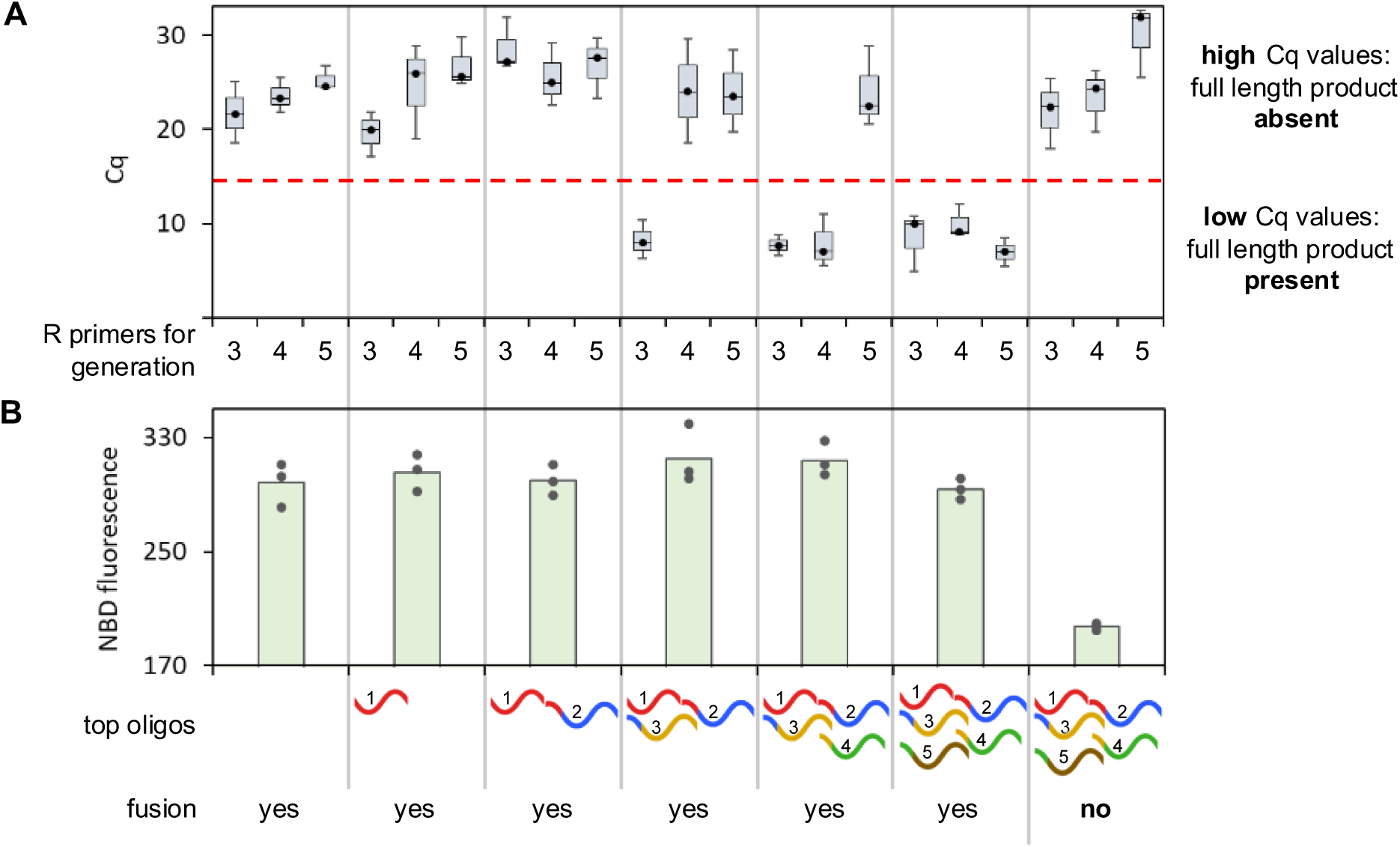
Generation counter tests in fusion experiments. Two populations of liposomes were prepared: one population encapsulated the bottom oligos for all generations (five bottom RNA oligos) and the RNA ligase enzyme. The second liposome population had top strands for one or more generations. Most samples contained a matching pair of DNA fusion tags, resulting in liposome fusion. In one set of experiments, the DNA tags were not complementary, so no fusion occurred. All liposomes in the population with the bottom strands and ligase were labeled with a FRET pair of NBD and rhodamine to monitor membrane fusion. After the fusion experiment and incubation for six hours, the NBD fluorescence was measured to quantify the membrane fusion (panel **B**) and samples were processed for RT qPCR experiments using a forward primer binding sequence at the beginning of the Gen1 oligo and a reverse primer binding in the sequence for Gen3, Gen4, or Gen5 (as indicated on the bottom of the figure legend), with Cq results shown on panel **A**. Experiments optimizing the ratio of liposome populations and ligase enzyme are shown on **Figure S48**. The dotted red line on panel **A** is an arbitrary cut-off at a Cq of 14 between distinct clusters of Cq values: a low Cq value indicates detection of the complete oligo containing sequence which indicates the fusion of the stated number of generation oligos, whereas high Cq values indicates the lack of the desired product. On panel **B**, high NBD signal indicates membrane mixing, low NBD signal indicates lack of membrane mixing (see **Figure S14** for schematic of the FRET experiment). The dots on the top of bars on panel **B** indicate all individual data points with the bars being the average value. Box and whisker data points on panel **A** are prepared with n=3 independently prepared and processed liposome samples. The RT qPCR data on panel **A** are obtained from the same samples as the NBD FRET data on panel **B**.

**Figure S48.**
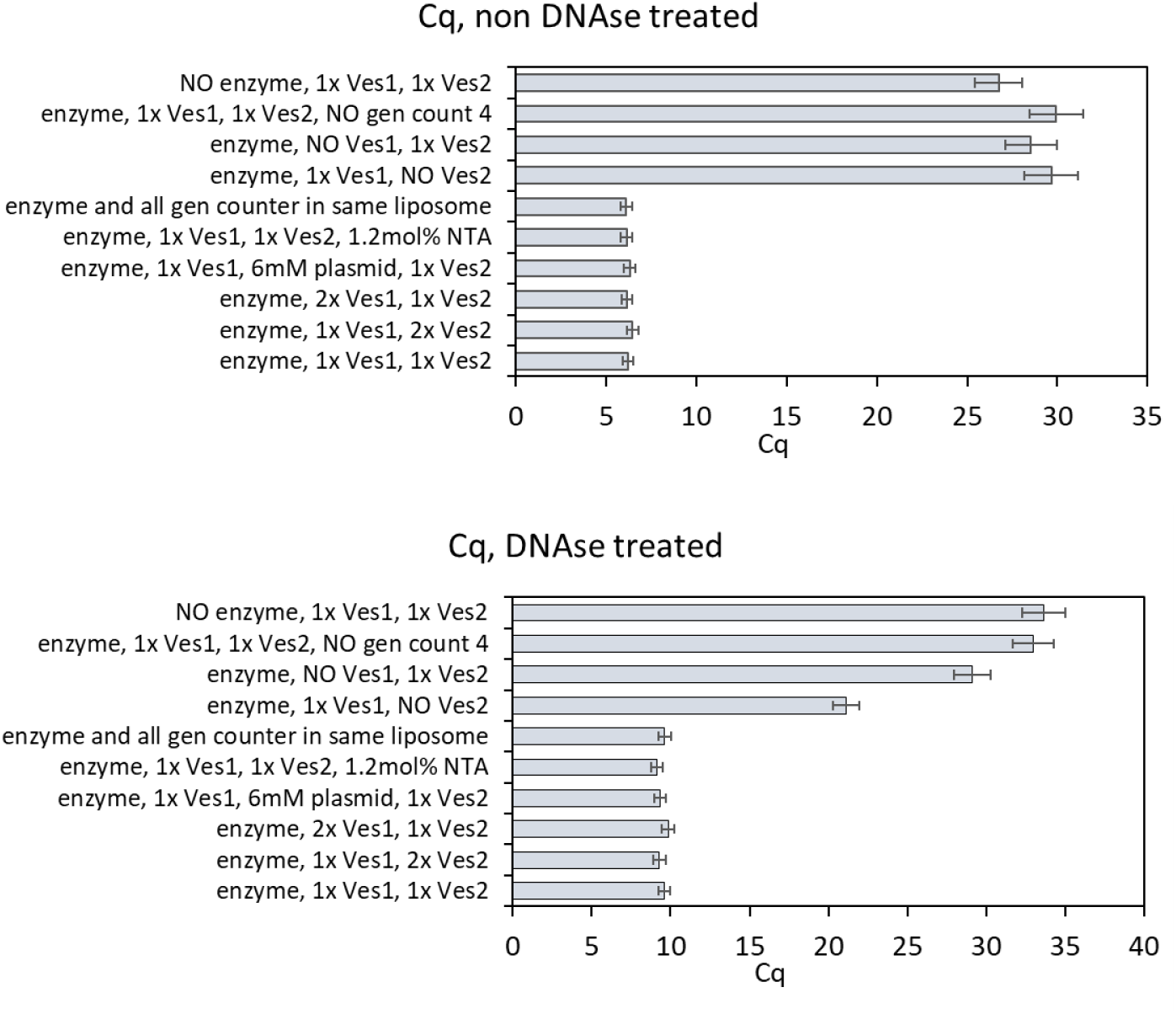
Optimizing the ratio of liposome populations and ligase enzyme. In all samples except one, all gen counter oligos were present. Ves1 is a liposome population with generation counter oligos 1 and and the ligase, nM αHL plasmid. Ves2 is liposome population with generation counter oligos 2, 3 and 4, 0.6mol% NTA lipids in the membrane.

**Figure S49.**
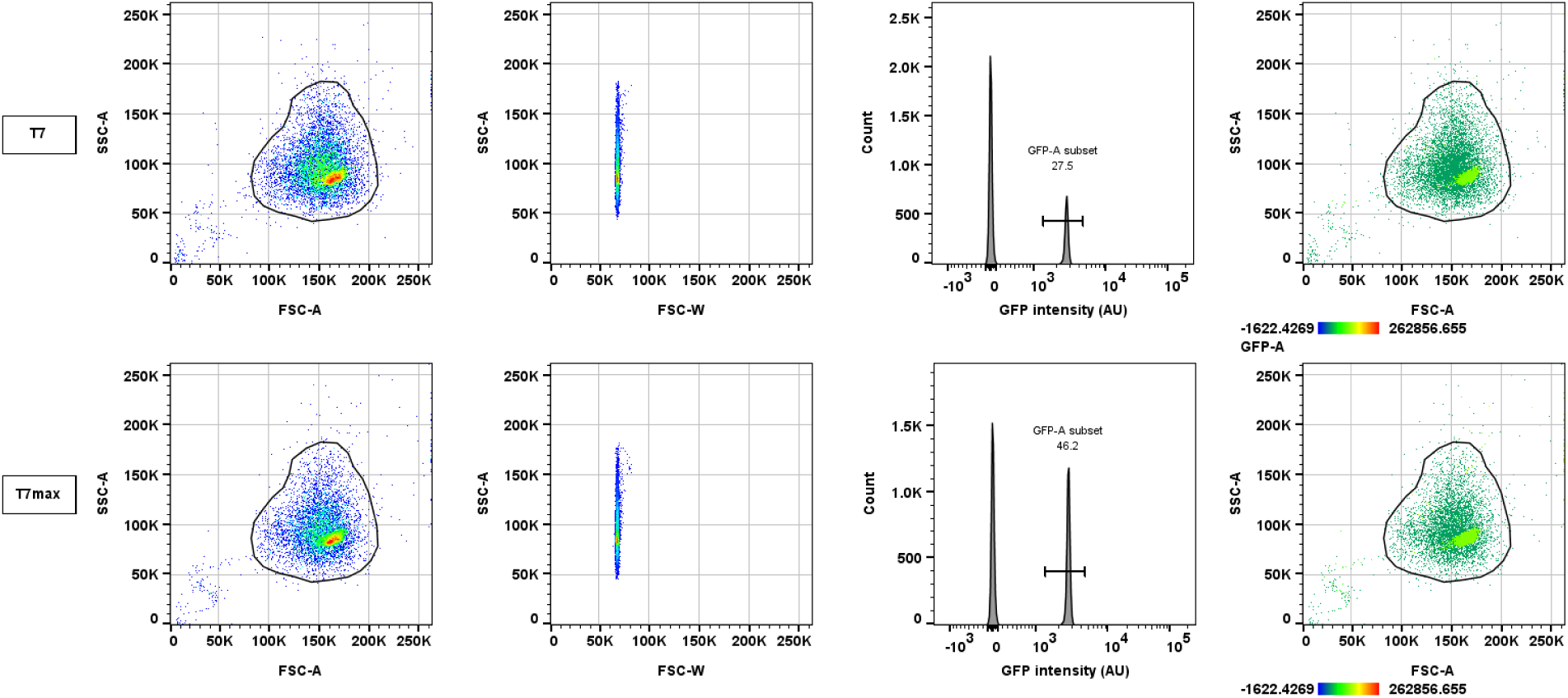
Flow cytometry analysis of fusion reactions, with either T7 αHL or T7Max αHL plasmids. The promoter on αHL plasmid was the only difference between the samples, the concentrations, feeder liposomes and all other parameters were identical. FSC- forward scatter, SSC- side scatter. See Methods section 17 “Flow cytometry”. This data was collected using PURE pREP.

**Figure S50.**
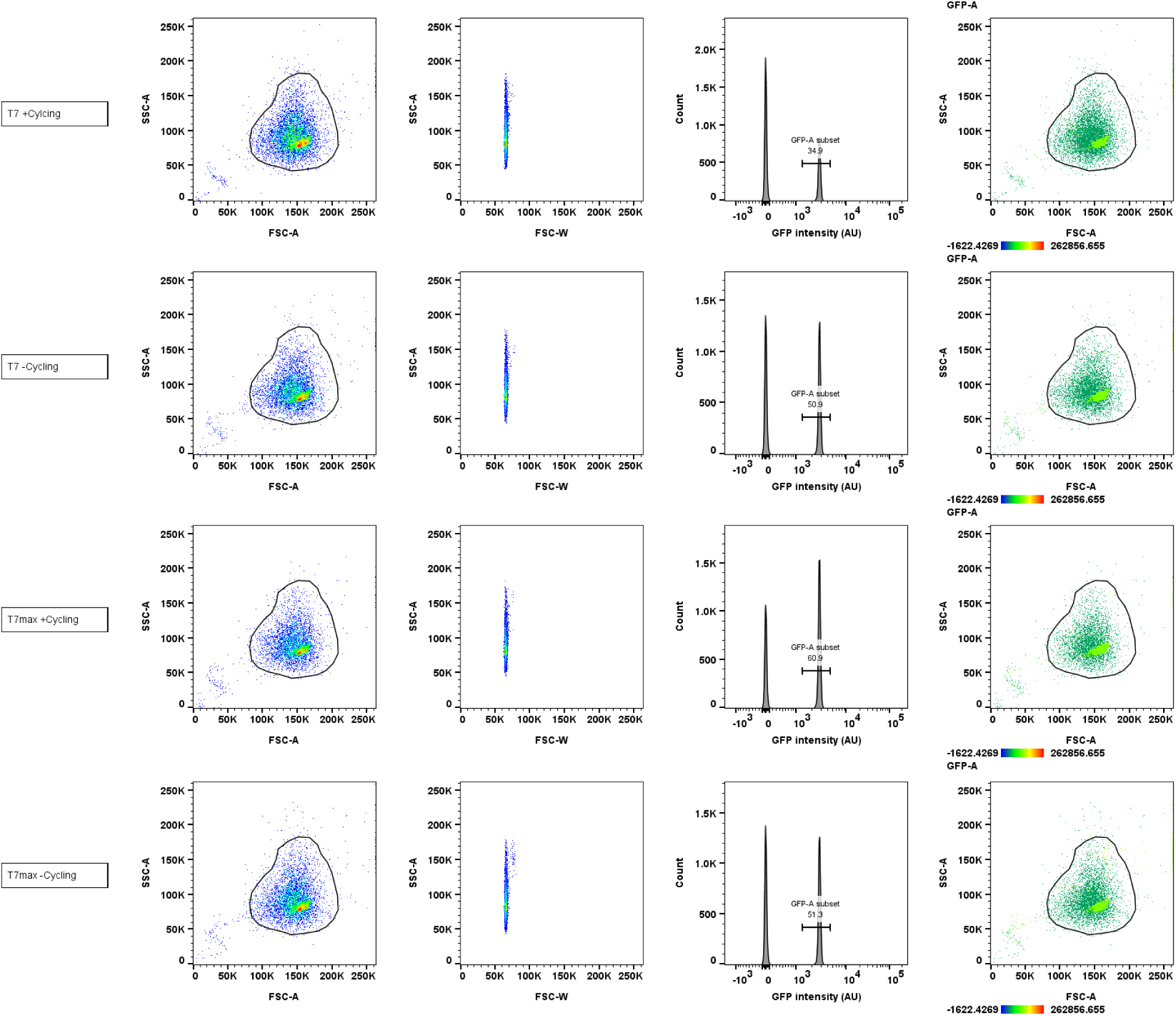
Flow cytometry analysis of synthetic cells after 5 generations of growth and division. In all samples, cells with T7 and T7Max αHL plasmids started at equal concentration (1 to 1 ratio of T7 population to T7Max population). The GFP plasmid was present either in samples with T7 αHL plasmid or in samples with T7Max αHL plasmid. +cycling label indicates samples that went through 5 generations of growth and division. -cycling label indicates samples that were incubated through 5 cycles, but the feeder liposomes had no Ni-NTA lipids, so the cells did not undergo feeding and growth. Negative (no GFP) and positive (all cells contain GFP) controls for this analysis are shown on **Figure S51**. FSC- forward scatter, SSC- side scatter. See Methods section 17 “Flow cytometry”. This data was collected using PURE pREP.

**Figure S51.**
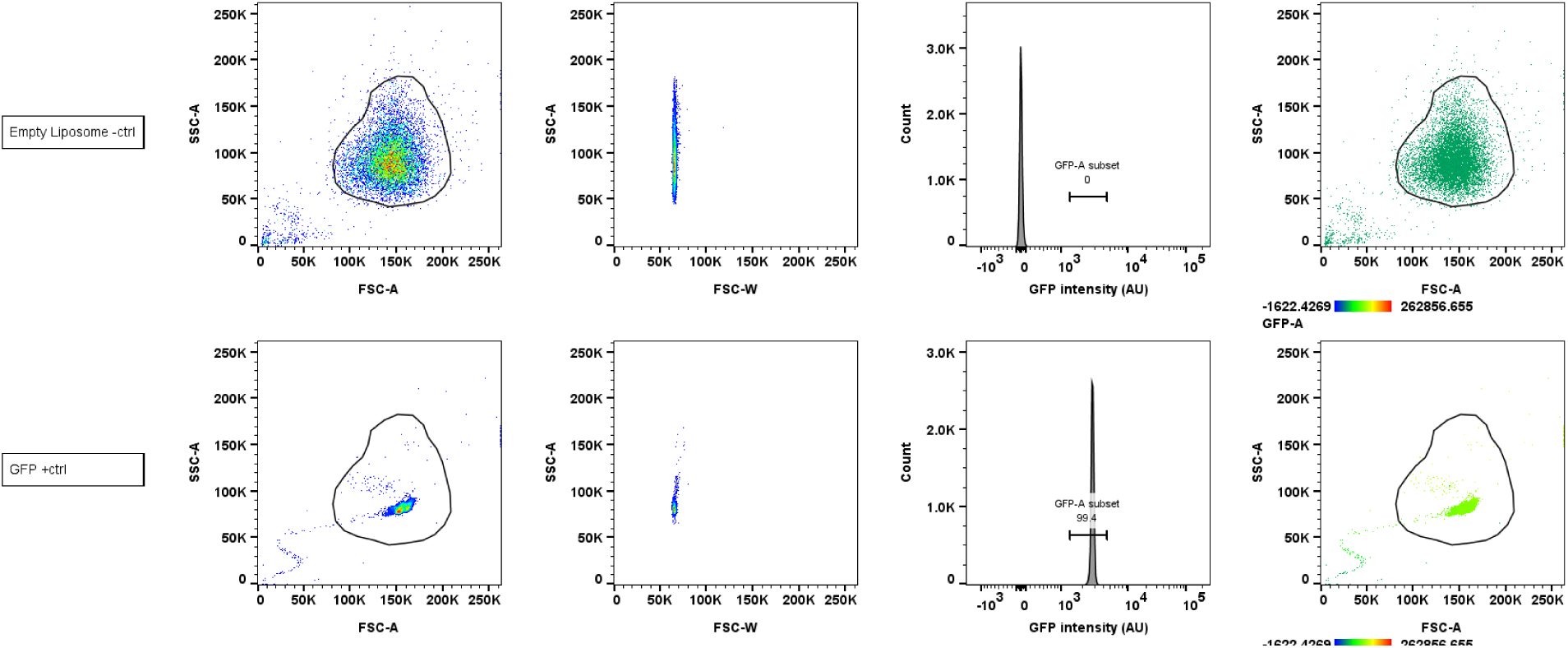
Flow cytometry analysis of positive and negative control samples after 5 generations of growth and division. Cells prepared like in the generation fusion experiments, but did not undergo cycling and generations. The samples either contained GFP plasmid in all cells (labeled GFP +ctrl), or did not contain GFP plasmid at all (labeled empty liposome -ctrl). FSC- forward scatter, SSC- side scatter. See Methods section 17 “Flow cytometry”. This data was collected using PURE pREP.

**Figure S52.**
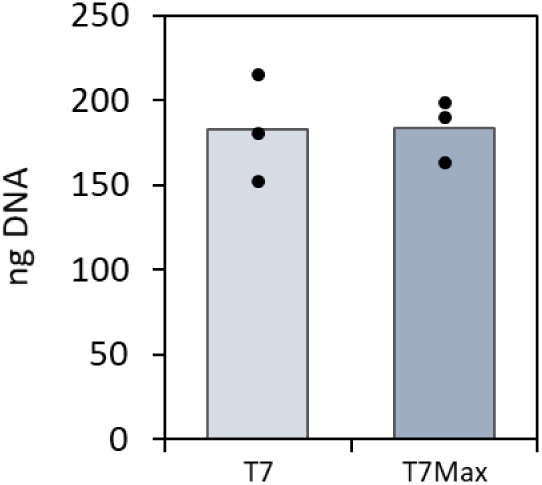
Phi29 replication for T7 αHL and T7Max αHL plasmids. Experiments were prepared according to the manufacturer protocol (NxGen Phi29, Lucigen 30221-2), and the resulting DNA was quantified using Quant-iT PicoGreen dsDNA Assay Kits. Numbers shown on this figure represent total yield isolated from the amplification reaction. Dots on top of the bars indicate all the individual data points, the bars are the average value.

**Figure S53.**
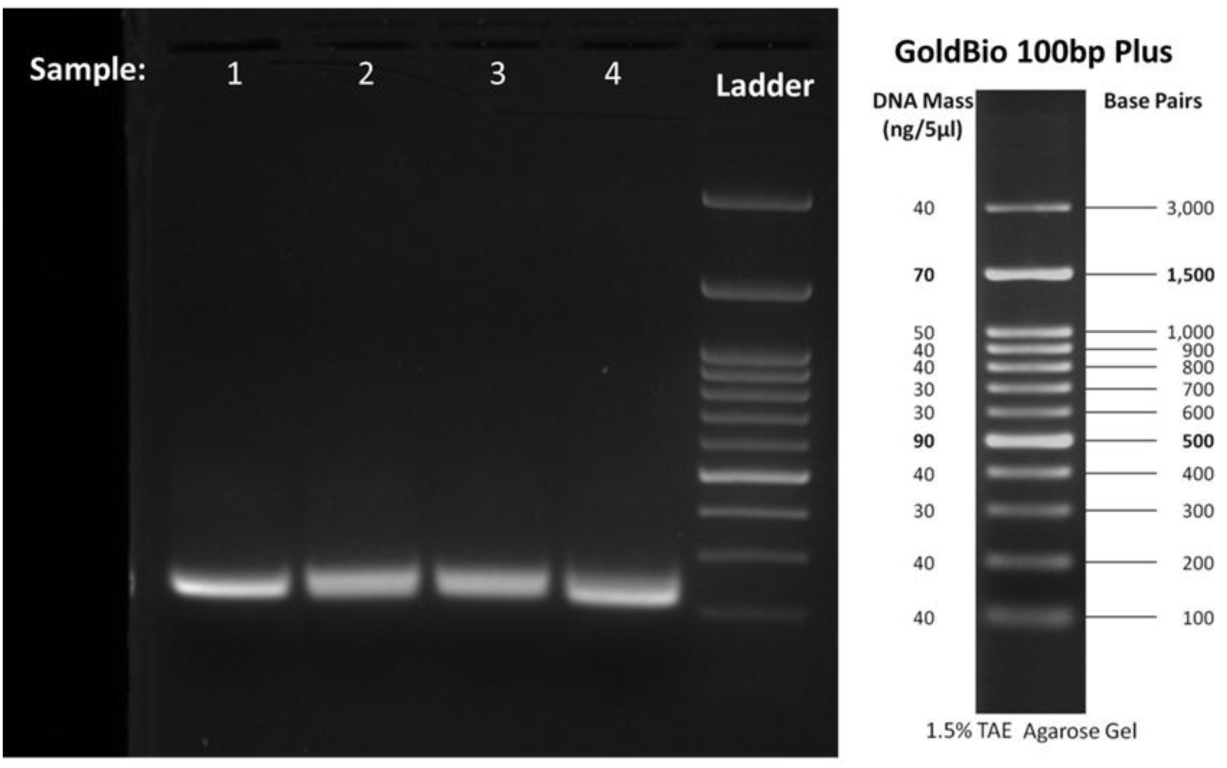
Amplicon DNA isolated from samples of competing cells after 5 generations. The DNA isolated from generation 5 samples was DpnI digested, and PCR amplified with partial Illumina sequencing adapters. The samples contained a mix of T7 and T7Max amplicons that migrated on the gel as single band. <u>Sample 1</u>: cells with T7 and T7Max αHL plasmids started at equal concentration (1 to 1 ratio of T7 population to T7Max population). Samples went through 5 generations of growth and division. <u>Sample 2</u>: cells with T7 αHL plasmid are 90%, and T7Max are 10% of total concentration (9 to 1 ratio of T7 population to T7Max population). Samples went through 5 generations of growth and division. <u>Sample 3</u>: cells with T7 and T7Max αHL plasmids started at equal concentration (1 to 1 ratio of T7 population to T7Max population). Samples were incubated through 5 cycles, but the feeder liposomes had no Ni-NTA lipids, so the cells did not undergo feeding and growth. <u>Sample 4</u>: cells with T7 αHL plasmid are 90%, and T7Max are 10% of total concentration (9 to 1 ratio of T7 population to T7Max population). Samples were incubated through 5 cycles, but the feeder liposomes had no Ni-NTA lipids, so the cells did not undergo feeding and growth. This data was collected using PURE pREP.

**Figure S54.**
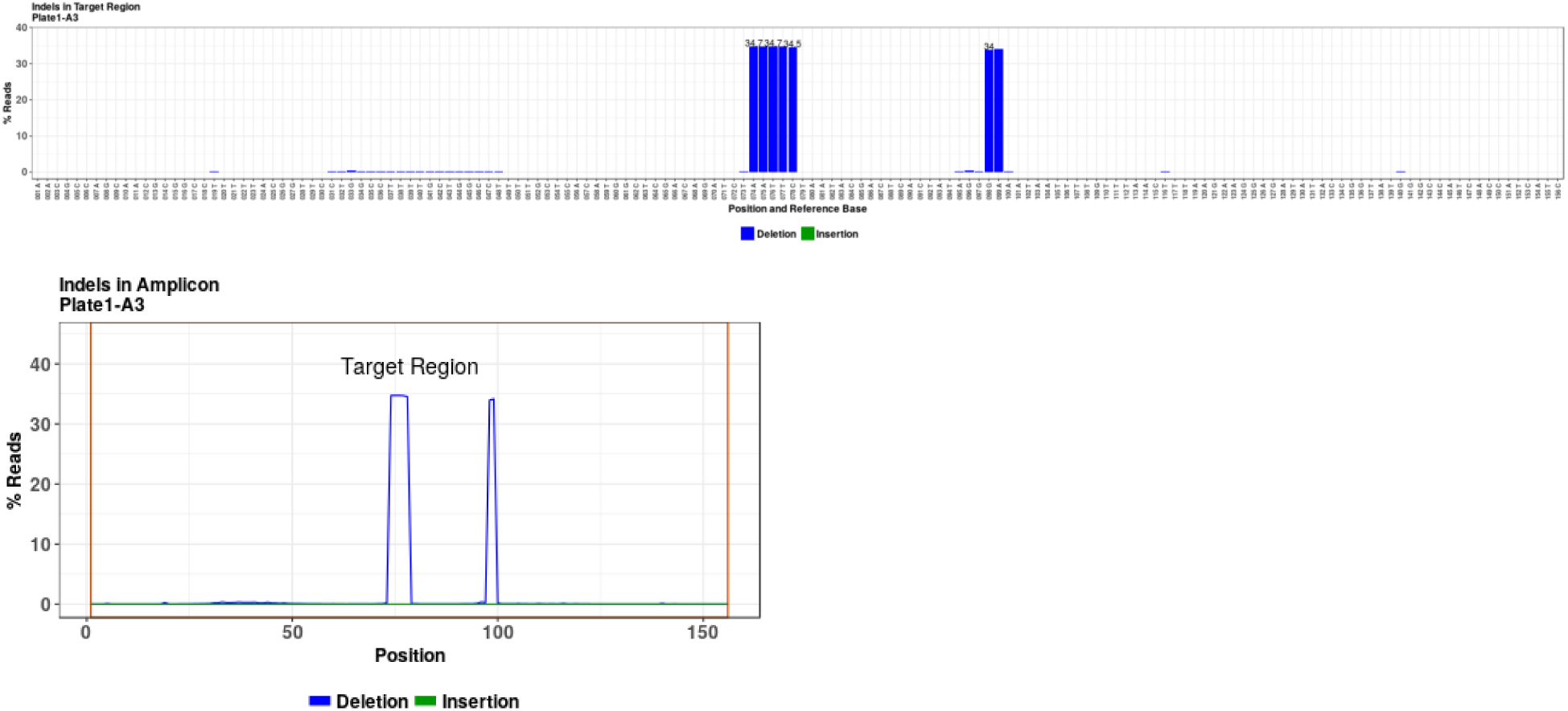
Example of analysis of NGS data. This is data for sample where T7 and T7Max cells started at 1 to 1 ratio and did undergo 5 cycles of growth and division. For analysis, T7Max was used as reference sequence, and T7 shows up as deflection mutant. This data was collected using PURE pREP.

**Figure S55.**
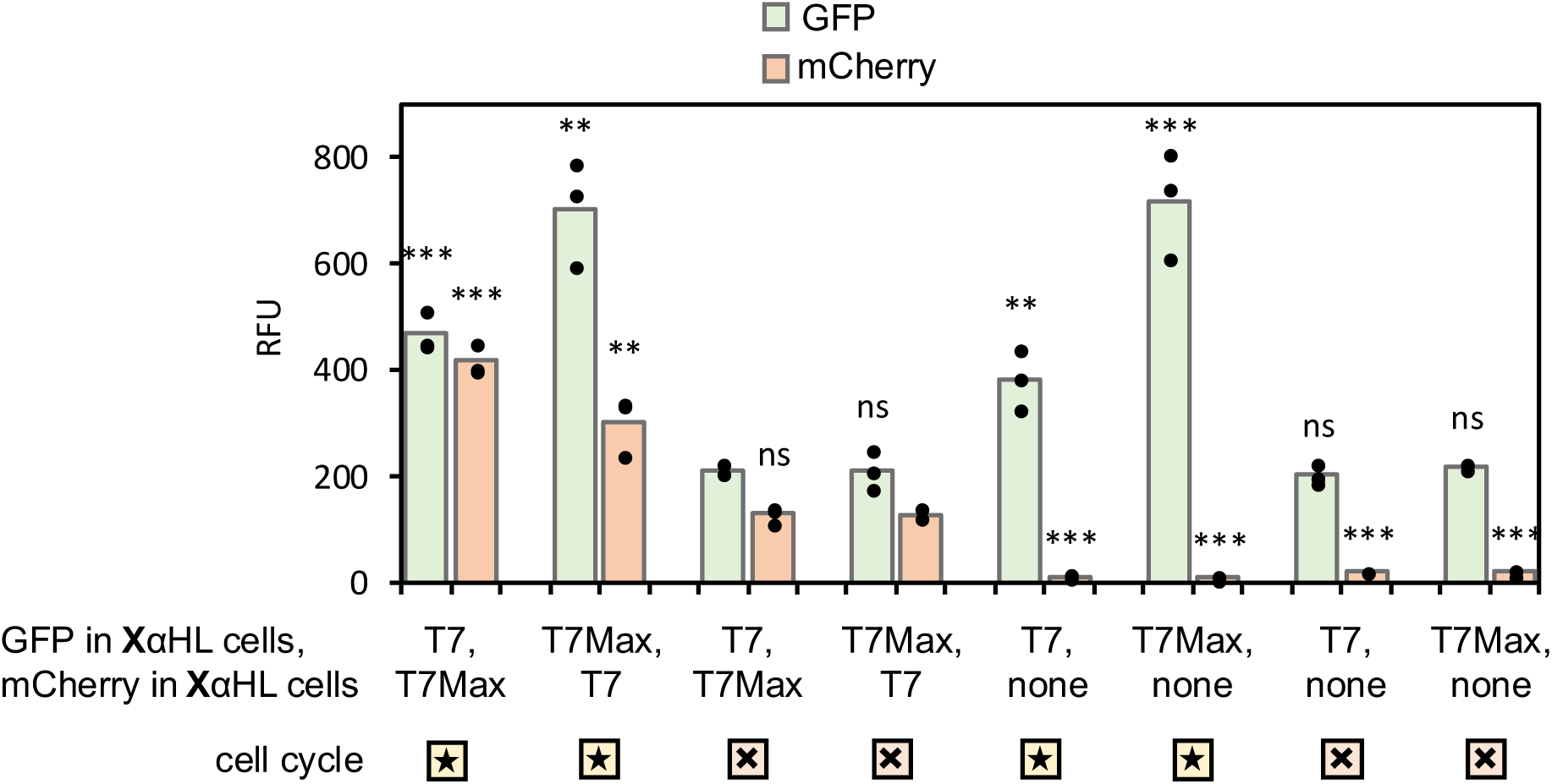
Selection experiments with normalized metabolic load. Raw fluorescence data selection experiments with all cells carrying a fluorescent protein of similar size (normalized metabolic load). Experiments performed similarly to those described on panels I-M of Figure 4, but both T7 and T7Max cells carried fluorescent protein (GFP or mCherry, as indicated in the legend under graph on panel o). After 5 generations, fluorescence of both GFP and mCherry was measured (see **Table S5** for wavelengths). Bar graph shows average value; individual replicates are represented by black dot data points. See **Table S10** for details of statistical analysis results. P values calculated for GFP and for mCherry compared to the negative control: incubation without DNA replication and without feeding and division (symbol 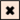) in the weaker promoter T7 cells. This data was collected using PURE pREP.

**Figure S56.**
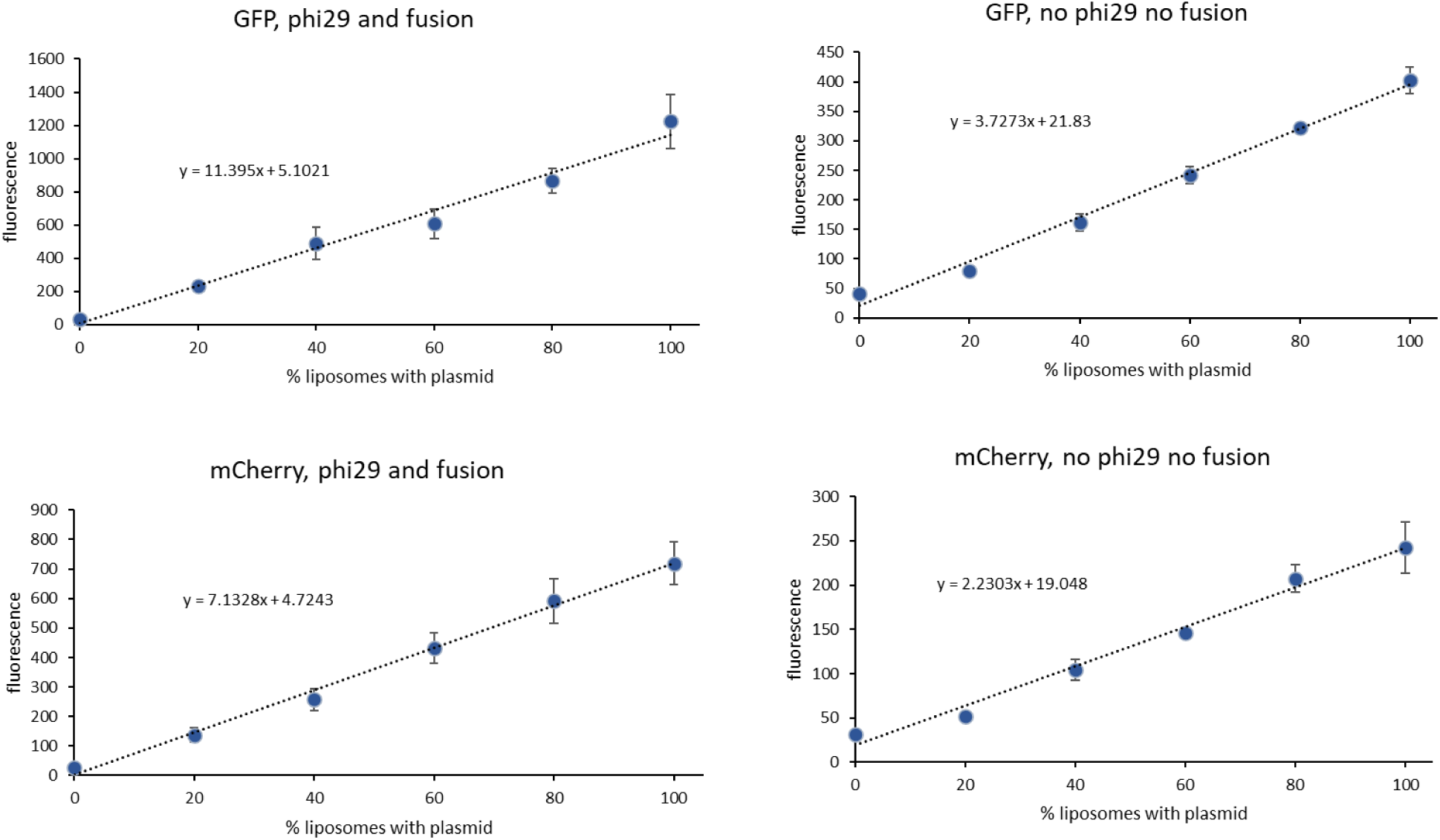
Calibration curves for GFP and mCherry production after 5 generations of synthetic cell cycle. All generation experiments were started with two populations of cells, carrying either GFP or mCherry plasmid at 5nM. After 5 generations of cell cycle fluorescence of both GFP and mCherry was measured. Separate control experiments were run without the Phi29 and fusion. The resulting calibration curves show the relationship between measured green (GFP) and red (mCherry) fluorescence to relative percent of the synthetic cell population carrying that fluorescent protein gene. All experiments were done in triplicate, error bars indicate SEM, n=3 independently prepared and processed liposome samples. This data was collected using PURE pREP.

**Figure S57.**
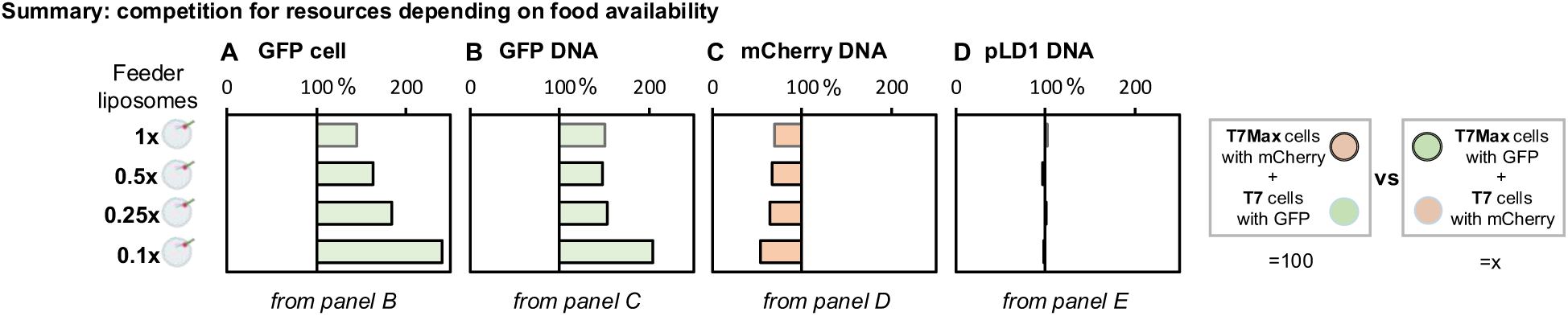
Summary of the results of competitive growth. Summary of measurements from panels B-E of Figure 5. Graphs show difference between T7Max αHL (fast growing) and T7 αHL (slower growing) populations. The value on X axis on all graphs represents what percent of the value measured in the sample containing T7Max αHL mCherry cells with T7 αHL GFP cells (labeled 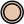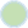) is the value measured in sample containing T7 αHL mCherry cells and T7Max αHL GFP cells (labeled 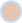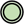) T7Max αHL mCherry cells with T7 αHL GFP cells (labeled 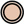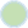) For example, a bar graph with GFP fluorescence value of 184 means GFP fluorescence in sample containing T7 αHL mCherry cells and T7Max αHL GFP cells (labeled 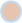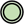) is 184% of the GFP value measured in sample of T7Max αHL mCherry cells with T7 αHL GFP cells (labeled 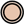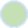). **A**: GFP cytometry, **B**: GFP DNA, **C**: mCherry DNA, and **D**: pLD1 “housekeeping gene” DNA. This data was collected using PURE pREP.

**Figure S58.**
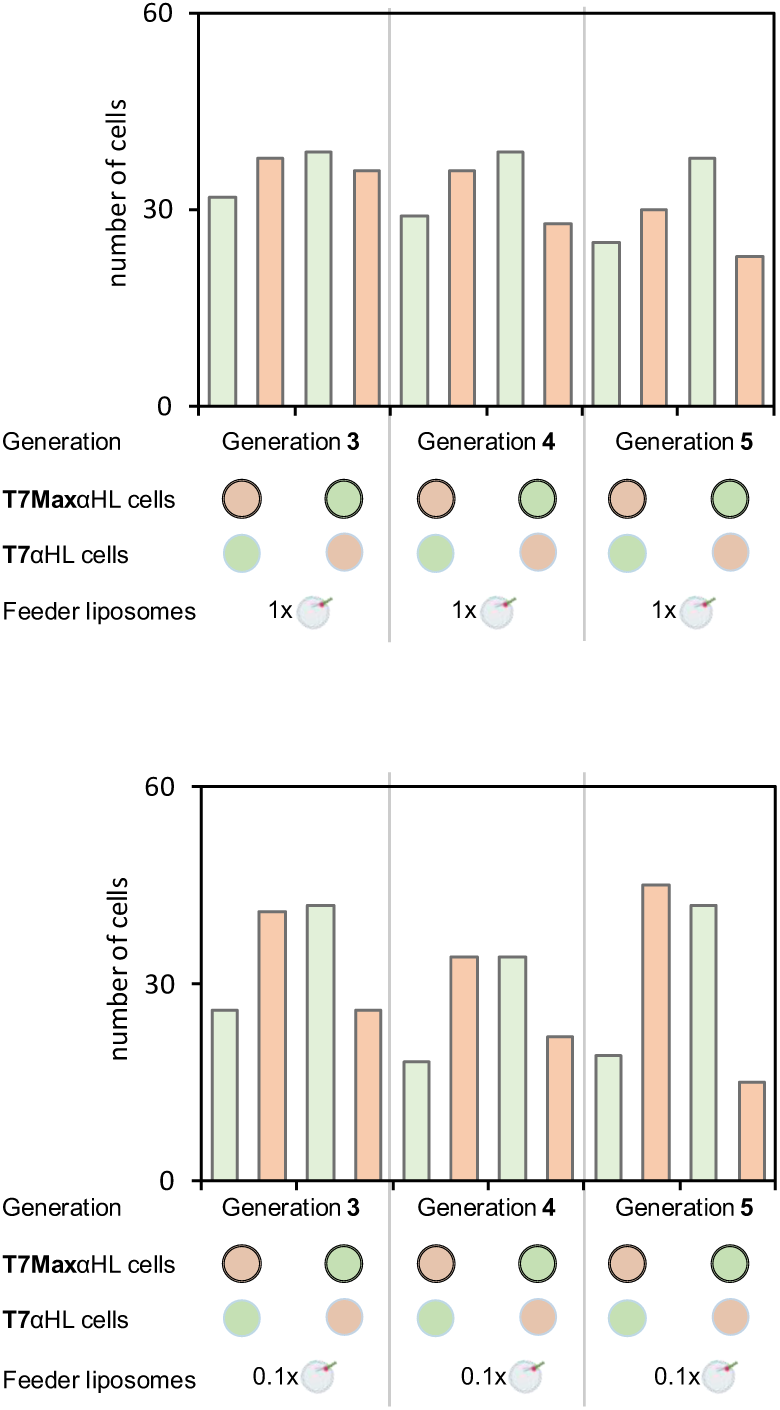
Number of cells containing GFP or mCherry plasmid during the subsequent generations of competition experiments. Single cell analysis of abundance of GFP plasmid and mCherry plasmid in synthetic cells undergoing rounds of competitive feeding. Two populations of synthetic cells were mixed in each competition experiment. One population expressed green marker (GFP) and the other population expressed red marker (mCherry). Each experiment was performed in two variants: either mCherry was in the T7Max αHL population (symbol 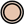) and GFP was in the T7 αHL population (symbol 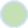), or the reverse with the GFP in the T7Max αHL population (symbol 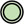) and mCherry in the T7 αHL population (symbol 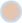). Both variants of each experiment had five generations of growth and feeding, with varying amount of feeder liposomes. After generation 3, 4 and 5 of the experiments with 1x feeder liposomes and with 0.1x feeder liposomes, samples were analyzed using single cell protocol (see Methods section 23). Cells were isolated by dilution. As described in Methods section 23, the only cells used for the analysis were the ones with detectable 16S ribosomal RNA – meaning samples that contained a cell, not empty samples that did not get a cell due to high dilution. The 1x feeder liposomes means the same conditions as cell cycle experiments on Figure 3. The 0.1x feeder liposomes refer to proportionally less feeder liposomes than in the original 1x conditions. Individual data points showing abundance of plasmid in each analyzed cell are on **Figure S59** for 1x feeding conditions and **Figure S60** for 0.1x feeding conditions. This data was collected using PURE pREP.

**Figure S59.**
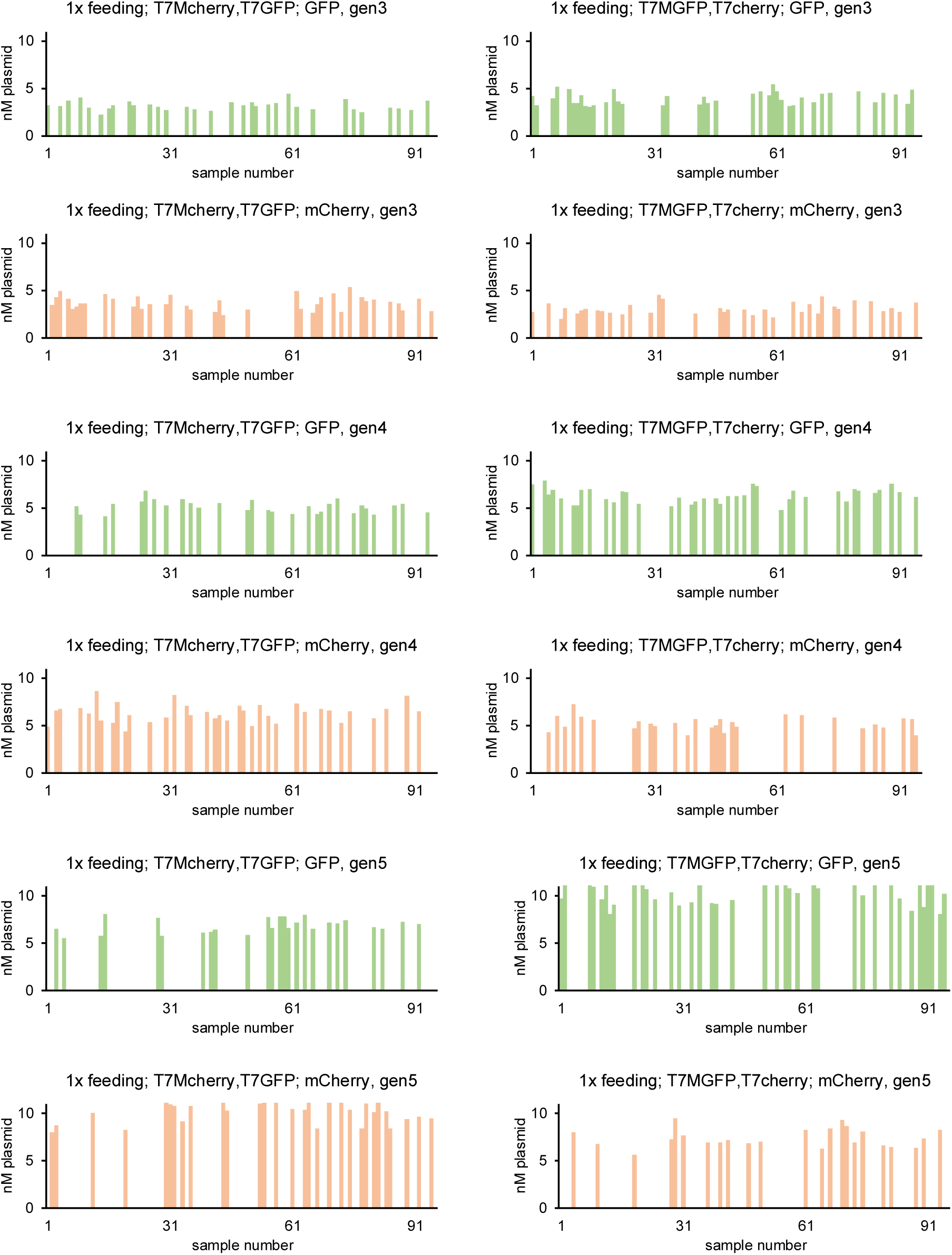
Abundance of GFP and mCherry plasmid in each individual cell after competitive growth experiments, with 1x feeder liposomes. Individual data points showing abundance of either mCherry or GFP plasmid in each of the analyzed cells, for competitive growth experiments as described on Figure 5. This figure shows individual data points for summary **Figure S58**. Sample labels indicate: 1x feeding – 1x feeding liposomes, the same feeding concentration conditions as cell cycle experiments on Figure 3. T7Mcherry – the T7Max cells contained mCherry T7MGFP – the T7Max cells contained GFP T7cherry – the T7 cells contained mCherry T7GFP – the T7 cells contained GFP GFP, green bars, indicate measurements of GFP plasmid mCherry, red bars, indicate measurements of mCherry plasmid gen3, gen4 and gen5 – indicate generation after which the samples were collected. The Y axis on all graphs on **Figure S59** and **Figure S60** is set to the same scale, to enable direct visual comparison of plasmid abundance. This data was collected using PURE pREP.

**Figure S60.**
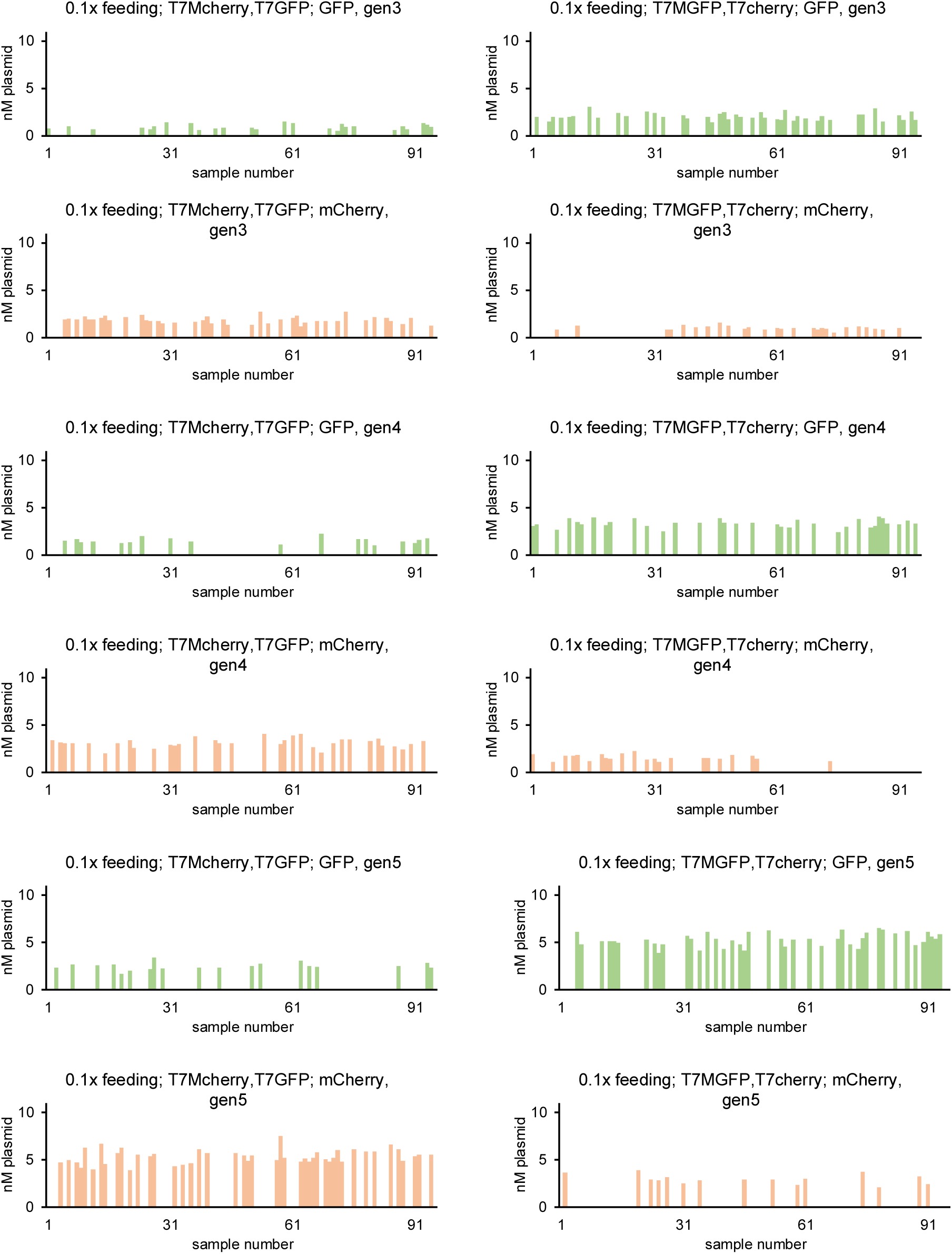
Abundance of GFP and mCherry plasmid in each individual cell after competitive growth experiments, with 0.1x feeder liposomes. Individual data points showing abundance of either mCherry or GFP plasmid in each of the analyzed cells, for competitive growth experiments as described on Figure 5. This figure shows individual data points for summary **Figure S58**. Sample labels indicate: 0.1x feeding – 0.1x feeding liposomes, the concentration of feeding liposomes that is 10% of the concentration used in conditions described as cell cycle experiments on Figure 3. T7Mcherry – the T7Max cells contained mCherry T7MGFP – the T7Max cells contained GFP T7cherry – the T7 cells contained mCherry T7GFP – the T7 cells contained GFP GFP, green bars, indicate measurements of GFP plasmid mCherry, red bars, indicate measurements of mCherry plasmid gen3, gen4 and gen5 – indicate generation after which the samples were collected. The Y axis on all graphs on **Figure S59** and **Figure S60** is set to the same scale, to enable direct visual comparison of plasmid abundance. This data was collected using PURE pREP.

**Figure S61.**
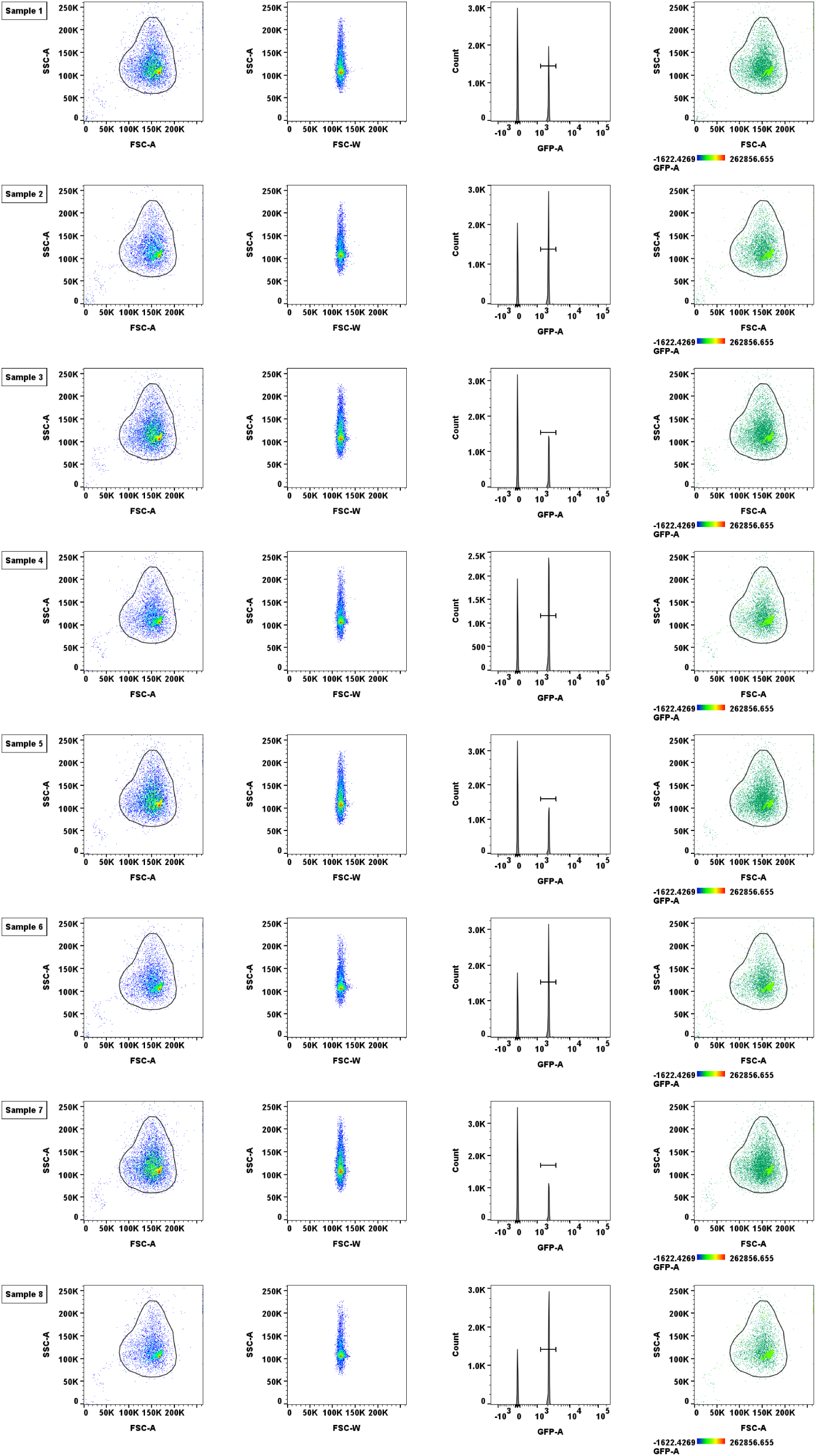
Flow cytometry data from feeding competition experiments. <u>sample 1</u>, T7 GFP, T7Max mCherry, 1x feeding <u>sample 2</u>, T7Max GFP, T7 mCherry, 1x feeding <u>sample 3</u>, T7 GFP, T7Max mCherry, 0.5x feeding <u>sample 4</u>, T7Max GFP, T7 mCherry, 0.5x feeding <u>sample 5</u>, T7 GFP, T7Max mCherry, 0.25x feeding <u>sample 6</u>, T7Max GFP, T7 mCherry, 0.25x feeding <u>sample 7</u>, T7 GFP, T7Max mCherry, 0.1x feeding <u>sample 8</u>, T7Max GFP, T7 mCherry, 0.1x feeding FSC- forward scatter, SSC- side scatter. See Methods section 17 “Flow cytometry”. This data was collected using PURE pREP.

**Figure S62.**
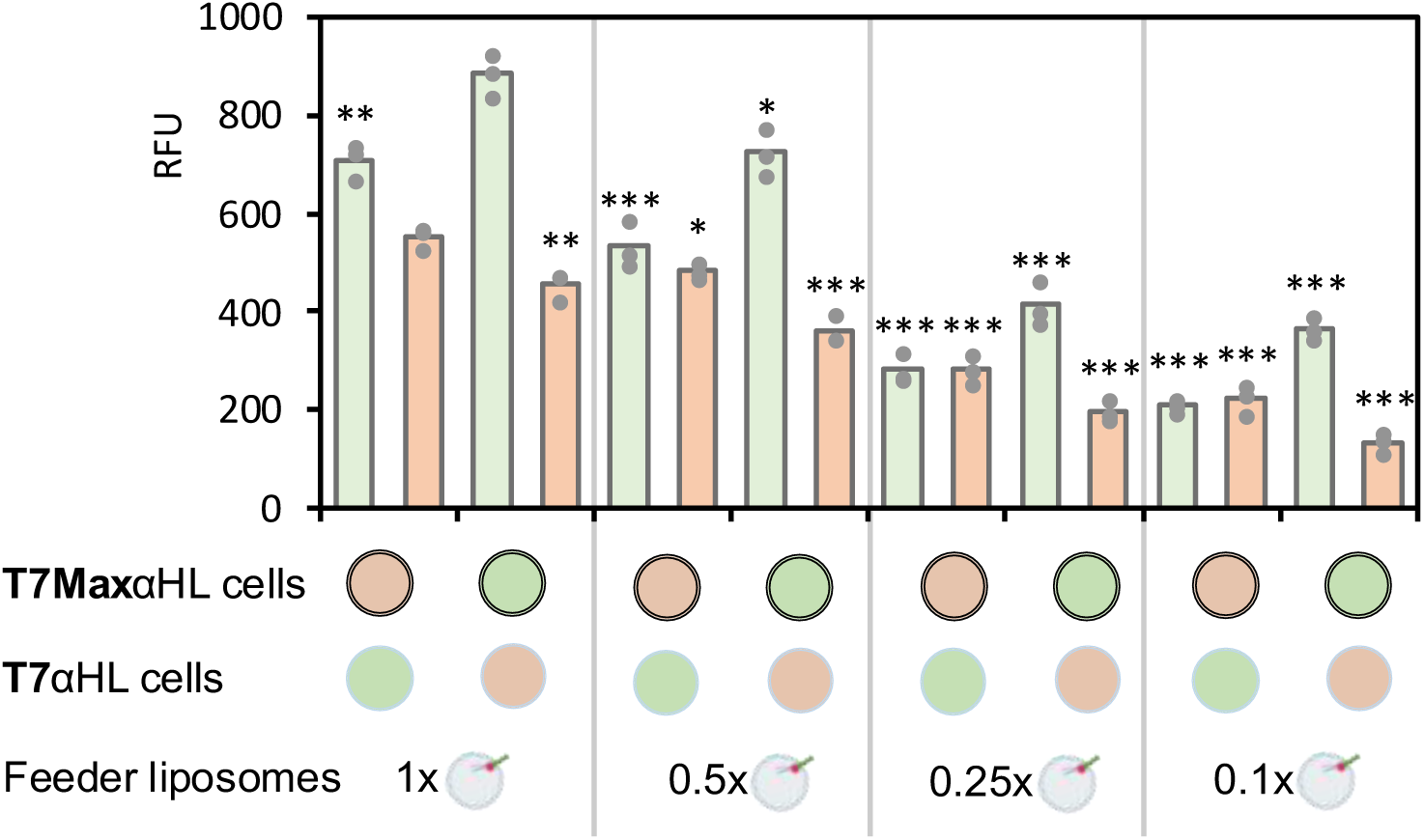
Fluorescence values for mCherry and GFP competitive feeding experiments. Absolute fluorescence values for mCherry (red bars) and GFP (green bars) in competitive feeding experiments. Y axis is RFU (relative fluorescent units). Dots on top of the bar graphs indicate individual values of three replicates. Statistical analysis shows P values for each experiment compared to 1x feeder liposome conditions. ns = P > 0.0; * = P ≤ 0.0, ** = P ≤ 0.01, *** = P ≤ 0.001. This data was collected using PURE pREP.

**Figure S63.**
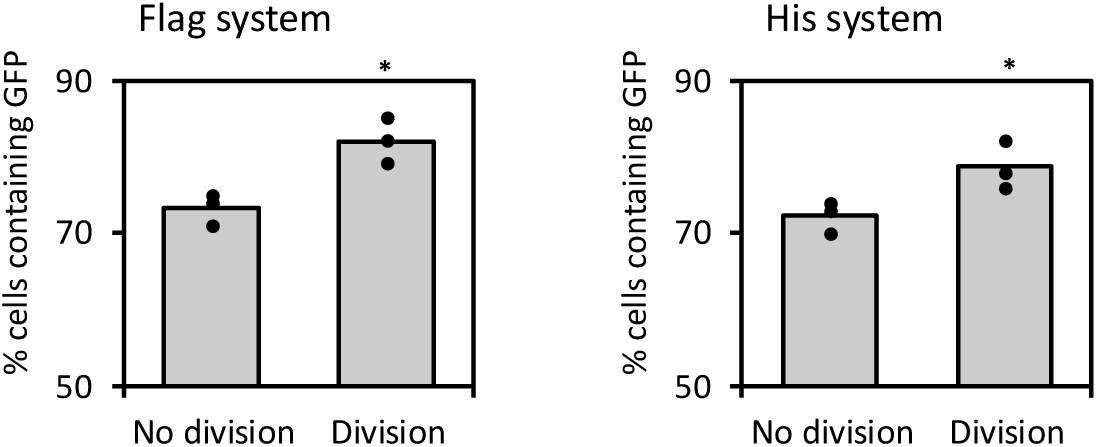
Flow cytometry of synthetic cells before and after division. Genetically encoded division experiments were performed, with synthetic cells expressing tagged membrane protein αHL; streptavidin and linker are added on the outside of the cells, binding to the αHL tag on the surface of the cells. The streptavidin induces curvature in the membrane, leading to cell division. Two systems were tested: αHL His tagged with Ni-NTA linker to bind streptavidin, and αHL FLAG tagged with biotin conjugated FLAG antibody to bind streptavidin. This experiment was performed by preparing synthetic cells with αHL and GFP plasmid (the dividing cells) and mixing them 70: 0 with “dark” cells, liposomes with PURE system but without any plasmid. This was done because for cytometry experiments, cells need to be diluted to certain lipid density, so measuring ratio of “dark” to “GFP” cells was more reliable way to calculate how many cells divided. The starting ratio was 70% cell had GFP. After division, presumably more cells had GFP (higher % of cells with GFP), which is the information that is not going to be lost during dilutions because both GFP and “dark” cells will be diluted equally. If we only used GFP cells, it would be difficult to estimate how many new GFP cells were made, because during dilution all samples would be brought to similar lipid density. Two conditions were tested for both Flag and His system. The synthetic cells in each experiment were: Flag system <u>No division</u>: αHL FLAG present, streptavidin absent, biotin FLAG antibody linker absent. <u>Division</u>: αHL FLAG present, streptavidin present, biotin FLAG antibody linker present. His system <u>No division</u>: αHL His tagged present, streptavidin absent, Ni-NTA linker absent. <u>Division</u>: αHL His tagged present, streptavidin present, Ni-NTA linker present. Dots on top of the bars indicate all the individual data points, the bars are the average value. This data was collected using PURE pREP.

**Figure S64.**
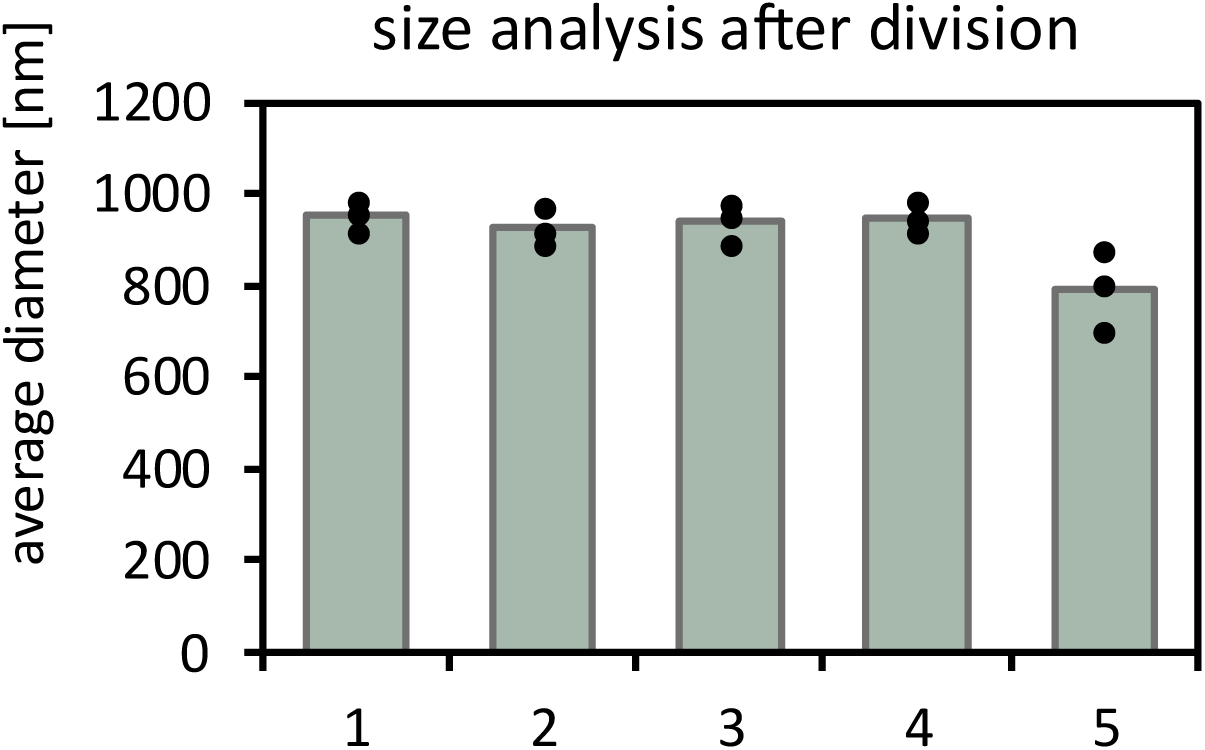
Size of synthetic cells after genetically encoded division using His tag system. Individual data points for each independent replicate experiment for size analysis after division (Figure 6B). Each of the three individual independent replicates (the black dot if an individual data point) is an average of six DLS measurements. <u>Sample 1</u>: αHL His tagged present, streptavidin absent, Ni-NTA linker absent; no division is expected. <u>Sample 2</u>: αHL His tagged present, streptavidin absent, Ni-NTA linker present; no division is expected. <u>Sample 3</u>: αHL His tagged present, streptavidin present, Ni-NTA linker absent; no division is expected. <u>Sample 4</u>: αHL His tagged absent, streptavidin present, Ni-NTA linker present; no division is expected. <u>Sample 5</u>: αHL His tagged present, streptavidin present, Ni-NTA linker present; division is expected. Dots on top of the bars indicate all the individual data points, the bars are the average value. This data was collected using PURE pREP.

**Figure S65.**
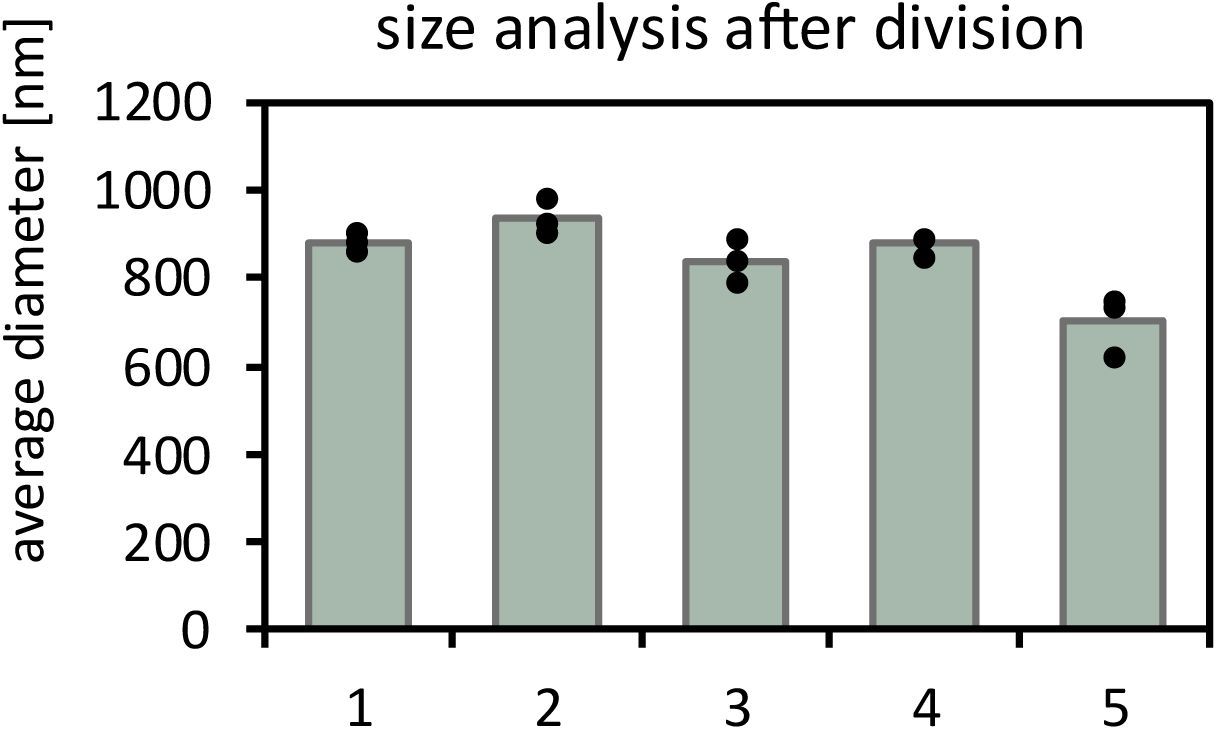
Size of synthetic cells after genetically encoded division using FLAG tag system. Individual data points for each independent replicate experiment for size analysis after division (Figure 6C). Each of the three individual independent replicates (the black dot if an individual data point) is an average of six DLS measurements. <u>Sample 1</u>: αHL FLAG present, streptavidin absent, biotin FLAG antibody linker absent; no division is expected. <u>Sample 2</u>: αHL FLAG present, streptavidin absent, biotin FLAG antibody linker present; no division is expected. <u>Sample 3</u>: αHL FLAG present, streptavidin present, biotin FLAG antibody linker absent; no division is expected. <u>Sample 4</u>: αHL FLAG absent, streptavidin present, biotin FLAG antibody linker present; no division is expected. <u>Sample 5</u>: αHL FLAG present, streptavidin present, biotin FLAG antibody linker present; division is expected. Dots on top of the bars indicate all the individual data points, the bars are the average value. This data was collected using PURE pREP.

**Figure S66.**
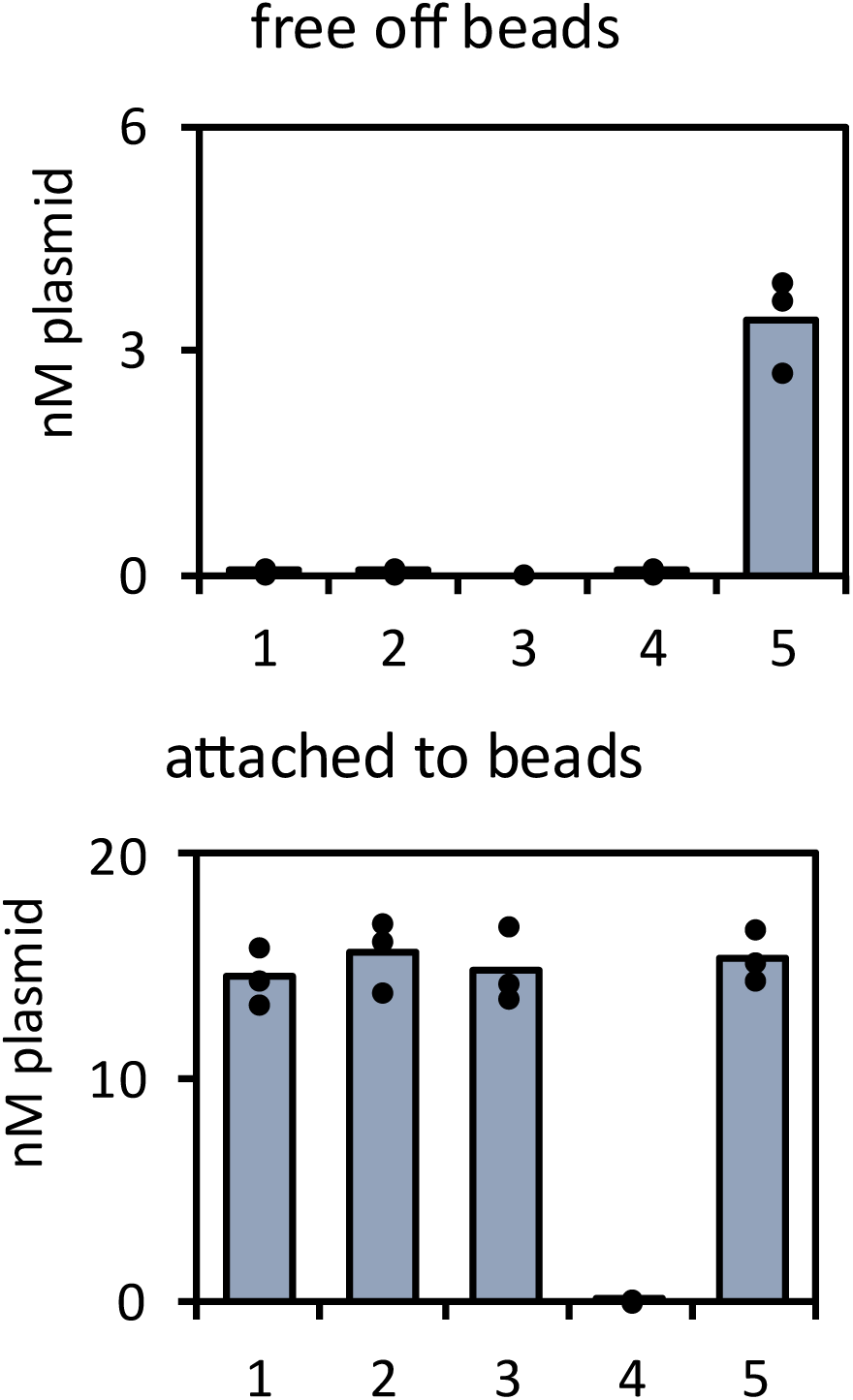
DNA abundance in synthetic cells after genetically encoded division on beads with the His tagged αHL. Individual data points for each independent replicate experiment of measurement of amount of αHL DNA in T7Max synthetic cells after division on beads (Figure 6). <u>Sample 1</u>: αHL present, streptavidin absent, Ni-NTA linker absent; no division is expected. <u>Sample 2</u>: αHL present, streptavidin absent, Ni-NTA linker present; no division is expected. <u>Sample 3</u>: αHL present, streptavidin present, Ni-NTA linker absent; no division is expected. <u>Sample 4</u>: αHL absent, streptavidin present, Ni-NTA linker present; no division is expected. <u>Sample 5</u>: αHL present, streptavidin present, Ni-NTA linker present; division is expected. Dots on top of the bars indicate all the individual data points, the bars are the average value. This data was collected using PURE pREP.

**Figure S67.**
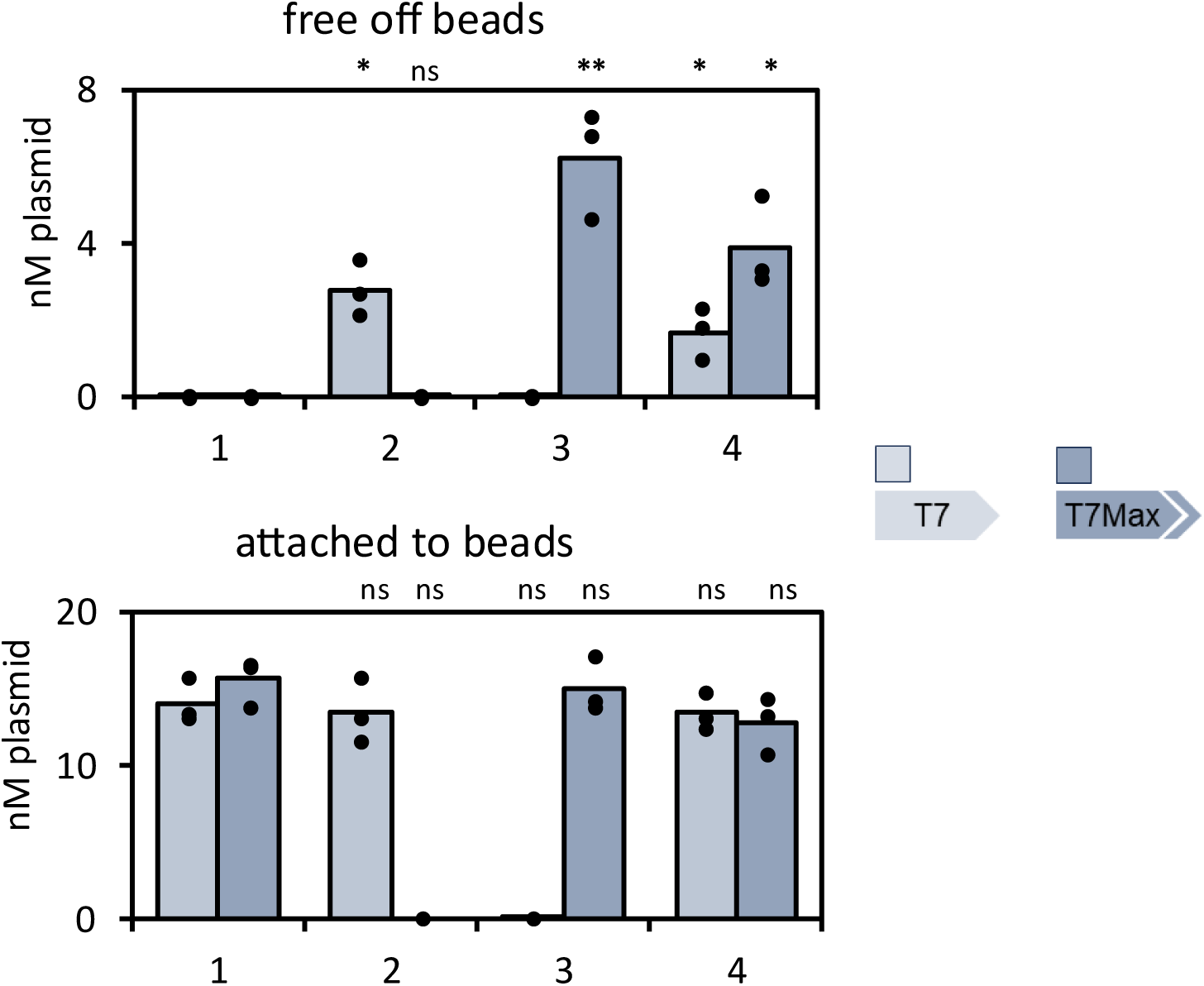
DNA abundance in synthetic cells after competitive genetically encoded division on beads with the His tagged αHL. Individual data points for each independent replicate experiment of measurement of amount of unique DNA marker in T7 and T7Max synthetic cells after competitive division on beads (Figure 6). The unique DNA marker was a plasmid not expressed inside synthetic cells (a gene under control of mammalian CMV promoter), it was used here only to mark otherwise identical synthetic cells, since T7 and T7Max promoters are similar enough that it’s difficult to reliably differentiate them with qPCR primers. See methods section 21 for details. <u>Sample 1</u>: T7 cells present, T7Max cells present; no division is expected. <u>Sample 2</u>: T7 cells present, T7Max cells absent; division is expected. <u>Sample 3</u>: T7 cells absent, T7Max cells present; division is expected. <u>Sample 4</u>: T7 cells present, T7Max cells present; division is expected. Light blue: T7 cells, darker blue: T7Max cells. Dots on top of the bars indicate all the individual data points, the bars are the average value. P values were calculated for samples compared to “Sample 1” T7 cells present, T7Max cells present; no division is expected. ns = P > 0.0; * = P ≤ 0.0, ** = P ≤ 0.01, *** = P ≤ 0.001. Exact P values are listed in **Table S12**. This data was collected using PURE pREP.

**Figure S68.**
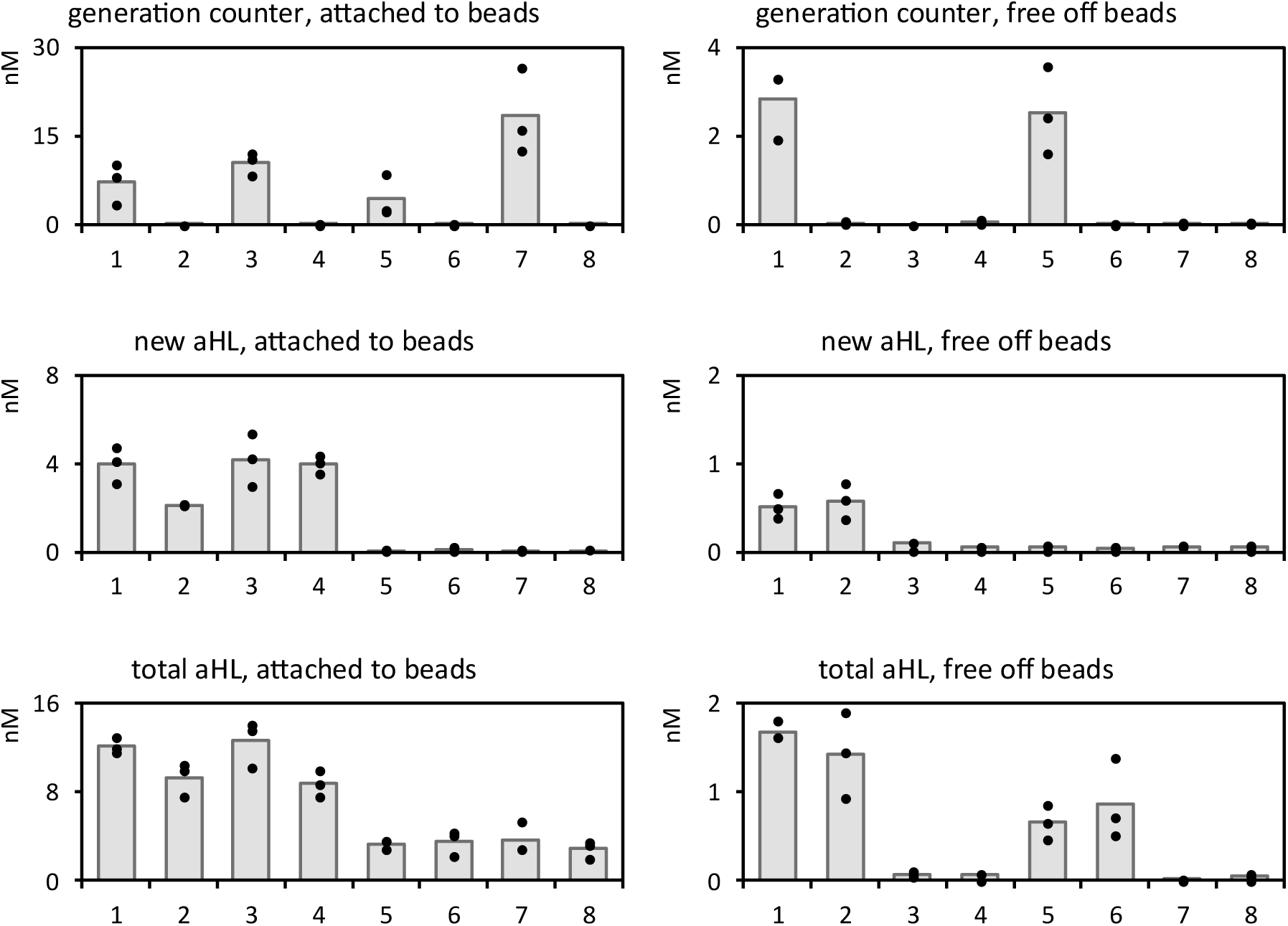
Individual data points for genetically encoded growth and division. Samples are in the same order as corresponding box and whiskers plots on Figure 6: <u>Sample 1</u>: DNA replication, growth, division <u>Sample 2</u>: DNA replication, no growth, division <u>Sample 3</u>: DNA replication, growth, no division <u>Sample 4</u>: DNA replication, no growth, no division <u>Sample 5</u>: no DNA replication, growth, division <u>Sample 6</u>: no DNA replication, no growth, division <u>Sample 7</u>: no DNA replication, growth, no division <u>Sample 8</u>: no DNA replication, no growth, no division Bar graphs represent average value, dots are individual data points for each replicate. Statistical analysis: P values are listed in **Table S12**. This data was collected using PURE pREP.

**Figure S69.**
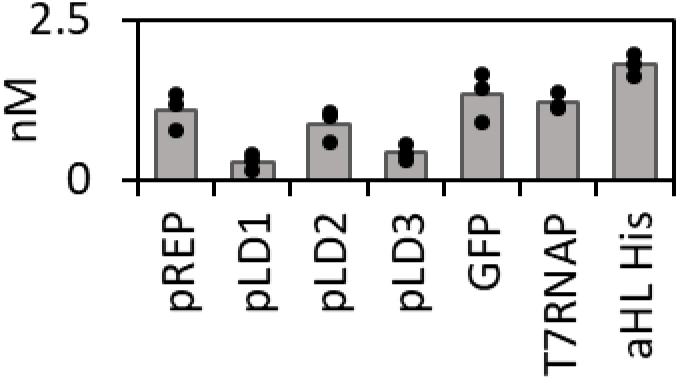
Detection of each plasmid in the synthetic cell genome after growth and division. qPCR analysis of each of the synthetic cell genome plasmids in “daughter” cell fraction of synthetic cells, the fraction that was collected off the magnetic beads, after growth and division. Those cells had DNA replication via Phi29. The αHL quantification did not differentiate between His and FLAG labeled proteins. Because some other proteins in the genome are also His tagged, it was impossible to create a qPCR primer for His tag that would be specific only for the αHL plasmid. Therefore, here we quantified αHL gene in the untagged region, so the results demonstrate combined quantification for both FLAG tagged and His tagged αHL plasmids. The bar graphs represent average value, dots on top of the bar graphs indicate individual values of three replicates. This data was collected using PURE pREP.

**Figure S70.**
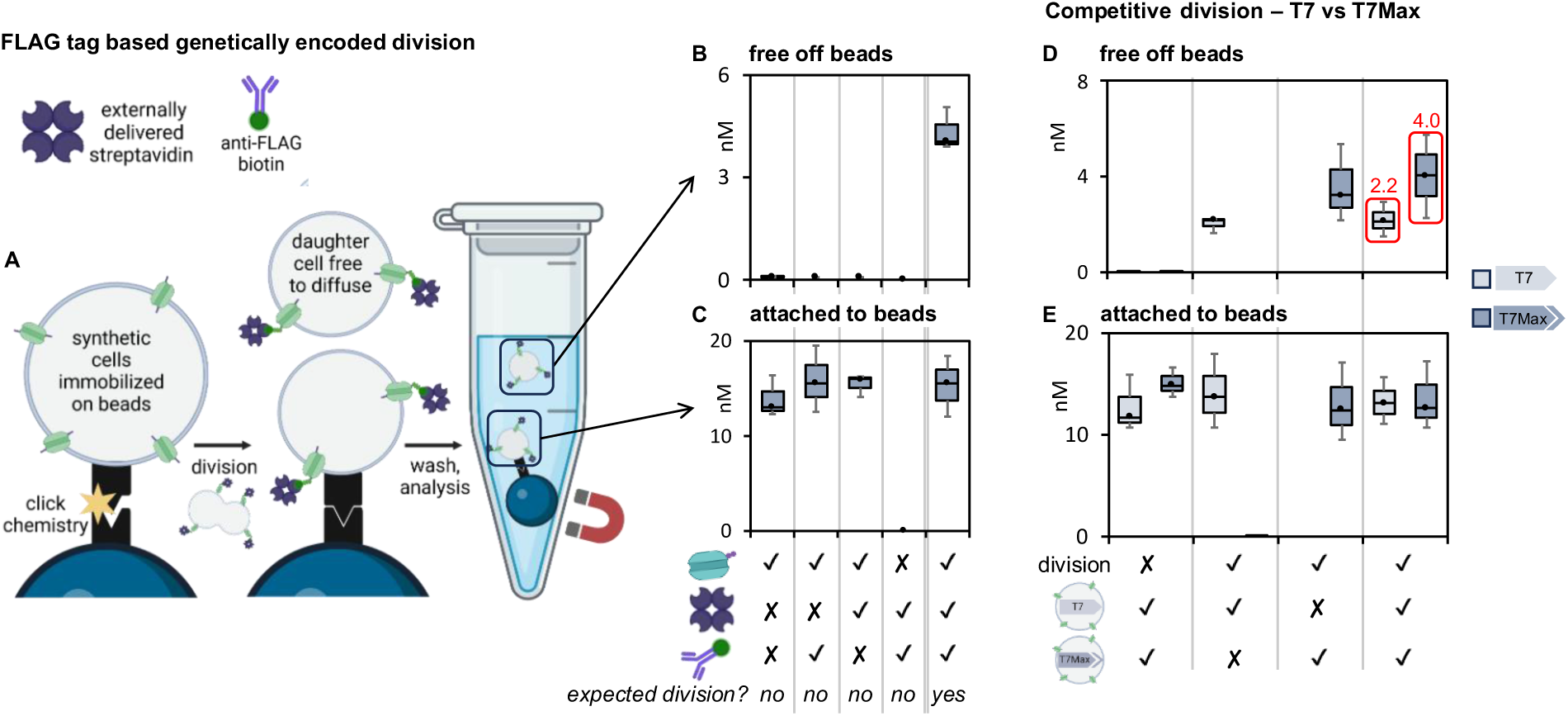
Genetically encoded division using FLAG system. **A**: Synthetic cells expressing FLAG tagged αHL are immobilized on magnetic beads. Streptavidin and biotin conjugated anti-FLAG antibody linker are added to the outside of the cells. If division occurs, the daughter cells are free to separate from the beads. After the experiment, free synthetic cells are collected separately from the synthetic cells that remained on the beads. **B** and **C**: quantification of αHL DNA in synthetic cells freed from the beads (the “daughter cells”, panel **B**) and cells that remained attached to the beads (the “parent” cells, panel **C**), in experiments excluding one of the essential division components (αHL, streptavidin or the biotin conjugated anti-FLAG antibody linker), and with all division elements present. Individual data points are in **Figure S71**. **D** and **E**: Competitive division. Synthetic cells containing αHL under “slow” promoter T7 and αHL under “fast” promoter T7Max are dividing in the same reaction. Samples were analyzed for the presence of unique DNA marker (different for T7 and T7Max cells). Label “division” indicates either presence of streptavidin and biotin conjugated anti-FLAG antibody linker (division label “yes”), or absence of both streptavidin and linker (labeled “no” division). In each experiment, the sample contained either a mix of both synthetic cells containing T7 and T7Max αHL in equal amounts, or only one of the populations (only T7 or only T7Max). After incubation and division, unique DNA marker was quantified, identifying T7 and T7Max populations in “daughter” cells free off the beads (panel **D**) and in cells that remained attached to beads (panel **E**). Individual data points are in **Figure S72**. This data was collected using PURE pREP.

**Figure S71.**
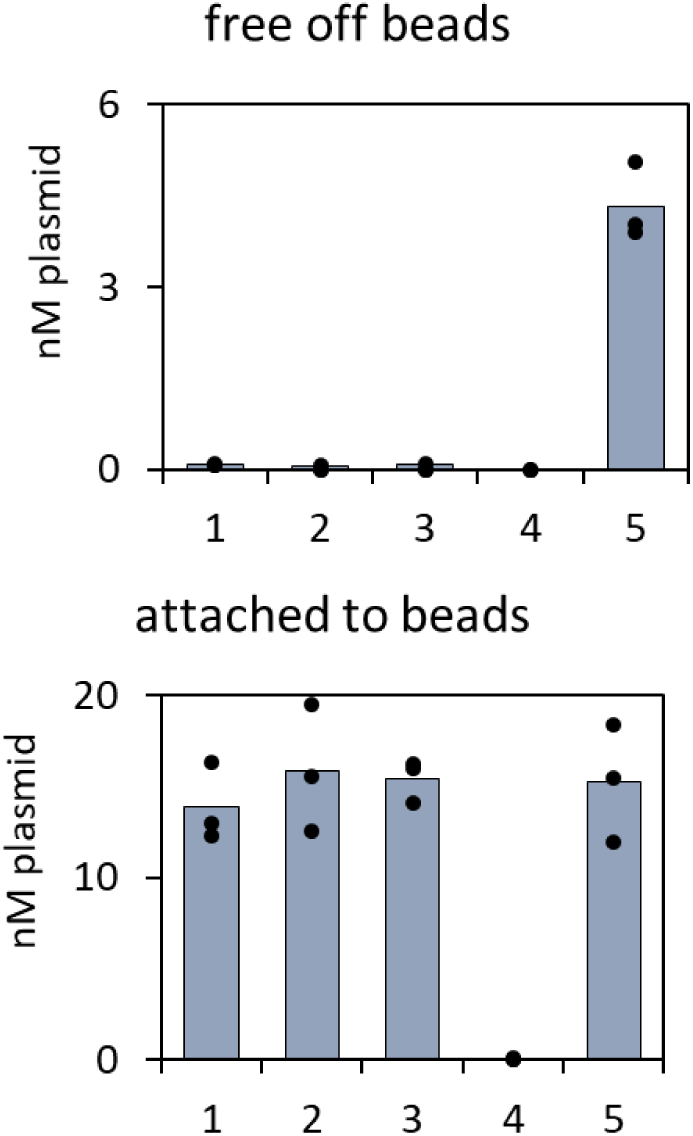
DNA abundance in synthetic cells after genetically encoded division on beads with the FLAG tagged αHL. Individual data points for each independent replicate experiment of measurement of amount of αHL DNA in T7Max synthetic cells after division on beads (**Figure S70**). <u>Sample 1</u>: αHL present, streptavidin absent, biotin conjugated anti-FLAG antibody linker absent; no division is expected. <u>Sample 2</u>: αHL present, streptavidin absent, biotin conjugated anti-FLAG antibody linker present; no division is expected. <u>Sample 3</u>: αHL present, streptavidin present, biotin conjugated anti-FLAG antibody linker absent; no division is expected. <u>Sample 4</u>: αHL absent, streptavidin present, biotin conjugated anti-FLAG antibody linker present; no division is expected. <u>Sample 5</u>: αHL present, streptavidin present, biotin conjugated anti-FLAG antibody linker present; division is expected. Dots on top of the bars indicate all the individual data points, the bars are the average value. This data was collected using PURE pREP.

**Figure S72.**
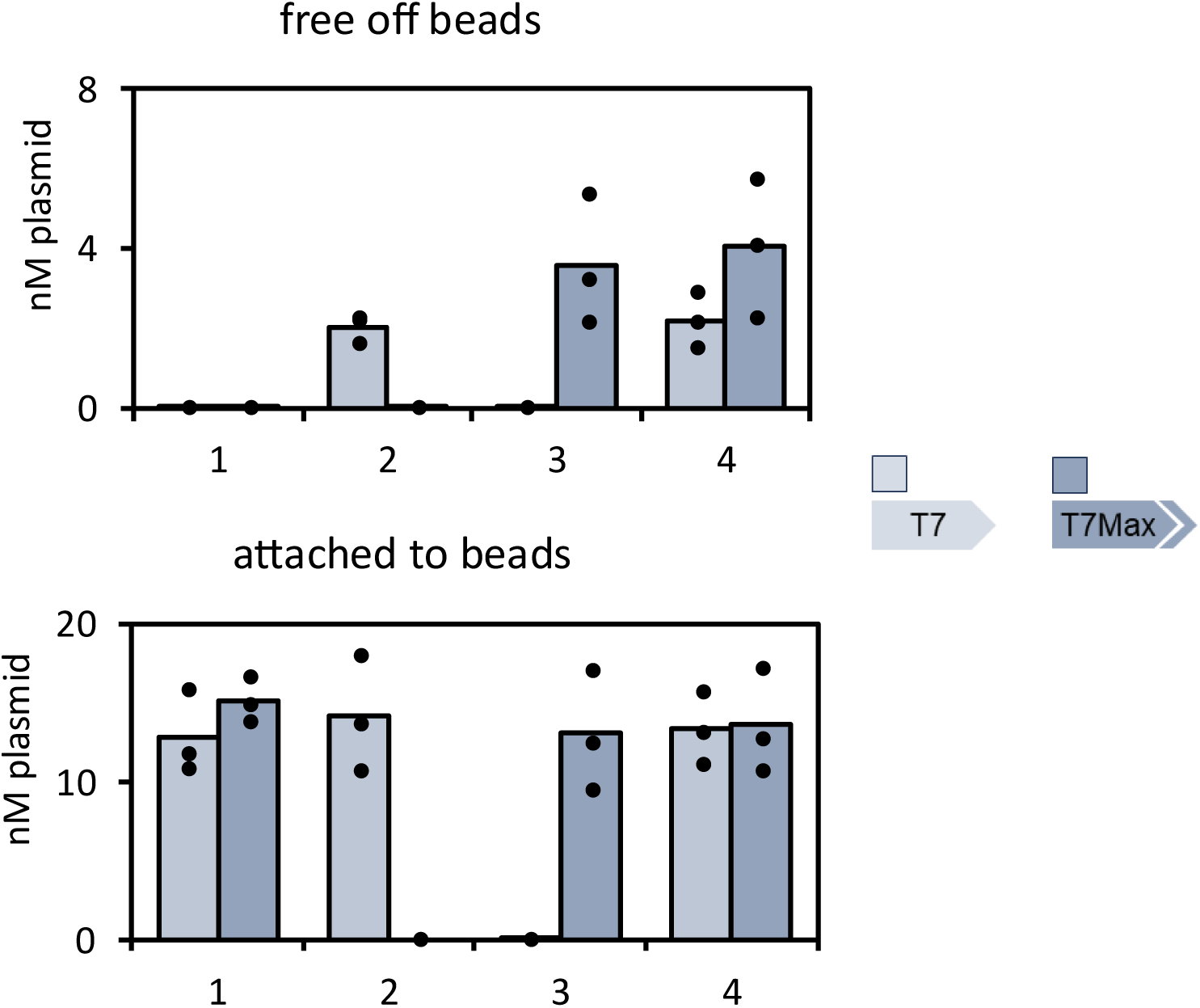
DNA abundance in synthetic cells after competitive genetically encoded division on beads with the FLAG tagged αHL. Individual data points for each independent replicate experiment of measurement of amount of unique DNA marker in T7 and T7Max synthetic cells after competitive division on beads (**Figure S70**). The unique DNA marker was a plasmid not expressed inside synthetic cells (a gene under control of mammalian CMV promoter), it was used here only to mark otherwise identical synthetic cells, since T7 and T7Max promoters are similar enough that it’s difficult to reliably differentiate them with qPCR primers. See methods section 21 for details. <u>Sample 1</u>: T7 cells present, T7Max cells present; no division is expected. <u>Sample 2</u>: T7 cells present, T7Max cells absent; division is expected. <u>Sample 3</u>: T7 cells absent, T7Max cells present; division is expected. <u>Sample 4</u>: T7 cells present, T7Max cells present; division is expected. Light blue: T7 cells, darker blue: T7Max cells. Dots on top of the bars indicate all the individual data points, the bars are the average value. This data was collected using PURE pREP.

**Figure S73.**
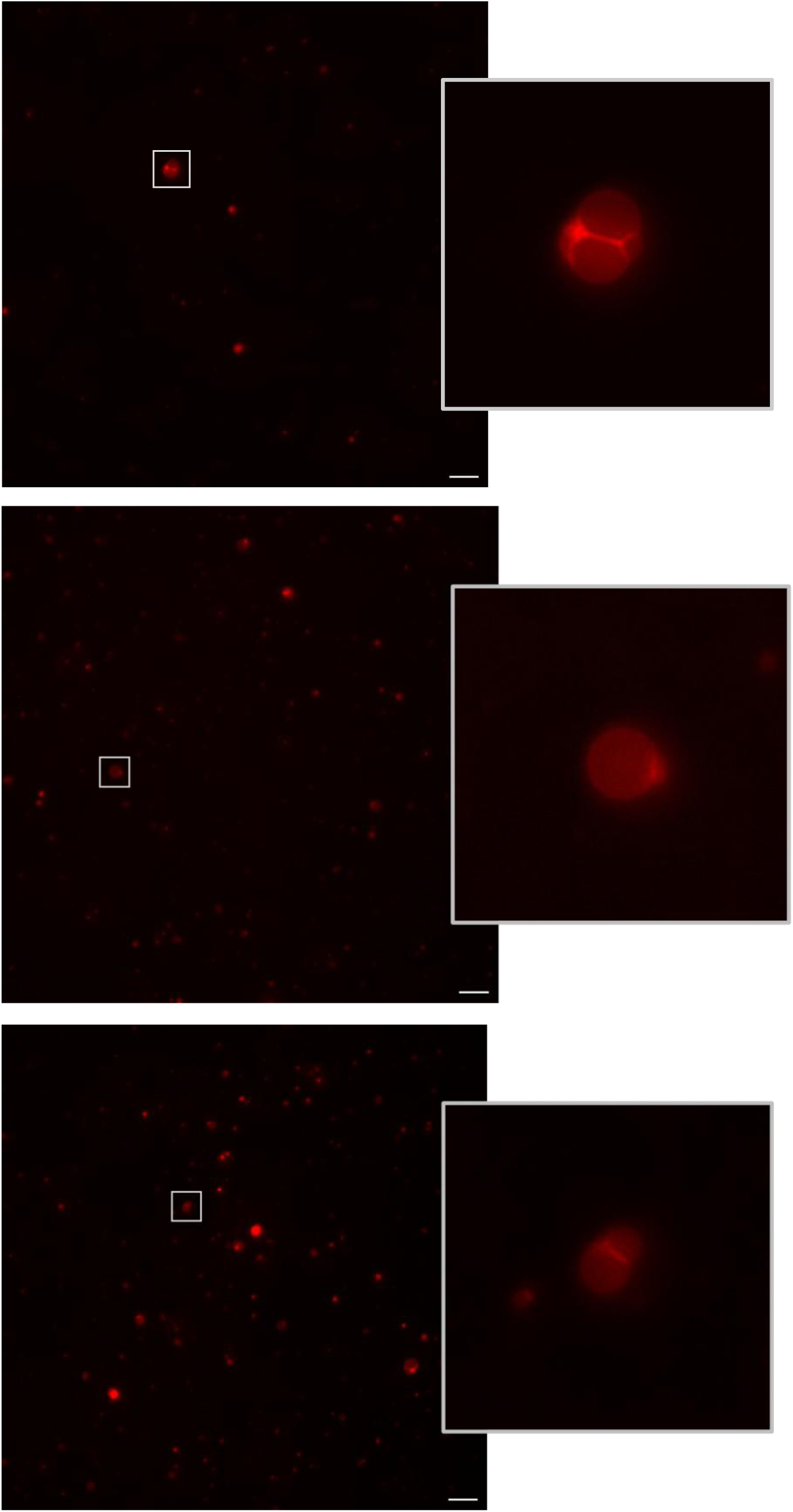
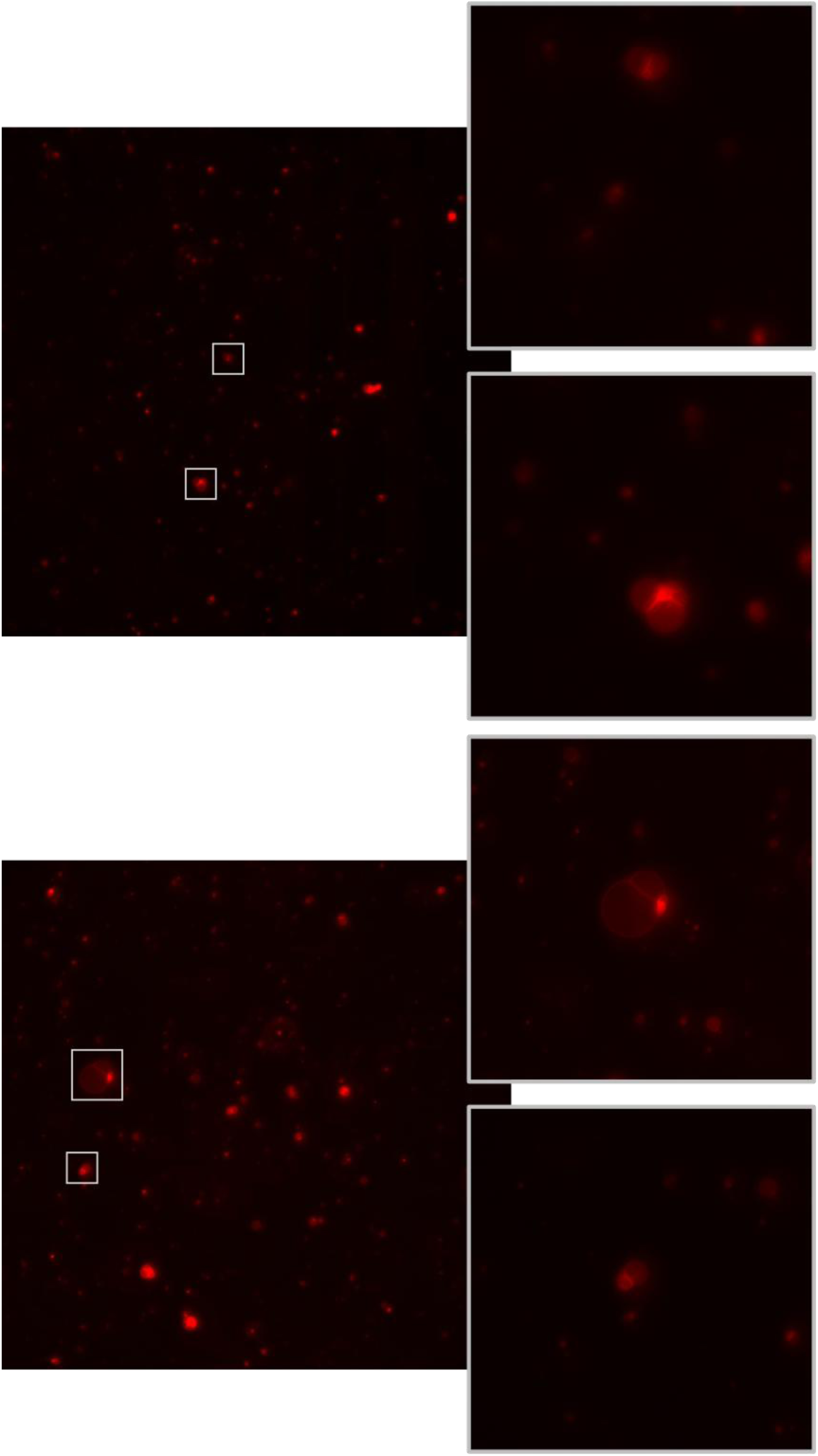
Fluorescent microscopy of synthetic cell undergoing a cycle of growth and division. (His tag mediated growth, Flag mediated division). The membrane of synthetic cell is labeled with Rhodamine. Scale bar is 20µm. The larger image is the entire field of view, the smaller image insert is zooming in on the cell of interest. This data was collected using PURE pREP.

**Figure S74.**
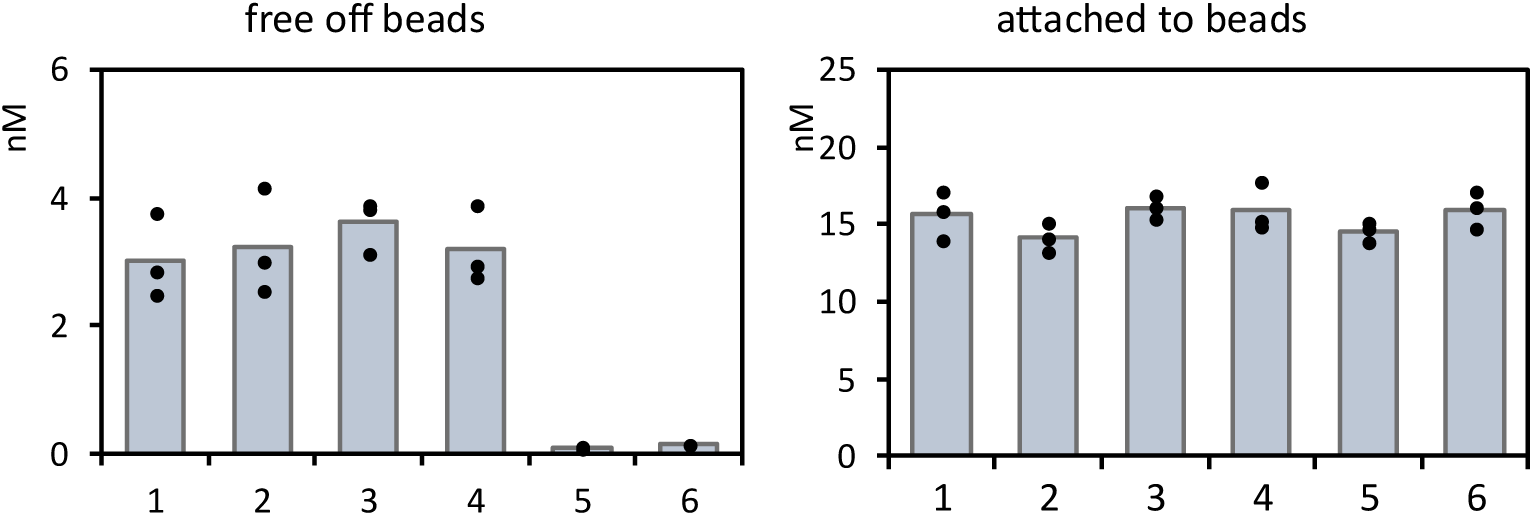
Measuring interference and cross-reactivity of His and Flag based division systems. The αHL plasmid abundance measured after 12h division experiment, performed according to protocol described in Methods section 21, with samples on beads according to protocol described in Methods section 20, under conditions indicated below in sample description. All samples contained streptavidin. <u>Sample 1</u>: Cells with αHL-FLAG and biotin FLAG antibody linker. Division is expected. <u>Sample 2</u>: Cells with αHL-His and Ni-NTA linker. Division is expected. <u>Sample 3</u>: Interference test - cells with αHL-FLAG and both biotin FLAG antibody and Ni-NTA linkers. Division is expected, if the His and Flag systems don’t interfere with each other. <u>Sample 4</u>: Interference test - cells with αHL-His and both biotin FLAG antibody and Ni-NTA linkers. Division is expected, if the His and Flag systems don’t interfere with each other. <u>Sample 5</u>: Cross-reactivity test - cells with αHL-FLAG and Ni-NTA linker. Division is not expected if the His and Flag systems don’t cross-react. <u>Sample 6</u>: Cross-reactivity test - cells with αHL-His and biotin FLAG antibody linker. Division is not expected if the His and Flag systems don’t cross-react. Bar graphs represent average values; dots represent individual data points. This data was collected using PURE pREP.

**Figure S75.**
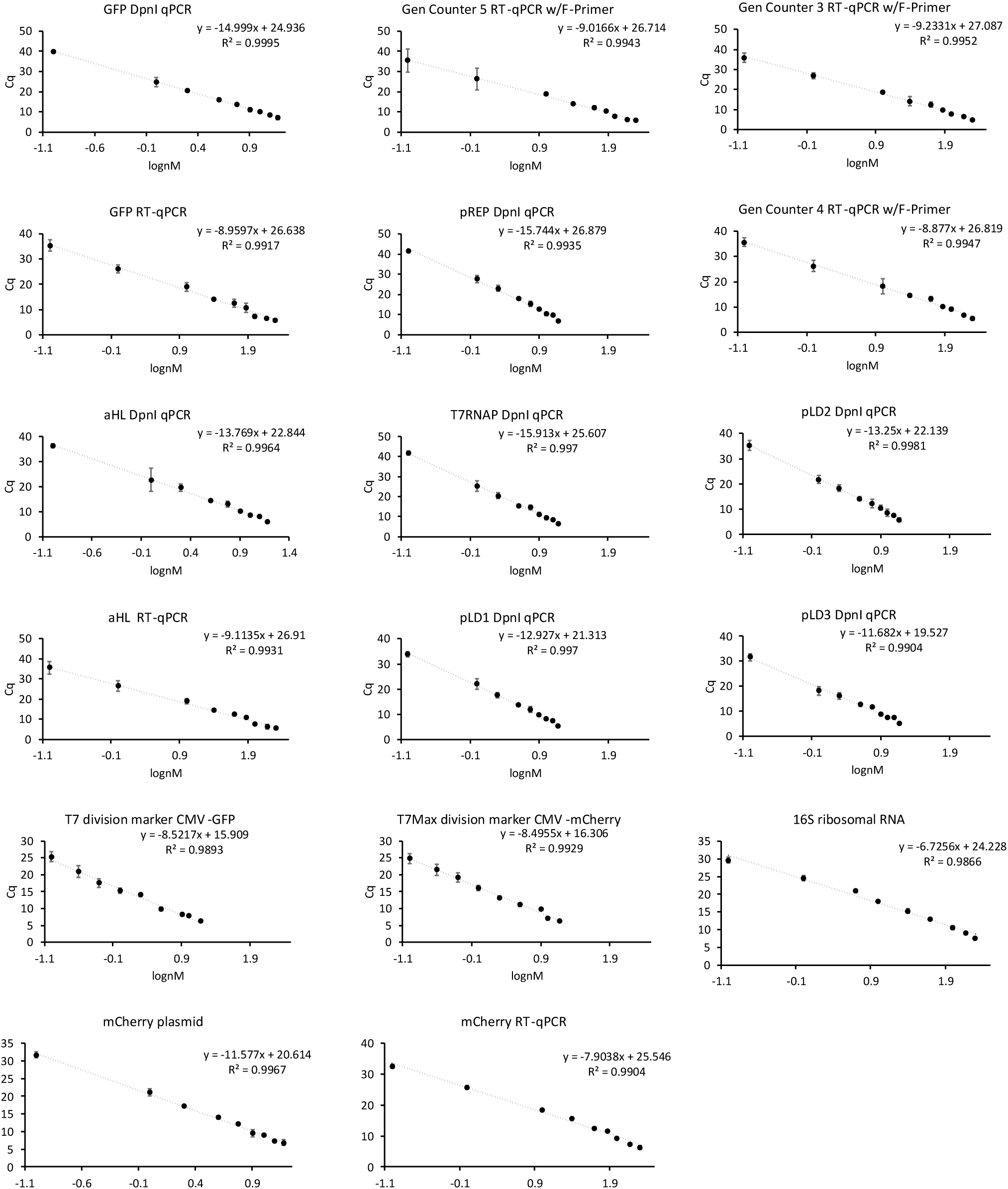
Calibration curves for qPCR experiments. Each data point is average of three experiments, error bars show SEM.

**Figure S76.**
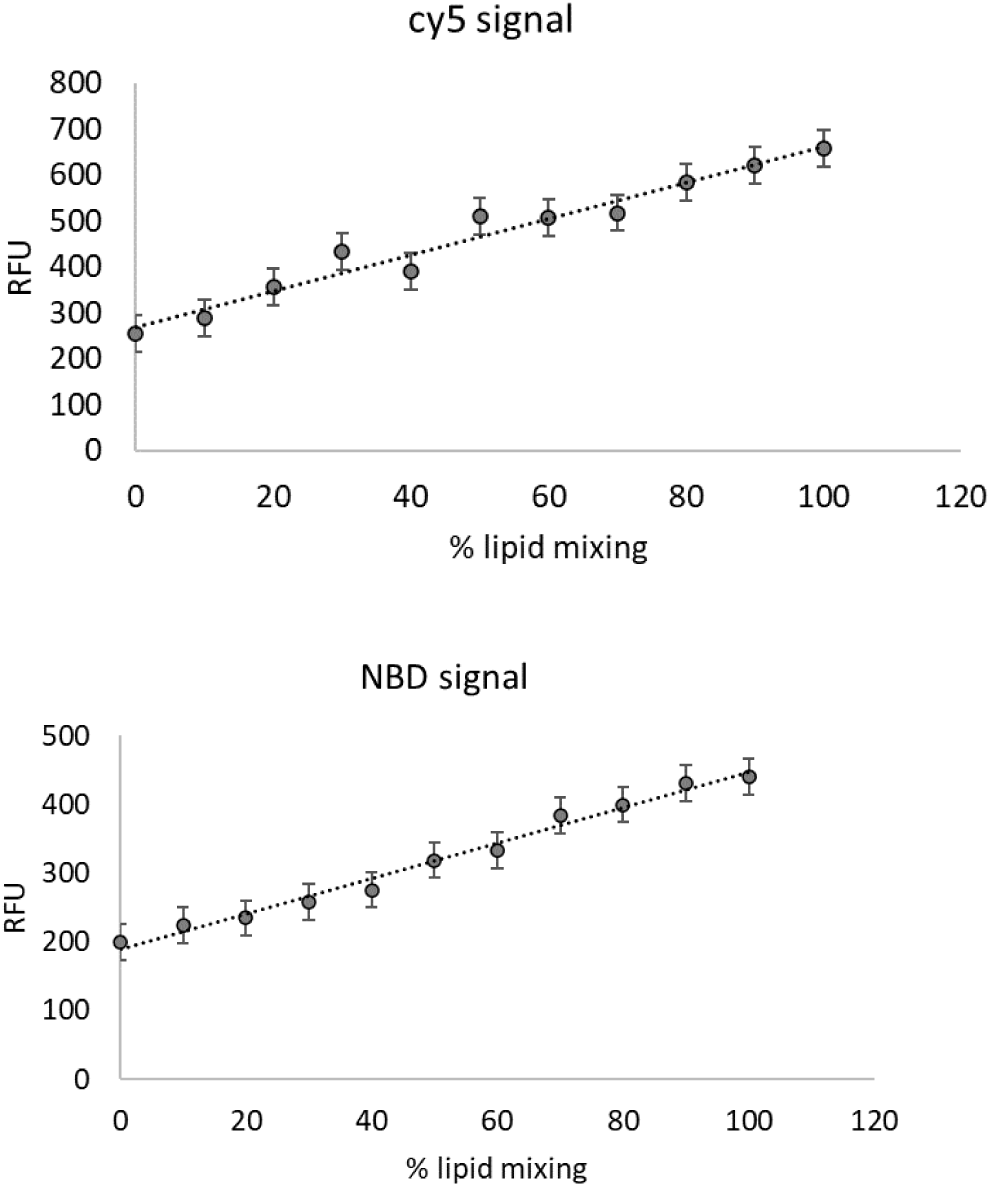
FRET calibration curves. Calibration for NBD fluorescence in NBD – rhodamine FRET experiments, and Cy5 fluorescence for Cy5 – Cy7 FRET experiments. Error bars indicate SEM, n=3 independently prepared and processed liposome samples.

**Table S1:**
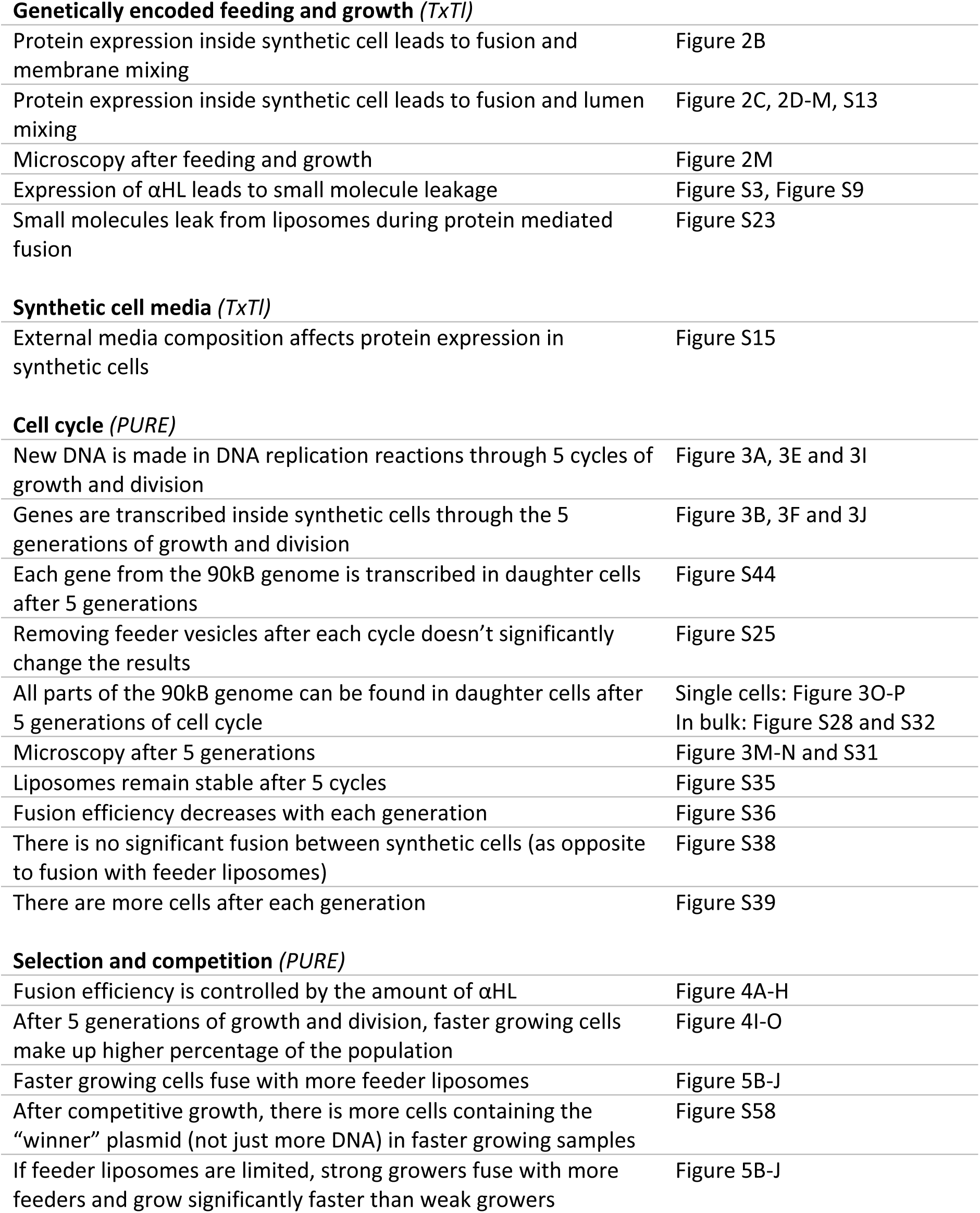
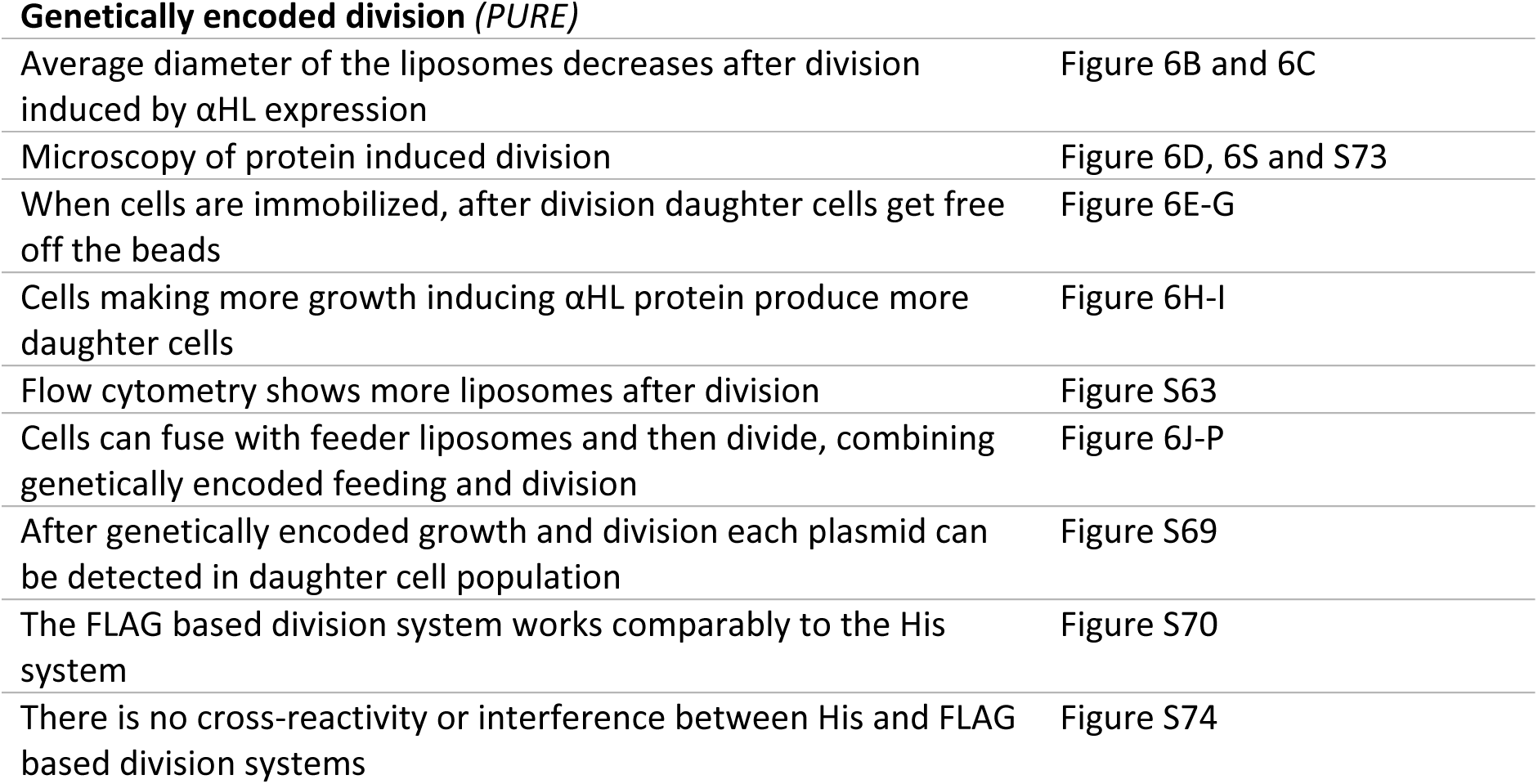
Summary of major results with reference to figures containing the information.

|  |  |
| --- | --- |
| Protein expression inside synthetic cell leads to fusion and membrane mixing | Figure 2B |
| Protein expression inside synthetic cell leads to fusion and lumen mixing | Figure 2C, 2D-M, S13 |
| Microscopy after feeding and growth | Figure 2M |
| Expression of αHL leads to small molecule leakage | Figure S3, Figure S9 |
| Small molecules leak from liposomes during protein mediated fusion | Figure S23 |
| External media composition affects protein expression in synthetic cells | Figure S15 |
| **Cell cycle (*PURE*)** |  |
| New DNA is made in DNA replication reactions through 5 cycles of growth and division | Figure 3A, 3E and 3I |
| Genes are transcribed inside synthetic cells through the 5 generations of growth and division | Figure 3B, 3F and 3J |
| Each gene from the 90kB genome is transcribed in daughter cells after 5 generations | Figure S44 |
| Removing feeder vesicles after each cycle doesn't significantly change the results | Figure S25 |
| All parts of the 90kB genome can be found in daughter cells after 5 generations of cell cycle | Single cells: Figure 3O-P<br>In bulk: Figure S28 and S32 |
| Microscopy after 5 generations | Figure 3M-N and S31 |
| Liposomes remain stable after 5 cycles | Figure S35 |
| Fusion efficiency decreases with each generation | Figure S36 |
| There is no significant fusion between synthetic cells (as opposite to fusion with feeder liposomes) | Figure S38 |
| There are more cells after each generation | Figure S39 |
| **Selection and competition (*PURE*)** |  |
| Fusion efficiency is controlled by the amount of αHL | Figure 4A-H |
| After 5 generations of growth and division, faster growing cells make up higher percentage of the population | Figure 4I-O |
| Faster growing cells fuse with more feeder liposomes | Figure 5B-J |
| After competitive growth, there is more cells containing the "winner" plasmid (not just more DNA) in faster growing samples | Figure S58 |
| If feeder liposomes are limited, strong growers fuse with more feeders and grow significantly faster than weak growers | Figure 5B-J |
| Average diameter of the liposomes decreases after division induced by αHL expression | Figure 6B and 6C |
| Microscopy of protein induced division | Figure 6D, 6S and S73 |
| When cells are immobilized, after division daughter cells get free off the beads | Figure 6E-G |
| Cells making more growth inducing αHL protein produce more daughter cells | Figure 6H-I |
| Flow cytometry shows more liposomes after division | Figure S63 |
| Cells can fuse with feeder liposomes and then divide, combining genetically encoded feeding and division | Figure 6J-P |
| After genetically encoded growth and division each plasmid can be detected in daughter cell population | Figure S69 |
| The FLAG based division system works comparably to the His system | Figure S70 |
| There is no cross-reactivity or interference between His and FLAG based division systems | Figure S74 |

**Table S2:**
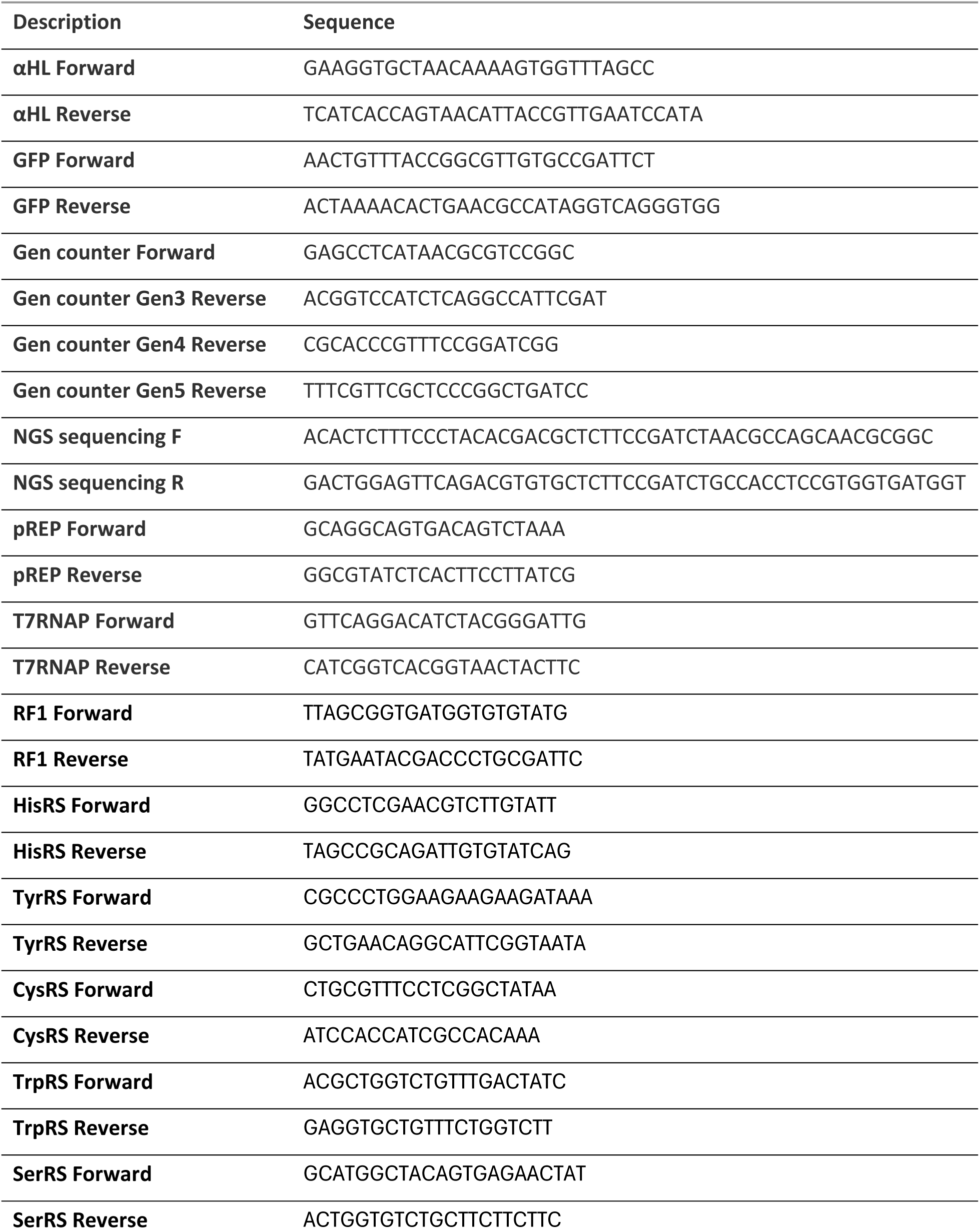
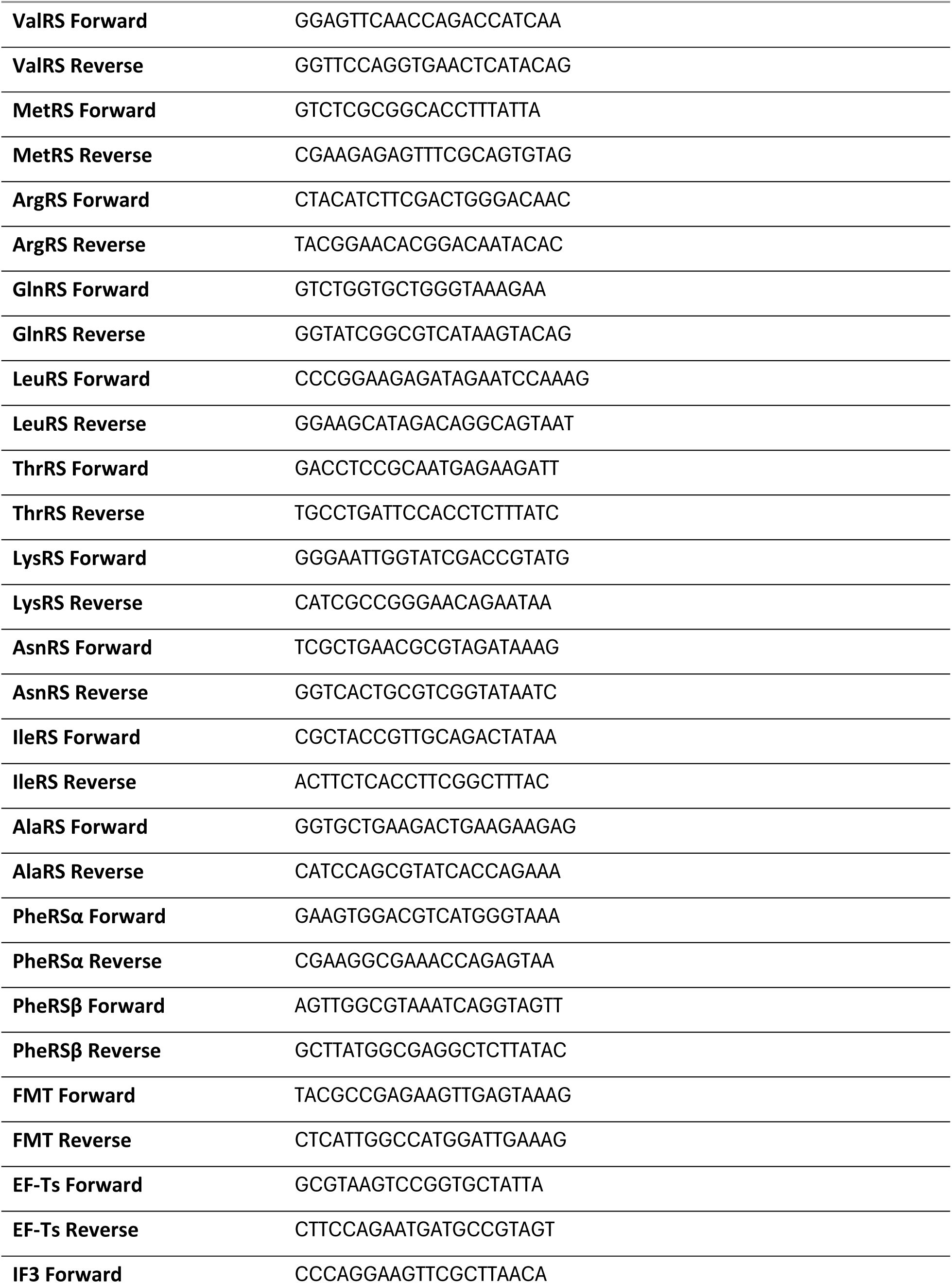
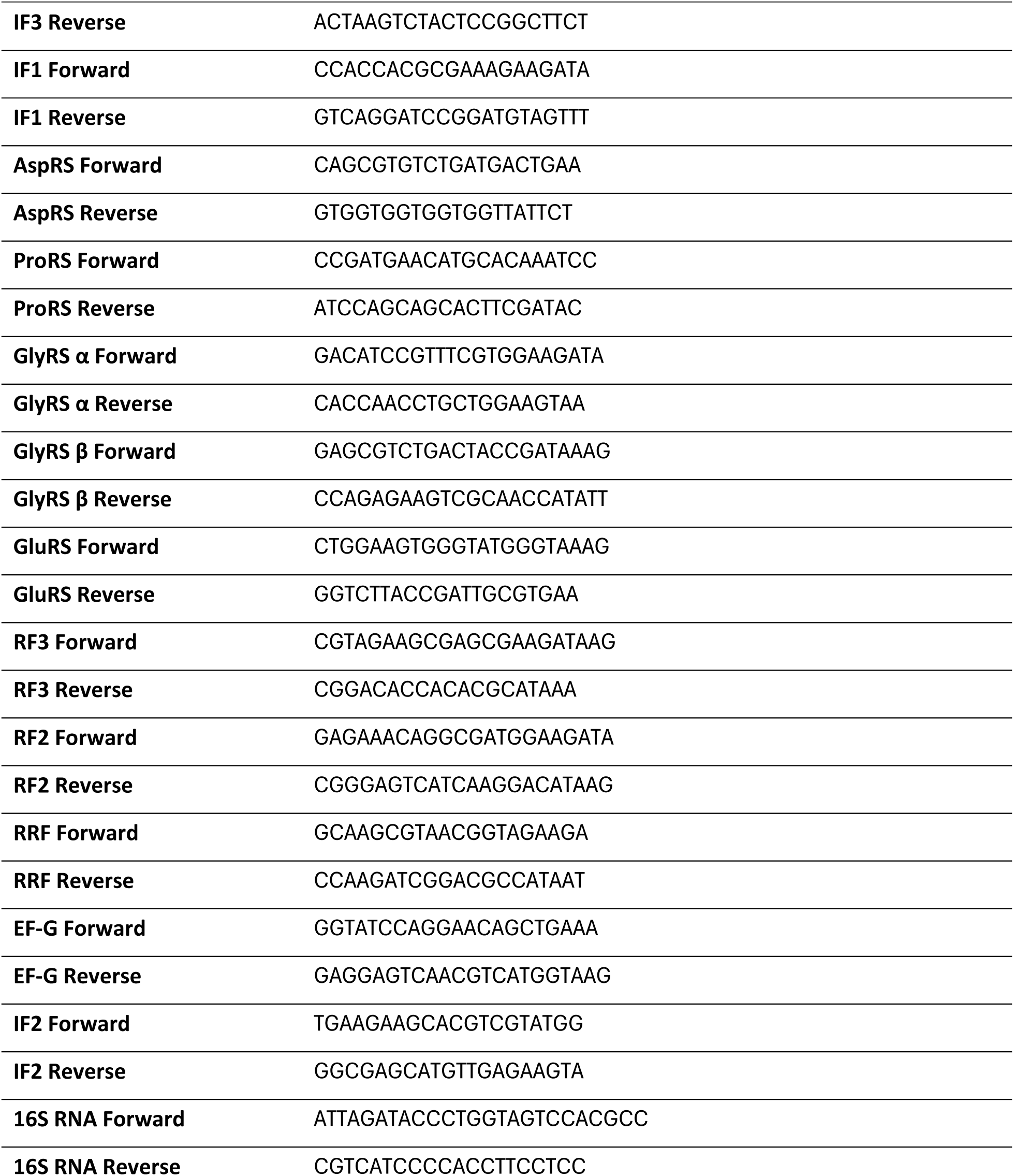
Sequences of primers used in this work.

**Table S3:**
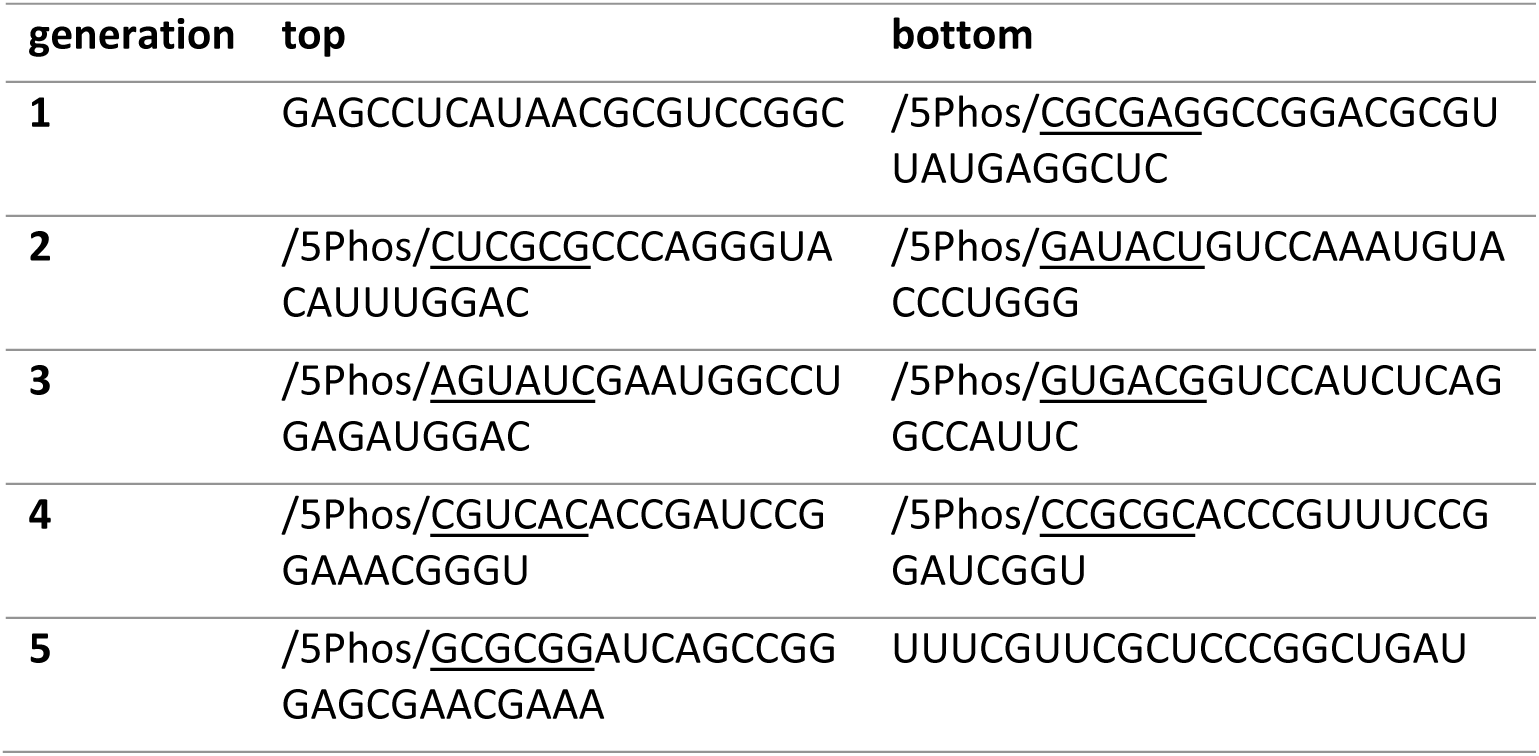
Sequences of RNA generation counter. Complementary overhangs are underlined. /5Phos/ is a 5’ Phosphorylation modification.

**Table S4:** Summary of trends in mRNA, DNA, and generation counter abundance in cell cycle, for data presented on Figure 3. Labels indicate: **∼** The result was ambiguous (no clear trend) **–** The analyte was not detected **+** The analyte was detected

|  |  | mRNA |  | *new* DNA |  | GFP | Generation counter |  |  |
| --- | --- | --- | --- | --- | --- | --- | --- | --- | --- |
|  |  | αHL | GFP | αHL | GFP | fluorescence | Gen3 | Gen4 | Gen5 |
| **Three generations** | Phi29, fusion ★ | ~ | ~ | + | + | + | + | - | - |
|  | no Phi29, fusion ◆ | ~ | ~ | - | - | + | + | - | - |
|  | Phi29, no fusion ▲ | ~ | ~ | + | + | + | - | - | - |
|  | no Phi29, no fusion ✖ | ~ | ~ | - | - | - | - | - | - |
| **Four generations** | Phi29, fusion ★ | ~ | ~ | + | + | + | + | + | - |
|  | no Phi29, fusion ◆ | ~ | ~ | - | - | + | + | + | - |
|  | Phi29, no fusion ▲ | ~ | ~ | + | + | + | - | - | - |
|  | no Phi29, no fusion ✖ | ~ | ~ | - | - | - | - | - | - |
| **Five generations** | Phi29, fusion ★ | ~ | ~ | + | + | + | + | + | + |
|  | no Phi29, fusion ◆ | ~ | ~ | - | - | + | + | + | + |
|  | Phi29, no fusion ▲ | ~ | ~ | + | + | + | - | - | - |
|  | no Phi29, no fusion ✖ | ~ | ~ | - | - | - | - | - | - |

**Table S5:**
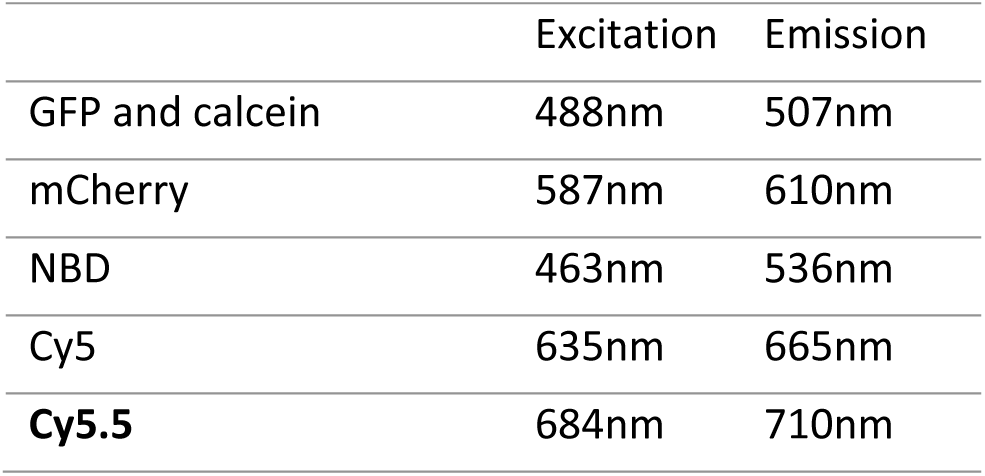
Wavelengths used in fluorescent measurements in this work.

**Table S6:** Summary of the composition of the whole 90.3kbp genome of the synthetic cell.

| plasmid | size | Proteins encoded |
| --- | --- | --- |
| pREP | 4.6kbp | Phi29 polymerase<sup>20</sup> |
| pLD1 | 30.1kbp | RF1, HisRS, TyrRS, CysRS, TrpRS, SerRS, ValRS, MetRS, ArgRS, GlnRS, LeuRS, ThrRS, LysRS<sup>65</sup> |
| pLD2 | 20.0kbp | AsnRS, IleRS, AlaRS, PheRS1, PheRS2, MTF, EF-Ts, IF3, IF1<sup>65</sup> |
| pLD3 | 23.4kbp | AspRS, ProRS, GlyRS1, GlyRS2, GluRS, RF3, RF2, RRF, EF-G, IF2<sup>65</sup> |
| GFP | 2.5kbp | Green fluorescent protein (GFP) |
| T7RNAP | 5.4kbp | T7 RNA polymerase<sup>89</sup> |
| αHL | 4.3kbp | alpha hemolysin (αHL) |

**Table S7:**
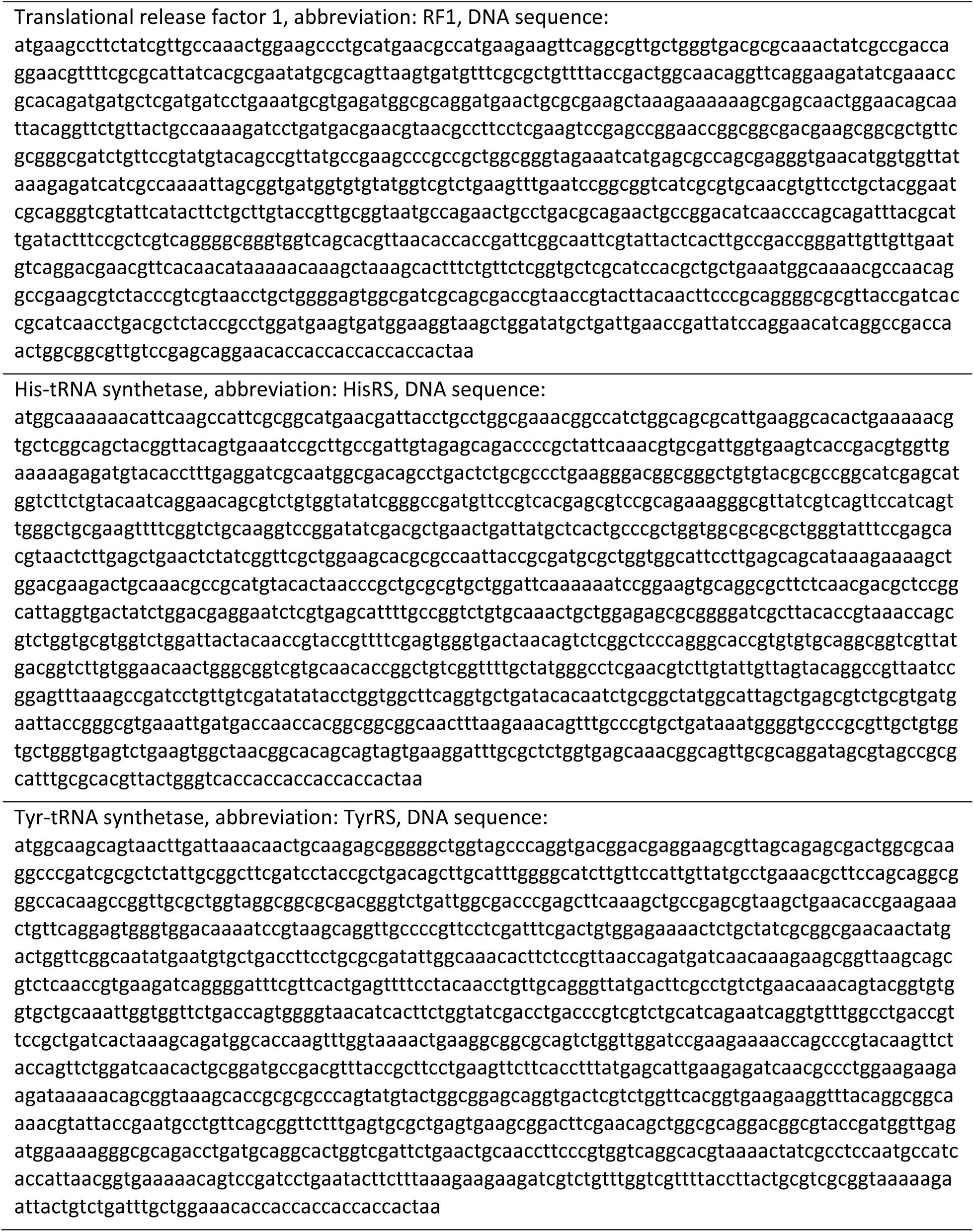
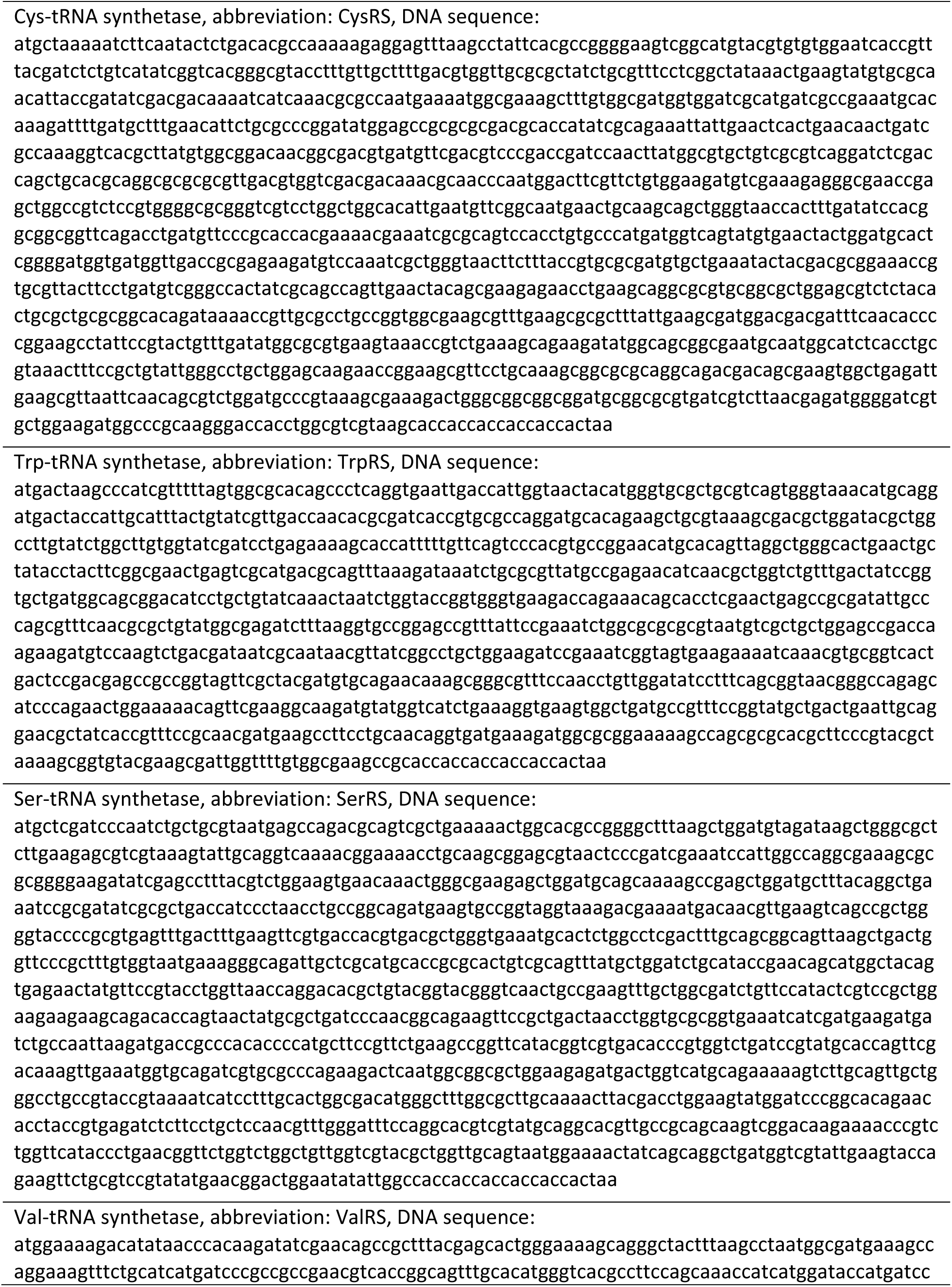
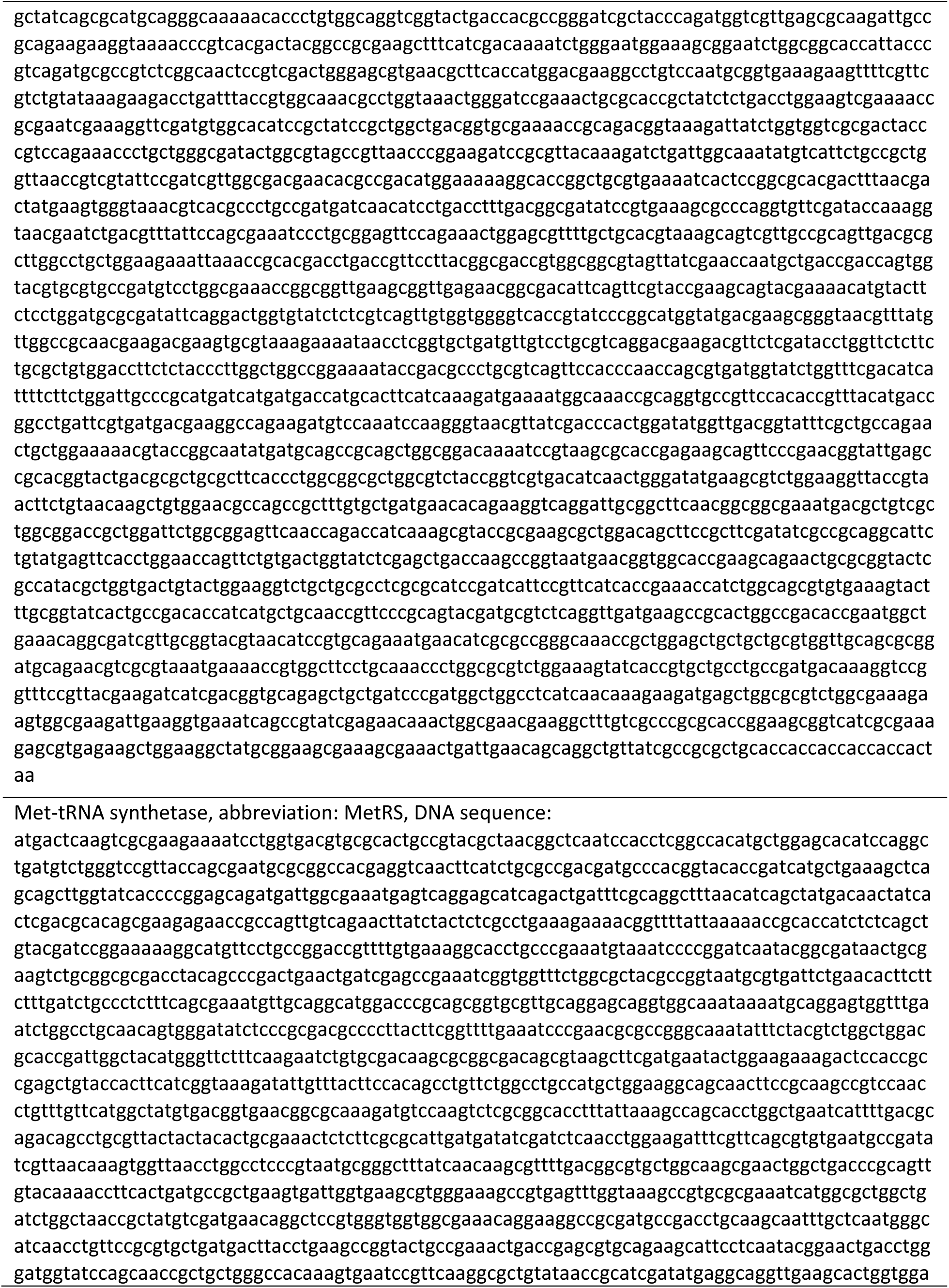
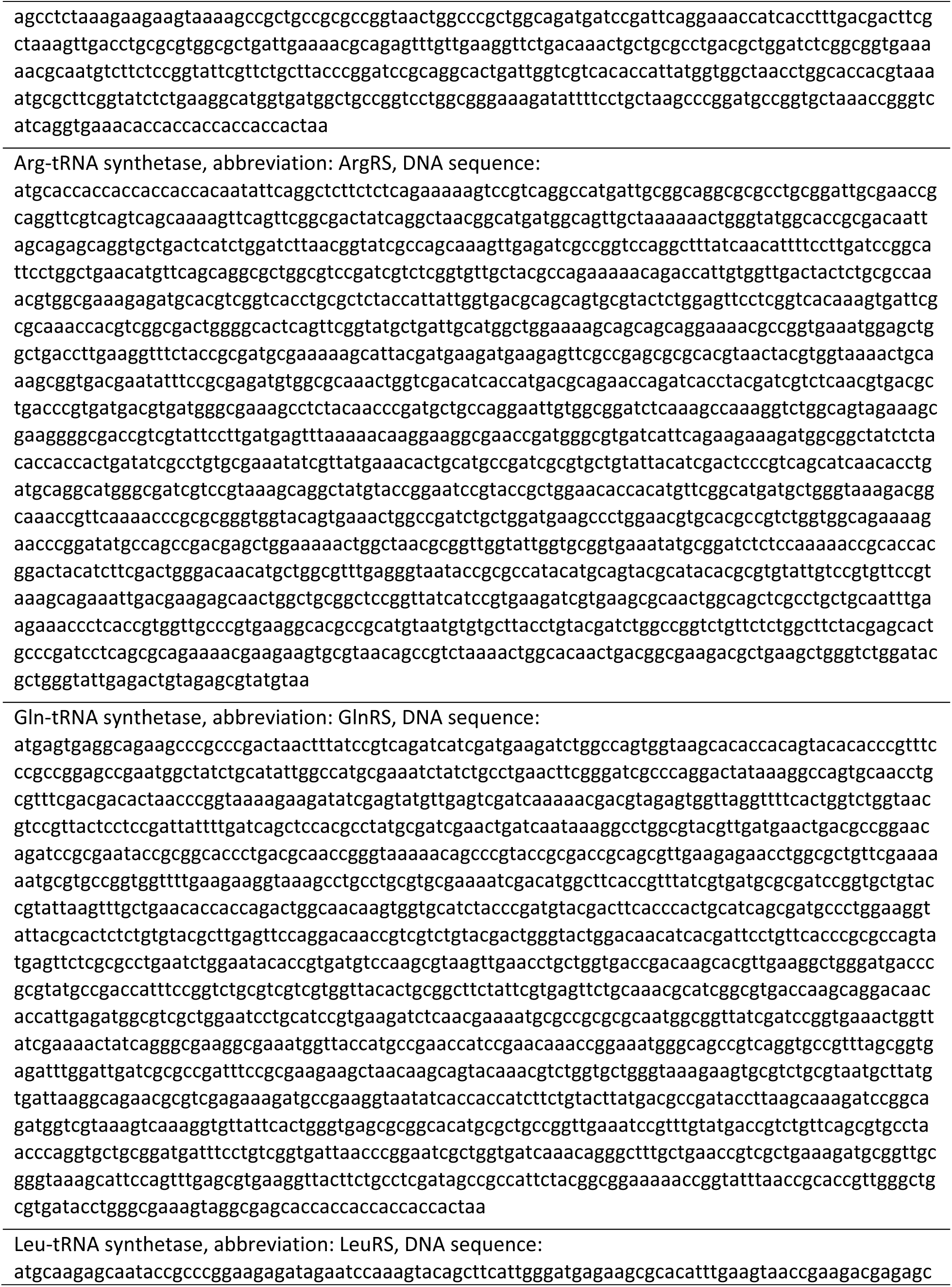
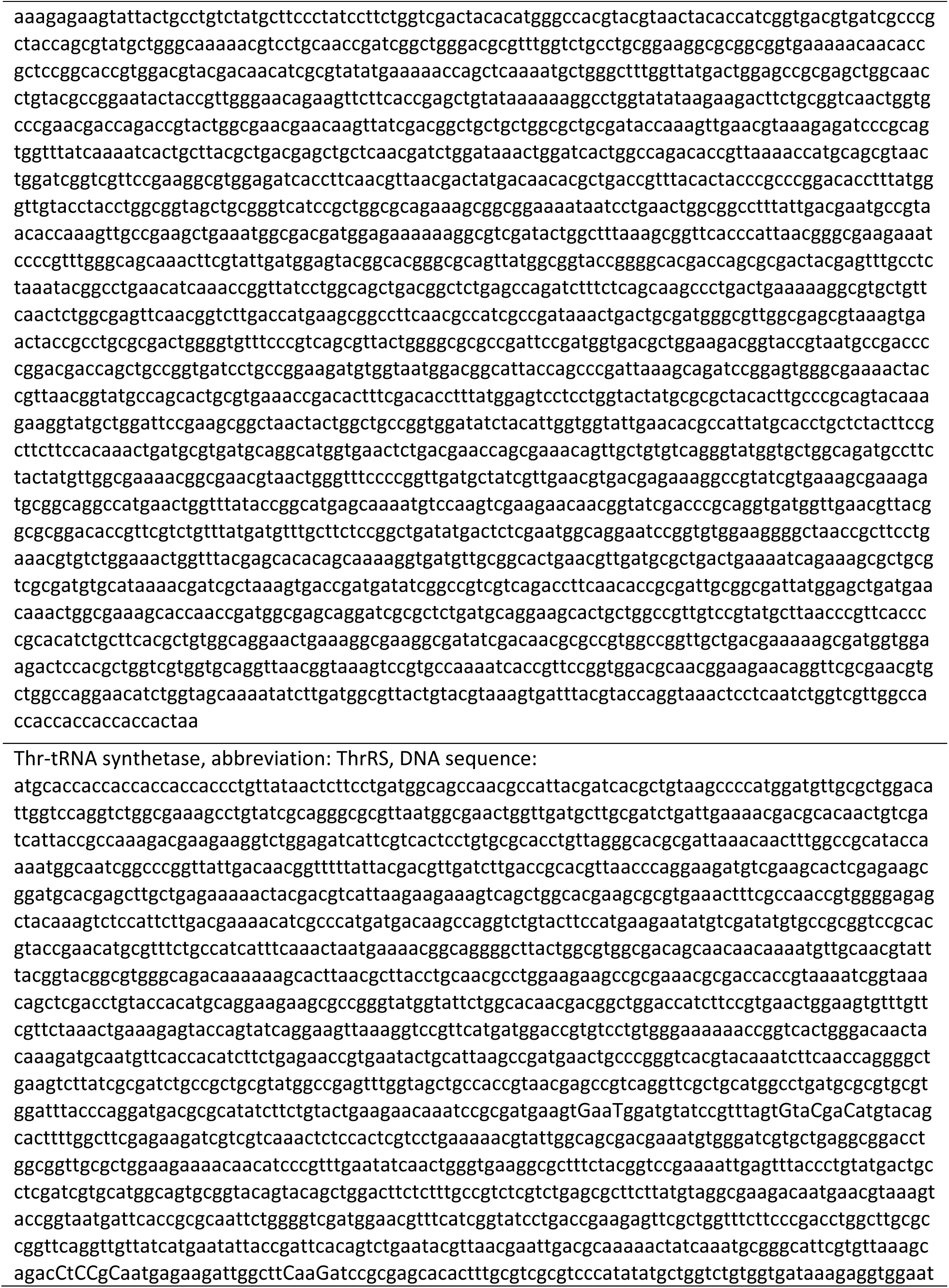
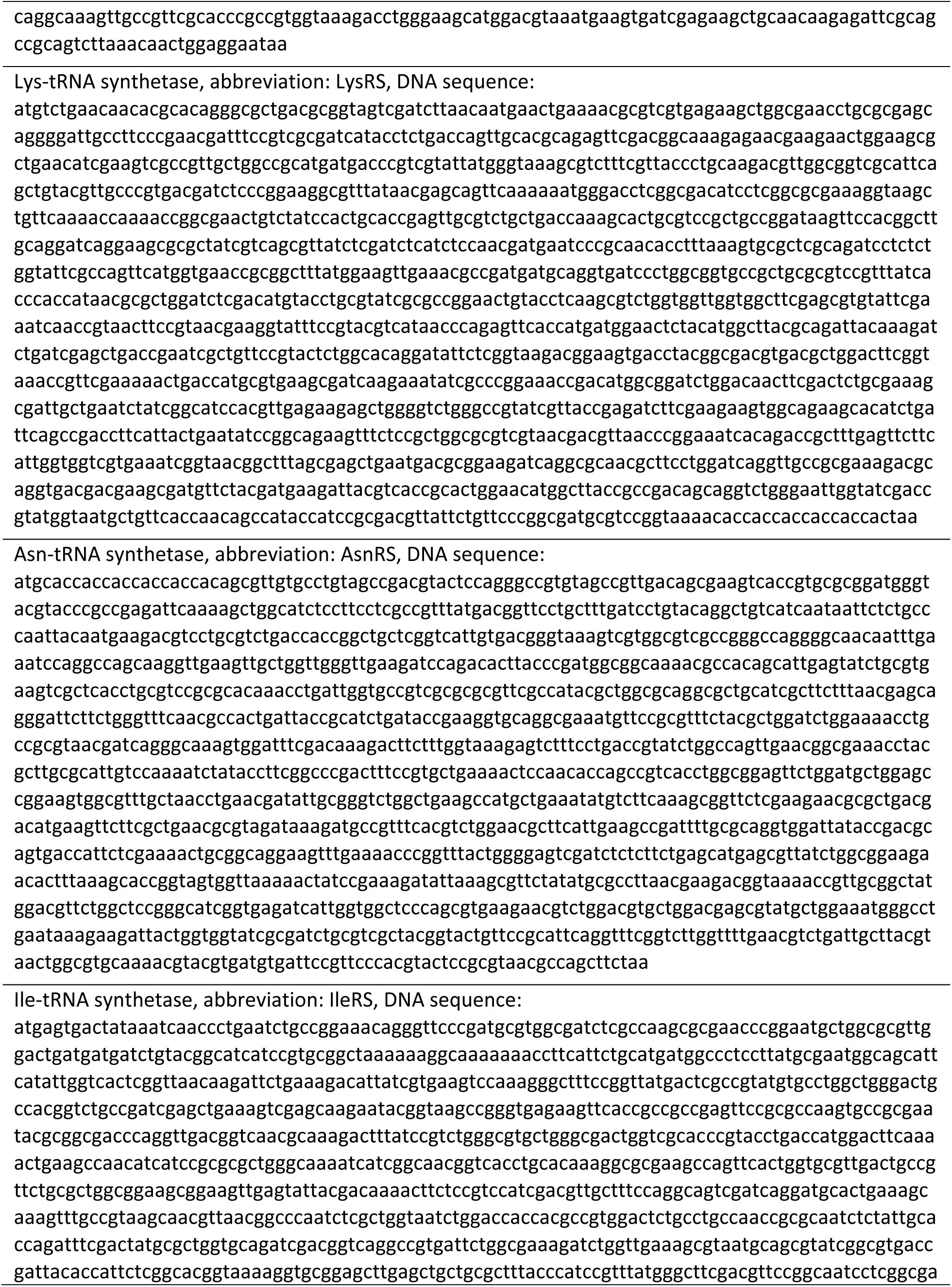
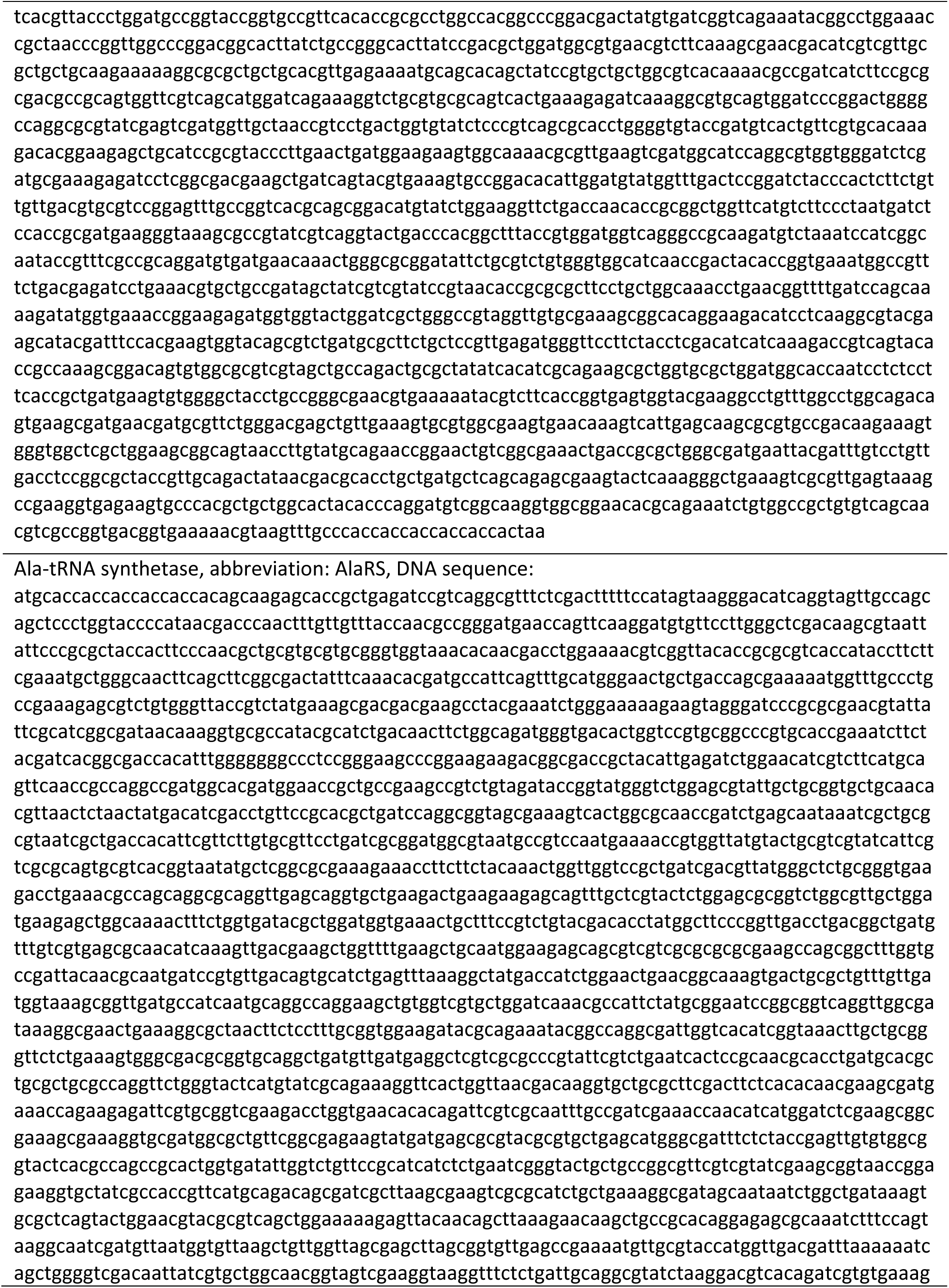
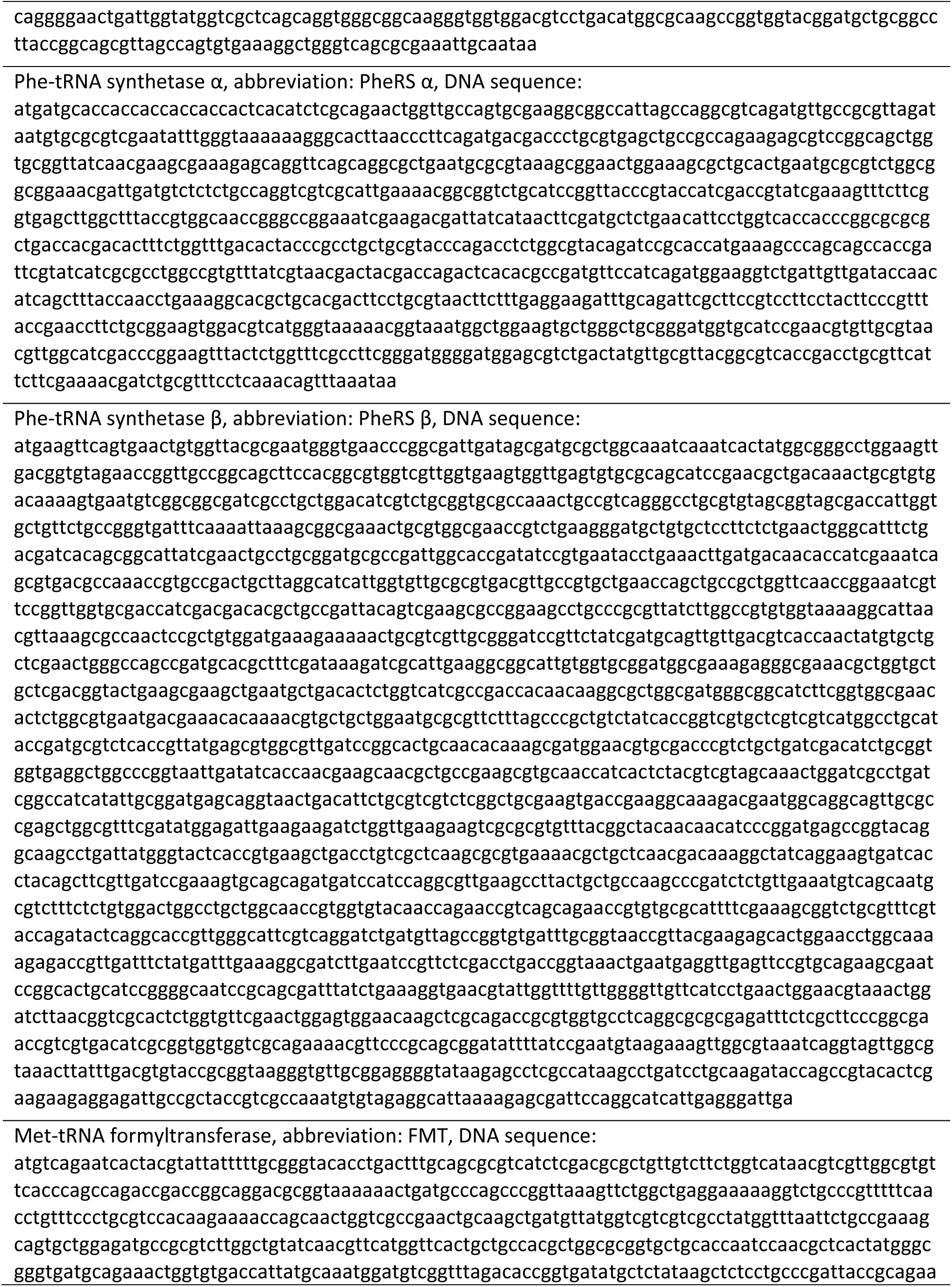
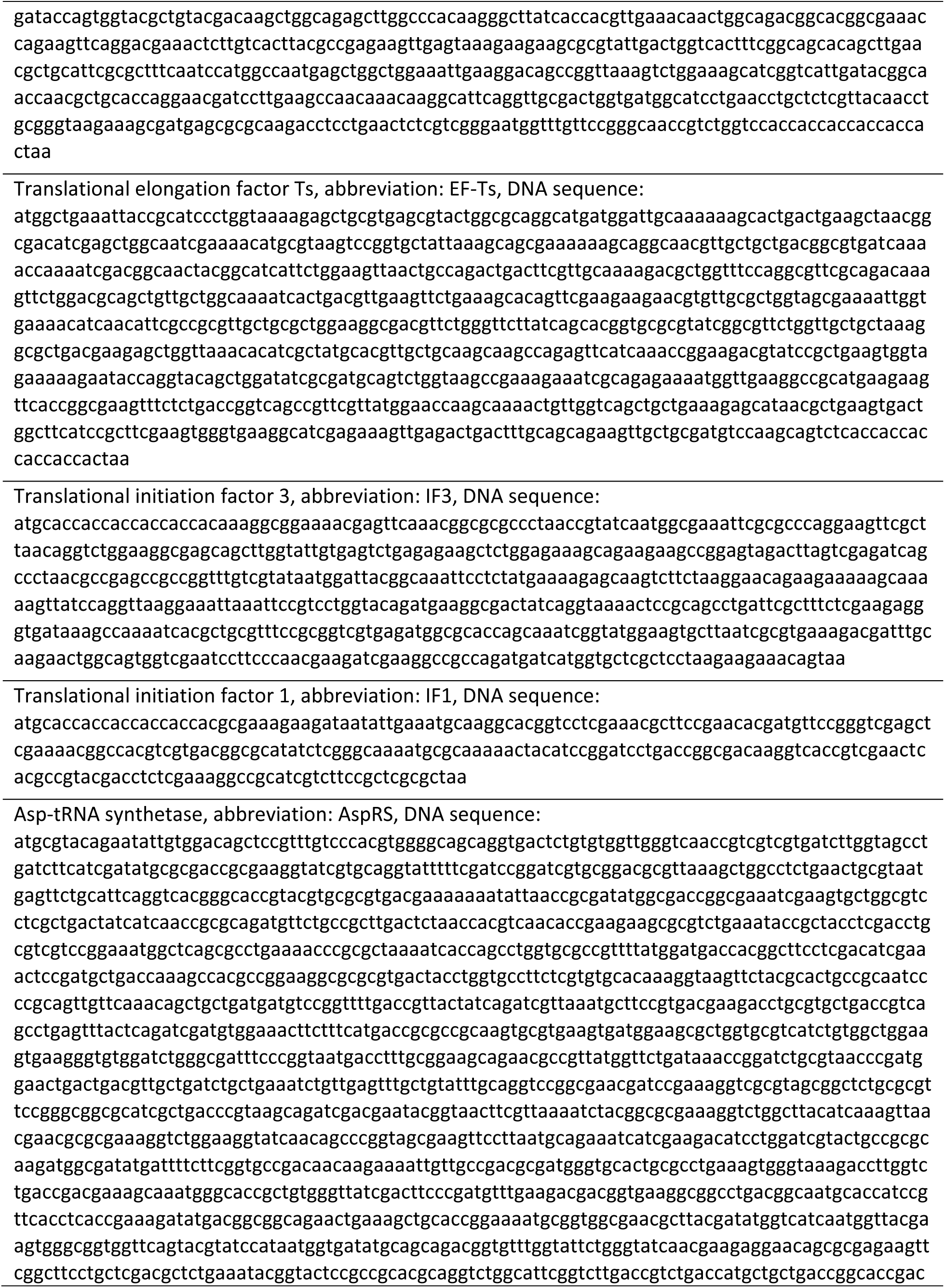
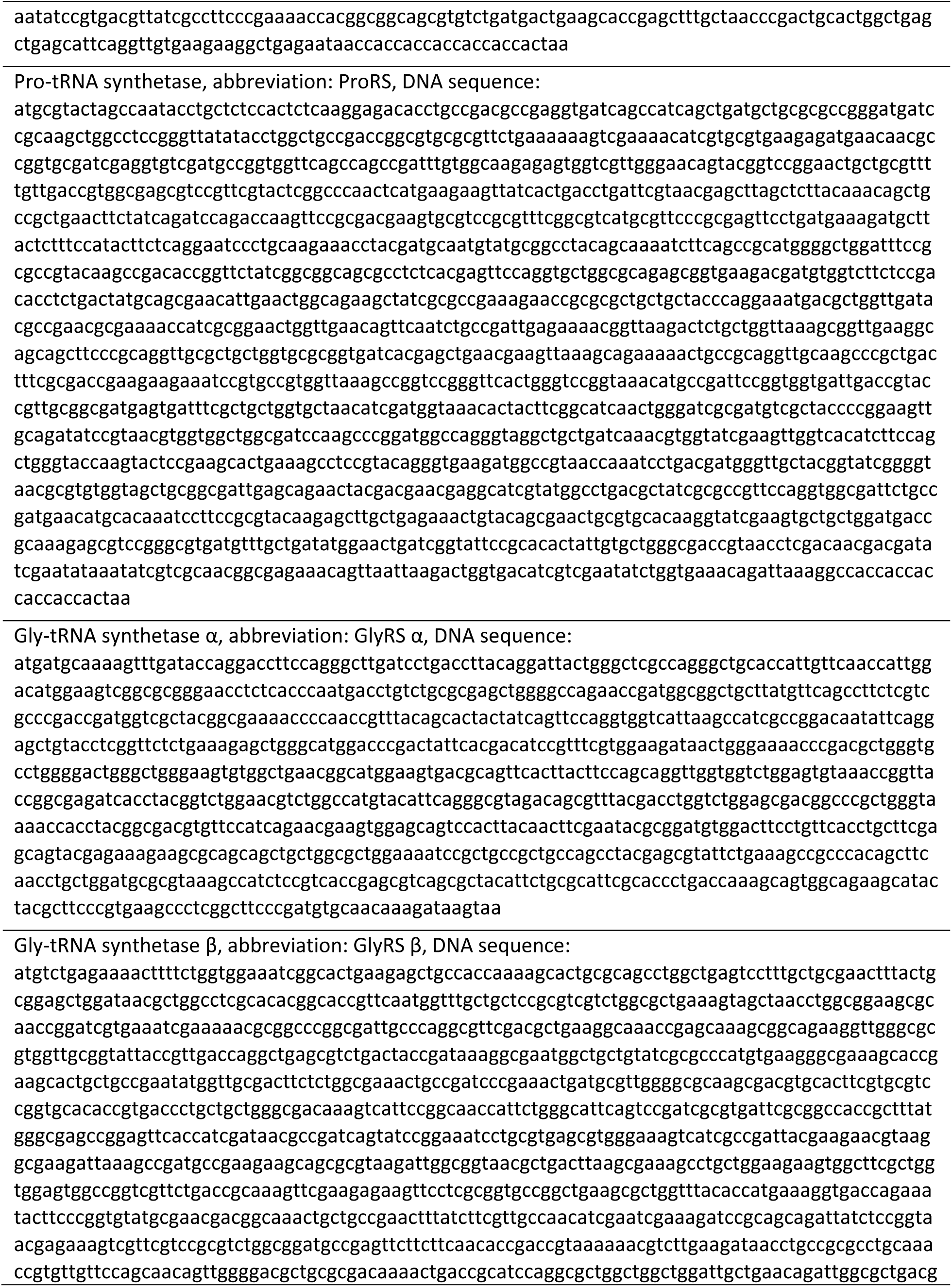
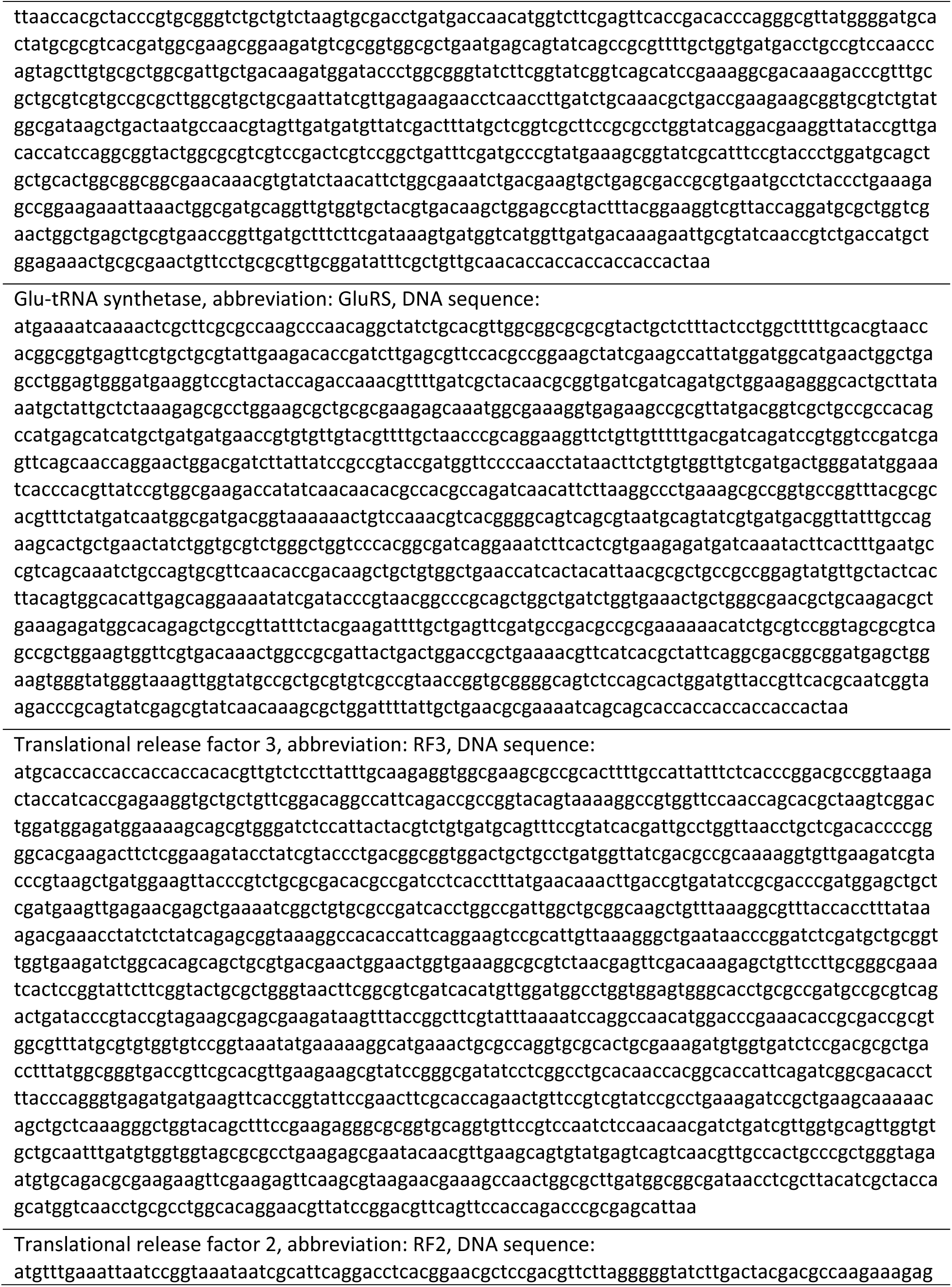
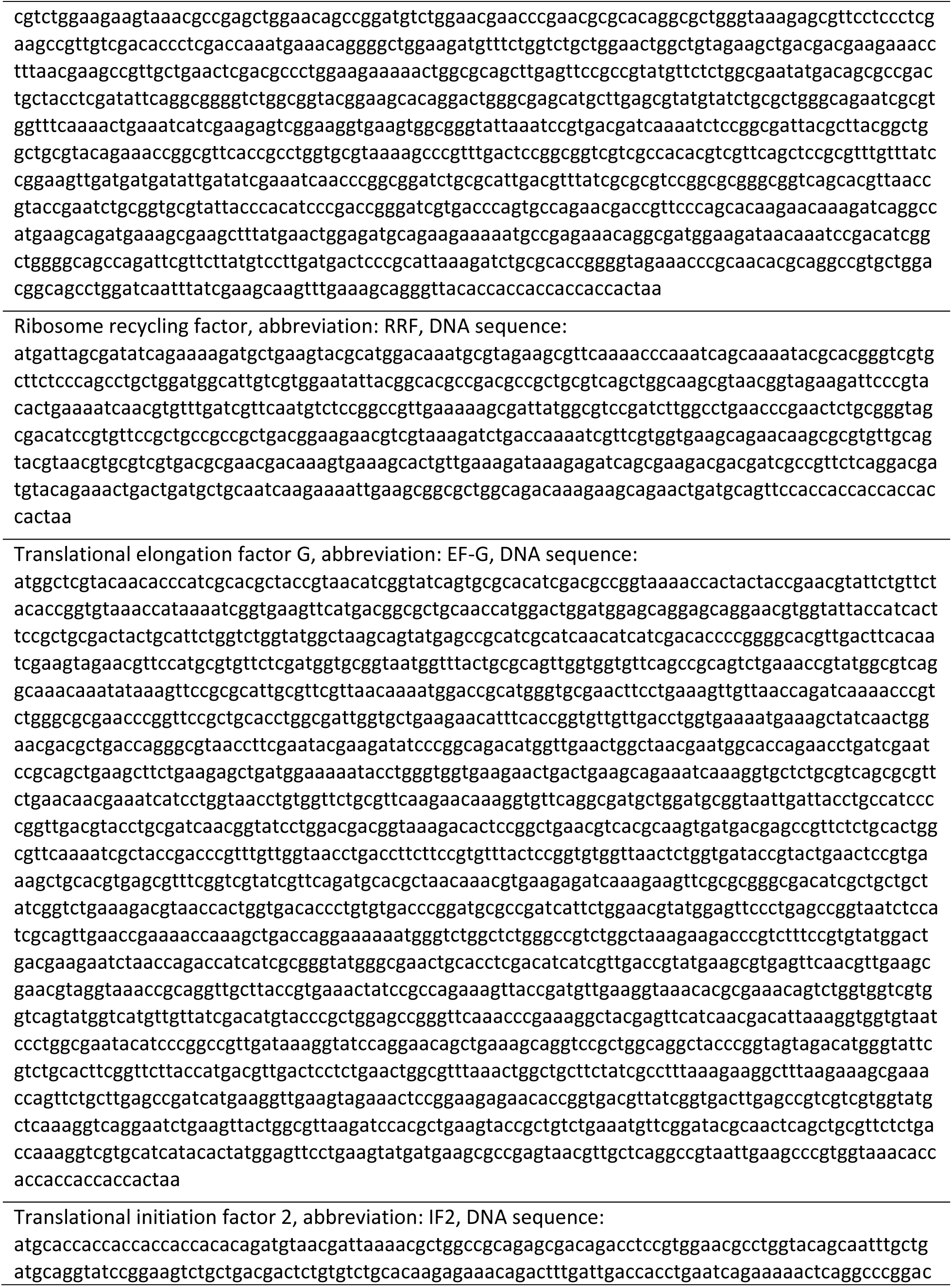
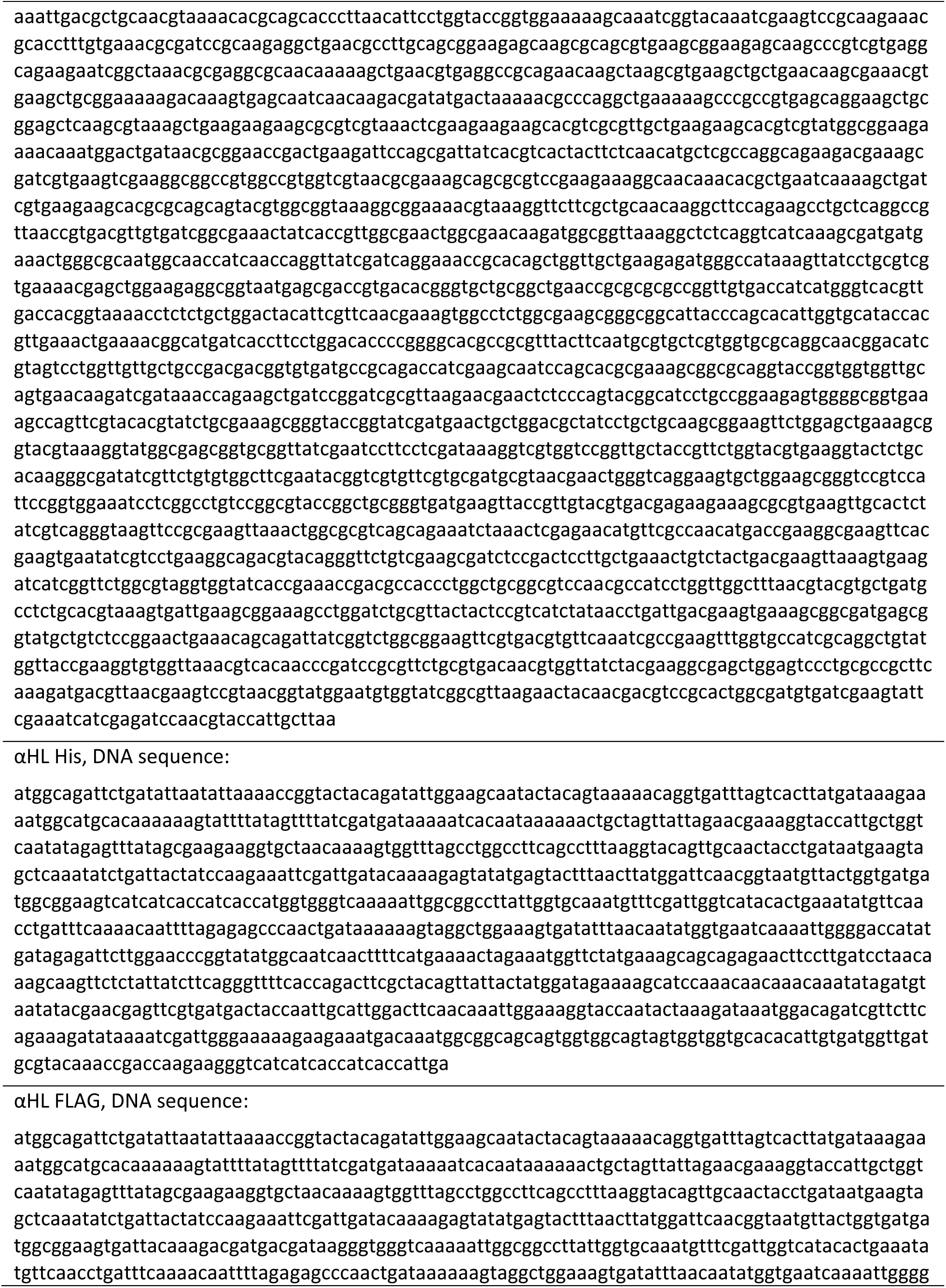
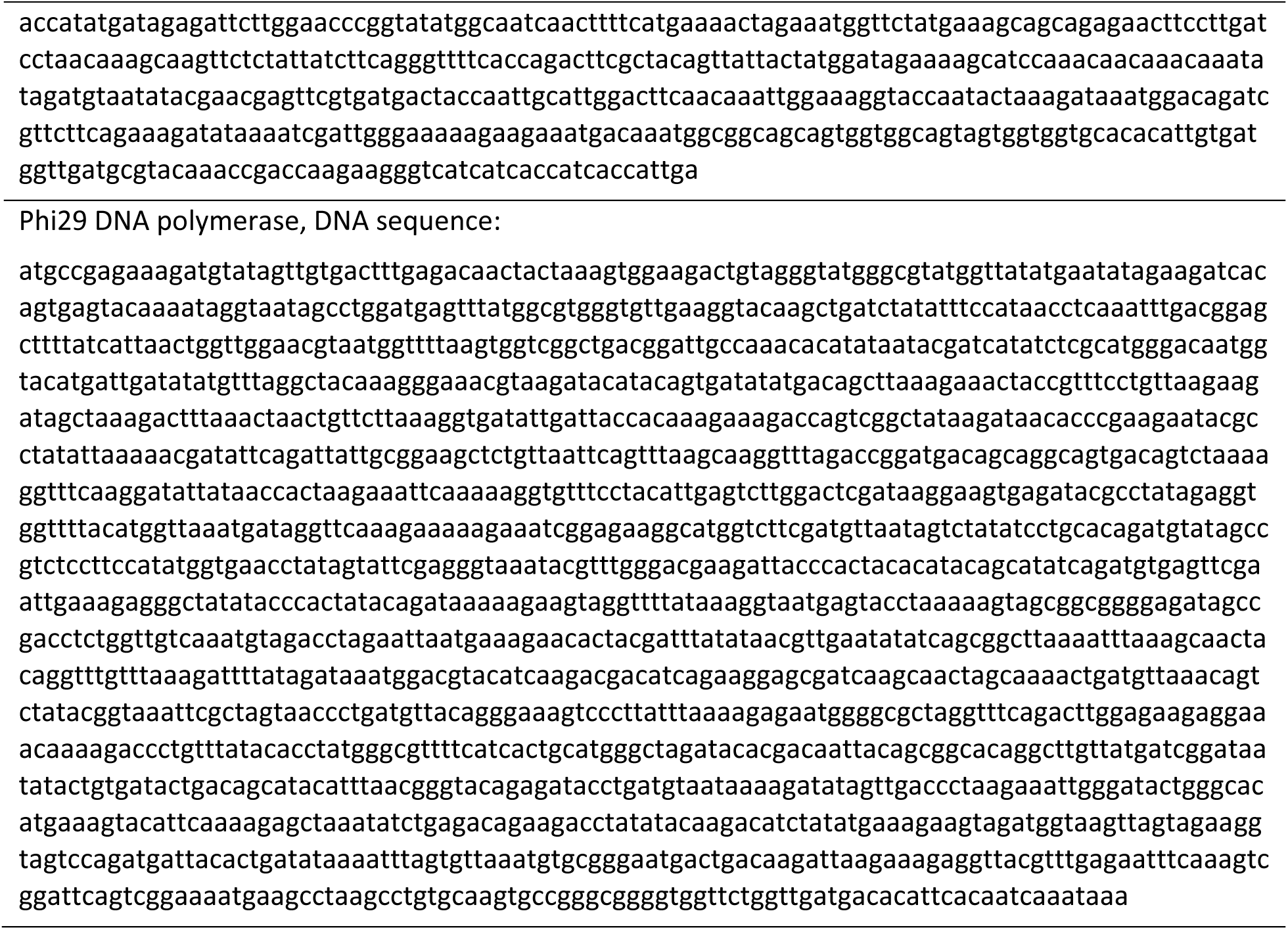
DNA sequences of synthetic cell genome translation factor genes. The sequences for pLD1, pLD2 and pLD3 are from ^65^.

**Table S8.** Conversion of plasmid concentration to plasmid copy number. The table is prepared for 1.5μm liposomes with 1mM lipids, assuming volume of 1.77 fL per liposome.

| plasmid<br>concentration | 1nM | 5nM | 10nM | 15nM |
| --- | --- | --- | --- | --- |
| average copy<br>number | 1.06 | 5.32 | 10.64 | 15.96 |

**Table S9.**
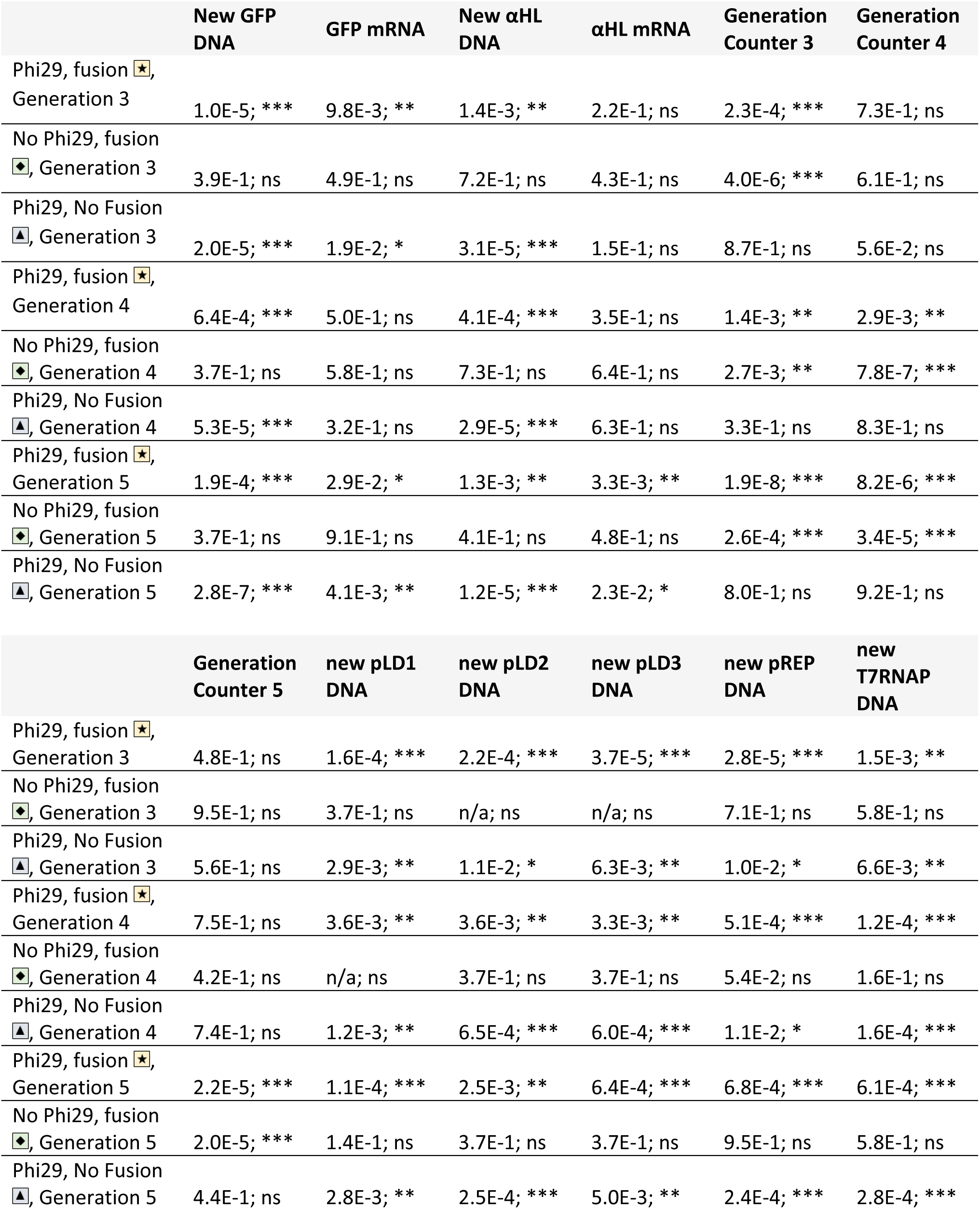
Statistical analysis of cell cycle data. Presented as plots on **Figure 3**, and individual data points on **Figure S34**. P values are calculated for all conditions compared to the negative control of no Phi29 genome replication and no feeding conditions (symbol 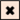 through the text). ns = P > 0.0; * = P ≤ 0.0, ** = P ≤ 0.01, *** = P ≤ 0.001; n/a when results are undetectable and P value was not calculated.

|  | New GFP DNA | GFP mRNA | New αHL DNA | αHL mRNA | Generation Counter 3 | Generation Counter 4 |
| --- | --- | --- | --- | --- | --- | --- |
| Phi29, fusion , Generation 3 | 1.0E-5; *** | 9.8E-3; ** | 1.4E-3; ** | 2.2E-1; ns | 2.3E-4; *** | 7.3E-1; ns |
| No Phi29, fusion , Generation 3 | 3.9E-1; ns | 4.9E-1; ns | 7.2E-1; ns | 4.3E-1; ns | 4.0E-6; *** | 6.1E-1; ns |
| Phi29, No Fusion , Generation 3 | 2.0E-5; *** | 1.9E-2; * | 3.1E-5; *** | 1.5E-1; ns | 8.7E-1; ns | 5.6E-2; ns |
| Phi29, fusion , Generation 4 | 6.4E-4; *** | 5.0E-1; ns | 4.1E-4; *** | 3.5E-1; ns | 1.4E-3; ** | 2.9E-3; ** |
| No Phi29, fusion , Generation 4 | 3.7E-1; ns | 5.8E-1; ns | 7.3E-1; ns | 6.4E-1; ns | 2.7E-3; ** | 7.8E-7; *** |
| Phi29, No Fusion , Generation 4 | 5.3E-5; *** | 3.2E-1; ns | 2.9E-5; *** | 6.3E-1; ns | 3.3E-1; ns | 8.3E-1; ns |
| Phi29, fusion , Generation 5 | 1.9E-4; *** | 2.9E-2; * | 1.3E-3; ** | 3.3E-3; ** | 1.9E-8; *** | 8.2E-6; *** |
| No Phi29, fusion , Generation 5 | 3.7E-1; ns | 9.1E-1; ns | 4.1E-1; ns | 4.8E-1; ns | 2.6E-4; *** | 3.4E-5; *** |
| Phi29, No Fusion , Generation 5 | 2.8E-7; *** | 4.1E-3; ** | 1.2E-5; *** | 2.3E-2; * | 8.0E-1; ns | 9.2E-1; ns |
|  | Generation Counter 5 | new pLD1 DNA | new pLD2 DNA | new pLD3 DNA | new pREP DNA | new T7RNAP DNA |
| Phi29, fusion , Generation 3 | 4.8E-1; ns | 1.6E-4; *** | 2.2E-4; *** | 3.7E-5; *** | 2.8E-5; *** | 1.5E-3; ** |
| No Phi29, fusion , Generation 3 | 9.5E-1; ns | 3.7E-1; ns | n/a; ns | n/a; ns | 7.1E-1; ns | 5.8E-1; ns |
| Phi29, No Fusion , Generation 3 | 5.6E-1; ns | 2.9E-3; ** | 1.1E-2; * | 6.3E-3; ** | 1.0E-2; * | 6.6E-3; ** |
| Phi29, fusion , Generation 4 | 7.5E-1; ns | 3.6E-3; ** | 3.6E-3; ** | 3.3E-3; ** | 5.1E-4; *** | 1.2E-4; *** |
| No Phi29, fusion , Generation 4 | 4.2E-1; ns | n/a; ns | 3.7E-1; ns | 3.7E-1; ns | 5.4E-2; ns | 1.6E-1; ns |
| Phi29, No Fusion , Generation 4 | 7.4E-1; ns | 1.2E-3; ** | 6.5E-4; *** | 6.0E-4; *** | 1.1E-2; * | 1.6E-4; *** |
| Phi29, fusion , Generation 5 | 2.2E-5; *** | 1.1E-4; *** | 2.5E-3; ** | 6.4E-4; *** | 6.8E-4; *** | 6.1E-4; *** |
| No Phi29, fusion , Generation 5 | 2.0E-5; *** | 1.4E-1; ns | 3.7E-1; ns | 3.7E-1; ns | 9.5E-1; ns | 5.8E-1; ns |
| Phi29, No Fusion , Generation 5 | 4.4E-1; ns | 2.8E-3; ** | 2.5E-4; *** | 5.0E-3; ** | 2.4E-4; *** | 2.8E-4; *** |

**Table S10.**
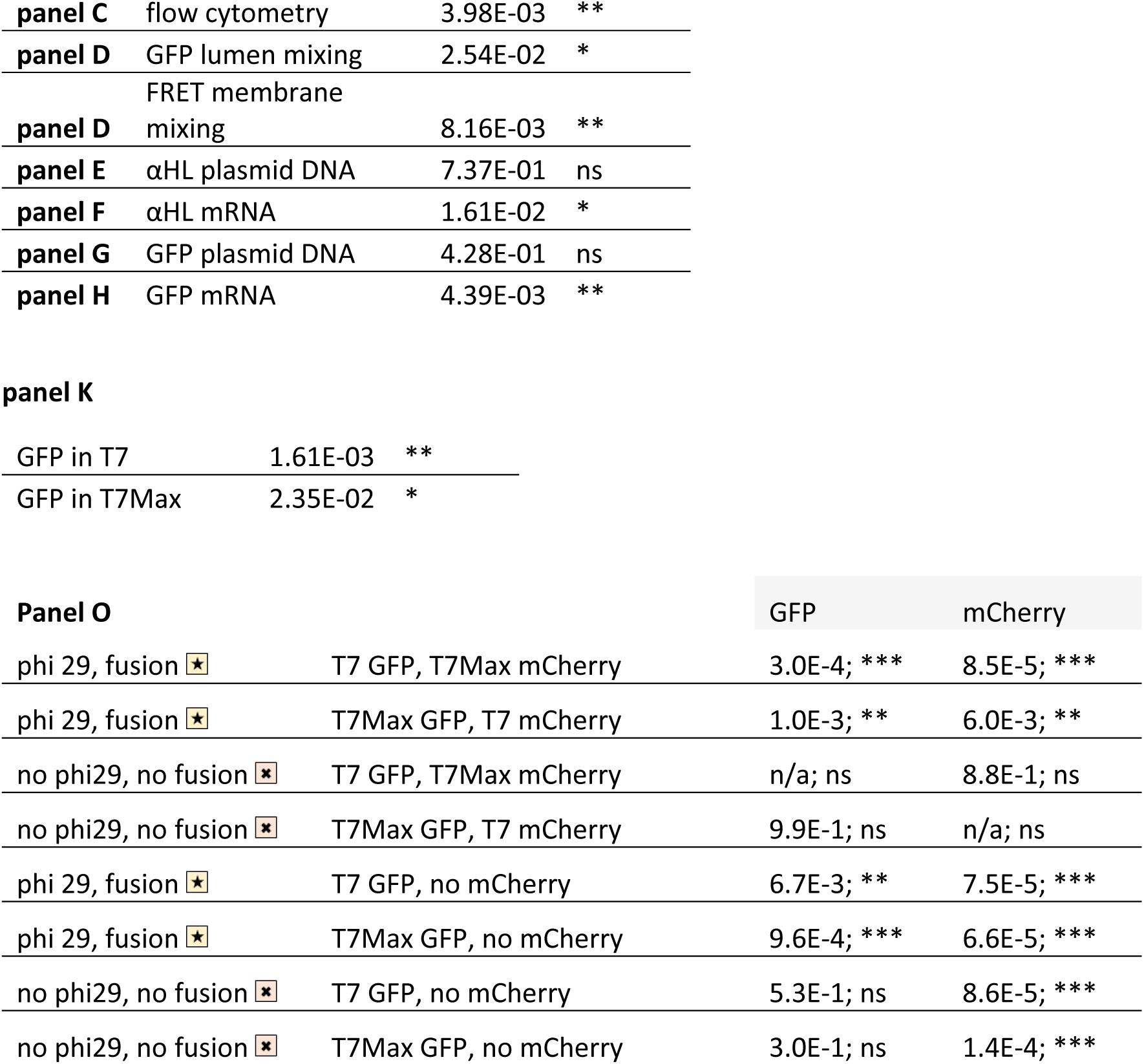
Statistical analysis of data for selection experiments. For data on panels C to H: Since those experiments measure competition between T7 and T7Max cells, the P values were calculated comparing sample set of T7αHL to T7MaxαHL (not compared to negative control). For data on panel K: the P values were calculated comparing samples with and without growth and division, directly comparing samples with GFP in T7 cells with and without cell cycle, and comparing samples with GFP in T7Max cells with and without cell cycle. For data on panel O: the P values calculated for GFP and for mCherry compared to the negative control: incubation without DNA replication and without feeding and division (symbol 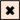) in the weaker promoter T7 cells. ns = P > 0.0; * = P ≤ 0.0, ** = P ≤ 0.01, *** = P ≤ 0.001, n/a when results are undetectable and P value was not calculated.

**Table S11.** Statistical analysis of data for two populations of synthetic cells mixed in each competition experiment, results shown on panels C – E of Figure 5. ns = P > 0.0; * = P ≤ 0.0, ** = P ≤ 0.01, *** = P ≤ 0.001; n/a when results are undetectable and P value was not calculated.

| feeder liposomes | synthetic cells | GFP new DNA, panel 5C | mCherry new DNA, panel 5D | pLD1 new DNA, panel 5E |
| --- | --- | --- | --- | --- |
| 1x | T7 GFP, T7Max mCherry | 2.8E-3; ** | n/a; ns | n/a; ns |
| 1x | T7Max GFP, T7 mCherry | n/a; ns | 5.5E-2; ns | 8.7E-1; ns |
| 0.5x | T7 GFP, T7Max mCherry | 8.6E-4; *** | 3.3E-1; ns | 8.3E-1; ns |
| 0.5x | T7Max GFP, T7 mCherry | 6.5E-2; ns | 5.7E-3; ** | 5.2E-1; ns |
| 0.25x | T7 GFP, T7Max mCherry | 1.5E-4; *** | 1.3E-2; * | 1.0E-2; * |
| 0.25x | T7Max GFP, T7 mCherry | 3.5E-3; ** | 2.0E-3; ** | 9.4E-3; ** |
| 0.1x | T7 GFP, T7Max mCherry | 7.0E-6; *** | 6.9E-3; ** | 3.4E-3; ** |
| 0.1x | T7Max GFP, T7 mCherry | 1.5E-4; *** | 9.0E-4; *** | 2.5E-3; ** |

**Table S12.** Statistical analysis of data for synthetic cells growth and division, data shown on Figure 6. ns = P > 0.0; * = P ≤ 0.0, ** = P ≤ 0.01, *** = P ≤ 0.001; n/a when results are undetectable and P value was not calculated.

|  |  |  |
| --- | --- | --- |
| washed off beads,<br>panel **6H** | T7 | T7Max |
| T7 cells present, T7Max cells<br>absent; division is expected | 1.5E-2; * | 4.8E-1; ns |
| T7 cells absent, T7Max cells<br>present; division is expected. | n/a; ns | 9.4E-3; ** |
| T7 cells present, T7Max cells<br>present; division is expected. | 4.4E-2; * | 2.3E-2; * |
| Attached to beads,<br>panel **6I** | T7 | T7Max |
| T7 cells present, T7Max cells<br>absent; division is expected | 7.2E-1; ns | n/a; ns |
| T7 cells absent, T7Max cells<br>present; division is expected. | n/a; ns | 7.1E-1; ns |
| T7 cells present, T7Max cells<br>present; division is expected. | 5.8E-1; ns | 1.1E-1; ns |

| sample | new αHL,<br>attached to<br>beads,<br>panel **6L** | total αHL,<br>attached to<br>beads,<br>panel **6M** | generation<br>counter,<br>attached to<br>beads,<br>panel **6K** | new αHL,<br>free off<br>beads,<br>panel **6O** | total αHL,<br>free off<br>beads,<br>panel **6P** | generation<br>counter,<br>free off<br>beads,<br>panel **6N** |
| --- | --- | --- | --- | --- | --- | --- |
| DNA replication; no<br>growth; division | 1.7E-2; * | 4.7E-2; * | n/a; ns | 6.9E-1; ns | 2.8E-1; ns | 3.5E-3; ** |
| DNA replication; growth;<br>no division | 8.4E-1; ns | 7.5E-1; ns | 2.2E-1; ns | n/a; ns | 7.0E-5; *** | n/a; ns |
| DNA replication; no<br>growth; no division | 9.8E-1; ns | 1.3E-2; * | 6.6E-2; ns | 2.5E-2; * | n/a; ns | 3.7E-3; ** |
| no DNA replication;<br>growth; division | n/a; ns | 4.7E-5; *** | 4.0E-1; ns | 2.6E-2; * | 1.6E-3; ** | 7.0E-1; ns |
| no DNA replication; no<br>growth; division | 1.2E-3; ** | 3.7E-4; *** | 6.5E-2; ns | 2.2E-2; * | 3.0E-2; * | n/a; ns |
| no DNA replication;<br>growth; no division | 7.7E-3; ** | 8.4E-4; *** | 7.4E-2; ns | 5.4E-3; ** | 8.0E-4; *** | n/a; ns |
| no DNA replication; no<br>growth; no division | 1.2E-3; ** | 1.3E-4; *** | 6.4E-2; ns | 2.5E-2; * | 8.6E-4; *** | 3.5E-3; ** |

